# Mutational signature catalogue *(MUSIC)* of DNA double-strand break repair

**DOI:** 10.1101/2025.04.29.650617

**Authors:** Marco Barazas, Robin van Schendel, Marcel Tijsterman

**Affiliations:** Human Genetics, Leiden University Medical Center, Leiden, The Netherlands; Institute of Biology Leiden, Leiden University, Leiden, The Netherlands

## Abstract

Genome alterations arise from inaccurate DNA repair, accumulating into distinct mutational signatures. Here, we investigate the role of every genomically encoded gene in double-strand break (DSB) repair by generating high-resolution outcome profiles following gene knockouts. Using a CRISPR/Cas9-based, massive-parallel bulk library approach, we construct a comprehensive, user-explorable mutational signature catalogue (MUSIC), mapping the full repertoire of DSB repair factors. Our analysis identifies and validates gene clusters – including nearly all known and several novel genes – linked to non-homologous end-joining, 53BP1 sub-pathways, homology-directed repair, and polymerase Theta (POLQ)-mediated end-joining. By focusing on pathway-specific repair outcomes, we uncover a previously unrecognized role for the WRN helicase in suppressing inverted templated insertions, a poorly understood POLQ-associated mutational signature also found in human disease alleles. Furthermore, in-depth analysis of MUSIC’s scar features reveals unexpected distinctions among genes within the same pathway, providing mechanistic insight and opening multiple new avenues for investigation into chromosomal break repair.

## Introduction

Cancer is fundamentally a genetic disease, with genomic instability standing as one of its defining hallmarks^1^. Next-generation sequencing (NGS) of tumours has provided comprehensive maps of genomic alterations^2–6^, revealing how mutations can serve as telltale signs of underlying cellular defects, a cell’s history of exposure to DNA-damaging agents, and potential cellular vulnerabilities. For example, breast tumours can be classified as homologous recombination (HR) proficient or deficient based on their mutational landscape, and even defects in BRCA1 or BRCA2 can be distinguished solely by their mutation profile^7^.

The abundance of deletions characterized by microhomology (MH) at their junctions is the strongest predictor of the HR-deficient (HRD) mutational signature^7,8^. This scar is also a hallmark of polymerase theta (POLQ)-mediated end-joining (TMEJ) of DNA double-strand breaks (DSBs)^9–11^. Indeed, the HRD signature depends on POLQ functionality^12^, and accordingly, HRD tumour cells rely on POLQ for their survival^13,14^. In TMEJ, POLQ utilizes complementary bases in 3’ single-stranded DNA (ssDNA) overhangs at DSB ends to establish a stable connection via DNA synthesis. As a result, the intervening nucleotides between the complementary MH stretches, as well as one of the two MH tracts, are lost. Detailed analysis of thousands of repair products across a wide range of species – including worms, flies, plants, fish, and mammals – have revealed another characteristic TMEJ outcome: templated insertions (TINS)^15^. These events involve a deletion accompanied by an insertion that is, at least in part, identical to sequences flanking the DSB. Even complex combinatorial insertions resembling a patchwork of nearby sequences have been observed. Such POLQ footprints are also found in *de novo* disease alleles responsible for human genetic syndromes^15^. The prevailing model for the aetiology of TINS involves iterative rounds of TMEJ: after initial DNA synthesis by POLQ, the repair reaction – due to unknown contributing factors – fails to complete. The extended DSB end then dissociates from its substrate, followed by a second repair attempt at a different MH^15,16^. Consequently, the DNA synthesized in the earlier repair attempt is incorporated as an insertion within the final TINS outcome.

To employ these genomic scars as biomarkers for mutagenic DSB repair, we previously developed software that establishes high resolution repair outcome profiles for CRISPR/Cas9-induced DSBs via targeted NGS^17^. Highly specific profile redistributions were observed upon knockout of core repair factors^18^, as was also recently demonstrated using a library-based approach for a set of over 400 genes linked to DNA repair^19^. This work showed that altered repair profiles – and the direction of these alterations – can help disentangle the determinants of DSB repair, the underlying mechanisms, the proteins involved, and how they collectively contribute to the spectrum of deleterious mutations arising at chromosomal breaks.

Here, we established a methodology for the simultaneous detection of CRISPR/Cas9-induced DSB repair outcomes mediated by canonical non-homologous end-joining (c-NHEJ), homology-directed repair (HDR) and TMEJ pathways. We systematically profiled repair outcomes across genome-wide gene knockouts in bulk and validated the top outliers through the construction and high-resolution testing of a candidate library, enabling their comprehensive exploration. We identified key factors driving each repair pathway and uncovered distinct patterns to which specific genes acting within one DSB repair pathway differentially contribute. This high-resolution mutational profiling provides deeper insight into the biochemical steps underlying DSB repair mechanisms. To demonstrate that even subtle deviations in repair outcome profiles can yield novel mechanistic insight, we investigated WRN helicase-deficient cells, which displayed elevated levels of a specific class of TINS: those in which the inserted sequence is inverted relative to its template in the DSB flank. Based on their sequence composition, we propose a model in which inverted TINS (iTINS) result from interaction and TMEJ activity between broken sister chromatids, a process suppressed by WRN. Finally, our data were compiled into a user-explorable mutational signature catalogue (MUSIC), offering a versatile, internally controlled, and high-resolution framework for analysing DSB repair at nucleotide resolution.

## Results

### A mutational signature catalogue (MUSIC) of DSB repair

We set out to develop a massive-parallel CRISPR/Cas9-based approach to construct DSB repair profiles in mouse embryonic stem (mES) cells following the knockout of each gene in the genome. DSBs are introduced into randomly integrated lentiviral vector molecules, which also encode single-guide RNAs (sgRNAs) that generate functional gene knockouts (Fig. 1A). By targeting a unique region near the sgRNA, paired-end sequencing reads can be obtained that couple the repair outcome to the genetic background of the cell in which the repair occurred. This allows for the compilation of mutational profiles reflecting the relative abundance of repair outcomes across different knockout conditions, provided that: (i) knockouts are induced at high frequencies, and (ii) sequencing is performed on sufficiently complex cell populations and at sufficient depth to ensure independent events within each genetic background. To test the experimental design, we transduced Cas9-expressing IB10 mES cells with lentiviruses targeting the c-NHEJ core factor *Lig4* and the TMEJ factor *Polq*; non-targeting sgRNAs (sgNT-1 and sgNT-2) served as controls^20^. Targeting efficiencies exceeding 90% were achieved (Supplementary Fig. S1A-B). These cell lines were subsequently transduced with one of three library backbone-targeting sgRNAs – hereafter referred to as sg*Library-1*, sg*Library-2*, and sg*Library-3.* Three days later, DNA was isolated, pooled, and subjected to NGS of the targeted region. Gene-specific mutational profiles were obtained by demultiplexing the pooled NGS data based on the sgRNA, which served as a barcode, followed by SIQ analysis^17^. This software deduces a repair outcome from each sequencing read based on the predicted CRISPR/Cas9 cut site and annotates numerous outcome characteristics crucial for future data analyses. Across all three target sites, mutational profiles derived from *Lig4*- and *Polq*-targeted populations each showed a characteristic outcome redistribution: altered frequencies of 1bp insertions and MH-driven deletions, respectively (Supplementary Fig. S1C-H). This redistribution pattern remains robust even after computationally down-sampling to a significantly reduced read depth (Supplementary Fig. S1I-J). Additionally, we verified that no data contamination occurred due to hybrid reads in which sgRNAs are randomly coupled to repair outcomes during library preparation, i.e. due to PCR jumping (Supplementary Fig. S1K). Collectively, these findings support the feasibility of a bulk whole-genome approach for generating gene-specific mutational signatures of DSB repair.

**Figure 1:**
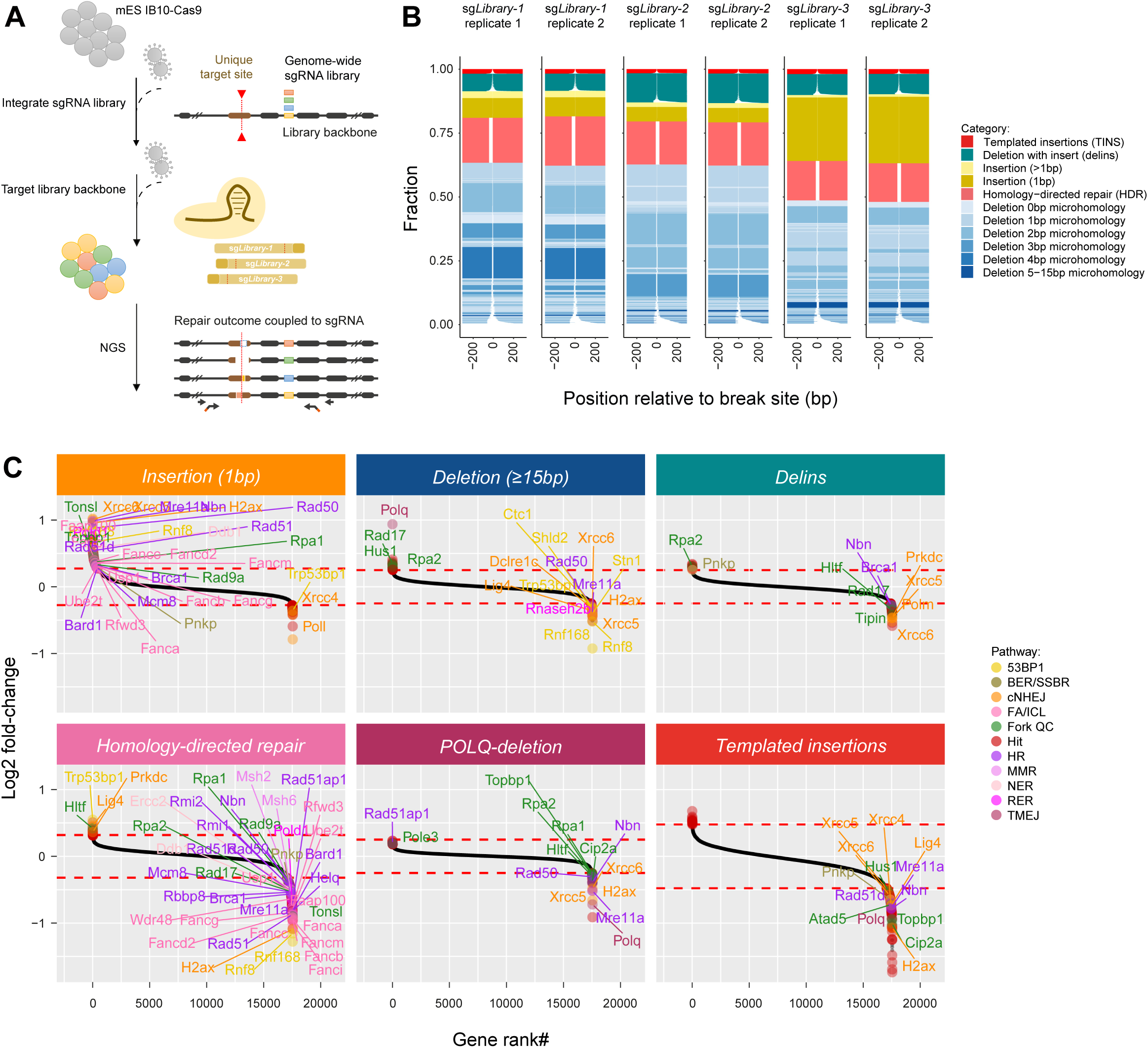
Mutational Signature Catalogue classifies known and novel DSB repair genes. ***(A)*** Schematic overview of the MUSIC workflow. Library sgRNAs both induce gene knockouts and serve as unique barcodes. Amplicons encompassing the sgRNA and DSB target site are subjected to massive parallel sequencing to generate repair profiles. ***(B)*** Deconvolution of NGS data using SIQ software produces tornado plots that display repair outcomes per target site and replicate. Outcomes are normalized to the total mutagenic reads and visualized horizontally, with colours indicating outcome categories. ***(C)*** SIQ data are demultiplexed into gene-specific profiles and outcomes are aggregated into defined mutational types. Genes are ranked by log2 fold-change (enrichment vs. depletion) and visualized in waterfall plots. A dashed horizontal line marks the 2.5x SD threshold; genes previously associated with DSB repair pathways are labelled and colour-coded.

The mouse genome-wide Yusa-v2 sgRNA-library^20^, which comprises ∼ 90,000 constructs (with 5 sgRNAs per gene), was used to perform a screen across three DSB target sites in duplicate (Fig. 1A-B). To manage, explore, and visualize the ∼500,000 repair profiles – representing ∼18,000 gene knockouts – we further developed the SIQ software^17^. The data were filtered and aggregated (see methods for full details) into overall gene-scores per mutational type – outcomes which likely are products of comparable biochemistry. This operation reveals the robust identification of core factors involved in various DSB repair pathways (Fig. 1C): (i) c-NHEJ factors XRCC4 and POLL facilitate 1-bp insertions, a typical outcome scar generated by CRISPR/Cas9 (ii) the 53BP1-pathway complexes^21–23^ SHLD^24–27^ and CST^28,29^ provokes a distinct class of c-NHEJ deletions characterized by their length (≥15bp). (iii) The Fanconi Anaemia (FA) pathway promotes homology-directed repair (HDR) outcomes. (iv) POLQ drives deletions characterized by MH (Supplementary Fig. S1F-H) and TINS.

For validation, higher resolution testing, and gene discovery, we constructed a focused candidate library comprising 2,400 sgRNAs targeting the top outlier genes (selecting the 3 most active sgRNAs per gene), 170 non-targeting control sgRNAs, and DDR genes identified from literature^19^. DSBs were introduced using sg*Library-1*, which generates a variety of c-NHEJ and TMEJ outcomes, achieving an average of ∼ 100,000 repair outcomes per sgRNA per replicate (n=3). The 40-fold increase in resolution enables many repair events to be analysed on both a per-barcode and per-outcome basis. For each of the 150 most abundant outcomes, sgRNAs corresponding to the top 100 deviating barcodes were selected and plotted in an unsupervised, hierarchically clustered heatmap (Fig. 2). Supporting the quality of the dataset, different sgRNAs targeting the same gene clustered closely together, as did the outcomes from the three replicates. A uniform manifold approximation and projection (UMAP) provides a concise summary and visualization of the data, clustering sgRNAs with similar outcome redistribution patterns (Fig. 3 and Supplementary Fig. S3). This UMAP reveals the major DSB repair pathways and nearly all their core factors. Furthermore, known complex members - such as SHLD, CST, XRCC4-LIG4, 9-1-1, CIP2A-TOPBP1, XRCC5-XRCC6, MRN, RNF8-RNF168, BRCA1-BARD1 and the FA complexes – exhibited close relative positions. Notably, we identified an unexpected gene cluster comprising METTL3, METTL14, WTAP, and ZC3H13, all of which regulate N6-methyladenosine (m6A) RNA methylation^30,31^, implicating these genes in DSB repair.

**Figure 2:**
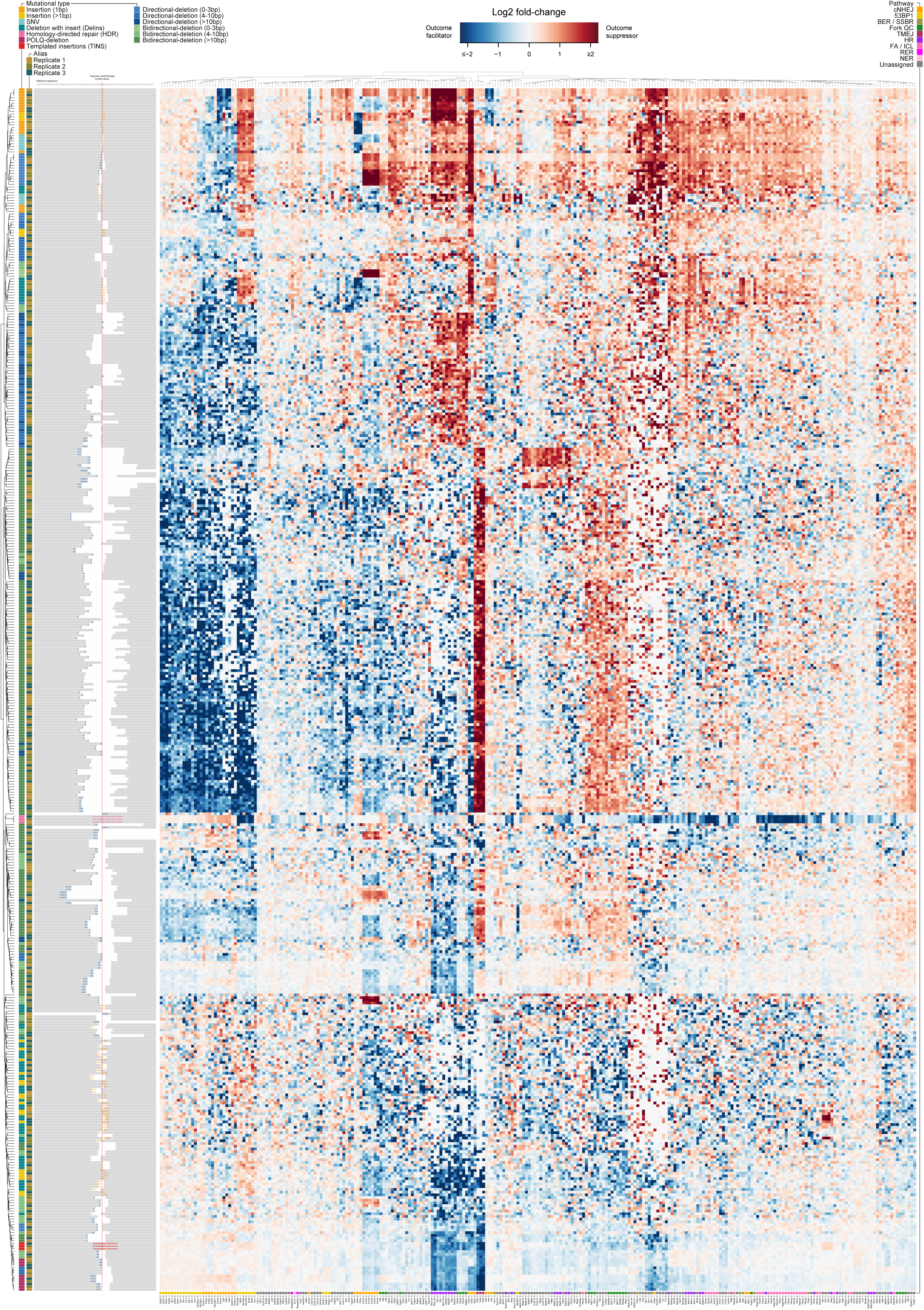
Heatmap representation of a focused MUSIC. For each of the 150 most abundant outcomes (y-axis), the top-100 sgRNA outliers were selected. The mutation redistribution profiles of these sgRNAs (x-axis) are presented in a heatmap, with relative positions determined by unsupervised hierarchical clustering. Outcomes and genes are also colour coded according to mutational type and DNA repair pathway.

**Figure 3:**
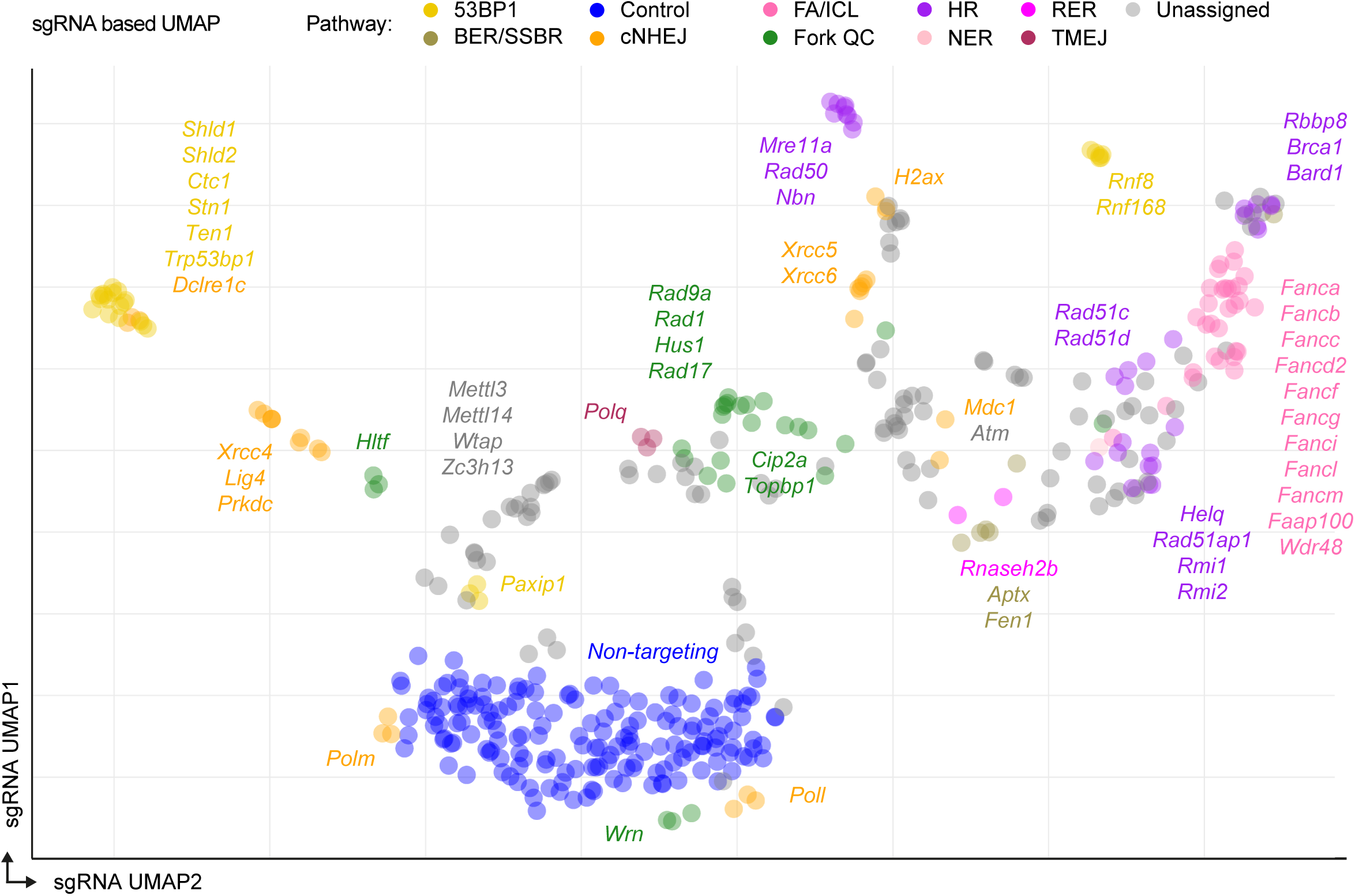
UMAP representation of MUSIC data. UMAP projection of the top outlier sgRNAs/genes. Each dot represents an individual sgRNA, positioned based on the similarity of its repair outcome distribution to that of other outlier or non-targeting sgRNAs. Genes are colour-coded by the DSB repair pathways in which they are involved.

A complimentary approach to summarise and extract mechanistic insight from MUSIC is to generate a UMAP that clusters repair outcomes based on shared genetic dependencies (Fig. 4A). Outcomes are colour-coded according to: (i) their mutational types, i.e. insertion, deletion – further annotated based on whether DNA loss occurs at one or both DSB ends (directional (D) versus bi-directional (B) deletions) –, deletions with insertion (delins), TINS, SNV and HDR (Fig. 4A), (ii) specific characteristics, such as deletion size and usage of microhomology (Fig. 4B-C), or (iii) the extent and direction of deviation for any given gene knock out (Fig. 4D). Highlighting the top 10 gene knockouts that induce the strongest profile changes illustrates how proteins that are representative of specific DSB repair activity, influence outcome distributions, and how these visualizations can be used to compare genes (Fig. 4D). For example, RNF8 and RNF168 knockouts produce nearly identical profiles, resembling but not fully matching that of 53BP1. Similarly, knockouts of MRN complex members display a reduction in specific outcomes also seen in POLQ knockouts, whilst distinct redistribution patterns are observed.

**Figure 4:**
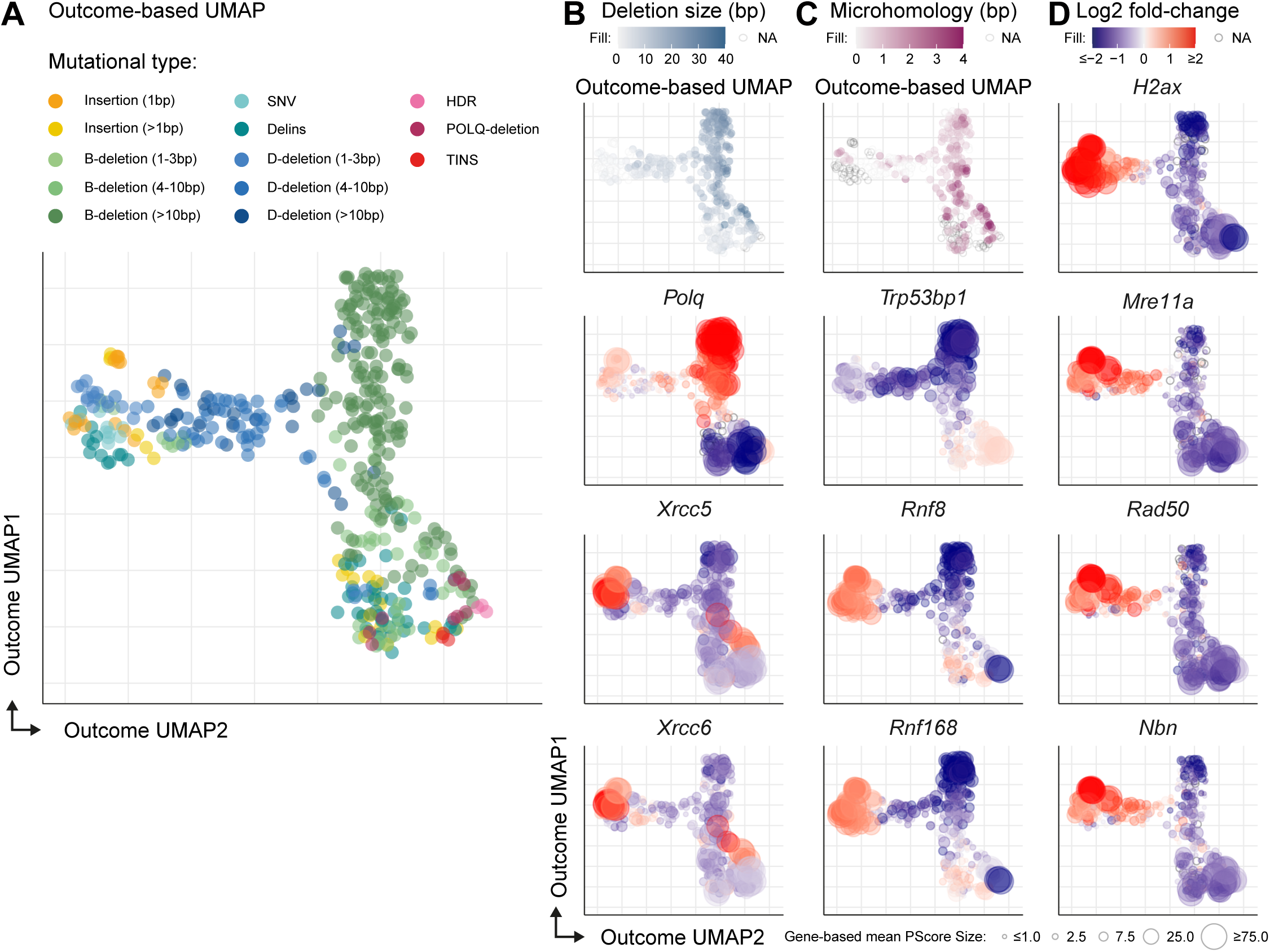
UMAP-based separation of repair outcomes according to distinct genetic dependencies. ***(A)*** UMAP projection of the top 100 outlier sgRNAs (consistent across replicates for at least one of the 150 most frequent repair outcomes), with each outcome plotted per replicate. Dots are colour coded by mutational type. ***(B)*** Same data as in (A), with colour intensity indicating deletion size (in bp). Outcomes without deletions are shown as open circles. ***(C)*** Same data as in (A), with colour intensity reflecting the length of microhomology (in bp). Only deletions are coloured; other outcomes appear as open circles. ***(D)*** The 10 genes with the strongest overall redistribution of repair outcomes. Dots are colour-coded by log2 fold-change relative to non-targeting sgRNAs and sized by Pscore. Outcomes absent in all sgRNAs targeting the indicated gene are represented as open circles.

### Three distinct aetiologies for insertional scars

Within the outcome UMAP we noticed clearly divergent positions for subsets of insertions and delins, suggesting the involvement of distinct repair mechanisms. One outcome cluster consists of small (1-3bp) insertions, deletions and delins and is driven by c-NHEJ factors (Fig. 5A). While c-NHEJ is generally accurate when acting on blunt DSB ends, it can produce small deletions or insertions due to additional processing before ligation or when acting on DSBs with protruding ends. Indeed, we find that a 1bp deletion event – consistent with processing a 1bp 5’ staggered DSB occasionally generated by Cas9 – requires the ARTEMIS (*Dclre1c*) nuclease in conjunction with the LIG4-XRCC4 DNA ligase (Fig. 5B). Conversely, the insertion of a single adenine, likely resulting from fill-in of the 1bp staggered DSB, depends on POLL (Fig. 5B-C), aligning with previous findings^19,32^. Interestingly, POLM, the other c-NHEJ polymerase^33^, contributes specifically to +1bp insertions of guanine - a bona fide c-NHEJ outcome requiring LIG4-XRCC4 - and to certain delins (Fig. 5B,D). These scars also require DNA-PKcs and ARTEMIS and are more prevalent in cells lacking POLL, which thus acts to supress these mutations. Unexpectedly, we also find that APTX, along with the ribonucleotide excision repair proteins RNASEH2B and FEN1, act as suppressors (Fig. 5A-B)^34^, even though these genes are not required for POLL-mediated insertional mutagenesis. It is tempting to speculate that these outcomes stem from the dNTP misincorporation properties previously described for POLM^35–37^. The POLL- and POLM-independent insertion events are brought about by POLQ and do not represent c-NHEJ action (Fig. 5B). The MUSIC thus highlights three distinct aetiologies for the generation of small (1-3bp) insertions, each with its own polymerase dependency.

**Figure 5:**
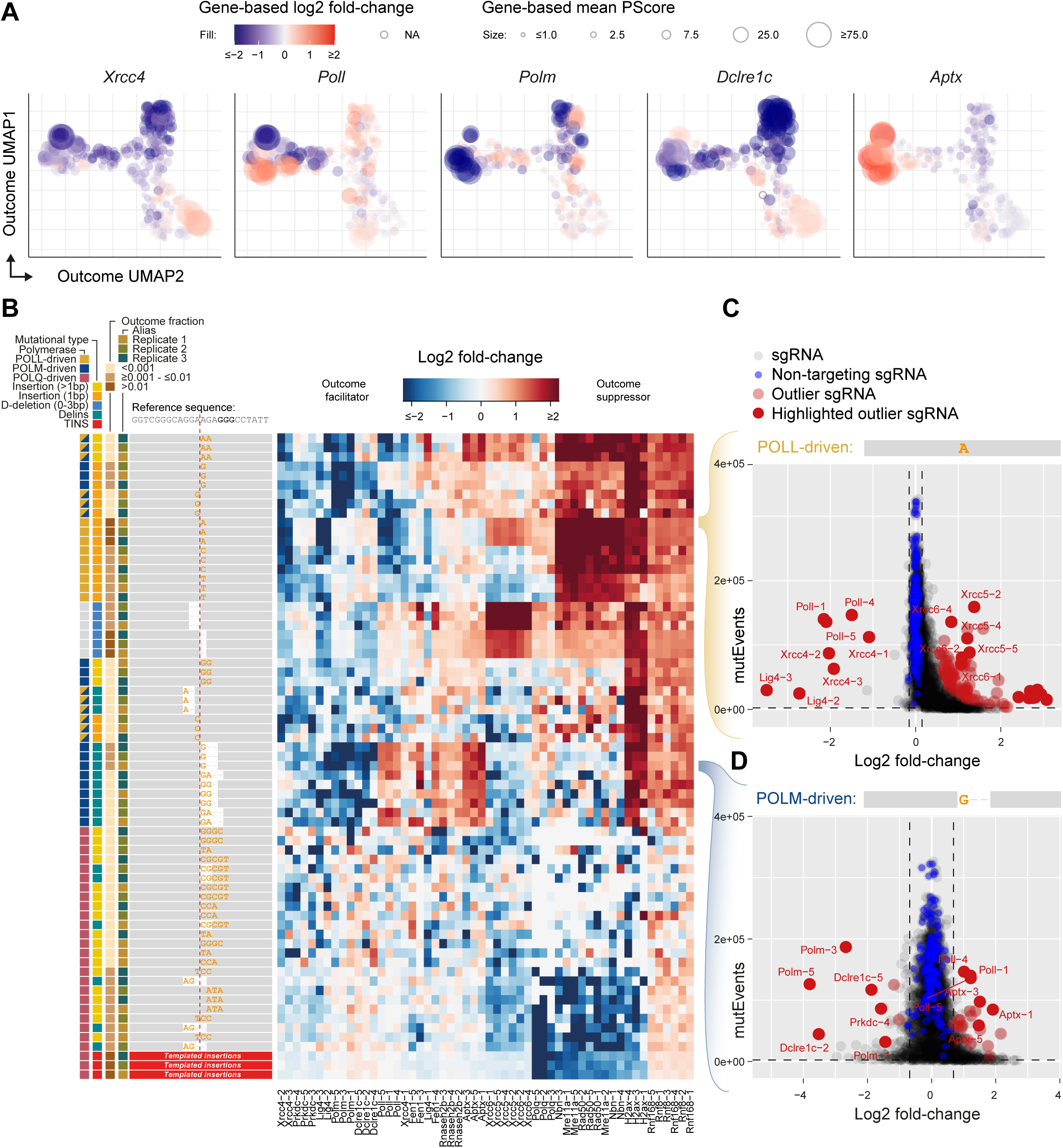
MUSIC identifies three distinct aetiologies for small (1-3bp) insertions. ***(A)*** UMAP projections (as in Fig. 4) of outcomes for the indicated genes. Dots are colour-coded by log2 fold-change relative to non-targeting sgRNAs and scaled according to P-Score. Outcomes absent in all sgRNAs targeting the indicated gene are represented as open circles. ***(B)*** Unsupervised hierarchical clustering of selected outcomes and sgRNAs, with each replicate shown separately. The heatmap displays log2 fold-change relative to non-targeting sgRNAs. Left panel shows schematic representations of outcomes, with coloured squares indicating the inferred polymerase, outcome category, frequency (as a fraction) and replicate. ***(C-D)*** Volcano plots for the indicated outcome, showing log2 fold-change (x-axis) versus total number of mutational events (y-axis) in replicate 1. Vertical dashed lines mark 3* SD from the non-targeting sgRNAs (blue), and the horizontal dashed line indicates a minimum of 2,000 mutational events per sgRNAs. sgRNAs consistently ranked in the top-100 most deviating (by Manhattan distance across all replicates) are labelled as ‘outliers’ (red); selected sgRNAs are further highlighted and annotated.

### The Fanconi Anaemia complex drives homology directed DSB repair

Many of the other outcomes follow from DSB end resection by members of the MRN complex (Fig. 4D). We unexpectedly discovered that in one specific, prominent outcome, the vector used to express Cas9 served as a template for HDR of DSBs at the target locus. The targeting vector and targeted vector are sufficiently different for library-specific PCR amplification to monitor repair outcomes yet have stretches of homologous sequence apparently sufficient to support HDR (Fig. 6A). We observed a reduction in HDR for the Fanconi Anaemia (FA) core complex members FANCA, FANCB, FANCC, FANCF, FANCG, FANCM, FAAP100, the FANCID2 complex and modifying factors FANCL, UBE2T, WDR48, USP1, as well as the downstream and/or associated repair factors BRCA1, BARD1, HELQ, RAD51AP1, RAD51C, RAD51D (Fig. 6B-C). A small number of novel factors, such as DUT, cluster tightly together with this set, providing the motivation for follow-up studies. Interestingly, c-NHEJ outcomes are specifically elevated in FA-targeted cells arguing against DSBs being rerouted to TMEJ (Fig. 6D). A potential upstream role of FA factors in HDR is in line with single-stranded template repair (SSTR) of CRISPR-induced DSBs, where the FA pathway directs repair away from c-NHEJ towards SSTR^38^.

**Figure 6:**
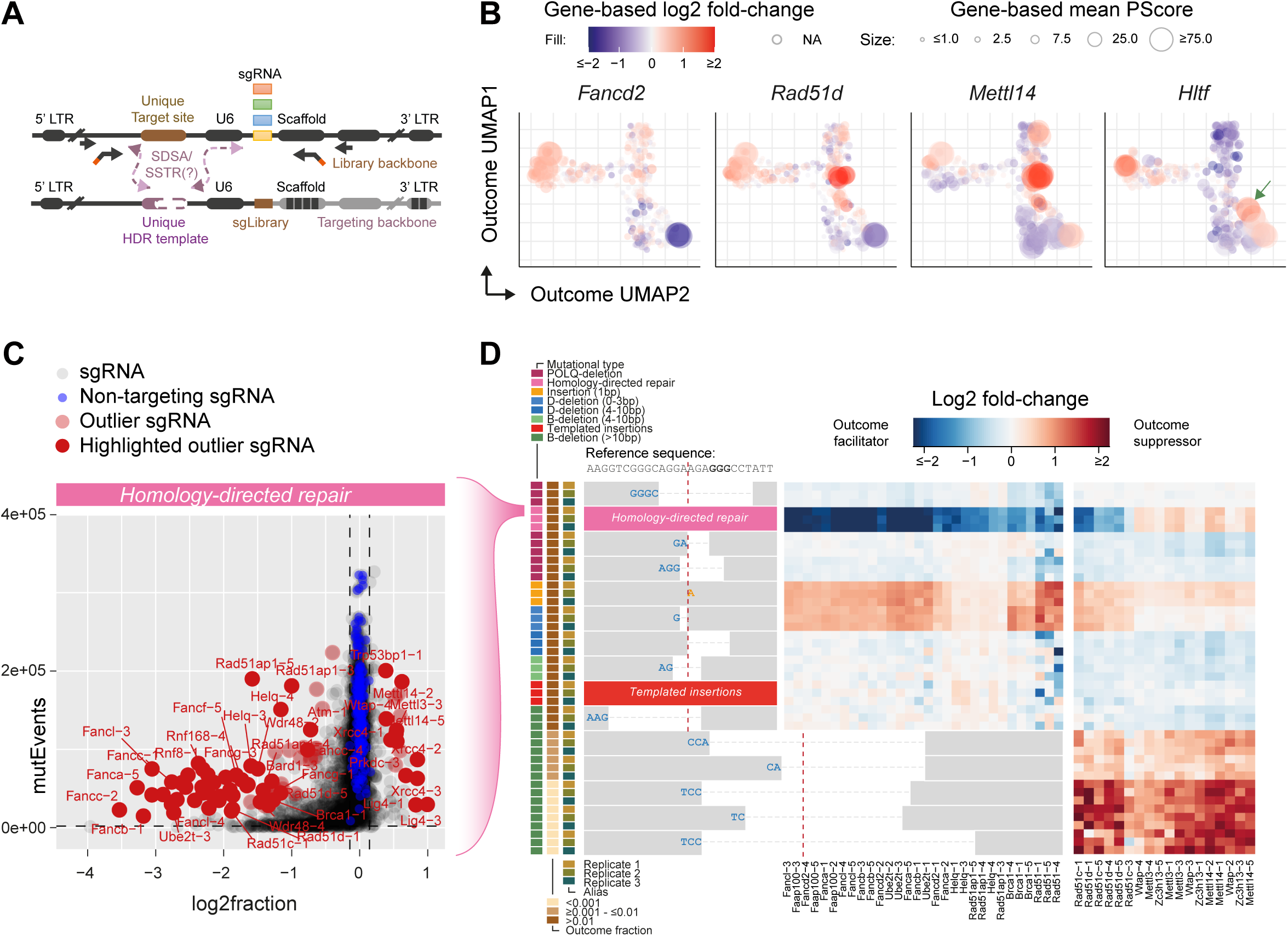
MUSIC of homology-directed repair implicates the Fanconi Anaemia pathway. ***(A)*** Schematic representation of the homology-directed repair reaction within MUSIC. ***(B)*** Outcome UMAPs for the indicated genes (as in Fig. 4). The green arrow points to a specific deletion having one of its junctions immediately upstream of the Cas9-binding site. ***(C)*** Volcano plot for the HDR outcome, showing log2 fold-change (x-axis) versus total number of mutational events (y-axis) in replicate 1. Vertical dashed lines mark 3* SD from the non-targeting sgRNAs (blue), and the horizontal dashed line indicates a minimum of 2,000 events per sgRNAs. sgRNAs consistently ranked in the top-100 most deviating (by Manhattan distance across all replicates) are labelled (red); selected sgRNAs are further highlighted and annotated. ***(D)*** Heatmap representation of the top-10 most frequent outcomes for a selected set of sgRNAs, clustered based on outcome similarity, as well as five infrequent outcomes (bottom) that collectively link METTL3-METTL14-WTAP-ZC3H13 to specific outcome redistributions.

We further noticed that HDR was suppressed by c-NHEJ factors LIG4, XRCC4 and DNA-PK (*Prkdc*) as well as by the TMEJ core factor POLQ (Fig. 2 and Fig. 6C), augmenting recent work^39–41^ that interference with these pathways can be utilized to synergistically enhance gene-targeting efficiency. HLTF also acted as a suppressor of HDR, which may be explained by its recently described activity in clearance of Cas9 from DNA upon DSB formation^42^. Further support for this notion is an increased frequency of a very specific deletion outcome with the junction located just upstream of the Cas9 binding site (Fig. 2 and Fig. 6B). Finally, we identified METTL3, METTL14, WTAP and ZC3H13 as suppressors of the HDR reaction and of very specific deletions (Fig. 6B,D).

### Genes involved in TMEJ

Besides HDR, many outcomes following MRN-dependent end resection can be classified as products that result either from 53BP1 pathway activity (Fig. 4D) – including its downstream complexes SHLD and CST (Fig. 2) – or from TMEJ pathway activity (Fig. 4D). This opposing behaviour has been observed previously^19^ and may indicate that the 53BP1-pathway repairs DSBs that could not be joined through POLQ action. Products of the TMEJ pathway includes deletions directed by microhomology, templated insertions, and certain delins and insertions (Fig. 4); POLQ is the most outstanding driver of these outcomes (Fig. 7A-B). Interestingly, the outcome UMAP reveals POLQ involvement in deletions that we *a priori* did not anticipate to result from TMEJ. For example, the 7^th^ most abundant repair outcome is POLQ-dependent, yet lacks MH at the junction (Fig. 7A). A potential explanation may lie in MH being concealed: this repair product has a 3bp MH 1bp away from the junction, fitting with the notion that recombinant POLQ can extend from unpaired termini *in vitro*^43^. These types of outcomes underscore the necessity of genetically examining mutational profiles at nucleotide resolution to fully understand the complex nature of POLQ activity *in vivo*.

**Figure 7:**
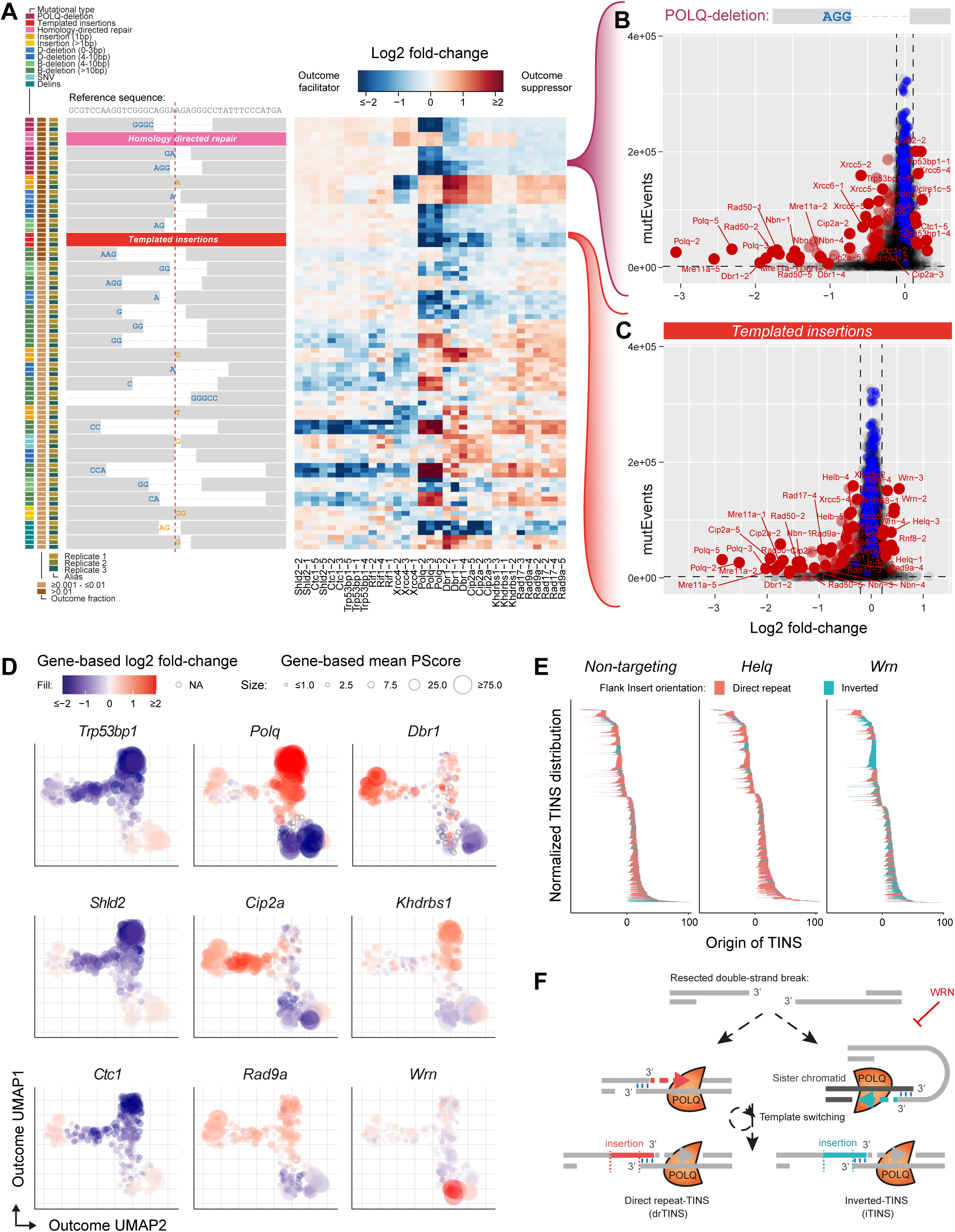
TMEJ genetics and outcomes; a role for WRN in suppressing inverted templated insertions. ***(A)*** Heatmap of the most abundant mutagenic repair outcomes (schematically depicted from top to bottom) for a selected set of sgRNAs, clustered based on outcome similarity. Each replicate (n=3) and each sgRNA are shown individually. ***(B-C)*** Volcano plots showing log2 fold-change (x-axis) versus total number of mutational events (y-axis) for a MH-driven deletion ***(B)*** and templated insertions outcomes ***(C)***. Vertical dashed lines mark 3* SD from the non-targeting sgRNAs (blue), and the horizontal dashed line indicates a minimum of 2,000 events per sgRNAs. sgRNAs consistently ranked as top outliers across all replicated (by Manhattan distance) are labelled in red; selected sgRNAs are further highlighted and annotated. ***(D)*** UMAP projections of repair outcomes for the indicated genes (as in Fig. 4). ***(E)*** TINS distribution plots for the indicated genotypes. Lines represent template origin within the flanks of the DSB, with the colour indicating TINS orientation. ***(F)*** Conceptual model for alternative TINS configurations. Direct-repeat TINS may result from interactions between opposing ends of a single DSB, whereas inverted-TINS may originate from engagement with a broken sister chromatid. WRN may suppress inverted TINS by limiting TMEJ activity on DSBs present in sister chromatid.

Beyond POLQ, the RNA lariat debranching enzyme DBR1^44^ and the splicing factor KHDRBS1 (also known as SAM68)^45^ – both not previously associated with DSB repair – as well as the RPA complex scored as modulators of TMEJ-driven outcomes (Fig. 7A-D). Also, all members of the MRN complex are found to promote TMEJ (Fig. 2 and Fig. 4) in line with end resection of blunt breaks needed to expose micro-homologous sequences. MUSIC also identifies components of the RAD9-RAD1-HUS1 complex and CIP2A-TOPBP1 as contributors to TMEJ outcomes (Fig. 2, Fig. 3, and Fig. 7A-D), aligning well with recent reports^19,46–48^. Interestingly, however, these proteins do not affect all outcomes equally. This is particularly evident for CIP2A, which specifically promotes TINS but has a much lesser effect on MH-mediated deletions (Fig. 7A). This finding also highlights the depth and resolution of our data, which allows for a detailed dissection of steps within a specific pathway. Moreover, the identified proteins provide hints towards the underlying mechanisms, in this case, it is tempting to speculate that the proposed tethering activity of CIP2A, keeping chromosome fragments together during mitosis^47,49–51^, may be essential for the recurrent engagement of DSB ends required for TMEJ iterations that generate TINS.

### WRN protects against a specific TMEJ-driven scar pattern

The finding that CIP2A loss differentially affects the TMEJ hallmark outcomes prompted us to zoom in on genes whose loss specifically affect TINS, a still poorly understood product of TMEJ. Beyond CIP2A, the most notable outliers are TOPBP1, TOP2A and HELB^52^. Conversely, TINS were specifically increased upon targeting HELQ^53,54^ and WRN^55–57^, identifying these factors as TINS suppressors (Fig. 7C). Previously, we found that HELQ, independently of POLQ, facilitates the repair of DSBs containing small flanking direct repeats^53^. Such repeat configurations also arise from TMEJ if the DSB ends are released upon initial POLQ-driven DNA synthesis (Supplementary Fig. S3), thus creating a preferred substrate for HELQ to reanneal the dissociated DSB ends for a renewed repair attempt. In the absence of HELQ, the strands from opposite DSB ends fail to anneal at the now available complementary sequences, instead, they initiate TMEJ at a new MH site, thereby generating a TINS (Supplementary Fig. S3).

The disruption of the RecQ DNA helicase WRN had a more intriguing effect on TINS formation: here, not only the frequency increased but most of the newly appearing TINS have insertions in the reverse-complement orientation (Fig. 7D,E), hereafter described as inverted-TINS (iTINS)^58^. To substantiate this phenotype, we generated monoclonal *Wrn*-knockout mES cell lines, and established DSB repair profiles for three different sequence contexts, including one in the endogenous X-linked *Hprt* locus. For all three sites we observed a profoundly increased ratio of iTINS to forward oriented direct-repeat TINS (drTINS) in WRN knockout cells (Supplementary Fig. S4 and S5). Also for HEK293, Hela, U2OS and RPE1*^hTert^* human cell lines, we found increased numbers of iTINS upon depletion of WRN (Supplementary Fig. S6), indicating that iTINS suppression by WRN is evolutionarily conserved.

### Inverted TINS are templated from the sister-chromatid

Using WRN as an exemplary case, we demonstrate that the resolution of the MUSIC is high enough to detect even the most marginal deviations from the norm (Fig. 3), thereby revealing novel aspects of DSB repair biology. To support this claim, we sought to address the question: what is WRN suppressing? According to prevailing models (e.g.^59^), iTINS generation by TMEJ occurs through the initial formation of an intramolecular hairpin, where a 3’ protruding tail folds back on itself stabilized by MH. After limited DNA extension and yet-unknown perturbation leading to dissociation, MH-mediated end joining to the opposite DSB end would generate an iTINS. However, our *in vivo* data contradicts this model. By aligning the template of each iTINS with the corresponding deleted sequence (Supplementary Fig. S7A), we reveal that iTINS cannot be constructed via a hairpin intermediate: extension from a fold-back structure cannot be templated by a sequence that has already been lost (see Supplementary Fig. S7B for schematic representation). Instead, as illustrated in Figure S7C, successful iTINS reconstruction through TMEJ requires an intact template, with the only logical candidate being the sister chromatid that also sustained a DSB, therefore being a viable substrate for TMEJ. While efficient Cas9 activity could generate DSBs in both sister chromatids (in S/G2 phases), alternatively, converging replication forks may collide with a DSB or a stably bound Cas9 (in S phase), breaking both sister chromatids. This latter scenario offers a potential mechanistic model for why WRN suppresses iTINS: when a fork encounters a DSB, it creates a 3’-tailed end on the lagging strand – ideal for TMEJ – and a blunt end on the leading strand, favouring KU binding and blocking TMEJ. WRN stabilizes KU thereby preventing resection^60,61^, potentially inhibiting TMEJ of the same ends of the broken sister chromatid. Without WRN, these ends may join, forming dicentric or acentric chromosomes, or iTINS in case of abrogated TMEJ. This explanation suggests that WRN plays a crucial role in preventing sister chromatid fusion, particularly in cases where DNA replication encounters a DSB, thus maintaining genomic stability.

## Discussion

In this study, we mapped – for each gene in the genome – the consequence of its genetic depletion on mutational DSB repair outcome patterns, creating a mutational signature catalogue we termed MUSIC. By directly sequencing the end-products of DSB repair using standard NGS, and analysing them through an updated SIQ software pipeline^17^, we gained insight into discrete DSB repair mechanisms. This approach, recently applied to smaller gene-sets^19^, complements traditional survival- or reporter-based methods, which, while powerful, lack nucleotide resolution.

We present a straightforward and adaptable lentiviral CRISPR/Cas9-based method, suitable for genome-wide analysis as well as focused sgRNA libraries or monoclonal knockout populations. Importantly, our method captures HDR outcomes as part of the mutational profile. MUSIC not only reconstructs canonical DSB repair pathways and core components from outcome data, but also reveals new details and nuances within those pathways. Some findings augment existing models – for example, staggered 5’ overhang removal by ARTEMIS versus fill-in by POLL, as divergent activities within c-NHEJ^18,19,62^, or the involvement of the FA pathway proteins in HDR^19,38^. Others support emerging concepts, such as mitotic tethering by CIP2A to promote TMEJ^47^ or HELQ-mediated suppression of TINS^53^.

We also identified several new factors that modulate repair pathway activity. While follow-up on each is beyond this study’s scope, we focused on WRN, which emerged as a conserved suppressor of iTINS. This served both as support for the claim that subtle changes in the overall profile are biologically meaningful, and as an entry point into the mechanism of (i)TINS formation – a hallmark of TMEJ that remains poorly understood. Structural analysis of iTINS argues against a fold-back model^58,59^, and instead points towards an interaction with the sister-chromatid. This hypothesis raises intriguing implications for CRISPR applications. To what extent does CRISPR induce sister chromatid exchanges (SCEs) or generate acentric/dicentric chromosomes that may trigger breakage-fusion-bridge cycles, anaphase bridges, or micronuclei formation? While (i)TINS appear to be a minor subset of repair outcomes, recent work identified TINS at replication-stress-induced complex structural variations in common fragile sites^63^, driven by mitotic POLQ. Moreover, (i)TINS have been observed in various disease alleles^15^, and function as an independent prognostic biomarker for survival in ovarian cancer^64^ underscoring the relevance of the underlying biology to contexts beyond CRISPR/Cas9.

In summary, MUSIC, alongside Repair-seq^19^ and a complementary c-NHEJ-focused study (released as a preprint during our manuscript preparation^32^), provides a rich dataset for DSB repair research. Transitioning from indirect phenotypic assays to direct mutational mapping of repair outcomes enables detailed investigation, especially when performed in genetic knockouts, or when combined with chemical inhibitors. To support this effort, we have developed a Shiny web-app (used to create all visualizations presented in this study) enabling full exploration of any gene-of-interest within MUSIC, to be released upon publication.

## Methods

### Resource availability

#### Lead contact

Further information and requests for resources and reagents should be directed to the Lead Contact, Marcel Tijsterman.

#### Materials Availability

Newly generated materials are made available upon request by contacting the lead contact

#### Data and Code availability

The targeted sequence data generated in this study have been deposited at NCBI SRA and will be made available upon publication of this manuscript. Original code is available on github https://github.com/RobinVanSchendel/MUSIC/. Data can be interactively explored through a web tool which will be made available upon publication of this manuscript.

### Experimental model and subject details

The 129/Ola-derived IB10 mouse embryonic stem (mES) cell line^18^ and HPRT-eGFP stable mES cell lines^39^ were maintained on 0.1% gelatin-coated dishes in complete medium, e.g. Buffalo rat liver (BRL)-conditioned KnockOut DMEM (Gibco) supplemented with 100 U/mL penicillin, 100 μg/mL streptomycin, 2 mM GlutaMAX, 1 mM sodium pyruvate, 1× non-essential amino acids, 100 μM β-mercaptoethanol (all from Gibco), 10% fetal calf serum (Capricorn Scientific) and leukemia inhibitory factor, as previously described^39,65^. Human RPE1-hTERT (gift from Rob Wolthuis), HEK293T, HeLa and U2OS cell lines^39^ were authenticated by means of the Promega PowerPlex Fusion 5c autosomal STR kit and were cultured in DMEM supplemented with GlutaMAX (Gibco), 10% fetal calf serum (Bodinco BV) and penicillin/streptomycin (Sigma), as previously described^39^. All cell lines were grown at 37°C and 5% CO2 and were tested negative for mycoplasma contamination.

### Method details

#### Plasmids

pLenti-Cas9-2A-Blast was a gift from Jason Moffat (Addgene# 73310). Mouse Improved Genome-wide Knockout CRISPR Library v2 was a gift from Kosuke Yusa (Addgene# 67988). pKLV2-U6gRNA5(BbsI)-PGKpuro2ABFP-W was a gift from Kosuke Yusa (Addgene# 67974). Lenti-multi-Guide was a gift from Qin Yan (Addgene# 85401). pLentiGuide-eGFP^24^ was a gift from Sylvie Noordermeer. Lentiviral packaging constructs pMDLg/pRRE, pRSV-Rev and pMD2.G were a gift from Didier Trono (Addgene# 12251, 12253, 12259). Plasmid pU6-(BbsI)_CBh-Cas9-T2A-mCherry was a gift from Ralf Kuehn (Addgene# 64324).

SgRNA sequences were cloned into the pKLV2-U6gRNA5(BbsI)-PGKpuro2ABFP-W backbone, the pLenti-multi-Guide backbone, or into the pU6-(BbsI)_CBh-Cas9-T2A-mCherry backbone using two custom complementary DNA oligos (IDT) consisting of the target sequence and *BbsI* overhangs, which were melted, annealed and ligated (Promega, Catalogue# M180B) into the relevant *BbsI*-digested (NEB, Catalogue# R3539S) plasmid backbone as described previously^66^. All constructs were sequence verified by Sanger sequencing using the following sequencing primer: (Seq)5’-GAGGGCCTATTTCCCATGATT-3’. sgRNA sequences are provided in Supplementary Table S1-S2.

#### Proof-of-principle setup

Stable Cas9-expressing IB10 (IB10-Cas9) mES cells were obtained through lentiviral transduction of the pLenti-Cas9-2A-Blast plasmid. Cells were kept under blasticidin selection (5µg/mL) for at least three passages prior to subsequent experiments. For the proof-of-principle experiment, IB10-Cas9 cells were transduced with lentiviruses to express pKLV2-U6gRNA5(BbsI)-PGKpuro2ABFP-W constructs in which the indicated sgRNAs (sg*Lig4*-2(JS), sg*Polq*-2(JS), sgNT-1 or sgNT-2) were cloned individually. Hereto, 5e^5^ cells were seeded on gelatin-coated 6-well plates containing complete medium supplemented with 8µg/mL polybrene at an MOI ≥ 1. The next morning, cells were put on selection with 2µg/mL puromycin for three days including one passage, after which the medium was refreshed without puromycin for recovery. By this time, the non-infected control well did not contain any surviving cells. To generate repair outcome profiles within the library backbone, 5e^5^ cells from each of these backgrounds were seeded on gelatin-coated 6-well plates and separately transduced with a library-targeting sgRNA expressed from the pLenti-multi-Guide backbone at an MOI of ∼2-5. The medium was refreshed the following day and cells were harvested three days after transduction. Genomic DNA was isolated from frozen cell pellets as described under *Genomic DNA isolation from cell lines*, and sequencing libraries were prepared as described under *MUSIC sequencing library preparation*. To determine outcome spectra per sg*Library*, libraries were prepared from each sample separately, wherein genotypic backgrounds targeted with the same sg*Library* were pooled after PCRI. To test for the occurrence of PCR jumping, samples targeted with sg*Library-1* were mixed with DNA from the untargeted sgRNA library prior to library preparation as written under *MUSIC sequencing library preparation*. PCR jumping events would present themselves as mutational repair outcomes linked to sgRNAs coming from the untargeted library. Results were demultiplexed and analysed as described under **Data processing pipeline**. Downsampling was done with Seqtk^67^.

#### Lentivirus production

Lentivirus production was carried out in 293T cells by calcium phosphate transfection using previously established methods^68^. Briefly, cells were seeded on 15-cm dishes one day before transfection. Medium was replaced without pen/strep 2h prior to transfection with 9µg ENV plasmid (VSV-G); 12.5µg pMDLg/pRRE plasmid; 6.25µg REV plasmid; 32µg transfer vector. Plasmids were added to a final volume of 1125µl 0.1×TE (1×TE: 10mM Tris [pH8.0] plus 1mM EDTA). Precipitate was formed by adding and mixing 125µl of 2.5M CaCl_2_ to the plasmids mixture, which was incubated for 5’ at RT prior to dropwise addition while vortexing to 1250µl of freshly prepared 2× HBS pH7.12. Precipitates were immediately added to the cultures. Medium was replaced 16h after transfection, and viral batches were collected 30h later, filtered with 0.45µm Acrodisc sterile syringe filters with a Supor PES membrane (PALL; Catalogue# 4614) and aliquoted. Aliquots were stored at -80°C and viral titers were determined by serial dilution prior to use.

#### MUSIC cell culture

IB10-Cas9 mES cells were transduced in bulk infections with the Mouse Improved Genome-wide Knockout CRISPR Library v2 (hereafter also referred to as Yusa-v2), comprising 90,230 sgRNAs targeting 18,424 genes. Hereto, 650e^6^ cells supplemented with 8µg/mL polybrene were transduced at an MOI of 0.7-1.0 and distributed over 0.1% gelatin-coated 15-cm cell culture dishes (12.5e^6^ cells per dish). The next morning, cells were put on selection with 2µg/mL puromycin for three days including one passage, after which the medium was refreshed without puromycin for recovery. By this time, the non-infected control plate did not contain any surviving cells. The MUSIC screen was performed five days after transduction with the library by collecting and splitting the cells over the different conditions (Supplementary Table S3), each condition containing 225e^6^ cells originating from the same bulk transduction.

Repair outcome spectra were generated by a second round of lentivirus transduction to express one of three library-targeting sgRNAs from the pLentiGuide-eGFP backbone. This second round of transduction was performed at an MOI of ∼2-5 supplemented with 8µg/mL polybrene per condition, in the presence of NU7441 (2µM) or ART-558 (10µM) when applicable. The medium was refreshed the next morning, and cells were harvested and aliquoted into 225e^6^ cell pellets three days after transduction, stored in -80°C until library preparation.

For the focused candidate library, the same approach was taken with the following changes: 300e^6^ cells were transduced at MOI 0.5 with the focused candidate library to generate a bulk knockout population, which was split on day 5 over three conditions containing 240e^6^ cells and hereafter processed independently. Targeting of the library backbone with sg*Library-1* to generate repair outcome spectra was performed as before, except harvested cells were aliquoted and stored as 300e^6^ cell pellets until library preparation (Supplementary Table S3).

#### MUSIC sequencing library preparation

Genomic DNA was isolated from screening samples using the Blood & Cell Culture DNA Kit (Qiagen; Catalogue# 13362, 10262, 19060), following manufacturers’ instructions for the preparation of cell culture samples and adhering to the stated capacity of 100e^6^ cells per 500/G genomic tip.

A nested PCR strategy was taken to amplify and retrieve the targeted region from genomic DNA by PCR, using sufficient parallel reactions to maintain coverage and hereby assuming that 6µg of DNA input is required per 1e^6^ cells/mouse genomes. The first round of PCR reactions was performed using NEBNext Ultra II Q5 Master Mix (NEB, Catalogue# M0544), containing 5µg genomic DNA, 0.2µM Fwd oligo 5’- TAGTGAACGGATCGGCACTG-3’, 0.2µM Rev oligo 5’-CTCCCCTACCCGGTAGAATTG-3’ per 50µl total reaction volume and run on a thermocycler with the following program: (1) 98°C for 3’, (2) 18 cycles of 98°C for 10”, 62°C for 30”, 72°C for 45”, (3) 72°C for 5’. Following PCR1 all parallel reactions were pooled and at least 600µl was purified using magnetic AMPure XP Beads (Beckman Coulter, Catalogue# A63882) in a 0.8X ratio and eluted in MQ. The second (nested) round of PCR was designed to generate the desired PCR fragment covering the target region and the sgRNA-region within paired-end short-read sequencing reads and simultaneously contained adaptors for the p5 and p7 index primers, p5 and p7 indexes, and flow-cell adaptor sequences. Hereto, PCR2 reactions were performed similarly to the first round, but contained 5µl of purified PCR1 template, 0.2-0.4µM Fwd oligo 5’- AATGATACGGCGACCACCGAGATCTACACnnnnnnnnTCGTCGGCAGCGTCAGATGTGTATAAGAGACAGttcgggtttat tacagggacag*c-3’, 0.2-0.4µM Rev oligo 5’- CAAGCAGAAGACGGCATACGAGATnnnnnnnnGTGACTGGAGTTCAGACGTGTGCTCTTCCGATCTaacttgctatgctgtt tccag*c-3 per 50µl reaction and for a total of at least 8 parallel reactions per sample, with the following program: (1) 98°C for 2’, (2) 10-12 cycles of 98°C for 10”, 65°C for 75”, (3) 65°C for 5’. Nnnnnnnn represents one of the following eight-nucleotide P5 barcodes: TGGTAATT, TTCCATGG, GAGAGTCA, GCGTCTAT, GGCGATAA, GTTCAGGT or P7 barcodes: GAGTCATG, GCCGGAAC, GGAGGTCC, TATAAGGC, TTATCCAT, TTGCCAGA (Supplementary Table S3). All parallel PCR2 reactions were pooled and purified using magnetic AMPure XP Beads in a 0.8X ratio and eluted in MQ. The quality and yield were assessed throughout library preparation using either a NanoDrop Spectrophotometer (Thermo Fisher Scientific, ND-1000), a Qubit 2.0 Fluorometer (Invitrogen, Catalogue# Q32866) using the dsDNA high sensitivity assay kit (Invitrogen, Catalogue# Q32854) or by gel electrophoresis. Finally, samples were measured for purity using a High Sensitivity DNA chip (Agilent, Catalogue# 5067-4626) on a 2100 Bioanalyzer (Agilent) and pooled in equimolar ratios prior to 150bp paired-end sequencing on a NovaSeq6000 (Illumina) or 300bp paired-end sequencing on a Miseq (Illumina).

#### Validation sgRNA library construction

To construct the custom library containing 2,400 sgRNAs, we identified the top outliers from our screen, 170 non-targeting controls and genes based on literature (including top genes from the previously published Repair-seq dataset^19^) (Supplementary Table S4). For each gene, we selected the three best performing sgRNAs on outcome redistributions from the Yusa-v2 library. Non-targeting sgRNA sequences and sgRNAs targeting genes not covered in the Yusa-v2 library were retrieved from the mouse-geckov2-library-b^69^. To each sgRNA sequence, we added the following 26-base overhang 5’- TATCTTGTGGAAAGGACGAAACACCG-3’ upstream and the 25-base overhang 5’- GTTTAAGAGCTATGCTGGAAACAGC-3’ downstream to enable bulk cloning. These 70-mer sequences were subsequently ordered as oPools synthesized by IDT and cloned into the pKLV2-U6gRNA5(BbsI)-PGKpuro2ABFP-W backbone (Addgene# 67974) using NEBuilder® HiFi DNA Assembly Master Mix (NEB, Catalogue# E2621L) according to manufacturers’ protocol. Briefly, the plasmid was digested with BbsI-HF (NEB, Catalogue# R3539S) followed by gel extraction and purification. Assembly was performed in five parallel reactions, wherein each assembly mixture was composed of 40-50ng digested and purified plasmid, 5µl of diluted ssODN (0.2µM in 1x NEBuffer 2.0), 2x NEB HiFi assembly mixture and MQ to a total volume of 20µl per reaction followed by a 1h incubation at 50°C and subsequent cleanup using the QIAquick PCR Purification Kit (Qiagen; Catalogue# 28106) following manufacturers’ protocol. The assembly mix was electroporated (Biorad; program for bacteria, Ec1, one pulse) into Endura Duos Electrocompetent Cells (Lucigen; Catalogue# 60242-1; Lot# 30680) using Gene Pulser/MicroPulser Electroporation Cuvettes (Biorad; Catalogue# 1652083) following manufacturers’ instructions and using ROC Expression Recovery Medium (Lucigen; Catalogue# 80030-1). Total transformation efficiency was determined by serial dilution of 10µl onto ampicillin-containing agar plates and was calculated to exceed >2000X coverage per sgRNA. The remainder volume was incubated in ampicillin-containing 2x LB liquid cultures in a shaker o/n at 32°C. The culture volume was doubled the next day for another 2-4 hours of incubation before harvesting bacterial pellets. Plasmid libraries were isolated using column-based purification methods (Macherey-Nagel; Catalogue# 74041050).

#### Transfections

Transfections were performed using previously established methods^39^. Briefly, mES cells were trypsinised, counted and resuspended in complete medium without pen/strep and transfected in suspension using Lipofectamine 2000 (Invitrogen, Catalogue# 11668019):DNA ratio of 2.4:1. Generally, 2µg DNA in Opti-MEM (Gibco) and 4.8µl Lipofectamine in Opti-MEM (Gibco) were both resuspended and incubated for 5’ at RT prior to mixing together and incubating for 20’ at RT. This mixture was added to an equal volume cell suspension containing 1×10^6^ cells and incubated in round-bottom tubes at 37°C for 30 minutes, which was subsequently seeded on gelatin-coated 12-well plates containing complete medium. The medium was refreshed the next day.

#### Generation of Wrn-knockout cell lines

IB10-Cas9 mES cells and two independent clones of *Hprt-eGFP* parental mES cells, which were generated previously^39^, were transfected with Cas9-WT-2A-mCherry constructs expressing indicated sgRNAs (Supplementary Table S5). 48h post transfection, cells were seeded at low density on gelatin-coated 10-cm dishes and maintained until colonies formed, changing medium regularly. Colonies were picked, trypsinised and transferred to gelatin-coated flat-bottom 96-well plates, and at (semi)-confluence split 1:3. One plate was used for freezing; the other for DNA isolation and genotyping by PCR. Successful knockout clones were retrieved from the frozen stock, subcultured, and used in experiments as indicated.

#### Genomic DNA isolation from cell lines

Unless stated otherwise, genomic DNA was isolated from frozen cell pellets as previously described^39^. Briefly, up to 4e^6^ cells were lysed in 500μl 10mM Tris-HCL pH7.5, 10mM EDTA, 200mM NaCl, 1% SDS and 0.4mg/mL Proteinase K, incubated at 55°C for 3-16 hours. Lysis was neutralized by 150μl saturated NaCl followed by centrifugation at 20,000g for 10’. DNA was precipitated and extracted by adding one volume of isopropanol to the supernatant, followed by centrifugation, washing with 70% ethanol and resuspension in TE.

#### Tide analysis

Target loci for sgmm*Lig4-2(JS)* were amplified by PCR using GoTaq (Promega, Catalogue# M7848) according to manufacturer’s protocol with the following program: (1) 95°C for 10’, (2) 35 cycles of 95°C for 30”, 60°C for 30”, 72°C for 60”, (3) 72°C for 5’. Amplified products were analysed by gel electrophoresis, diluted 50X in MQ and 2µl of diluted product was submitted for Sanger sequencing. Primers used were: (Fwd)5’- CAACGTCACCACAGATCTGG-3’, (Rev)5’-CTAAGGATCTCATACCTCTTC-3’, (Seq)5’- CAACGTCACCACAGATCTGG-3’.

Target loci for sgmm*Polq-2(JS)* were amplified by PCR using GoTaq (Promega, Catalogue# M7848) according to manufacturer’s protocol with the following program: (1) 95°C for 10’, (2) 35 cycles of 95°C for 30”, 60°C for 30”, 72°C for 60”, (3) 72°C for 5’. Amplified products were analysed by gel electrophoresis, diluted 50X in MQ and 2µl of diluted product was submitted for Sanger sequencing. Primers used were: (Fwd)5’-GCTAGCATTTTTCTGTATAGG-3’, (Rev)5’-CAAATGTCCTTGTTTCCCTCC-3’, (Seq)5’- GCTAGCATTTTTCTGTATAGG-3’.

Target loci for *Wrn* were amplified by PCR using GoTaq (Promega, Catalogue# M7848) according to manufacturer’s protocol with the following program: (1) 95°C for 10’, (2) 35 cycles of 95°C for 30”, 60°C for 30”, 72°C for 60”, (3) 72°C for 5’. Amplified products were analysed by gel electrophoresis, diluted 50X in MQ and 2µl of diluted product was submitted for Sanger sequencing. Primers used to amplify sg*Wrn*-2 were: (Fwd)5’-GAAAAATAGAACCGCTTCCTTTCC-3’, (Rev)5’-CTAAAGCGGGCCACAAGGAT-3’, (Seq)5’- GAAAAATAGAACCGCTTCCTTTCC-3’; sg*Wrn-4* was amplified using (Fwd)5’-CGAACGCCACCTCTAGTAACA-3’, (Rev)5’-AGCCTAGACTTTGAACAGACCTT-3’, (Seq)5’-CGAACGCCACCTCTAGTAACA-3’.

Sanger sequences were analysed with TIDE^70^ using the PCR product from parental control gDNA as reference for alignment. Genotypes of samples that could not be decomposed by TIDE (e.g. due to the presence of deletions that exceeded the TIDE decomposition window) were determined through cloning of the PCR products using pGEM®-T Easy Vector Systems (Promega, Catalogue# A1360), followed by Sanger sequencing and manual alignment.

#### Immuno blotting

Protein samples were generated by lysing cells in SNTBS lysis buffer (2% SDS, 1% NP-40, 150mM NaCl, 50mM Tris pH7.5). Lysates were passed through a syringe and heated at 95°C for 10 minutes, centrifuged, and supernatants transferred to new tubes. Protein concentrations were determined using the Pierce™ BCA Protein Assay Kits (ThermoFisher Scientific, Catalogue# 23225). Equal protein quantities were prepared in 4x NuPAGE LDS Sample buffer (Invitrogen, Catalogue# NP0007) and 50mM DTT, and incubated at 70°C for 10’ prior to loading and separation on Novex 4-12% Bis-Tris gradient gels (Invitrogen, Catalogue# NW04125BOX) using MOPS SDS running buffer and transferred onto Immobilon-FL membranes (Merck Millipore, Catalogue# #IPFL00010). Membranes were blocked using Blocking Buffer for Fluorescent Western Blotting (Rockland, Catalogue# MB-070) diluted 1:1 in PBS for at least 30’. The primary antibodies used to analyse protein expression were: anti-WRN (8H3, 1:1000) mouse monoclonal (Cell Signalling, Lot# 5) and anti-Tubulin (DM1a, 1:10000) mouse monoclonal (Sigma, Lot# 029M4842V). The secondary antibody was CF770 goat anti-mouse IgG (Biotium, Catalogue# 20077, Lot# 21C0302, 1:5000). Antibodies were diluted in 1:1 mixtures of Blocking Buffer for Fluorescent Western Blotting (Rockland, Catalogue# MB-070) and PBS-T. The Odyssey infrared imaging scanning system (LI-COR biosciences) was used to detect protein expression.

#### Targeted sequencing of Cas9-induced repair outcomes

IB10-Cas9 mES *Wrn*-knockout clones or parental cells were separately transduced with lentivirus to express either the sgNT-1, sgNT-2 or sg*hsLig4-2(JS)* (acts as non-targeting on the mouse genome) from the pKLV2-U6gRNA5(BbsI)-PGKpuro2ABFP-W backbone using 5e^5^ cells per transduction and seeded on gelatin-coated 6-well plates containing complete medium supplemented with 8µg/mL polybrene. In each of these populations, the pKLV2-U6gRNA5(BbsI)-PGKpuro2ABFP-W backbone was subsequently targeted in a second round of lentivirus transduction to express sg*Library-1* or sg*Library-3* from the pLentiGuide-GFP backbone, again using 5e^5^ cells per transduction and seeded on gelatin-coated 6-well plates containing complete medium supplemented with 8µg/mL polybrene. Samples were harvested and processed as described under *Genomic DNA isolation from cell lines* and *MUSIC sequencing library preparation*. A coverage of at least 5e^5^ was maintained throughout all steps. For each genetic background, the three different non-targeting sgRNAs, which were generated, cultured and processed independently, act as biological replicates. The NGS data were demultiplexed based on the non-targeting sgRNAs prior to further analysis (see below).

*Hprt-eGFP* stable parental or *Wrn*-knockout mES cell lines were transfected with a Cas9-WT-2A-mCherry construct to express the indicated *Hprt*-targeting sgRNA, and split two days after transfection. On day five post transfection, cells ware harvested and frozen before gDNA isolation as described under *Genomic DNA isolation from cell lines*. Samples for Illumina sequencing on the *Hprt* locus were prepared as previously described^65^. Briefly, specific primers were used to amplify the targeted region, yielding ∼150-200bp products and containing adaptors for the p5 and p7 index primers (sg*Hprt-eGFPex3*(Fwd)5’- GATGTGTATAAGAGACAGGGACTGAAAGACTTGCTCGAGAT-3’, sg*Hprt-eGFPex3*(Rev)5’- CGTGTGCTCTTCCGATCTACACAGTAGCTCTTCAGTCTGATAAAA-3’, respectively), using Phusion High-Fidelity DNA polymerase (Thermo Fisher Scientific, Catalogue# F-530L) with the following program: (1) 95°C for 10’, (2) 25 cycles of 95°C for 10”, 60°C for 30”, 72°C for 30”, (3) 72°C for 5’. Products were purified using 0.8X volume of magnetic AMPure XP beads (Beckman Coulter, Catalogue# A63882) according to the manufacturers protocol and eluting DNA in 20µl MQ. A second round of PCR was used to add flow-cell adaptor sequences by 0.3µM p5 and p7 index primers p5(Fwd)5’- AATGATACGGCGACCACCGAGATCTACACnnnnnnnnTCGTCGGCAGCGTCAGATGTGTATAAGAGACA*G-3’ and p7(Rev)5’-CAAGCAGAAGACGGCATACGAGATnnnnnnnnGTGACTGGAGTTCAGACGTGTGCTCTTCCGATC*T-3’ with nnnnnnnn representing barcodes as indicated (Supplementary Table S3). PCR products were purified with 0.8X volume of magnetic AMPure XP beads and eluted in 20µl MQ. The quality and yield was assessed as described under *MUSIC sequencing library preparation* and samples were subsequently submitted for NGS. Experiments were performed in duplicate on two independent clonally derived knockout lines per genotype.

Human cell lines were first transduced with the pLenti-Cas9-2A-Blast plasmid and selected for at least three passages prior to infection with either non-targeting or *Wrn*-targeting sgRNAs expressed from the pKLV2-U6gRNA5(BbsI)-PGKpuro2ABFP-W backbone. Targeting of the KLV2 backbone was performed analogous to the MUSIC setup, but with the indicated library-targeting sgRNAs, split over two replicates and using the following concentrations for blasticidin and puromycin selection, respectively: HEK293T (5µg/mL; 3µg/mL), Hela (2µg/mL; 1µg/mL), U2OS (5µg/mL; 3µg/mL) and RPE1-hTERT (10µg/mL; 5µg/mL). Samples were harvested and processed as described under *Genomic DNA isolation from cell lines* and *MUSIC sequencing library preparation*. Experiments describing human cells were performed in a pooled manner, a pool either consisting of two non-targeting sgRNAs or of two non-targeting sgRNAs and four *WRN*-targeting sgRNAs, demultiplexed prior to further analysis. These samples were combined and reduced into either a sgNT control pool, or a sg*WRN* pool wherein only the *WRN*-targeting sgRNAs were included.

All NGS data from clonal knockout lines were processed by a custom JAVA program and the output was analysed and plotted using the corresponding SIQplotteR web tool^17^. The distribution of TINS was determined by subcategorising events of the type “TINS” into drTINS (isFirstHit equals “R0” or “0L”) or iTINS (isFirstHit equals “rcR0” or “0Lrc”). The ratio of iTINS was calculated to the total TINS using the total read counts. For plotting, TINS were normalized to 1. The mapping of inverted templated insertions was done by filtering reads derived from the parental *Hprt-eGFP* mES cell lines targeted with sg*Hprt-eGFPex3,* targeting a haploid locus, for all insertions that could be fully explained by a single iteration (isGetSubS2 equals “<rcR0>” or “<0Lrc>”). From these insertions, the position of the deletion (“delRelativeStart”, “delRelativeEnd”) was overlayed with the position of the insertion template (“isStartPosRel”, “isEndPosRel”) and plotted, in which all events that had at least 1bp overlap were scored as “hairpin-incompatible”.

#### Statistical analysis

GraphPad Prism software (GraphPad Prism 10.2.3) was used for statistical analysis and production of the bar graphs and to plot the origin of iTINS. Statistical significance was assessed by one-way ANOVA or unpaired two-tailed *t*-test as indicated.

### Bulk data processing pipeline

Reads derived from bulk library data were processed by a custom JAVA program^17^ (https://github.com/RobinVanSchendel/SIQ), but with the following adaptations to enable processing of bulk data: paired-end NGS was designed such that the repair outcome could be determined from one end of the sequencing read and the sgRNA could be determined from the other end of the sequencing read. For each sample, paired-reads were first demultiplexed into barcode and gene based on an exact match with an sgRNA from the Yusa-v2 library. The barcode was added to each read and binned into 187 files (containing multiple genes per file). The resulting files were then analysed by SIQ to classify reads into the following events: deletion, delins (deletion with an insertion of de novo DNA at the location of the deletion), insertion, tins (templated insertion; events containing a ≥6 nt insertion with a large enough match to a DNA sequence in the immediate vicinity of the break junction (100 bp in both directions and orientation) with the probability of finding such a match of the same size in a scrambled sequence being <10%; for more detailed information see^17^), HDR (homology-directed repair; a specific deletion-insertion event that can be mapped to the targeting backbone), SNV (single-nucleotide variant) or wild-type (i.e. identical to the reference sequence). Additional parameters such as deletion/insertion-size and microhomology length for deletions were included for each event. Per sgRNA, outcomes were only included in the final data if at least two reads support that event. Reads matching wild-type and SNV outcomes outside the DSB site were excluded from analysis: all data were analysed relative to the total amount of mutational reads.

For the genome-wide MUSIC, repair outcome distributions (demultiplexed per sgRNA in the previous step) were then compiled into a mutational signature catalogue, maintaining only those sgRNAs that were represented by at least 200 mutational reads. Note that although only the top 20 most prevalent outcomes were written to the catalogue to limit its file size, the mutational types and fractions were calculated against all mutational reads including those outcomes that do not belong to the top 20 outcomes. To increase statistical power on low-complex data (where infrequent outcomes will not be present in all sgRNAs), we further aggregated outcomes into the following mutational types which we presumed resulted from comparable biochemistry: (i) “Insertion_1bp” contained insertions of one base pair; (ii) “insertions_>1bp” contained all insertions – not accompanied by a deletion – larger than one base pair; (iii) “Deletion_PQ” contained the three most common deletions that were verified to be POLQ-dependent from the pilot experiment; (iv) “deletion_≥15bp” contained deletions of at least 15bp in size”; (v) “deletion_1-14bp” contained the remaining deletions that did not fall into any of the previous categories; (vi) “HDR” contained a specific delins (35bp deletion and 11bp insertion) that could be mapped entirely to the targeting backbone, hence suggesting homology-usage and template repair in the generation of this repair product; (vii) “delins” and (viii) “tins” were maintained without further subdivision. Due to the lack of non-targeting sgRNAs within the Yusa-v2 library, the average outcome distribution of the entire population was used as reference for downstream analysis and was termed “*SampleAvg*”. The log2 fold-change of each mutational type frequency was calculated per sgRNA relative to the *SampleAvg*, after which gene-scores were calculated by taking the mean change over all sgRNAs and conditions, e.g. sgRNAs and conditions were weighted equally irrespective of the read-count. Pvalues were calculated by a -log10 transformed binomial test (binom.test function from the R Stats Package) using the outcome distribution from the population mean *SampleAvg* as the hypothesized probability of success. The number of reads belong to each corresponding outcome was used as input for the number of successes and the total mutational reads for a sgRNA was used as input for the number of trials.

The list of candidate genes was established by calculating the mean gene-based redistribution profiles of the repair mutational types for all genes of which at least 3 sgRNAs passed the above filters. Outliers were defined per mutational type as genes that resulted in a mean redistribution of at least 2.5×SD (SD taken from the *SampleAvg*), a mean absolute log2 fold-change of at least 0.25 and a mean -log10 transformed Pval of at least 7.5, subsequently ranked by mean log2 fold-change. From this list, at least the top 20 and at most the top 150 deviating genes (ranked by mean Pval) were included per direction. This candidate gene list was supplemented with non-targeting controls and genes based on literature^19^ as described above.

The focused candidate library was processed using the same pipeline as MUSIC, but with the following changes: sgRNA sequences were matched to the focused candidate library (Supplementary Table S4), profile redistributions were calculated relative to the average from all non-targeting sgRNAs instead of the *SampleAvg*. Deletions were additionally separated into size (“1-3bp”, “4-10bp”, “>10bp”) and directionality (“B” – bidirectional, spanning both sides of the break junction; “D” – directional, spanning only one side of the break junction). Heatmaps were generated using heatmap.2 from the gplots package, umaps were generated using the umap package.

The MUSIC web tool was built as a user-explorable resource to enable any user to look up the behaviour of any gene of interest in the MUSIC and validation datasets. All settings have been predefined such as used in this manuscript, but full customization of filters is possible. This app will be made available upon publication of the manuscript.

## Acknowledgements

The authors thank S. Kloet and the staff of the Leiden Genome Technology Center (LGTC) for their support to sequencing and J.J. Goeman from the department of Biomedical Data Sciences for advice on statistics relating to the MUSIC data. This work was funded by a Unique High-Risk grant from The Dutch Cancer Society (KWF, 2021-2/13905) to M.T.

## Contributions

M.B. and M.T. conceived and designed the study. M.B. performed the experiments. R.v.S. adapted the Sequence Interrogation and Quantification program to handle bulk libraries, performed the bioinformatical analyses, and developed the corresponding Shiny web-tool with input from M.B. and M.T., M.B. and M.T. interpreted the experimental results and prepared the manuscript with input from R.v.S. The study was supervised by M.T.

## Ethics declarations

## Competing interests

The authors declare no competing interests.

## A Supplementary Figure legends

**Supplementary Figure S1:**
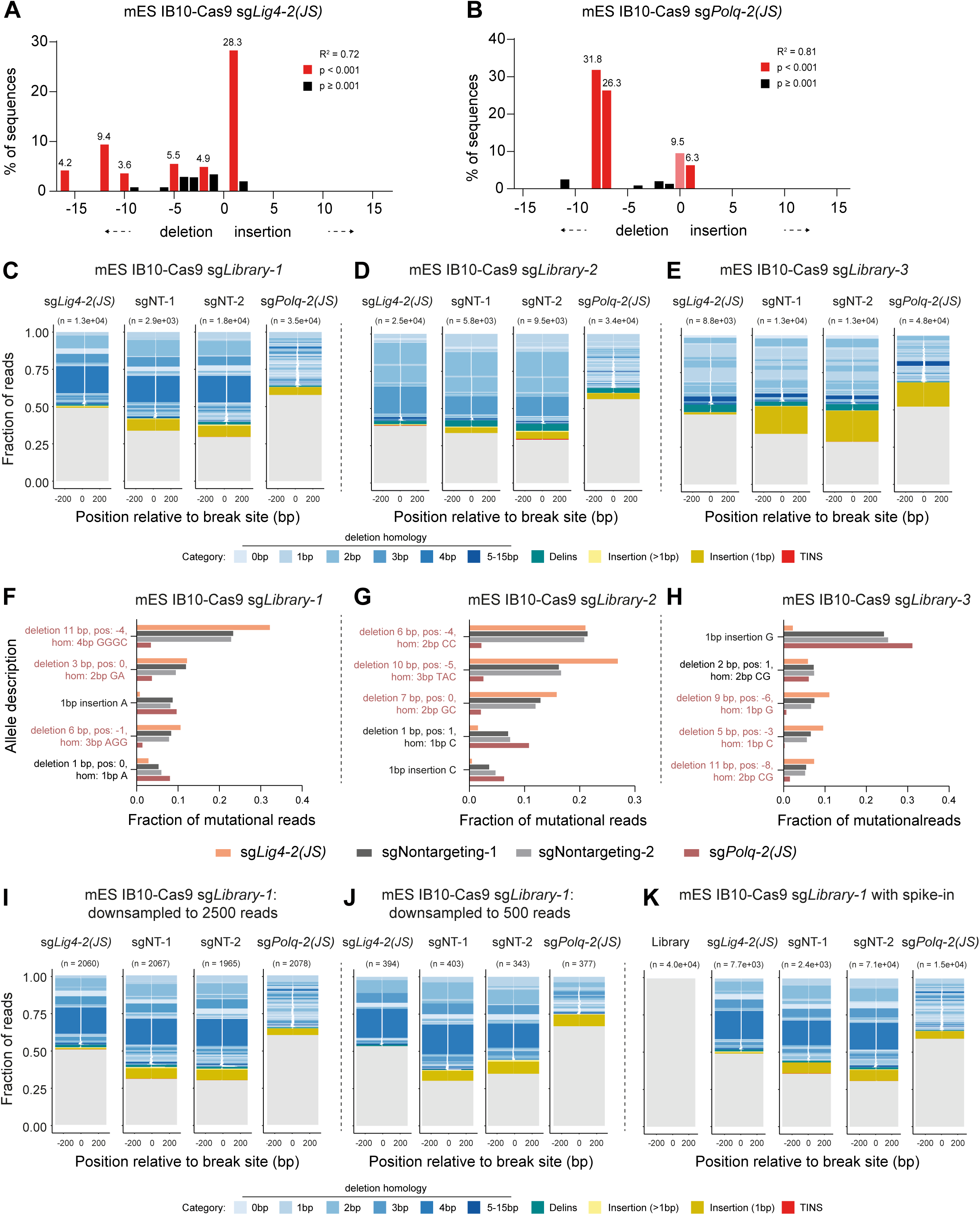
Gene-specific mutational signatures can be generated in a bulk-sequencing approach. ***(A,B)*** IB10-Cas9 mES cells were transduced with lentivirus to express the indicated sgRNA. Genomic DNA was isolated from puromycin resistant cells followed by PCR of the target locus, Sanger sequencing, and analysed using TIDE^70^ to determine mutation frequencies. The indel size range is plotted and colour coded according to the indicated statistical significance. ***(C-E)*** Repair profiles for Cas9-induced DSBs for the indicated mES cell lines. Cells were first transduced with a targetable vector, then with the indicated sgRNA, and processed analogous to the MUSIC workflow three days after transduction. Samples were pooled after the second PCR and after NGS demultiplexed based on the sgRNA sequence contained in each read. Tornado plots show repair outcomes (horizontal line interrupted by deleted sequence), scaled by read frequency (line thickness), and colour-coded by outcome category: deletion (shading indicating degree of MH), delins, insertion (>1bp), insertion (1bp), and TINS. Cas9-induced DSB is positioned at 0 (x-axis). Total read count is shown above each graph. ***(F-H)*** Quantification of the five most abundant mutated alleles in the non-targeting controls, normalized to the total mutagenic read count. Allele descriptions include the outcome category, and for deletions also the size, the starting position and the length of MH. The three most abundant POLQ-dependent deletions are coloured and annotated and used in Figure 1 as “PQ-Del”. ***(I,J)*** Computational down-sampling the data presented in *(C)* to the indicated read depth. ***(K)*** DSB repair profiles generated as in *(C)* but now also including a DNA sample from Yusav2 library-containing cells that were not targeted, illustrating a clear lack of potential mis annotation - the library sample is devoid of repair events.

**Supplementary Figure S2:**
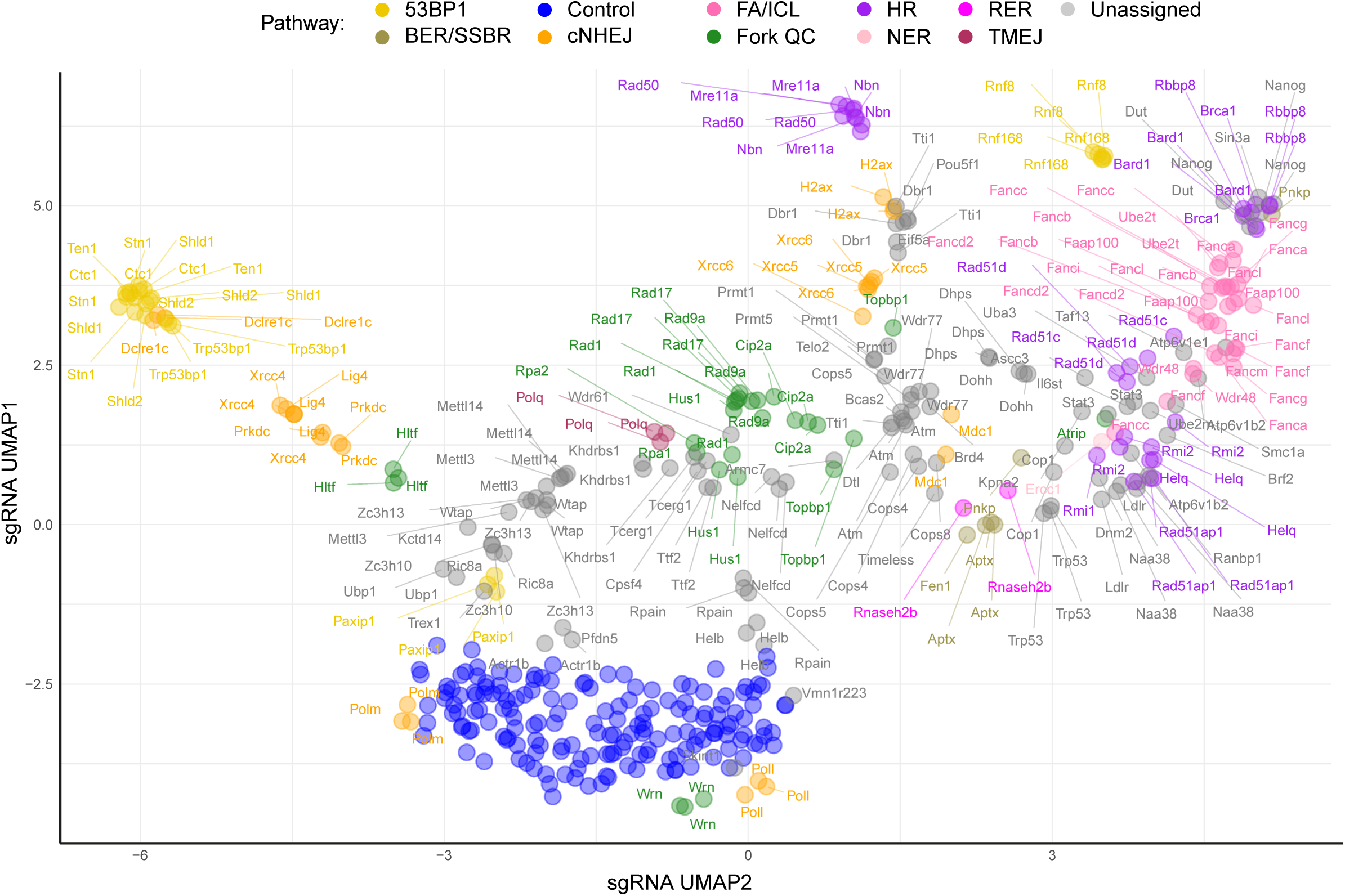
UMAP output including all labels, belonging to Fig 3. The same UMAP as shown in Figure 3, now including all gene labels.

**Supplementary Figure S3:**
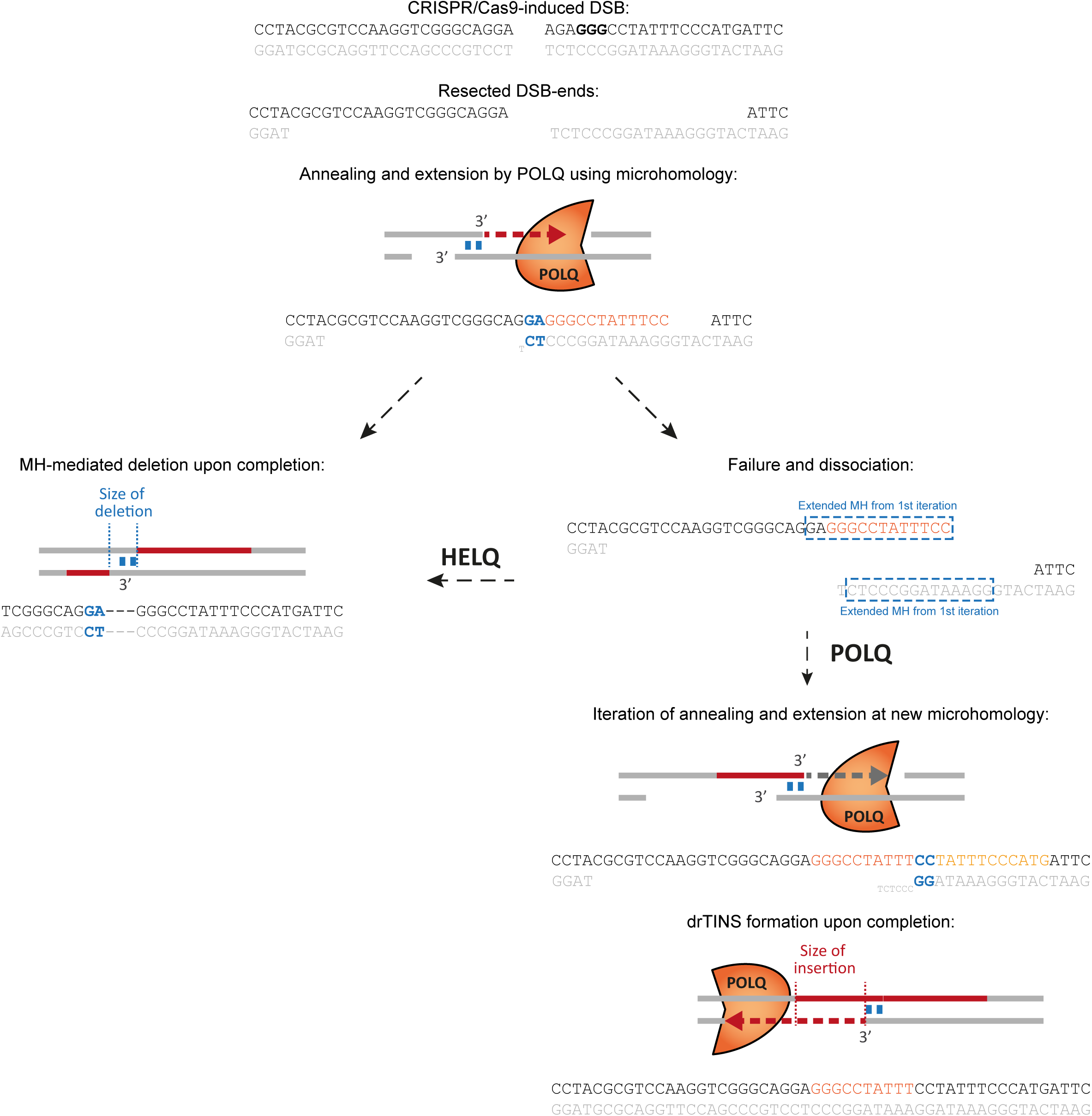
Proposed model for how HELQ suppresses templated insertions. Reconstruction of TMEJ outcomes initiated by a 2-bp MH. Successful annealing, extension, and gap filling typically results in a MH-mediated deletion. However, if TMEJ fails to complete and the opposing DNA ends dissociate, two alternative outcomes may occur: (i) annealing and extension initiate at a new MH site, and subsequent gap filling produces a templated insertion. (ii) the extended MH generated by POLQ can reanneal to the original position on the opposing DSB end, and gap filling then produces a deletion outcome indistinguishable from that of uninterrupted TMEJ. HELQ is proposed to suppress TINS formation by promoting reannealing of the extended MH, thereby favouring completion of canonical TMEJ.

**Supplementary Figure S4:**
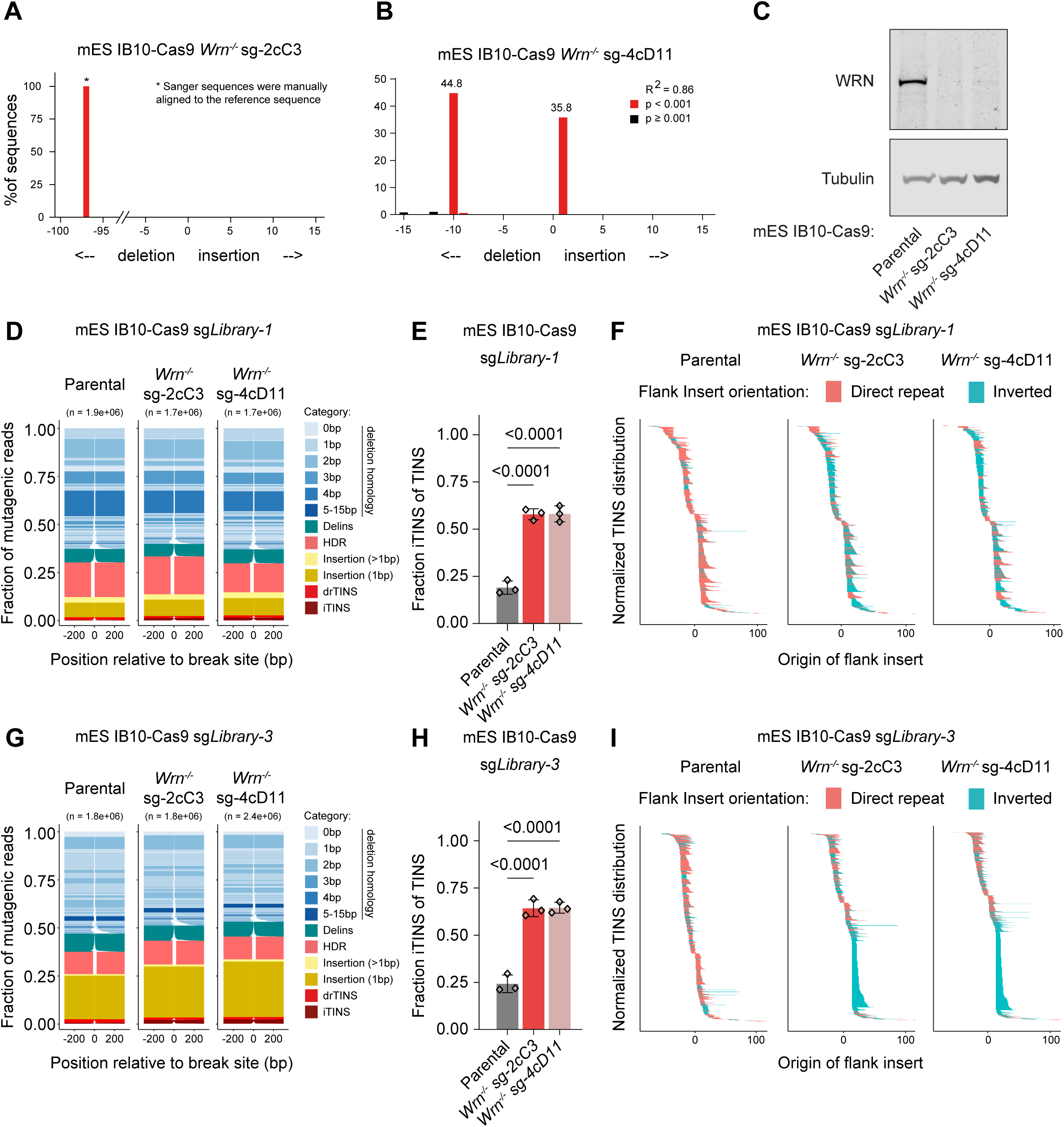
Clonal knockout lines validate inverted TINS suppression by WRN. ***(A, B)*** TIDE^70^ analysis of two independent IB10-Cas9 *Wrn*-knockout clones. ***(C)*** Western blot showing WRN protein levels in parental and *Wrn*-knockout clones. ***(D, G)*** DSB repair profiles for parental and *Wrn*-knockout IB10-Cas9 cells. Cells were first transduced with a targetable vector, then with the indicated targeting sgRNA, and processed analogous to the MUSIC workflow three days after transduction. Tornado plots show repair outcomes (horizontal line interrupted by deleted sequence), scaled by read frequency (line thickness), and colour-coded by outcome category: deletion (shading indicating degree of MH), delins, HDR, insertion (>1bp), insertion (1bp), drTINS, iTINS. Cas9-induced DSB is positioned at 0 (x-axis). Data represent the sum of three experiments. ***(E, H)*** Quantification of iTINS as a fraction of total TINS. Bars show mean ± SD, individual replicates are indicated. Statistical significance was tested by one-way ANOVA. ***(F, I)*** TINS distribution plots for the indicated genotypes. Lines represent template origin within the flanks of the DSB (position 0), with the colour indicating TINS orientation. The sum of three experiments is presented.

**Supplementary Figure S5:**
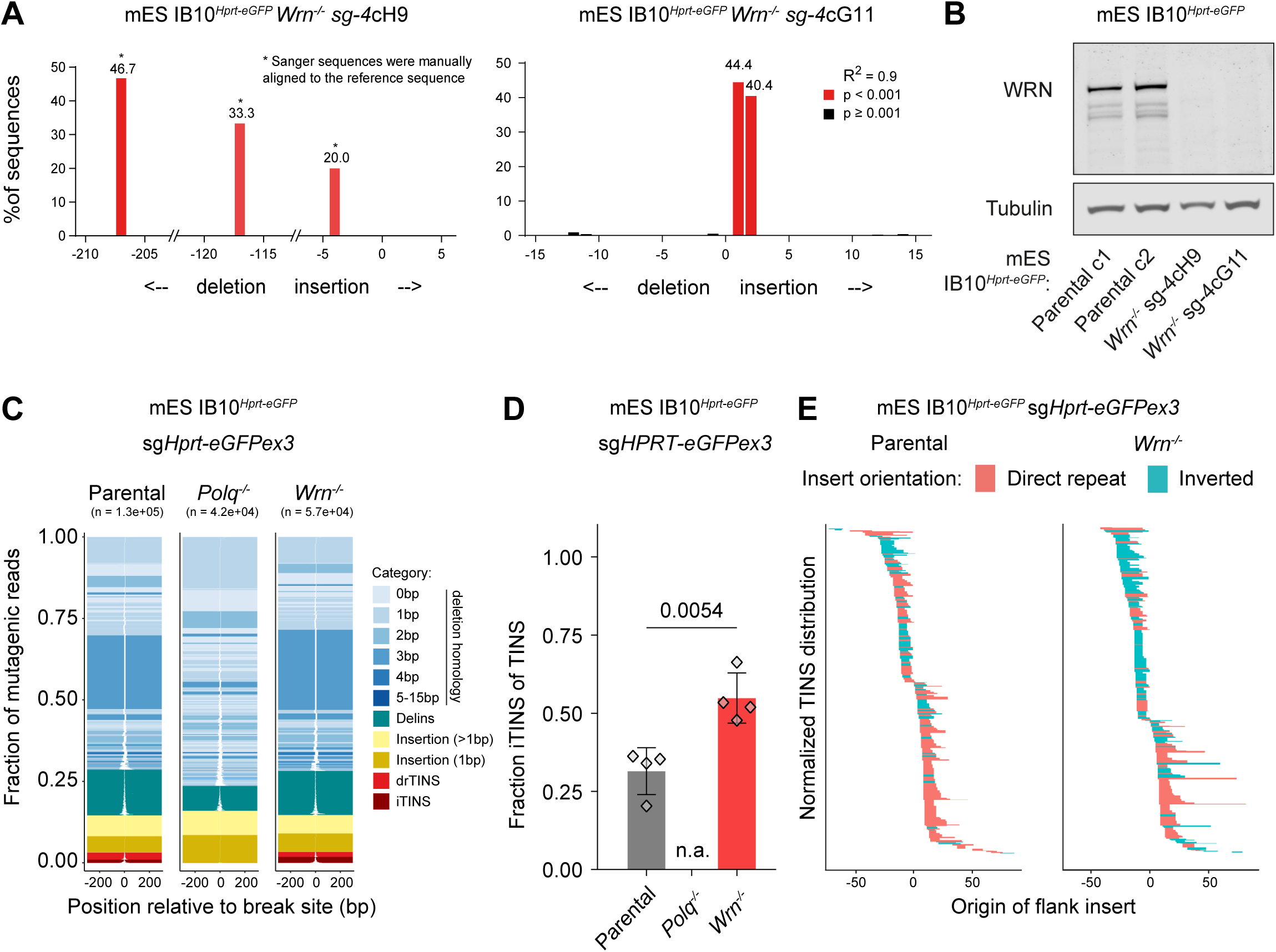
WRN suppresses inverted TINS at a genomic Hprt site. ***(A)*** Allele modifications of two independently derived *Wrn*-knockout IB10*^Hprt-eGFP^* mES clones. ***(B)*** Western blot showing WRN protein levels in parental and *Wrn*-knockout IB10*^Hprt-eGFP^* mES clones. ***(C)*** DSB repair profiles for the indicated mES cell lines transfected with Cas9 and sg*Hprt-eGFPex3*. Tornado plots show repair outcomes (horizontal line interrupted by deleted sequence), scaled by read frequency (line thickness), and colour-coded by outcome category: deletion (shading indicating degree of MH), delins, HDR, insertion (>1bp), insertion (1bp), drTINS, iTINS. Cas9-induced DSB is positioned at 0 (x-axis). Data represent the sum of four experiments (two independent clones per genotype). ***(D)*** Quantification of iTINS as a fraction of total TINS. Bars show mean ± SD, individual replicates are indicated. Statistical significance was assessed using an unpaired two-tailed t-test. ***(E)*** TINS distribution plots for the indicated genotypes. Lines represent template origin within the flanks of the DSB (position 0), with the colour indicating TINS orientation.

**Supplementary Figure S6:**
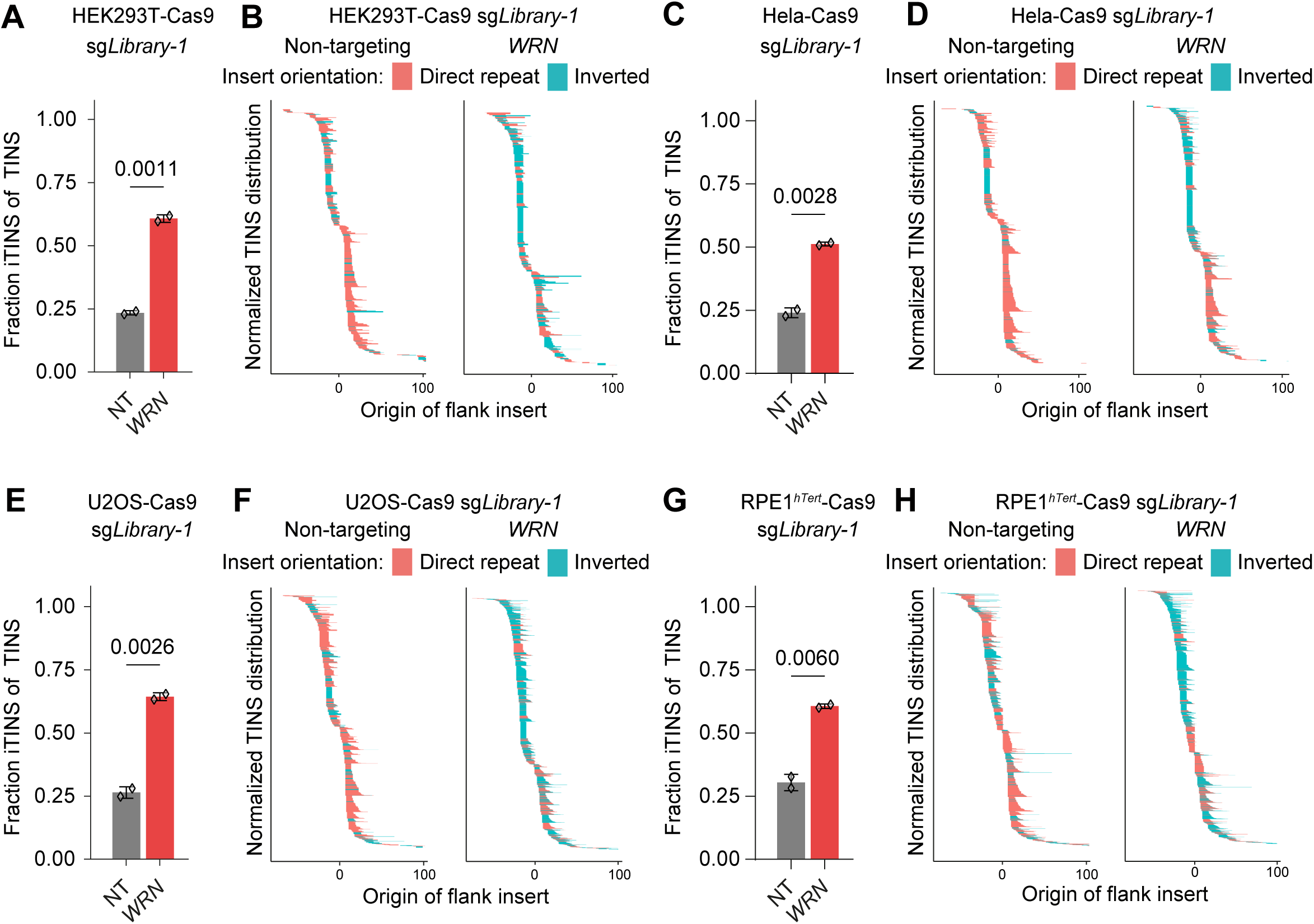
Suppression of inverted TINS by WRN in human cells. ***(A-H)*** Cas9-expressing human cell lines transduced to either express a non-targeting or a WRN-targeting sgRNA, followed by a library-targeting sgRNA to generate repair profiles. ***(A,C,E,G)*** Quantification of iTINS as a fraction of total TINS. Bars show mean ± SD, individual replicates are indicated. Statistical significance was assessed using an unpaired two-tailed t-test. ***(B,D,F,H)*** TINS distribution plots. Lines represent template origin within the flanks of the DSB (position 0), with the colour indicating TINS orientation. Replicates of the same genotype were combined and normalized to 1.

**Supplementary Figure S7:**
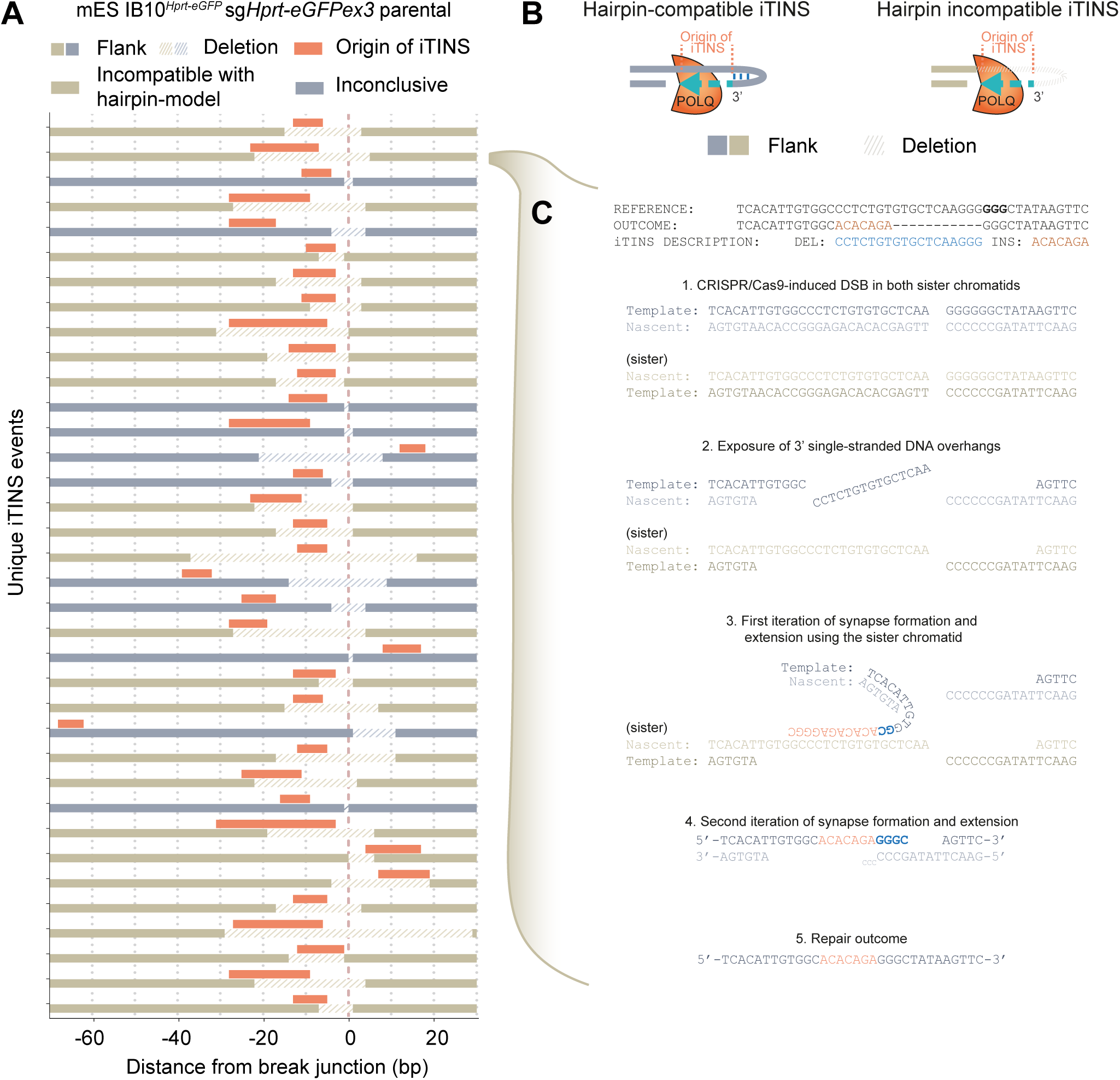
Aetiology of iTINs. ***(A)*** The data presented in Figure S5 was first filtered for iTINS that contained an insertion that completely mapped to the flank of the DSB (i.e. devoid of non-templated bases or multiple matching stretches). For these iTINS, the flanks are plotted as a solid bar, the deletion as a dashed bar and the templated inverted insertions as orange bars. All segments are plotted relative to the DSB at position 0 (bp). Only events from parental cells are plotted for clarity. Only iTINS without overlap between the insertion and the deletion can be explained with a hairpin model, schematically illustrated in ***(B,left panel)***; these are marked “inconclusive” as they can also be explained through an alternative sister chromatid-based model. iTINS containing insertions that overlap with the deleted sequence are “incompatible with a hairpin-model”, as illustrated in ***(B, right panel)***. Events in ***(A)*** are colour coded accordingly. The vast majority of iTINS are incompatible with a hairpin model, whereas all iTINS can be explained by invoking the engagement of a second molecule, i.e. the sister chromatid. ***(C)*** Step-by-step reconstruction of a hairpin-incompatible iTINS, explained with the sister chromatid as a temporal template for the insertion.

## Notes

### Competing Interest Statement

The authors have declared no competing interest.

