## Supplementary Tables for "Mutational signature catalogue *(MUSIC)* of DNA double-strand break repair"

### Supplementary Table legends

***Supplementary Table S1****: List of the oligos used in this study.*

***Supplementary Table S2****: List of the individual sgRNAs used in this study.*

***Supplementary Table S3****: List of the NGS details.*

***Supplementary Table S4****: List of the sgRNAs that were included in the focused candidate library.*

***Supplementary Table S5****: List of the cell lines used in this study.*

***Supplementary Table S1***

| Oligo name | Sequence | NNNNNNNN |
| --- | --- | --- |
| Fwd_mmLig4-2(JS) | CAACGTCACCACAGATCTGG | n.a. |
| Rev_mmLig4-2(JS) | CTAAGGATCTCATACCTCTTC | n.a. |
| Fwd_mmPolq-2(JS) | GCTAGCATTTTTCTGTATAGG | n.a. |
| Rev_mmPolq-2(JS) | CAAATGTCCTTGTTTCCCTCC | n.a. |
| Fwd_Wrn-2 | GAAAAATAGAACCGCTTCCTTTCC | n.a. |
| Rev_Wrn-2 | CTAAAGCGGGCCACAAGGAT | n.a. |
| Fwd_Wrn-4 | CGAACGCCACCTCTAGTAACA | n.a. |
| Rev_Wrn-4 | AGCCTAGACTTTGAACAGACCTT | n.a. |
| Fwd1_Library | TAGTGAACGGATCGGCACTG | n.a. |
| Rev1_Library | CTCCCCTACCCGGTAGAATTG | n.a. |
| Fwd2_Library | TTCGGGTTTATTACAGGGACAGC | n.a. |
| Rev2_Library | AACTTGCTATGCTGTTTCCAGC | n.a. |
| Fwd2ngs_Library | AATGATACGGCGACCACCGAGATCTACACNNNNNNNNTCGTCGGCAGCGTCAGATGTGTATAAGAGACAGTTCGGGTTTATTACAGGGACAG*C | TGGTAATT, TTCCATGG, GAGAGTCA, GCGTCTAT, GGCGATAA, GTTCAGGT |
| Rev2ngs_Library | CAAGCAGAAGACGGCATACGAGATNNNNNNNNGTGACTGGAGTTCAGACGTGTGCTCTTCCGATCTAACTTGCTATGCTGTTTCCAG*C | GAGTCATG, GCCGGAAC, GGAGGTCC, TATAAGGC, TTATCCAT, TTGCCAGA |
| Fwdngs_sgHprt-eGFPex3 | GATGTGTATAAGAGACAGGGACTGAAAGACTTGCTCGAGAT | n.a. |
| Revngs_sgHprt-eGFPex3 | CGTGTGCTCTTCCGATCTACACAGTAGCTCTTCAGTCTGATAAAA | n.a. |
| p5_general | AATGATACGGCGACCACCGAGATCTACACNNNNNNNNTCGTCGGCAGCGTCAGATGTGTATAAGAGACA*G | AAGCGTTG, AGTATAAC, ATCCGCGC, ATTACGCT, CAGAGCTT, CGCTTCAT, GAATTCGC, GCAAGCAT, GGCATCGA, GTTACCGC, CCGCCATT, CGCATATG, CTCCGGTT, GAGCTAAT, GCTAATGG, GTAAGTTG, AACGCAAT, AGATGCCA, AGCTAGAG, ATTAACTG, TAGACTGG, TCAGTCCA, TGAAGCTC, TGGTAATT, TTCCATGG, GAGAGTCA, GCGTCTAT, GGCGATAA, GTTCAGGT |
| p7_general | CAAGCAGAAGACGGCATACGAGATNNNNNNNNGTGACTGGAGTTCAGACGTGTGCTCTTCCGATC*T | ATTAACTG, ATTACGCT, CTGCCGCC, CTGCTCGT, GTTCAGGT, GTTCCTCG, AGCGTAGC, AGCCTCG, TGCCTCTT, TCCTCTAC, AGTTACGT, ATACGACG, CCAACTCA, CGTACTTC, GAGTCATG, GCCGGAAC, GGAGGTCC, TATAAGGC, TTATCCAT, TTGCCAGA |
| U6-sequencing primer | GAGGGCCTATTTCCCATGATT | n.a. |
| Fwd_sgLibrary-1 | CACCGTCCAAGGTCGGGCAGGAAGA | n.a. |
| Rev_sgLibrary-1 | AAACTCTTCCTGCCCGACCTTGGAC | n.a. |
| Fwd_sgLibrary-2 | CACCGGACCTTGGACGCGTAGGGCC | n.a. |
| Rev_sgLibrary-2 | AAACGGCCCTACGCGTCCAAGGTCC | n.a. |
| Fwd_sgLibrary-3 | CACCGTGCCCGACCTTGGACGCGTA | n.a. |
| Rev_sgLibrary-3 | AAACTACGCGTCCAAGGTCGGGCAC | n.a. |
| Fwd_sgNon-targeting(NT)-1 | CACCGCGAGGTATTCGGCTCCGCG | n.a. |
| Fwd_sgNon-targeting(NT)-2 | CACCGTGCACCGGTACGGGCGACTC | n.a. |
| Rev_sgNon-targeting(NT)-1 | AAACCGCGGAGCCGAATACCTCGC | n.a. |
| Rev_Non-targeting(NT)-2 | AAACGAGTCGCCCGTACCGGTGCAC | n.a. |
| Fwd_sgmmLig4-2(JS) | CACCGTACAACTATACCGACCAGTT | n.a. |
| Rev_sgmmLig4-2(JS) | AAACAACTGGTCGGTATAGTTGTAC | n.a. |
| Fwd_sgmmPolq-2(JS) | CACCGTGTAAGAAAGATGTTCGAA | n.a. |
| Rev_sgmmPolq-2(JS) | AAACTTCGAACATCTTTCTTACAC | n.a. |
| Fwd_sgmmWrn-2 | CACCGGTTGGGTTCCGGCAATAAA | n.a. |
| Fwd_sgmmWrn-4 | CACCGGGCCGCCCATATACAAGCC | n.a. |
| Rev_sgmmWrn-2 | AAACTTTATTGCCGGAACCCAACC | n.a. |
| Rev_sgmmWrn-4 | AAACGGCTTGTATATGGGCGGCCC | n.a. |
| Fwd_sgHprt-eGFPex3 | CACCGAGCCCCCCTTGAGCACACAG | n.a. |
| Rev_sgHprt-eGFPex3 | AAACCTGTGTGCTCAAGGGGGGCTC | n.a. |
| Fwd_sghsLig4-2(JS) | CACCGTCTGCAATAGCAGCTAGCAT | n.a. |
| Rev_sghsLig4-2(JS) | AAACATGCTAGCTGCTATTGCAGAC | n.a. |
| Fwd_sghsWrn-ex02 | CACCGTGGAAACAACTGCACAGCAG | n.a. |
| Fwd_sghsWrn-ex05 | CACCGTAGGAATTGAAGGAGATCAG | n.a. |
| Fwd_sghsWrn-ex09 | CACCGCTAACATTGAGACTGAACTG | n.a. |
| Fwd_sghsWrn-ex19 | CACCGGAAATGGCTGCAGATCCTGA | n.a. |
| Rev_sghsWrn-ex02 | AAACCTGCTGTGCAGTTGTTTCCAC | n.a. |
| Rev_sghsWrn-ex05 | AAACCTGATCTCCTTCAATTCCTAC | n.a. |
| Rev_sghsWrn-ex09 | AAACCAGTTCAGTCTCAATGTTAGC | n.a. |
| Rev_sghsWrn-ex19 | AAACTCAGGATCTGCAGCCATTTCC | n.a. |

***Supplementary Table S2***

| sgRNA name | sgRNA sequence | Fwd_cloning | Rev_cloning | Fwd_genotype | Rev_genotype | Fwd(Nested)_Genotype | Rev(Nested)_Genotype | Sangerseq_Genotype |
|---|---|---|---|---|---|---|---|---|
| sgLibrary-1 | TCCAAGGTCGGGCAGGAAGA | Fwd_sgLibrary-1 | Rev_sgLibrary-1 | Fwd1_Library | Rev1_Library | Fwd2_Library | Rev2_Library | Fwd2_Library |
| sgLibrary-2 | GACCTTGGACGCGTAGGGCC | Fwd_sgLibrary-2 | Rev_sgLibrary-2 | Fwd1_Library | Rev1_Library | Fwd2_Library | Rev2_Library | Fwd2_Library |
| sgLibrary-3 | TGCCCGACCTTGGACGCGTA | Fwd_sgLibrary-3 | Rev_sgLibrary-3 | Fwd1_Library | Rev1_Library | Fwd2_Library | Rev2_Library | Fwd2_Library |
| sgNon-targeting(NT)-1 | CGAGGTATTCGGCTCCGCG | Fwd_sgmmNT-1 | Rev_sgmmNT-1 | n.a. | n.a. | n.a. | n.a. | n.a. |
| sgNon-targeting(NT)-2 | TGCACCGGTACGGGCGACTC | Fwd_sgmmNT-2 | Rev_sgmmNT-2 | n.a. | n.a. | n.a. | n.a. | n.a. |
| sgmmLig4-2(JS) | TACAACTATACCGACCAGTT | Fwd_sgmmLig4-2(JS) | Rev_sgmmLig4-2(JS) | Fwd_mmLig4-2(JS) | Rev_mmLig4-2(JS) | n.a. | n.a. | Fwd_mmLig4-2(JS) |
| sgmmPolq-2(JS) | CACCGTGTAAGAAAGATGTTCGAA | Fwd_sgmmPolq-2(JS) | Rev_sgmmPolq-2(JS) | Fwd_mmPolq-2(JS) | Rev_mmPolq-2(JS) | n.a. | n.a. | Fwd_mmPolq-2(JS) |
| sgmmWrn-2 | GTTGGGTTCCGGCAATAAA | Fwd_sgmmWrn-2 | Rev_sgmmWrn-2 | Fwd_Wrn-2 | Rev_Wrn-2 | n.a. | n.a. | Fwd_Wrn-2 |
| sgmmWrn-4 | GGCCGCCCATATACAAGCC | Fwd_sgmmWrn-4 | Rev_sgmmWrn-4 | Fwd_Wrn-4 | Rev_Wrn-4 | n.a. | n.a. | Fwd_Wrn-4 |
| sgHprt-eGFPex3 | AGCCCCCCTTGAGCACACAG | Fwd_sgHprt-eGFPex3 | Rev_sgHprt-eGFPex3 | Fwdngs_sgHprt-eGFPex3 | Revngs_sgHprt-eGFPex3 | n.a. | n.a. | n.a. |
| sghsLig4-2(JS) | TCTGCAATAGCAGCTAGCAT | Fwd_hsLig4-2(JS) | Rev_hsLig4-2(JS) | n.a. | n.a. | n.a. | n.a. | n.a. |
| sghsWrn-ex02 | TGGAAACAACTGCACAGCAG | Fwd_sghsWrn-ex02 | Rev_sghsWrn-ex02 | n.a. | n.a. | n.a. | n.a. | n.a. |
| sghsWrn-ex05 | TAGGAATTGAAGGAGATCAG | Fwd_sghsWrn-ex05 | Rev_sghsWrn-ex05 | n.a. | n.a. | n.a. | n.a. | n.a. |
| sghsWrn-ex09 | CTAACATTGAGACTGAACTG | Fwd_sghsWrn-ex09 | Rev_sghsWrn-ex09 | n.a. | n.a. | n.a. | n.a. | n.a. |
| sghsWrn-ex19 | GAAATGGCTGCAGATCCTGA | Fwd_sghsWrn-ex19 | Rev_sghsWrn-ex19 | n.a. | n.a. | n.a. | n.a. | n.a. |

***Supplementary Table S3***

| Experiment | Related to figure | Cell line(s) | Species | Target locus | Expression vector | Condition | Replicate | Title | Design | P5 | P7 | Platform |
|---|---|---|---|---|---|---|---|---|---|---|---|---|
| Pilot MUSIC in IB10-Cas9 cells | Fig. S1 | IB10-Cas9 | *M. musculus* | sgLibrary1_pilot | pLenti-multi-Guide | sgNT-1, sgNT-2, sgLig4-2(JS), sgPolq-2(JS) | Replicate 1 | Amplicon of MUSIC_Pilot_1 | Deep sequencing of amplicon | GGCGATAA | ATGGATAA | Illumina Miseq |
| Pilot MUSIC in IB10-Cas9 cells | Fig. S1 | IB10-Cas9 | *M. musculus* | sgLibrary2_pilot | pLenti-multi-Guide | sgNT-1, sgNT-2, sgLig4-2(JS), sgPolq-2(JS) | Replicate 1 | Amplicon of MUSIC_Pilot_2 | Deep sequencing of amplicon | GGCGATAA | TCTGGCAA | Illumina Miseq |
| Pilot MUSIC in IB10-Cas9 cells | Fig. S1 | IB10-Cas9 | *M. musculus* | sgLibrary3_pilot | pLenti-multi-Guide | sgNT-1, sgNT-2, sgLig4-2(JS), sgPolq-2(JS) | Replicate 1 | Amplicon of MUSIC_Pilot_3 | Deep sequencing of amplicon | GTTCAGGT | ATGGATAA | Illumina Miseq |
| Pilot MUSIC in IB10-Cas9 cells | Fig. S1 | IB10-Cas9 | *M. musculus* | sgLibrary1_pilot_spikein | pLenti-multi-Guide | sgNT-1, sgNT-2, sgLig4-2(JS), sgPolq-2(JS) + spiked-in wildtype Yusav2 library backbone | Replicate 1 | Amplicon of MUSIC_Pilot_4 | Deep sequencing of amplicon | GTTCAGGT | TCTGGCAA | Illumina Miseq |
| Genome-wide MUSIC in IB10-Cas9 cells | Fig. 1 | IB10-Cas9 | *M. musculus* | sgLibrary-1 | pLentiGuide_eGFP | Yusa_v2 + sgLibrary-1 | Replicate 1 | Amplicon of MUSIC_target_1_rep_1 | Deep sequencing of amplicon | GAGAGTCA | GGACCTCC | Illumina NovaSeq 6000 |
| Genome-wide MUSIC in IB10-Cas9 cells | Fig. 1 | IB10-Cas9 | *M. musculus* | sgLibrary-1 | pLentiGuide_eGFP | Yusa_v2 + sgLibrary-1 | Replicate 2 | Amplicon of MUSIC_target_1_rep_2 | Deep sequencing of amplicon | GCGTCTAT | GCCTTATA | Illumina NovaSeq 6000 |
| Genome-wide MUSIC in IB10-Cas9 cells | Fig. 1 | IB10-Cas9 | *M. musculus* | sgLibrary-2 | pLentiGuide_eGFP | Yusa_v2 + sgLibrary-2 | Replicate 1 | Amplicon of MUSIC_target_2_rep_1 | Deep sequencing of amplicon | GGCGATAA | ATGGATAA | Illumina NovaSeq 6000 |
| Genome-wide MUSIC in IB10-Cas9 cells | Fig. 1 | IB10-Cas9 | *M. musculus* | sgLibrary-2 | pLentiGuide_eGFP | Yusa_v2 + sgLibrary-2 | Replicate 2 | Amplicon of MUSIC_target_2_rep_2 | Deep sequencing of amplicon | GTTCAGGT | TCTGGCAA | Illumina NovaSeq 6000 |
| Genome-wide MUSIC in IB10-Cas9 cells | Fig. 1 | IB10-Cas9 | *M. musculus* | sgLibrary-3 | pLentiGuide_eGFP | Yusa_v2 + sgLibrary-3 | Replicate 1 | Amplicon of MUSIC_target_3_rep_1 | Deep sequencing of amplicon | GAGAGTCA | TCTGGCAA | Illumina NovaSeq 6000 |
| Genome-wide MUSIC in IB10-Cas9 cells | Fig. 1 | IB10-Cas9 | *M. musculus* | sgLibrary-3 | pLentiGuide_eGFP | Yusa_v2 + sgLibrary-3 | Replicate 2 | Amplicon of MUSIC_target_3_rep_2 | Deep sequencing of amplicon | GCGTCTAT | ATGGATAA | Illumina NovaSeq 6000 |
| Genome-wide MUSIC in IB10-Cas9 cells | Fig. 1 | IB10-Cas9 | *M. musculus* | sgLibrary-1 | pLentiGuide_eGFP | Yusa_v2 + sgLibrary-1 + NU7441 (2 µM) | Replicate 1 | Amplicon of MUSIC_target_1_rep_1_NU7441 | Deep sequencing of amplicon | GGCGATAA | GCCTTATA | Illumina NovaSeq 6000 |
| Genome-wide MUSIC in IB10-Cas9 cells | Fig. 1 | IB10-Cas9 | *M. musculus* | sgLibrary-1 | pLentiGuide_eGFP | Yusa_v2 + sgLibrary-1 + ART588 (10 µM) | Replicate 1 | Amplicon of MUSIC_target_1_rep_1_ART558 | Deep sequencing of amplicon | GTTCAGGT | GGACCTCC | Illumina NovaSeq 6000 |
| Focused candidate MUSIC in IB10-Cas9 cells | Fig. 2-7; Fig. S2-3 | IB10-Cas9 | *M. musculus* | sgLibrary-1 | pLentiGuide_eGFP | FocusedCandidateLibrary + sgLibrary-1 | Replicate 1 | Amplicon of MUSIC_candidate_rep_1 | Deep sequencing of amplicon | ACCTGAAC | GCCTTATA | Illumina NovaSeq 6000 |
| Focused candidate MUSIC in IB10-Cas9 cells | Fig. 2-7; Fig. S2-3 | IB10-Cas9 | *M. musculus* | sgLibrary-1 | pLentiGuide_eGFP | FocusedCandidateLibrary + sgLibrary-1 | Replicate 2 | Amplicon of MUSIC_candidate_rep_2 | Deep sequencing of amplicon | ACCTGAAC | ATGGATAA | Illumina NovaSeq 6000 |
| Focused candidate MUSIC in IB10-Cas9 cells | Fig. 2-7; Fig. S2-3 | IB10-Cas9 | *M. musculus* | sgLibrary-1 | pLentiGuide_eGFP | FocusedCandidateLibrary + sgLibrary-1 | Replicate 3 | Amplicon of MUSIC_candidate_rep_3 | Deep sequencing of amplicon | ACCTGAAC | TCTGGCAA | Illumina NovaSeq 6000 |
| Validation sgLibrary-1 in mES IB10-Cas9 Wrn-knockout clones | Fig. S4 | IB10-Cas9 | *M. musculus* | sgLibrary-1 | pLentiGuide_eGFP | IB10Cas9_parental_(-) + sgNT-1, sgNT-2, sghsLig4-2 in Addgene#67974 + sgLibrary-1 | Replicate 1-3 | Amplicon of MUSIC_validation_wt_lib_1 | Deep sequencing of amplicon | AATTACCA | CATGACTC | Illumina NovaSeq 6000 |
| Validation sgLibrary-1 in mES IB10-Cas9 Wrn-knockout clones | Fig. S4 | IB10-Cas9 Wrn-/- sg-2cC3 | *M. musculus* | sgLibrary-1 | pLentiGuide_eGFP | IB10Cas9_Wrn-2cC3_(-) + sgNT-1, sgNT-2, sghsLig4-2 in Addgene#67974 + sgLibrary-1 | Replicate 1-3 | Amplicon of MUSIC_validation_WRN2_lib_1 | Deep sequencing of amplicon | AATTACCA | GTTCCGGC | Illumina NovaSeq 6000 |
| Validation sgLibrary-1 in mES IB10-Cas9 Wrn-knockout clones | Fig. S4 | IB10-Cas9 Wrn-/- sg-4cD11 | *M. musculus* | sgLibrary-1 | pLentiGuide_eGFP | IB10Cas9_Wrn-4cD11_(-) + sgNT-1, sgNT-2, sghsLig4-2 in Addgene#67974 + sgLibrary-1 | Replicate 1-3 | Amplicon of MUSIC_validation_WRN4_lib_1 | Deep sequencing of amplicon | AATTACCA | TCTGGCAA | Illumina NovaSeq 6000 |
| Validation sgLibrary-3 in mES IB10-Cas9 Wrn-knockout clones | Fig. S4 | IB10-Cas9 | *M. musculus* | sgLibrary-3 | pLentiGuide_eGFP | IB10Cas9_parental_(-) + sgNT-1, sgNT-2, sghsLig4-2 in Addgene#67974 + sgLibrary-3 | Replicate 1-3 | Amplicon of MUSIC_validation_wt_lib_3 | Deep sequencing of amplicon | CCATGGAA | GCCTTATA | Illumina NovaSeq 6000 |
| Validation sgLibrary-3 in mES IB10-Cas9 Wrn-knockout clones | Fig. S4 | IB10-Cas9 Wrn-/- sg-2cC3 | *M. musculus* | sgLibrary-3 | pLentiGuide_eGFP | IB10Cas9_Wrn-2cC3_(-) + sgNT-1, sgNT-2, sghsLig4-2 in Addgene#67974 + sgLibrary-3 | Replicate 1-3 | Amplicon of MUSIC_validation_WRN2_lib_3 | Deep sequencing of amplicon | CCATGGAA | ATGGATAA | Illumina NovaSeq 6000 |
| Validation sgLibrary-3 in mES IB10-Cas9 Wrn-knockout clones | Fig. S4 | IB10-Cas9 Wrn-/- sg-4cD11 | *M. musculus* | sgLibrary-3 | pLentiGuide_eGFP | IB10Cas9_Wrn-4cD11_(-) + sgNT-1, sgNT-2, sghsLig4-2 in Addgene#67974 + sgLibrary-3 | Replicate 1-3 | Amplicon of MUSIC_validation_WRN4_lib_3 | Deep sequencing of amplicon | TGACTCTC | GGACCTCC | Illumina NovaSeq 6000 |
| Validation Hprt-locus in mES-Hprt-eGFP Wrn-knockout clones | Fig. S5 and S7 | MT01 Hprt-eGFP parental c1 | *M. musculus* | sgHprt-eGFPex3 | Plasmid pU6-(BbsI)_CBh-Cas9-T2A-mCherry | MT01 parental + sgHprt-eGFPex3 | Replicate 1 | Amplicon of HPRT-eGFPex3_MT01_rep_1 | Deep sequencing of amplicon | AGTATAAC | CAGTTAAT | Illumina Miseq |
| Validation Hprt-locus in mES-Hprt-eGFP Wrn-knockout clones | Fig. S5 and S7 | MT02 Hprt-eGFP parental c2 | *M. musculus* | sgHprt-eGFPex3 | Plasmid pU6-(BbsI)_CBh-Cas9-T2A-mCherry | MT02 parental+ sgHprt-eGFPex3 | Replicate 1 | Amplicon of HPRT-eGFPex3_MT02_rep_1 | Deep sequencing of amplicon | AGTATAAC | AGCGTAAT | Illumina Miseq |
| Validation Hprt-locus in mES-Hprt-eGFP Wrn-knockout clones | Fig. S5 and S7 | MT03 Hprt-eGFP;Polq-/- c1 | *M. musculus* | sgHprt-eGFPex3 | Plasmid pU6-(BbsI)_CBh-Cas9-T2A-mCherry | MT03_Polq-ko + sgHprt-eGFPex3 | Replicate 1 | Amplicon of HPRT-eGFPex3_MT03_rep_1 | Deep sequencing of amplicon | AGTATAAC | GGCGGCAG | Illumina Miseq |
| Validation Hprt-locus in mES-Hprt-eGFP Wrn-knockout clones | Fig. S5 and S7 | MT04 Hprt-eGFP;Polq-/- c2 | *M. musculus* | sgHprt-eGFPex3 | Plasmid pU6-(BbsI)_CBh-Cas9-T2A-mCherry | MT04_Polq-ko + sgHprt-eGFPex3 | Replicate 1 | Amplicon of HPRT-eGFPex3_MT04_rep_1 | Deep sequencing of amplicon | AGTATAAC | ACGAGCAG | Illumina Miseq |
| Validation Hprt-locus in mES-Hprt-eGFP Wrn-knockout clones | Fig. S5 and S7 | MT01 Hprt-eGFP;Wrn-/- sg-4cH9 | *M. musculus* | sgHprt-eGFPex3 | Plasmid pU6-(BbsI)_CBh-Cas9-T2A-mCherry | MT01_Wrn-ko_cH9 + sgHprt-eGFPex3 | Replicate 1 | Amplicon of HPRT-eGFPex3_MT01_WRN_H9_rep_1 | Deep sequencing of amplicon | AGTATAAC | ACCTGAAC | Illumina Miseq |
| Validation Hprt-locus in mES-Hprt-eGFP Wrn-knockout clones | Fig. S5 and S7 | MT02 Hprt-eGFP;Wrn-/- sg-4cG11 | *M. musculus* | sgHprt-eGFPex3 | Plasmid pU6-(BbsI)_CBh-Cas9-T2A-mCherry | MT02_Wrn-ko_cG11 + sgHprt-eGFPex3 | Replicate 1 | Amplicon of HPRT-eGFPex3_MT01_WRN_G11_rep_1 | Deep sequencing of amplicon | AGTATAAC | CGAGGAAC | Illumina Miseq |
| Validation Hprt-locus in mES-Hprt-eGFP Wrn-knockout clones | Fig. S5 and S7 | MT01 Hprt-eGFP parental c1 | *M. musculus* | sgHprt-eGFPex3 | Plasmid pU6-(BbsI)_CBh-Cas9-T2A-mCherry | MT01 parental + sgHprt-eGFPex3 | Replicate 2 | Amplicon of HPRT-eGFPex3_MT01_rep_2 | Deep sequencing of amplicon | AGTATAAC | ACGTAACT | Illumina Miseq |
| Validation Hprt-locus in mES-Hprt-eGFP Wrn-knockout clones | Fig. S5 and S7 | MT02 Hprt-eGFP parental c2 | *M. musculus* | sgHprt-eGFPex3 | Plasmid pU6-(BbsI)_CBh-Cas9-T2A-mCherry | MT02 parental + sgHprt-eGFPex3 | Replicate 2 | Amplicon of HPRT-eGFPex3_MT02_rep_2 | Deep sequencing of amplicon | AGTATAAC | CGTCGTAT | Illumina Miseq |
| Validation Hprt-locus in mES-Hprt-eGFP Wrn-knockout clones | Fig. S5 and S7 | MT03 Hprt-eGFP;Polq-/- c1 | *M. musculus* | sgHprt-eGFPex3 | Plasmid pU6-(BbsI)_CBh-Cas9-T2A-mCherry | MT03_Polq-ko + sgHprt-eGFPex3 | Replicate 2 | Amplicon of HPRT-eGFPex3_MT03_rep_2 | Deep sequencing of amplicon | AGTATAAC | TGAGTTGG | Illumina Miseq |
| Validation Hprt-locus in mES-Hprt-eGFP Wrn-knockout clones | Fig. S5 and S7 | MT04 Hprt-eGFP;Polq-/- c2 | *M. musculus* | sgHprt-eGFPex3 | Plasmid pU6-(BbsI)_CBh-Cas9-T2A-mCherry | MT04_Polq-ko + sgHprt-eGFPex3 | Replicate 2 | Amplicon of HPRT-eGFPex3_MT04_rep_2 | Deep sequencing of amplicon | AGTATAAC | GAAGTACG | Illumina Miseq |
| Validation Hprt-locus in mES-Hprt-eGFP Wrn-knockout clones | Fig. S5 and S7 | MT01 Hprt-eGFP;Wrn-/- sg-4cH9 | *M. musculus* | sgHprt-eGFPex3 | Plasmid pU6-(BbsI)_CBh-Cas9-T2A-mCherry | MT01_Wrn-ko_cH9 + sgHprt-eGFPex3 | Replicate 2 | Amplicon of HPRT-eGFPex3_MT01_WRN_H9_rep_2 | Deep sequencing of amplicon | AGTATAAC | CATGACTC | Illumina Miseq |
| Validation Hprt-locus in mES-Hprt-eGFP Wrn-knockout clones | Fig. S5 and S7 | MT02 Hprt-eGFP;Wrn-/- sg-4cG11 | *M. musculus* | sgHprt-eGFPex3 | Plasmid pU6-(BbsI)_CBh-Cas9-T2A-mCherry | MT02_Wrn-ko_cG11 + sgHprt-eGFPex3 | Replicate 2 | Amplicon of HPRT-eGFPex3_MT01_WRN_G11_rep_2 | Deep sequencing of amplicon | AGTATAAC | GTTCCGGC | Illumina Miseq |
| Validation sgLibrary-1 in Hela-Cas9 polyclonal *WRN* -targeted cells | Fig. S6 | Hela-Cas9 | *H. sapiens* | sgLibrary-1 | pLentiGuide_eGFP | sghsWrn-ex02, sghsWrn-ex05, sghsWrn-ex09, sghsWrn-ex19 in Addgene#67974 + sgLibrary-1 | Replicate 1 | Amplicon of Hela-Cas9_sgLibrary-1_Wrn_repl_1 | Deep sequencing of amplicon | GGACCTCC | CCATGGAA | Illumina NovaSeq 6000 |
| Validation sgLibrary-1 in Hela-Cas9 polyclonal Non-targeted cells | Fig. S6 | Hela-Cas9 | *H. sapiens* | sgLibrary-1 | pLentiGuide_eGFP | sgNT-1, sgNT-2 in Addgene#67974 + sgLibrary-1 | Replicate 1 | Amplicon of Hela-Cas9_sgLibrary-1_NT_repl_1 | Deep sequencing of amplicon | TCTGGCAA | CCATGGAA | Illumina NovaSeq 6000 |
| Validation sgLibrary-1 in U2OS-Cas9 polyclonal *WRN* -targeted cells | Fig. S6 | U2OS-Cas9 | *H. sapiens* | sgLibrary-1 | pLentiGuide_eGFP | sghsWrn-ex02, sghsWrn-ex05, sghsWrn-ex09, sghsWrn-ex19 in Addgene#67974 + sgLibrary-1 | Replicate 1 | Amplicon of U2OS-Cas9_sgLibrary-1_Wrn_repl_1 | Deep sequencing of amplicon | CATGACTC | TGACTCTC | Illumina NovaSeq 6000 |
| Validation sgLibrary-1 in U2OS-Cas9 polyclonal Non-targeted cells | Fig. S6 | U2OS-Cas9 | *H. sapiens* | sgLibrary-1 | pLentiGuide_eGFP | sgNT-1, sgNT-2 in Addgene#67974 + sgLibrary-1 | Replicate 1 | Amplicon of U2OS-Cas9_sgLibrary-1_NT_repl_1 | Deep sequencing of amplicon | GTTCCGGC | TGACTCTC | Illumina NovaSeq 6000 |
| Validation sgLibrary-1 in HEK-Cas9 polyclonal *WRN* -targeted cells | Fig. S6 | HEK-Cas9 | *H. sapiens* | sgLibrary-1 | pLentiGuide_eGFP | sghsWrn-ex02, sghsWrn-ex05, sghsWrn-ex09, sghsWrn-ex19 in Addgene#67974 + sgLibrary-1 | Replicate 1 | Amplicon of HEK-Cas9_sgLibrary-1_Wrn_repl_1 | Deep sequencing of amplicon | GGACCTCC | TGACTCTC | Illumina NovaSeq 6000 |
| Validation sgLibrary-1 in HEK-Cas9 polyclonal Non-targeted cells | Fig. S6 | HEK-Cas9 | *H. sapiens* | sgLibrary-1 | pLentiGuide_eGFP | sgNT-1, sgNT-2 in Addgene#67974 + sgLibrary-1 | Replicate 1 | Amplicon of HEK-Cas9_sgLibrary-1_NT_repl_1 | Deep sequencing of amplicon | GCCTTATA | TGACTCTC | Illumina NovaSeq 6000 |
| Validation sgLibrary-1 in RPE1hTert-Cas9 polyclonal *WRN* -targeted cells | Fig. S6 | RPE1hTert-Cas9 | *H. sapiens* | sgLibrary-1 | pLentiGuide_eGFP | sghsWrn-ex02, sghsWrn-ex05, sghsWrn-ex09, sghsWrn-ex19 in Addgene#67974 + sgLibrary-1 | Replicate 1 | Amplicon of RPE1hTert-Cas9_sgLibrary-1_Wrn_repl_1 | Deep sequencing of amplicon | ATGGATAA | TGACTCTC | Illumina NovaSeq 6000 |
| Validation sgLibrary-1 in RPE1hTert-Cas9 polyclonal Non-targeted cells | Fig. S6 | RPE1hTert-Cas9 | *H. sapiens* | sgLibrary-1 | pLentiGuide_eGFP | sgNT-1, sgNT-2 in Addgene#67974 + sgLibrary-1 | Replicate 1 | Amplicon of RPE1hTert-Cas9_sgLibrary-1_NT_repl_1 | Deep sequencing of amplicon | TCTGGCAA | TGACTCTC | Illumina NovaSeq 6000 |
| Validation sgLibrary-1 in Hela-Cas9 polyclonal *WRN* -targeted cells | Fig. S6 | Hela-Cas9 | *H. sapiens* | sgLibrary-1 | pLentiGuide_eGFP | sghsWrn-ex02, sghsWrn-ex05, sghsWrn-ex09, sghsWrn-ex19 in Addgene#67974 + sgLibrary-1 | Replicate 2 | Amplicon of Hela-Cas9_sgLibrary-1_Wrn_repl_2 | Deep sequencing of amplicon | CATGACTC | ATAGACGC | Illumina NovaSeq 6000 |
| Validation sgLibrary-1 in Hela-Cas9 polyclonal Non-targeted cells | Fig. S6 | Hela-Cas9 | *H. sapiens* | sgLibrary-1 | pLentiGuide_eGFP | sgNT-1, sgNT-2 in Addgene#67974 + sgLibrary-1 | Replicate 2 | Amplicon of Hela-Cas9_sgLibrary-1_NT_repl_2 | Deep sequencing of amplicon | GTTCCGGC | ATAGACGC | Illumina NovaSeq 6000 |
| Validation sgLibrary-1 in U2OS-Cas9 polyclonal *WRN* -targeted cells | Fig. S6 | U2OS-Cas9 | *H. sapiens* | sgLibrary-1 | pLentiGuide_eGFP | sghsWrn-ex02, sghsWrn-ex05, sghsWrn-ex09, sghsWrn-ex19 in Addgene#67974 + sgLibrary-1 | Replicate 2 | Amplicon of U2OS-Cas9_sgLibrary-1_Wrn_repl_2 | Deep sequencing of amplicon | GGACCTCC | ATAGACGC | Illumina NovaSeq 6000 |
| Validation sgLibrary-1 in U2OS-Cas9 polyclonal Non-targeted cells | Fig. S6 | U2OS-Cas9 | *H. sapiens* | sgLibrary-1 | pLentiGuide_eGFP | sgNT-1, sgNT-2 in Addgene#67974 + sgLibrary-1 | Replicate 2 | Amplicon of U2OS-Cas9_sgLibrary-1_NT_repl_2 | Deep sequencing of amplicon | GCCTTATA | ATAGACGC | Illumina NovaSeq 6000 |
| Validation sgLibrary-1 in HEK-Cas9 polyclonal *WRN* -targeted cells | Fig. S6 | HEK-Cas9 | *H. sapiens* | sgLibrary-1 | pLentiGuide_eGFP | sghsWrn-ex02, sghsWrn-ex05, sghsWrn-ex09, sghsWrn-ex19 in Addgene#67974 + sgLibrary-1 | Replicate 2 | Amplicon of HEK-Cas9_sgLibrary-1_Wrn_repl_2 | Deep sequencing of amplicon | ATGGATAA | ATAGACGC | Illumina NovaSeq 6000 |
| Validation sgLibrary-1 in HEK-Cas9 polyclonal Non-targeted cells | Fig. S6 | HEK-Cas9 | *H. sapiens* | sgLibrary-1 | pLentiGuide_eGFP | sgNT-1, sgNT-2 in Addgene#67974 + sgLibrary-1 | Replicate 2 | Amplicon of HEK-Cas9_sgLibrary-1_NT_repl_2 | Deep sequencing of amplicon | TCTGGCAA | ATAGACGC | Illumina NovaSeq 6000 |
| Validation sgLibrary-1 in RPE1hTert-Cas9 polyclonal *WRN* -targeted cells | Fig. S6 | RPE1hTert-Cas9 | *H. sapiens* | sgLibrary-1 | pLentiGuide_eGFP | sghsWrn-ex02, sghsWrn-ex05, sghsWrn-ex09, sghsWrn-ex19 in Addgene#67974 + sgLibrary-1 | Replicate 2 | Amplicon of RPE1hTert-Cas9_sgLibrary-1_Wrn_repl_2 | Deep sequencing of amplicon | CATGACTC | TTATCGCC | Illumina NovaSeq 6000 |
| Validation sgLibrary-1 in RPE1hTert-Cas9 polyclonal Non-targeted cells | Fig. S6 | RPE1hTert-Cas9 | *H. sapiens* | sgLibrary-1 | pLentiGuide_eGFP | sgNT-1, sgNT-2 in Addgene#67974 + sgLibrary-1 | Replicate 2 | Amplicon of RPE1hTert-Cas9_sgLibrary-1_NT_repl_2 | Deep sequencing of amplicon | GTTCCGGC | TTATCGCC | Illumina NovaSeq 6000 |

*Supplementary Table S4*

| Gene | Barcode | Sequence | Oligo |
|---|---|---|---|
| 1700025G04Rik | 1700025G04Rik-1 | GGGTTGATGAACTAATCCA | tatcttgtggaaaggacgaaacaccGGGGTTGATGAACTAATCCAgtttaagagctatgctggaaacagc |
| 1700025G04Rik | 1700025G04Rik-2 | AAACTAATAAAGAAATCCC | tatcttgtggaaaggacgaaacaccGAAACTAATAAAGAAATCCCgtttaagagctatgctggaaacagc |
| 1700025G04Rik | 1700025G04Rik-4 | GGTCAAATACATGAAAAAT | tatcttgtggaaaggacgaaacaccGGGTCAAATACATGAAAAATgtttaagagctatgctggaaacagc |
| 1700123O20Rik | 1700123O20Rik-1 | GGGAAGTGATGTAAGACAT | tatcttgtggaaaggacgaaacaccGGGGAAGTGATGTAAGACATgtttaagagctatgctggaaacagc |
| 1700123O20Rik | 1700123O20Rik-2 | CGCTGGGGACCTGACCAAC | tatcttgtggaaaggacgaaacaccGCGCTGGGGACCTGACCAACgtttaagagctatgctggaaacagc |
| 1700123O20Rik | 1700123O20Rik-5 | GCTGACACTCAAAGATACA | tatcttgtggaaaggacgaaacaccGGCTGACACTCAAAGATACAgtttaagagctatgctggaaacagc |
| 1810037I17Rik | 1810037I17Rik-1 | GGAAGGAGGAACCAAGATG | tatcttgtggaaaggacgaaacaccGGGAAGGAGGAACCAAGATGgtttaagagctatgctggaaacagc |
| 1810037I17Rik | 1810037I17Rik-2 | AAGATGAGGCACCTCAGTC | tatcttgtggaaaggacgaaacaccGAAGATGAGGCACCTCAGTCgtttaagagctatgctggaaacagc |
| 1810037I17Rik | 1810037I17Rik-3 | TTCGGAAGCCCGTTCATCC | tatcttgtggaaaggacgaaacaccGTTCGGAAGCCCGTTCATCCgtttaagagctatgctggaaacagc |
| 2610507B11Rik | 2610507B11Rik-1 | GAAGCTTCGAAGCGGCAGT | tatcttgtggaaaggacgaaacaccGGAAGCTTCGAAGCGGCAGTgtttaagagctatgctggaaacagc |
| 2610507B11Rik | 2610507B11Rik-3 | ATAGAGACGACGAGAGCGC | tatcttgtggaaaggacgaaacaccGATAGAGACGACGAGAGCGCgtttaagagctatgctggaaacagc |
| 2610507B11Rik | 2610507B11Rik-5 | ACGTCGACGGGAGCATGGC | tatcttgtggaaaggacgaaacaccGACGTCGACGGGAGCATGGCgtttaagagctatgctggaaacagc |
| 4921524J17Rik | 4921524J17Rik-2 | CATACAGAAAACAAGCACG | tatcttgtggaaaggacgaaacaccGCATACAGAAAACAAGCACGgtttaagagctatgctggaaacagc |
| 4921524J17Rik | 4921524J17Rik-3 | TTTCACAATTACCTGTAGA | tatcttgtggaaaggacgaaacaccGTTTCACAATTACCTGTAGAgtttaagagctatgctggaaacagc |
| 4921524J17Rik | 4921524J17Rik-5 | CTAATAAATATGTAACCAC | tatcttgtggaaaggacgaaacaccGCTAATAAATATGTAACCACgtttaagagctatgctggaaacagc |
| 4930407I10Rik | 4930407I10Rik-1 | AGTCCTCTTAACCAACAAT | tatcttgtggaaaggacgaaacaccGAGTCCTCTTAACCAACAATgtttaagagctatgctggaaacagc |
| 4930407I10Rik | 4930407I10Rik-4 | AATGATTCACGGACCAAAA | tatcttgtggaaaggacgaaacaccGAATGATTCACGGACCAAAAgtttaagagctatgctggaaacagc |
| 4930407I10Rik | 4930407I10Rik-5 | GTGATATGACCCACGGGAC | tatcttgtggaaaggacgaaacaccGGTGATATGACCCACGGGACgtttaagagctatgctggaaacagc |
| 4930503E14Rik | 4930503E14Rik-1 | TCATGTTGTGATTTGATGA | tatcttgtggaaaggacgaaacaccGTCATGTTGTGATTTGATGAgtttaagagctatgctggaaacagc |
| 4930503E14Rik | 4930503E14Rik-3 | CAGAAAGATAAATGCTGAG | tatcttgtggaaaggacgaaacaccGCAGAAAGATAAATGCTGAGgtttaagagctatgctggaaacagc |
| 4930503E14Rik | 4930503E14Rik-4 | AATGAAATCATTCTGGGAA | tatcttgtggaaaggacgaaacaccGAATGAAATCATTCTGGGAAgtttaagagctatgctggaaacagc |
| 4931406C07Rik | 4931406C07Rik-1 | TTGAACAATTGGCATGAAC | tatcttgtggaaaggacgaaacaccGTTGAACAATTGGCATGAACgtttaagagctatgctggaaacagc |
| 4931406C07Rik | 4931406C07Rik-4 | TTACAAGAGGCAATAAGTA | tatcttgtggaaaggacgaaacaccGTTACAAGAGGCAATAAGTAgtttaagagctatgctggaaacagc |
| 4931406C07Rik | 4931406C07Rik-5 | ATGGCTCCTTTGTTAAATC | tatcttgtggaaaggacgaaacaccGATGGCTCCTTTGTTAAATCgtttaagagctatgctggaaacagc |
| 9530002B09Rik | 9530002B09Rik-1 | AGATCTACAGCTAGAATCA | tatcttgtggaaaggacgaaacaccGAGATCTACAGCTAGAATCAgtttaagagctatgctggaaacagc |
| 9530002B09Rik | 9530002B09Rik-4 | CTTCTCAGTCTCAACAGAT | tatcttgtggaaaggacgaaacaccGCTTCTCAGTCTCAACAGATgtttaagagctatgctggaaacagc |
| 9530002B09Rik | 9530002B09Rik-5 | GCATGTGAGCTGACATACA | tatcttgtggaaaggacgaaacaccGGCATGTGAGCTGACATACAgtttaagagctatgctggaaacagc |
| A630023A22Rik | A630023A22Rik-2 | TGGTGGTGTGATGTCATAA | tatcttgtggaaaggacgaaacaccGTGGTGGTGTGATGTCATAAgtttaagagctatgctggaaacagc |
| A630023A22Rik | A630023A22Rik-4 | AGTGCCGAACAAGGCATGG | tatcttgtggaaaggacgaaacaccGAGTGCCGAACAAGGCATGGgtttaagagctatgctggaaacagc |
| A630023A22Rik | A630023A22Rik-5 | GGCTCCCACCTTTAAAAGT | tatcttgtggaaaggacgaaacaccGGGCTCCCACCTTTAAAAGTgtttaagagctatgctggaaacagc |
| AA467197 | AA467197-1 | GCATACAAAGCGAAAGATG | tatcttgtggaaaggacgaaacaccGGCATACAAAGCGAAAGATGgtttaagagctatgctggaaacagc |
| AA467197 | AA467197-2 | GCGTTGAAAAAAACCGACG | tatcttgtggaaaggacgaaacaccGGCGTTGAAAAAAACCGACGgtttaagagctatgctggaaacagc |
| AA467197 | AA467197-3 | GTTAGTACTTACACCACGT | tatcttgtggaaaggacgaaacaccGGTTAGTACTTACACCACGTgtttaagagctatgctggaaacagc |
| Aaas | Aaas-1 | TTGGCTTGGGCCCCCAATG | tatcttgtggaaaggacgaaacaccGTTGGCTTGGGCCCCCAATGgtttaagagctatgctggaaacagc |
| Aaas | Aaas-2 | ATGACTCACCTCGTAGATA | tatcttgtggaaaggacgaaacaccGATGACTCACCTCGTAGATAgtttaagagctatgctggaaacagc |
| Aaas | Aaas-3 | CAGCGAAATGTAGCAGCCT | tatcttgtggaaaggacgaaacaccGCAGCGAAATGTAGCAGCCTgtttaagagctatgctggaaacagc |
| AAdacl4fm3 | AAdacl4fm3-1 | TGAAGAACCTTCAAACCTA | tatcttgtggaaaggacgaaacaccGTGAAGAACCTTCAAACCTAgtttaagagctatgctggaaacagc |
| AAdacl4fm3 | AAdacl4fm3-2 | GGCAATCCAGGTAAACAAC | tatcttgtggaaaggacgaaacaccGGGCAATCCAGGTAAACAACgtttaagagctatgctggaaacagc |
| AAdacl4fm3 | AAdacl4fm3-5 | AACTGTAAAATTGCAGGCA | tatcttgtggaaaggacgaaacaccGAACTGTAAAATTGCAGGCAgtttaagagctatgctggaaacagc |
| Abraxas1 | Abraxas1-2 | CCATTGGTGGTTACCAATC | tatcttgtggaaaggacgaaacaccGCCATTGGTGGTTACCAATCgtttaagagctatgctggaaacagc |
| Abraxas1 | Abraxas1-3 | CGCCTTATATAAACCTCAA | tatcttgtggaaaggacgaaacaccGCGCCTTATATAAACCTCAAgtttaagagctatgctggaaacagc |
| Abraxas1 | Abraxas1-5 | CCTAAAAAGCCGGTAGCAT | tatcttgtggaaaggacgaaacaccGCCTAAAAAGCCGGTAGCATgtttaagagctatgctggaaacagc |
| Acadl | Acadl-2 | GAACGAACGCCAAAAGATC | tatcttgtggaaaggacgaaacaccGGAACGAACGCCAAAAGATCgtttaagagctatgctggaaacagc |
| Acadl | Acadl-3 | TCATCCCCCAGATGACGGC | tatcttgtggaaaggacgaaacaccGTCATCCCCCAGATGACGGCgtttaagagctatgctggaaacagc |
| Acadl | Acadl-5 | ATTAACTGACATTGGAATC | tatcttgtggaaaggacgaaacaccGATTAACTGACATTGGAATCgtttaagagctatgctggaaacagc |
| Acbd7 | Acbd7-3 | CCAGCAATGTTAGATCTAA | tatcttgtggaaaggacgaaacaccGCCAGCAATGTTAGATCTAAgtttaagagctatgctggaaacagc |
| Acbd7 | Acbd7-4 | CTAAAGGGTAAGGCCAAAT | tatcttgtggaaaggacgaaacaccGCTAAAGGGTAAGGCCAAATgtttaagagctatgctggaaacagc |
| Acbd7 | Acbd7-5 | AAGCTTGGAACCTCCAAAA | tatcttgtggaaaggacgaaacaccGAAGCTTGGAACCTCCAAAAgtttaagagctatgctggaaacagc |
| Acot9 | Acot9-2 | AGGCGTATCCACAGTCTGG | tatcttgtggaaaggacgaaacaccGAGGCGTATCCACAGTCTGGgtttaagagctatgctggaaacagc |
| Acot9 | Acot9-4 | GGACTGTAACTATTGATAA | tatcttgtggaaaggacgaaacaccGGGACTGTAACTATTGATAAgtttaagagctatgctggaaacagc |
| Acot9 | Acot9-5 | GCTCGAGATTCTGAAAATA | tatcttgtggaaaggacgaaacaccGGCTCGAGATTCTGAAAATAgtttaagagctatgctggaaacagc |
| Actl6a | Actl6a-2 | GATGACGGAAGTACAATGA | tatcttgtggaaaggacgaaacaccGGATGACGGAAGTACAATGAgtttaagagctatgctggaaacagc |
| Actl6a | Actl6a-4 | TCGTGGACTGGAATCGCAG | tatcttgtggaaaggacgaaacaccGTCGTGGACTGGAATCGCAGgtttaagagctatgctggaaacagc |
| Actl6a | Actl6a-5 | TTGTGAAATCCCCTCTGGC | tatcttgtggaaaggacgaaacaccGTTGTGAAATCCCCTCTGGCgtttaagagctatgctggaaacagc |
| Actr1b | Actr1b-2 | GTTTTAGATTCGGGGGACG | tatcttgtggaaaggacgaaacaccGGTTTTAGATTCGGGGGACGgtttaagagctatgctggaaacagc |
| Actr1b | Actr1b-4 | TGTTCCATGGGGTATCGGA | tatcttgtggaaaggacgaaacaccGTGTTCCATGGGGTATCGGAgtttaagagctatgctggaaacagc |
| Actr1b | Actr1b-5 | TGCTTTCGGTCCAATGAAC | tatcttgtggaaaggacgaaacaccGTGCTTTCGGTCCAATGAACgtttaagagctatgctggaaacagc |
| Actr6 | Actr6-2 | TAAGGAACAATGTGCGTGA | tatcttgtggaaaggacgaaacaccGTAAGGAACAATGTGCGTGAgtttaagagctatgctggaaacagc |
| Actr6 | Actr6-3 | GAACTGATATTCTTCGAAT | tatcttgtggaaaggacgaaacaccGGAACTGATATTCTTCGAATgtttaagagctatgctggaaacagc |
| Actr6 | Actr6-5 | GATTACCTTTTGAAAAGGA | tatcttgtggaaaggacgaaacaccGGATTACCTTTTGAAAAGGAgtttaagagctatgctggaaacagc |
| Adam34l | Adam34l-2 | GCACTAAAAACAATAACAC | tatcttgtggaaaggacgaaacaccGGCACTAAAAACAATAACACgtttaagagctatgctggaaacagc |
| Adam34l | Adam34l-3 | TTGGAGAAATATATGCATA | tatcttgtggaaaggacgaaacaccGTTGGAGAAATATATGCATAgtttaagagctatgctggaaacagc |
| Adam34l | Adam34l-5 | ATAATCACTATGAAACCCA | tatcttgtggaaaggacgaaacaccGATAATCACTATGAAACCCAgtttaagagctatgctggaaacagc |
| Adam6a | Adam6a-3 | CTCATGACTGTGTATACCG | tatcttgtggaaaggacgaaacaccGCTCATGACTGTGTATACCGgtttaagagctatgctggaaacagc |
| Adam6a | Adam6a-4 | TAATTAACAAGAATTCGCC | tatcttgtggaaaggacgaaacaccGTAATTAACAAGAATTCGCCgtttaagagctatgctggaaacagc |
| Adam6a | Adam6a-5 | TAATTTGTCTTGGTCAACA | tatcttgtggaaaggacgaaacaccGTAATTTGTCTTGGTCAACAgtttaagagctatgctggaaacagc |
| Adam6b | Adam6b-1 | CTGTATATAATTCTGCTGC | tatcttgtggaaaggacgaaacaccGCTGTATATAATTCTGCTGCgtttaagagctatgctggaaacagc |
| Adam6b | Adam6b-3 | AATTATAAGCCGACCCTCC | tatcttgtggaaaggacgaaacaccGAATTATAAGCCGACCCTCCgtttaagagctatgctggaaacagc |
| Adam6b | Adam6b-4 | ATAAATTTAAGCCTGATGC | tatcttgtggaaaggacgaaacaccGATAAATTTAAGCCTGATGCgtttaagagctatgctggaaacagc |
| Adap1 | Adap1-1 | TCCCCCCAGGCTACCGTGA | tatcttgtggaaaggacgaaacaccGTCCCCCCAGGCTACCGTGAgtttaagagctatgctggaaacagc |
| Adap1 | Adap1-3 | CCTGACTCATACCTCAACT | tatcttgtggaaaggacgaaacaccGCCTGACTCATACCTCAACTgtttaagagctatgctggaaacagc |
| Adap1 | Adap1-5 | GACTGGGCGTCCTACACCC | tatcttgtggaaaggacgaaacaccGGACTGGGCGTCCTACACCCgtttaagagctatgctggaaacagc |
| Ahctf1 | Ahctf1-2 | GACCTACGTGGATGCTAGC | tatcttgtggaaaggacgaaacaccGGACCTACGTGGATGCTAGCgtttaagagctatgctggaaacagc |

| | | | |
|---|---|---|---|
| Ahctf1 | Ahctf1-3 | ATCAATCATCAAGCAGTCG | tatcttgtggaaaggacgaaacaccGATCAATCATCAAGCAGTCGgtttaagagctatgctggaaacagc |
| Ahctf1 | Ahctf1-4 | GGCCACGCCAAAAAGCCAC | tatcttgtggaaaggacgaaacaccGGGCCACGCCAAAAAGCCACgtttaagagctatgctggaaacagc |
| Aldoa | Aldoa-1 | GAACATACTCCCTCGGCCC | tatcttgtggaaaggacgaaacaccGGAACATACTCCCTCGGCCCgtttaagagctatgctggaaacagc |
| Aldoa | Aldoa-2 | TTCCCCTTCAGGGCTGGAT | tatcttgtggaaaggacgaaacaccGTTCCCCTTCAGGGCTGGATgtttaagagctatgctggaaacagc |
| Aldoa | Aldoa-3 | AATGGCGAGACAACTACCC | tatcttgtggaaaggacgaaacaccGAATGGCGAGACAACTACCCgtttaagagctatgctggaaacagc |
| Alg1 | Alg1-2 | CGCGTGGTACTGCATGCGG | tatcttgtggaaaggacgaaacaccGCGCGTGGTACTGCATGCGGgtttaagagctatgctggaaacagc |
| Alg1 | Alg1-4 | GAGCAAATTGGTCATCGAC | tatcttgtggaaaggacgaaacaccGGAGCAAATTGGTCATCGACgtttaagagctatgctggaaacagc |
| Alg1 | Alg1-5 | GTGACCAATGCTATGCGGG | tatcttgtggaaaggacgaaacaccGGTGACCAATGCTATGCGGGgtttaagagctatgctggaaacagc |
| Alg2 | Alg2-1 | TATTCCTCGATCCAGTCGA | tatcttgtggaaaggacgaaacaccGTATTCCTCGATCCAGTCGAgtttaagagctatgctggaaacagc |
| Alg2 | Alg2-4 | AGCTCTCGGTGCAATGCGC | tatcttgtggaaaggacgaaacaccGAGCTCTCGGTGCAATGCGCgtttaagagctatgctggaaacagc |
| Alg2 | Alg2-5 | TGGTTCGGGTCGTAGTGCG | tatcttgtggaaaggacgaaacaccGTGGTTCGGGTCGTAGTGCGgtttaagagctatgctggaaacagc |
| Alkbh3 | Alkbh3-1 | CGACGATGAACCATCCCTG | tatcttgtggaaaggacgaaacaccGCGACGATGAACCATCCCTGgtttaagagctatgctggaaacagc |
| Alkbh3 | Alkbh3-3 | TAAGGGTGTGCTTATATCC | tatcttgtggaaaggacgaaacaccGTAAGGGTGTGCTTATATCCgtttaagagctatgctggaaacagc |
| Alkbh3 | Alkbh3-4 | AACAACCATTTACCAGCTG | tatcttgtggaaaggacgaaacaccGAACAACCATTTACCAGCTGgtttaagagctatgctggaaacagc |
| Anapc11 | Anapc11-3 | CGAGGGGGCAGTCATCACC | tatcttgtggaaaggacgaaacaccGCGAGGGGGCAGTCATCACCgtttaagagctatgctggaaacagc |
| Anapc11 | Anapc11-4 | GAGCACTGTCCCCACACGA | tatcttgtggaaaggacgaaacaccGGAGCACTGTCCCCACACGAgtttaagagctatgctggaaacagc |
| Anapc11 | Anapc11-5 | CTTGAGGATGCAGTGCATG | tatcttgtggaaaggacgaaacaccGCTTGAGGATGCAGTGCATGgtttaagagctatgctggaaacagc |
| Ang5 | Ang5-1 | ACTATGATGCCAAGCCAAC | tatcttgtggaaaggacgaaacaccGACTATGATGCCAAGCCAACgtttaagagctatgctggaaacagc |
| Ang5 | Ang5-2 | ATCTGTAATCCCGGCCAGT | tatcttgtggaaaggacgaaacaccGATCTGTAATCCCGGCCAGTgtttaagagctatgctggaaacagc |
| Ang5 | Ang5-5 | ACAACATCAAGGCCATCTG | tatcttgtggaaaggacgaaacaccGACAACATCAAGGCCATCTGgtttaagagctatgctggaaacagc |
| Ankrd66 | Ankrd66-2 | CCCTGCTTGGTAACGGATG | tatcttgtggaaaggacgaaacaccGCCCTGCTTGGTAACGGATGgtttaagagctatgctggaaacagc |
| Ankrd66 | Ankrd66-3 | TCCATCTCCTGATACAATA | tatcttgtggaaaggacgaaacaccGTCCATCTCCTGATACAATAgtttaagagctatgctggaaacagc |
| Ankrd66 | Ankrd66-4 | TGGTATATCTAAGGACAAA | tatcttgtggaaaggacgaaacaccGTGGTATATCTAAGGACAAAgtttaagagctatgctggaaacagc |
| Ap1s2 | Ap1s2-1 | TTGGGTTTCCGTGCTAAAA | tatcttgtggaaaggacgaaacaccGTTGGGTTTCCGTGCTAAAAgtttaagagctatgctggaaacagc |
| Ap1s2 | Ap1s2-4 | GACAATGAACTGATTACCC | tatcttgtggaaaggacgaaacaccGGACAATGAACTGATTACCCgtttaagagctatgctggaaacagc |
| Ap1s2 | Ap1s2-5 | AAGTAATTCCACGTAACGA | tatcttgtggaaaggacgaaacaccGAAGTAATTCCACGTAACGAgtttaagagctatgctggaaacagc |
| Arhgap24 | Arhgap24-1 | CCGATATGAAAAGAGGTAC | tatcttgtggaaaggacgaaacaccGCCGATATGAAAAGAGGTACgtttaagagctatgctggaaacagc |
| Arhgap24 | Arhgap24-3 | TCGGAGATACAATTTCAGC | tatcttgtggaaaggacgaaacaccGTCGGAGATACAATTTCAGCgtttaagagctatgctggaaacagc |
| Arhgap24 | Arhgap24-5 | CCTAACATCCTGCGCCCAA | tatcttgtggaaaggacgaaacaccGCCTAACATCCTGCGCCCAAgtttaagagctatgctggaaacagc |
| Arl2 | Arl2-1 | TTGCTCCAGCCAGGCGCTA | tatcttgtggaaaggacgaaacaccGTTGCTCCAGCCAGGCGCTAgtttaagagctatgctggaaacagc |
| Arl2 | Arl2-2 | GTCTCTGCGCTCCTACTGG | tatcttgtggaaaggacgaaacaccGGTCTCTGCGCTCCTACTGGgtttaagagctatgctggaaacagc |
| Arl2 | Arl2-4 | ACTCACCCGCGGTGCTCCA | tatcttgtggaaaggacgaaacaccGACTCACCCGCGGTGCTCCAgtttaagagctatgctggaaacagc |
| Arl4d | Arl4d-2 | TCCAGAGCGTCCCCACCAA | tatcttgtggaaaggacgaaacaccGTCCAGAGCGTCCCCACCAAgtttaagagctatgctggaaacagc |
| Arl4d | Arl4d-3 | TTCTCAGTATTAAAGCCCT | tatcttgtggaaaggacgaaacaccGTTCTCAGTATTAAAGCCCTgtttaagagctatgctggaaacagc |
| Arl4d | Arl4d-4 | GAGAAGATCCGCGTGCCCC | tatcttgtggaaaggacgaaacaccGGAGAAGATCCGCGTGCCCCgtttaagagctatgctggaaacagc |
| Armc7 | Armc7-1 | GAAGTTGGCGAGGTTGGCG | tatcttgtggaaaggacgaaacaccGGAAGTTGGCGAGGTTGGCGgtttaagagctatgctggaaacagc |
| Armc7 | Armc7-2 | CCCGGGTCATAAGCGAAGT | tatcttgtggaaaggacgaaacaccGCCCGGGTCATAAGCGAAGTgtttaagagctatgctggaaacagc |
| Armc7 | Armc7-5 | GTTGGCCTTGTCCGCGCAC | tatcttgtggaaaggacgaaacaccGGTTGGCCTTGTCCGCGCACgtttaagagctatgctggaaacagc |
| Art4 | Art4-1 | GTAGTGCAGATACTTGAAG | tatcttgtggaaaggacgaaacaccGGTAGTGCAGATACTTGAAGgtttaagagctatgctggaaacagc |
| Art4 | Art4-2 | ATGACTGACTGTACTGCCC | tatcttgtggaaaggacgaaacaccGATGACTGACTGTACTGCCCgtttaagagctatgctggaaacagc |
| Art4 | Art4-5 | CCTTTGATGATCAATACCA | tatcttgtggaaaggacgaaacaccGCCTTTGATGATCAATACCAgtttaagagctatgctggaaacagc |
| Asf1a | Asf1a-1 | CACTCGCTCACCTTCAGAC | tatcttgtggaaaggacgaaacaccGCACTCGCTCACCTTCAGACgtttaagagctatgctggaaacagc |
| Asf1a | Asf1a-4 | TGATTACTTGCACCTACCG | tatcttgtggaaaggacgaaacaccGTGATTACTTGCACCTACCGgtttaagagctatgctggaaacagc |
| Asf1a | Asf1a-5 | GTCAAGAATTTATTAGAGT | tatcttgtggaaaggacgaaacaccGGTCAAGAATTTATTAGAGTgtttaagagctatgctggaaacagc |
| Atad5 | Atad5-1 | CGCCAGACACGCTCAAAAA | tatcttgtggaaaggacgaaacaccGCGCCAGACACGCTCAAAAAgtttaagagctatgctggaaacagc |
| Atad5 | Atad5-2 | GCGTGTAGAGACATACTCC | tatcttgtggaaaggacgaaacaccGGCGTGTAGAGACATACTCCgtttaagagctatgctggaaacagc |
| Atad5 | Atad5-4 | AGCGTGATTTTCTAATGAG | tatcttgtggaaaggacgaaacaccGAGCGTGATTTTCTAATGAGgtttaagagctatgctggaaacagc |
| Atm | Atm-1 | ATAACATCGCGAAGGACAA | tatcttgtggaaaggacgaaacaccGATAACATCGCGAAGGACAAgtttaagagctatgctggaaacagc |
| Atm | Atm-3 | ATCGCTCACAGAACTAGCG | tatcttgtggaaaggacgaaacaccGATCGCTCACAGAACTAGCGgtttaagagctatgctggaaacagc |
| Atm | Atm-4 | GGATCACGGAGTACATCCA | tatcttgtggaaaggacgaaacaccGGGATCACGGAGTACATCCAgtttaagagctatgctggaaacagc |
| Atp2a2 | Atp2a2-2 | GGGTCAGTATGCTTGATGA | tatcttgtggaaaggacgaaacaccGGGGTCAGTATGCTTGATGAgtttaagagctatgctggaaacagc |
| Atp2a2 | Atp2a2-3 | CAATTTCCACTATATCACC | tatcttgtggaaaggacgaaacaccGCAATTTCCACTATATCACCgtttaagagctatgctggaaacagc |
| Atp2a2 | Atp2a2-5 | ATTACAAACGGCTCTACAA | tatcttgtggaaaggacgaaacaccGATTACAAACGGCTCTACAAgtttaagagctatgctggaaacagc |
| Atp5a1 | Atp5a1-2 | ATTCGGGGGATAATTCCAG | tatcttgtggaaaggacgaaacaccGATTCGGGGGATAATTCCAGgtttaagagctatgctggaaacagc |
| Atp5a1 | Atp5a1-3 | CCGATGCAGACCGGCATCA | tatcttgtggaaaggacgaaacaccGCCGATGCAGACCGGCATCAgtttaagagctatgctggaaacagc |
| Atp5a1 | Atp5a1-5 | ACTGCATCTACGTCGCGAT | tatcttgtggaaaggacgaaacaccGACTGCATCTACGTCGCGATgtttaagagctatgctggaaacagc |
| Atp5mpl | Atp5mpl-1 | TAAAATCAGGAGTGCTGGT | tatcttgtggaaaggacgaaacaccGTAAAATCAGGAGTGCTGGTgtttaagagctatgctggaaacagc |
| Atp5mpl | Atp5mpl-2 | CCTCATCGTATATAAAATC | tatcttgtggaaaggacgaaacaccGCCTCATCGTATATAAAATCgtttaagagctatgctggaaacagc |
| Atp5mpl | Atp5mpl-3 | GGAAATTTGGGTAGGAGTG | tatcttgtggaaaggacgaaacaccGGGAAATTTGGGTAGGAGTGgtttaagagctatgctggaaacagc |
| Atp5pb | Atp5pb-1 | ATCCAGGATGCAATCGACA | tatcttgtggaaaggacgaaacaccGATCCAGGATGCAATCGACAgtttaagagctatgctggaaacagc |
| Atp5pb | Atp5pb-3 | GACCTTATGTGCTTGGAAC | tatcttgtggaaaggacgaaacaccGGACCTTATGTGCTTGGAACgtttaagagctatgctggaaacagc |
| Atp5pb | Atp5pb-5 | CGAGGCTGTCCTGTGTGAA | tatcttgtggaaaggacgaaacaccGCGAGGCTGTCCTGTGTGAAgtttaagagctatgctggaaacagc |
| Atp6v0b | Atp6v0b-1 | ATTATCTTCTGTGAAGCGG | tatcttgtggaaaggacgaaacaccGATTATCTTCTGTGAAGCGGgtttaagagctatgctggaaacagc |
| Atp6v0b | Atp6v0b-3 | CCGGTTATATAGATTCCCC | tatcttgtggaaaggacgaaacaccGCCGGTTATATAGATTCCCCgtttaagagctatgctggaaacagc |
| Atp6v0b | Atp6v0b-5 | GAAGTTTCCGTCAGGAACC | tatcttgtggaaaggacgaaacaccGGAAGTTTCCGTCAGGAACCgtttaagagctatgctggaaacagc |
| Atp6v1a | Atp6v1a-2 | TTACGGGAAAAGGGCATCG | tatcttgtggaaaggacgaaacaccGTTACGGGAAAAGGGCATCGgtttaagagctatgctggaaacagc |
| Atp6v1a | Atp6v1a-4 | AACCTCTCTCGGTCGAGCT | tatcttgtggaaaggacgaaacaccGAACCTCTCTCGGTCGAGCTgtttaagagctatgctggaaacagc |
| Atp6v1a | Atp6v1a-5 | GAGACCCCGTACTCCGCAC | tatcttgtggaaaggacgaaacaccGGAGACCCCGTACTCCGCACgtttaagagctatgctggaaacagc |
| Atp6v1b2 | Atp6v1b2-2 | TCCACTTGACATTACCAGA | tatcttgtggaaaggacgaaacaccGTCCACTTGACATTACCAGAgtttaagagctatgctggaaacagc |
| Atp6v1b2 | Atp6v1b2-3 | TGGATAGATCCGACACTGA | tatcttgtggaaaggacgaaacaccGTGGATAGATCCGACACTGAgtttaagagctatgctggaaacagc |
| Atp6v1b2 | Atp6v1b2-5 | GCTGGATTACCACACAACG | tatcttgtggaaaggacgaaacaccGGCTGGATTACCACACAACGgtttaagagctatgctggaaacagc |
| Atp6v1e1 | Atp6v1e1-1 | GGCATCCAAGCATACCTGA | tatcttgtggaaaggacgaaacaccGGGCATCCAAGCATACCTGAgtttaagagctatgctggaaacagc |
| Atp6v1e1 | Atp6v1e1-3 | TTGTAGGATCTGCTAAATG | tatcttgtggaaaggacgaaacaccGTTGTAGGATCTGCTAAATGgtttaagagctatgctggaaacagc |
| Atp6v1e1 | Atp6v1e1-5 | CAGTCTTTGCGTTTGCACA | tatcttgtggaaaggacgaaacaccGCAGTCTTTGCGTTTGCACAgtttaagagctatgctggaaacagc |
| Avl9 | Avl9-1 | GATAGTTATGTGCACCATC | tatcttgtggaaaggacgaaacaccGGATAGTTATGTGCACCATCgtttaagagctatgctggaaacagc |
| Avl9 | Avl9-2 | CTTGAAGTAAGCCATAAAG | tatcttgtggaaaggacgaaacaccGCTTGAAGTAAGCCATAAAGgtttaagagctatgctggaaacagc |

| | | | |
|---|---|---|---|
| Avl9 | Avl9-5 | GACGTCTTAAATACCAGCT | tatcttgtggaaaggacgaaacaccGGACGTCTTAAATACCAGCTgtttaagagctatgctggaaacagc |
| B3gnt7 | B3gnt7-3 | ACACAGCATGACCGCCGAG | tatcttgtggaaaggacgaaacaccGACACAGCATGACCGCCGAGgtttaagagctatgctggaaacagc |
| B3gnt7 | B3gnt7-4 | AGAGGTCATTCGTCAGACC | tatcttgtggaaaggacgaaacaccGAGAGGTCATTCGTCAGACCgtttaagagctatgctggaaacagc |
| B3gnt7 | B3gnt7-5 | CTTGGAGGCCGTGCCGAGC | tatcttgtggaaaggacgaaacaccGCTTGGAGGCCGTGCCGAGCgtttaagagctatgctggaaacagc |
| B9d1 | B9d1-3 | TTTAAAGGTCACATCGATG | tatcttgtggaaaggacgaaacaccGTTTAAAGGTCACATCGATGgtttaagagctatgctggaaacagc |
| B9d1 | B9d1-4 | TTAAAAGTACCAACCCCTA | tatcttgtggaaaggacgaaacaccGTTAAAAGTACCAACCCCTAgtttaagagctatgctggaaacagc |
| B9d1 | B9d1-5 | TTGGGAATGATGTCGTCCG | tatcttgtggaaaggacgaaacaccGTTGGGAATGATGTCGTCCGgtttaagagctatgctggaaacagc |
| Bambi | Bambi-1 | TGTGATGCTGCTCACTGTG | tatcttgtggaaaggacgaaacaccGTGTGATGCTGCTCACTGTGgtttaagagctatgctggaaacagc |
| Bambi | Bambi-2 | CTGGACTCACTTGCAAGCA | tatcttgtggaaaggacgaaacaccGCTGGACTCACTTGCAAGCAgtttaagagctatgctggaaacagc |
| Bambi | Bambi-3 | GACATCTGCCGAGCCAAAC | tatcttgtggaaaggacgaaacaccGGACATCTGCCGAGCCAAACgtttaagagctatgctggaaacagc |
| Banp | Banp-1 | CAGACGATATGTTTGCGGT | tatcttgtggaaaggacgaaacaccGCAGACGATATGTTTGCGGTgtttaagagctatgctggaaacagc |
| Banp | Banp-4 | GATGTGCAACATGTCGGAA | tatcttgtggaaaggacgaaacaccGGATGTGCAACATGTCGGAAgtttaagagctatgctggaaacagc |
| Banp | Banp-5 | TTCTCGGCCGTGCGACAGT | tatcttgtggaaaggacgaaacaccGTTCTCGGCCGTGCGACAGTgtttaagagctatgctggaaacagc |
| Bard1 | Bard1-2 | AAATCGTAAAGGCTGCCAC | tatcttgtggaaaggacgaaacaccGAAATCGTAAAGGCTGCCACgtttaagagctatgctggaaacagc |
| Bard1 | Bard1-3 | CTATTTCACCATTTACCCG | tatcttgtggaaaggacgaaacaccGCTATTTCACCATTTACCCGgtttaagagctatgctggaaacagc |
| Bard1 | Bard1-5 | CTCAAGATAAACCGACAAT | tatcttgtggaaaggacgaaacaccGCTCAAGATAAACCGACAATgtttaagagctatgctggaaacagc |
| Bbx | Bbx-1 | CAATGATCATAATTCCGTG | tatcttgtggaaaggacgaaacaccGCAATGATCATAATTCCGTGgtttaagagctatgctggaaacagc |
| Bbx | Bbx-2 | CTAATTATTATCTTGCTAC | tatcttgtggaaaggacgaaacaccGCTAATTATTATCTTGCTACgtttaagagctatgctggaaacagc |
| Bbx | Bbx-5 | CACCTCGGTTATCAAGCCT | tatcttgtggaaaggacgaaacaccGCACCTCGGTTATCAAGCCTgtttaagagctatgctggaaacagc |
| Bcas2 | Bcas2-3 | TCATGCTGAGTAATTCAAT | tatcttgtggaaaggacgaaacaccGTCATGCTGAGTAATTCAATgtttaagagctatgctggaaacagc |
| Bcas2 | Bcas2-4 | AACAATTCTATGGCTCAGT | tatcttgtggaaaggacgaaacaccGAACAATTCTATGGCTCAGTgtttaagagctatgctggaaacagc |
| Bcas2 | Bcas2-5 | GCGGTCCGGATCGAGAATC | tatcttgtggaaaggacgaaacaccGGCGGTCCGGATCGAGAATCgtttaagagctatgctggaaacagc |
| Bckdk | Bckdk-3 | AGGATTGCTCACCGCATCA | tatcttgtggaaaggacgaaacaccGAGGATTGCTCACCGCATCAgtttaagagctatgctggaaacagc |
| Bckdk | Bckdk-4 | GGTTGCAACCAATGATGAA | tatcttgtggaaaggacgaaacaccGGGTTGCAACCAATGATGAAgtttaagagctatgctggaaacagc |
| Bckdk | Bckdk-5 | CAGCACGAGCTATACATCC | tatcttgtggaaaggacgaaacaccGCAGCACGAGCTATACATCCgtttaagagctatgctggaaacagc |
| Bcl2a1d | Bcl2a1d-1 | TGGCATCATTAACTGGGGA | tatcttgtggaaaggacgaaacaccGTGGCATCATTAACTGGGGAgtttaagagctatgctggaaacagc |
| Bcl2a1d | Bcl2a1d-2 | TGGAAAAAGAGTTTGAAGA | tatcttgtggaaaggacgaaacaccGTGGAAAAAGAGTTTGAAGAgtttaagagctatgctggaaacagc |
| Bcl2a1d | Bcl2a1d-3 | ATTATTCTGGCGGTATCTA | tatcttgtggaaaggacgaaacaccGATTATTCTGGCGGTATCTAgtttaagagctatgctggaaacagc |
| Bid | Bid-2 | CATACTTACTTCCTCCGAC | tatcttgtggaaaggacgaaacaccGCATACTTACTTCCTCCGACgtttaagagctatgctggaaacagc |
| Bid | Bid-3 | ATGAATGGCAGCCTGTCGG | tatcttgtggaaaggacgaaacaccGATGAATGGCAGCCTGTCGGgtttaagagctatgctggaaacagc |
| Bid | Bid-5 | GGCTGTACTCGCCAAGAGC | tatcttgtggaaaggacgaaacaccGGGCTGTACTCGCCAAGAGCgtttaagagctatgctggaaacagc |
| Bop1 | Bop1-2 | TGCATCCACGCCGAGCGCT | tatcttgtggaaaggacgaaacaccGTGCATCCACGCCGAGCGCTgtttaagagctatgctggaaacagc |
| Bop1 | Bop1-3 | AGCTCATCTCGTGTCCGCA | tatcttgtggaaaggacgaaacaccGAGCTCATCTCGTGTCCGCAgtttaagagctatgctggaaacagc |
| Bop1 | Bop1-4 | GTAGATACGTTTGCCATCC | tatcttgtggaaaggacgaaacaccGGTAGATACGTTTGCCATCCgtttaagagctatgctggaaacagc |
| Brca1 | Brca1-1 | TTAAGCGCGTGTCTCAAGG | tatcttgtggaaaggacgaaacaccGTTAAGCGCGTGTCTCAAGGgtttaagagctatgctggaaacagc |
| Brca1 | Brca1-4 | GTCTACATTGAACTAGGTA | tatcttgtggaaaggacgaaacaccGGTCTACATTGAACTAGGTAgtttaagagctatgctggaaacagc |
| Brca1 | Brca1-5 | GCGTCGATCATCCAGAGCG | tatcttgtggaaaggacgaaacaccGGCGTCGATCATCCAGAGCGgtttaagagctatgctggaaacagc |
| Brd4 | Brd4-2 | GTCTTAATGGCAGAAGCTC | tatcttgtggaaaggacgaaacaccGGTCTTAATGGCAGAAGCTCgtttaagagctatgctggaaacagc |
| Brd4 | Brd4-3 | GGAACAATAAAGAAGCGCT | tatcttgtggaaaggacgaaacaccGGGAACAATAAAGAAGCGCTgtttaagagctatgctggaaacagc |
| Brd4 | Brd4-5 | CTGGAAAGGCCACGCAAAC | tatcttgtggaaaggacgaaacaccGCTGGAAAGGCCACGCAAACgtttaagagctatgctggaaacagc |
| Brf2 | Brf2-2 | GGCCAGTTATGTTGCCGAC | tatcttgtggaaaggacgaaacaccGGGCCAGTTATGTTGCCGACgtttaagagctatgctggaaacagc |
| Brf2 | Brf2-4 | AAGTTGCCCTCGTCGCTAA | tatcttgtggaaaggacgaaacaccGAAGTTGCCCTCGTCGCTAAgtttaagagctatgctggaaacagc |
| Brf2 | Brf2-5 | CGGCTGCGTGGTCACCGAA | tatcttgtggaaaggacgaaacaccGCGGCTGCGTGGTCACCGAAgtttaagagctatgctggaaacagc |
| Brix1 | Brix1-2 | TGGCTGAATTAAAGATGAC | tatcttgtggaaaggacgaaacaccGTGGCTGAATTAAAGATGACgtttaagagctatgctggaaacagc |
| Brix1 | Brix1-4 | AAGTACCTGCTTTAGAATG | tatcttgtggaaaggacgaaacaccGAAGTACCTGCTTTAGAATGgtttaagagctatgctggaaacagc |
| Brix1 | Brix1-5 | TTACCTTGCAAACCGGACC | tatcttgtggaaaggacgaaacaccGTTACCTTGCAAACCGGACCgtttaagagctatgctggaaacagc |
| Btaf1 | Btaf1-1 | GGGATACCCGGATTGCAGC | tatcttgtggaaaggacgaaacaccGGGGATACCCGGATTGCAGCgtttaagagctatgctggaaacagc |
| Btaf1 | Btaf1-4 | TAAACGCGGTTGCACAGCT | tatcttgtggaaaggacgaaacaccGTAAACGCGGTTGCACAGCTgtttaagagctatgctggaaacagc |
| Btaf1 | Btaf1-5 | TTAGATAGTAAACGGCATC | tatcttgtggaaaggacgaaacaccGTTAGATAGTAAACGGCATCgtttaagagctatgctggaaacagc |
| Bysl | Bysl-3 | TTAAGCCTTACCTCACGGG | tatcttgtggaaaggacgaaacaccGTTAAGCCTTACCTCACGGGgtttaagagctatgctggaaacagc |
| Bysl | Bysl-4 | ACCGCGAGAGCGCGCCACG | tatcttgtggaaaggacgaaacaccGACCGCGAGAGCGCGCCACGgtttaagagctatgctggaaacagc |
| Bysl | Bysl-5 | GGACCCGAGAAAAACGGCG | tatcttgtggaaaggacgaaacaccGGGACCCGAGAAAAACGGCGgtttaagagctatgctggaaacagc |
| Bzw1 | Bzw1-1 | TTCAAGGCTTAACTGAAAC | tatcttgtggaaaggacgaaacaccGTTCAAGGCTTAACTGAAACgtttaagagctatgctggaaacagc |
| Bzw1 | Bzw1-2 | TTAGCCCCGGGTGGTACAC | tatcttgtggaaaggacgaaacaccGTTAGCCCCGGGTGGTACACgtttaagagctatgctggaaacagc |
| Bzw1 | Bzw1-3 | GCTACAAATACCTGGAGAA | tatcttgtggaaaggacgaaacaccGGCTACAAATACCTGGAGAAgtttaagagctatgctggaaacagc |
| C1galt1c1 | C1galt1c1-2 | TCAATAACAGCGAAAGTAG | tatcttgtggaaaggacgaaacaccGTCAATAACAGCGAAAGTAGgtttaagagctatgctggaaacagc |
| C1galt1c1 | C1galt1c1-3 | TATGCTTATGATCAATACA | tatcttgtggaaaggacgaaacaccGTATGCTTATGATCAATACAgtttaagagctatgctggaaacagc |
| C1galt1c1 | C1galt1c1-4 | AAGAACTATACAGTATACC | tatcttgtggaaaggacgaaacaccGAAGAACTATACAGTATACCgtttaagagctatgctggaaacagc |
| Castor2 | Castor2-1 | TACACCATCATCGTAGATG | tatcttgtggaaaggacgaaacaccGTACACCATCATCGTAGATGgtttaagagctatgctggaaacagc |
| Castor2 | Castor2-4 | GATGTTCTGGTCGGCGAGT | tatcttgtggaaaggacgaaacaccGGATGTTCTGGTCGGCGAGTgtttaagagctatgctggaaacagc |
| Castor2 | Castor2-5 | CCATTGACAACGCGTAGGA | tatcttgtggaaaggacgaaacaccGCCATTGACAACGCGTAGGAgtttaagagctatgctggaaacagc |
| Ccdc137 | Ccdc137-3 | CCAAGTTCAAGCAAAGGAA | tatcttgtggaaaggacgaaacaccGCCAAGTTCAAGCAAAGGAAgtttaagagctatgctggaaacagc |
| Ccdc137 | Ccdc137-4 | TAGCAAGAACCAGGCCCCT | tatcttgtggaaaggacgaaacaccGTAGCAAGAACCAGGCCCCTgtttaagagctatgctggaaacagc |
| Ccdc137 | Ccdc137-5 | TGCACCTCTGGCTGCCGAG | tatcttgtggaaaggacgaaacaccGTGCACCTCTGGCTGCCGAGgtttaagagctatgctggaaacagc |
| Ccdc175 | Ccdc175-1 | GTGACTCAAAAGCAAAATC | tatcttgtggaaaggacgaaacaccGGTGACTCAAAAGCAAAATCgtttaagagctatgctggaaacagc |
| Ccdc175 | Ccdc175-2 | TCTTTTCTCTTATCGTCAC | tatcttgtggaaaggacgaaacaccGTCTTTTCTCTTATCGTCACgtttaagagctatgctggaaacagc |
| Ccdc175 | Ccdc175-5 | AGCGCATTGGCCGCTACCG | tatcttgtggaaaggacgaaacaccGAGCGCATTGGCCGCTACCGgtttaagagctatgctggaaacagc |
| Ccdc7b | Ccdc7b-1 | ATTTTATCAGGTGTCGACT | tatcttgtggaaaggacgaaacaccGATTTTATCAGGTGTCGACTgtttaagagctatgctggaaacagc |
| Ccdc7b | Ccdc7b-3 | TGAAGAAAAACCTGCTAGA | tatcttgtggaaaggacgaaacaccGTGAAGAAAAACCTGCTAGAgtttaagagctatgctggaaacagc |
| Ccdc7b | Ccdc7b-5 | GGCTGAGAGATCCCGAGGT | tatcttgtggaaaggacgaaacaccGGGCTGAGAGATCCCGAGGTgtttaagagctatgctggaaacagc |
| Ccdc84 | Ccdc84-1 | GAGGACGAGGTGATTAAAG | tatcttgtggaaaggacgaaacaccGGAGGACGAGGTGATTAAAGgtttaagagctatgctggaaacagc |
| Ccdc84 | Ccdc84-2 | AACTCACCGTGCATAATCC | tatcttgtggaaaggacgaaacaccGAACTCACCGTGCATAATCCgtttaagagctatgctggaaacagc |
| Ccdc84 | Ccdc84-3 | AACCTGACAGTGCTACACG | tatcttgtggaaaggacgaaacaccGAACCTGACAGTGCTACACGgtttaagagctatgctggaaacagc |
| Ccdc91 | Ccdc91-1 | AGGAGGTAGTAGTACCTGC | tatcttgtggaaaggacgaaacaccGAGGAGGTAGTAGTACCTGCgtttaagagctatgctggaaacagc |
| Ccdc91 | Ccdc91-3 | AGATGAGTGGTCATGGTCC | tatcttgtggaaaggacgaaacaccGAGATGAGTGGTCATGGTCCgtttaagagctatgctggaaacagc |
| Ccdc91 | Ccdc91-4 | GCTCATCCGTTAAACCAAG | tatcttgtggaaaggacgaaacaccGGCTCATCCGTTAAACCAAGgtttaagagctatgctggaaacagc |

| | | | |
|---|---|---|---|
| Ccnjl | Ccnjl-1 | AATGTCTACAAAGAATCGA | tatcttgtggaaaggacgaaacaccGAATGTCTACAAAGAATCGAgtttaagagctatgctggaaacagc |
| Ccnjl | Ccnjl-3 | CGTAGGTAAGTTTGAAGAC | tatcttgtggaaaggacgaaacaccGCGTAGGTAAGTTTGAAGACgtttaagagctatgctggaaacagc |
| Ccnjl | Ccnjl-5 | TCCAGAAAGTGGGCGGGCG | tatcttgtggaaaggacgaaacaccGTCCAGAAAGTGGGCGGGCGgtttaagagctatgctggaaacagc |
| Ccnl1 | Ccnl1-2 | ATGCGCAGATCCGTCTCGC | tatcttgtggaaaggacgaaacaccGATGCGCAGATCCGTCTCGCgtttaagagctatgctggaaacagc |
| Ccnl1 | Ccnl1-4 | GAGCGACACCTCCGAATAC | tatcttgtggaaaggacgaaacaccGGAGCGACACCTCCGAATACgtttaagagctatgctggaaacagc |
| Ccnl1 | Ccnl1-5 | ATCAGGATCCCTCCGGTCG | tatcttgtggaaaggacgaaacaccGATCAGGATCCCTCCGGTCGgtttaagagctatgctggaaacagc |
| Ccnq | Ccnq-1 | TCGCACAGGTACTTTAACC | tatcttgtggaaaggacgaaacaccGTCGCACAGGTACTTTAACCgtttaagagctatgctggaaacagc |
| Ccnq | Ccnq-2 | CATCATCAACGTGTCGCAC | tatcttgtggaaaggacgaaacaccGCATCATCAACGTGTCGCACgtttaagagctatgctggaaacagc |
| Ccnq | Ccnq-4 | GGTGCAAGCGGTGGCGATG | tatcttgtggaaaggacgaaacaccGGGTGCAAGCGGTGGCGATGgtttaagagctatgctggaaacagc |
| Cd180 | Cd180-1 | AGATTGATAAGTCTGCTGA | tatcttgtggaaaggacgaaacaccGAGATTGATAAGTCTGCTGAgtttaagagctatgctggaaacagc |
| Cd180 | Cd180-2 | TACCTACCTGGTTAAATCC | tatcttgtggaaaggacgaaacaccGTACCTACCTGGTTAAATCCgtttaagagctatgctggaaacagc |
| Cd180 | Cd180-3 | TTCATGTATCCAGTAAATC | tatcttgtggaaaggacgaaacaccGTTCATGTATCCAGTAAATCgtttaagagctatgctggaaacagc |
| Cd36 | Cd36-1 | ATCCTATAAGTACCACAGT | tatcttgtggaaaggacgaaacaccGATCCTATAAGTACCACAGTgtttaagagctatgctggaaacagc |
| Cd36 | Cd36-4 | CAAAACTGTCTGTACACAG | tatcttgtggaaaggacgaaacaccGCAAAACTGTCTGTACACAGgtttaagagctatgctggaaacagc |
| Cd36 | Cd36-5 | ACCACTGCTTTCAAAAACT | tatcttgtggaaaggacgaaacaccGACCACTGCTTTCAAAAACTgtttaagagctatgctggaaacagc |
| Cd46 | Cd46-2 | GGGTCCCTATTTCAGATGC | tatcttgtggaaaggacgaaacaccGGGGTCCCTATTTCAGATGCgtttaagagctatgctggaaacagc |
| Cd46 | Cd46-3 | AGGTAGCAATCATCAGGTA | tatcttgtggaaaggacgaaacaccGAGGTAGCAATCATCAGGTAgtttaagagctatgctggaaacagc |
| Cd46 | Cd46-5 | TTGAAGCTATGGAACTCAA | tatcttgtggaaaggacgaaacaccGTTGAAGCTATGGAACTCAAgtttaagagctatgctggaaacagc |
| Cd74 | Cd74-2 | TAACATGCTCCTTGGGGTA | tatcttgtggaaaggacgaaacaccGTAACATGCTCCTTGGGGTAgtttaagagctatgctggaaacagc |
| Cd74 | Cd74-3 | TGAAGAACGTTACCAAGTA | tatcttgtggaaaggacgaaacaccGTGAAGAACGTTACCAAGTAgtttaagagctatgctggaaacagc |
| Cd74 | Cd74-4 | CGTGAGCAGATGCATCACA | tatcttgtggaaaggacgaaacaccGCGTGAGCAGATGCATCACAgtttaagagctatgctggaaacagc |
| Cd8a | Cd8a-1 | GCCGAACTTGGTCAGAAGG | tatcttgtggaaaggacgaaacaccGGCCGAACTTGGTCAGAAGGgtttaagagctatgctggaaacagc |
| Cd8a | Cd8a-3 | GCCATATAGACAACGAAGG | tatcttgtggaaaggacgaaacaccGGCCATATAGACAACGAAGGgtttaagagctatgctggaaacagc |
| Cd8a | Cd8a-5 | TTATTATTCGTGTCCCTCA | tatcttgtggaaaggacgaaacaccGTTATTATTCGTGTCCCTCAgtttaagagctatgctggaaacagc |
| Cdadc1 | Cdadc1-2 | CATGTAACACACCAACGGC | tatcttgtggaaaggacgaaacaccGCATGTAACACACCAACGGCgtttaagagctatgctggaaacagc |
| Cdadc1 | Cdadc1-3 | TGGAGTAAGACACACACAT | tatcttgtggaaaggacgaaacaccGTGGAGTAAGACACACACATgtttaagagctatgctggaaacagc |
| Cdadc1 | Cdadc1-5 | TTAATTAGGTGAAGAAAAC | tatcttgtggaaaggacgaaacaccGTTAATTAGGTGAAGAAAACgtttaagagctatgctggaaacagc |
| Cdc16 | Cdc16-2 | TACGTGGCATCTAGCTGCA | tatcttgtggaaaggacgaaacaccGTACGTGGCATCTAGCTGCAgtttaagagctatgctggaaacagc |
| Cdc16 | Cdc16-3 | GCTTTAGATAATCGCACCC | tatcttgtggaaaggacgaaacaccGGCTTTAGATAATCGCACCCgtttaagagctatgctggaaacagc |
| Cdc16 | Cdc16-5 | TAGGTGTCTTGGTTTGCGG | tatcttgtggaaaggacgaaacaccGTAGGTGTCTTGGTTTGCGGgtttaagagctatgctggaaacagc |
| Cdc20 | Cdc20-1 | CCGTTCAGGTTGAGAGACC | tatcttgtggaaaggacgaaacaccGCCGTTCAGGTTGAGAGACCgtttaagagctatgctggaaacagc |
| Cdc20 | Cdc20-2 | AGGGGGACGCCCACCAAAA | tatcttgtggaaaggacgaaacaccGAGGGGGACGCCCACCAAAAgtttaagagctatgctggaaacagc |
| Cdc20 | Cdc20-3 | GGATAAAGCGGTCACCTCC | tatcttgtggaaaggacgaaacaccGGGATAAAGCGGTCACCTCCgtttaagagctatgctggaaacagc |
| Cdipt | Cdipt-3 | CAGCTCGAGCCCTTAATCA | tatcttgtggaaaggacgaaacaccGCAGCTCGAGCCCTTAATCAgtttaagagctatgctggaaacagc |
| Cdipt | Cdipt-4 | TCGAGCATGGCCCCAAACC | tatcttgtggaaaggacgaaacaccGTCGAGCATGGCCCCAAACCgtttaagagctatgctggaaacagc |
| Cdipt | Cdipt-5 | AGTTCATTTCCAGCACACA | tatcttgtggaaaggacgaaacaccGAGTTCATTTCCAGCACACAgtttaagagctatgctggaaacagc |
| Cdk9 | Cdk9-2 | TTAGCTAGGCTGAAGGCAC | tatcttgtggaaaggacgaaacaccGTTAGCTAGGCTGAAGGCACgtttaagagctatgctggaaacagc |
| Cdk9 | Cdk9-3 | GTGCTCATTACCCGGGATG | tatcttgtggaaaggacgaaacaccGGTGCTCATTACCCGGGATGgtttaagagctatgctggaaacagc |
| Cdk9 | Cdk9-5 | TTGAGATTTGTCGGACCAA | tatcttgtggaaaggacgaaacaccGTTGAGATTTGTCGGACCAAgtttaagagctatgctggaaacagc |
| Cdon | Cdon-2 | AAAATGTACGGTGGCCCCC | tatcttgtggaaaggacgaaacaccGAAAATGTACGGTGGCCCCCgtttaagagctatgctggaaacagc |
| Cdon | Cdon-3 | CTGGTGCGGTGACAATAAC | tatcttgtggaaaggacgaaacaccGCTGGTGCGGTGACAATAACgtttaagagctatgctggaaacagc |
| Cdon | Cdon-4 | TTCATTGGTACGGCCGCCA | tatcttgtggaaaggacgaaacaccGTTCATTGGTACGGCCGCCAgtttaagagctatgctggaaacagc |
| Celf2 | Celf2-1 | GGCTGCTCTTCACCAATCG | tatcttgtggaaaggacgaaacaccGGGCTGCTCTTCACCAATCGgtttaagagctatgctggaaacagc |
| Celf2 | Celf2-3 | TAGTGAAAAGTCAAACGGT | tatcttgtggaaaggacgaaacaccGTAGTGAAAAGTCAAACGGTgtttaagagctatgctggaaacagc |
| Celf2 | Celf2-5 | TTGTACCTTTACTCTGGGG | tatcttgtggaaaggacgaaacaccGTTGTACCTTTACTCTGGGGgtttaagagctatgctggaaacagc |
| Cenpk | Cenpk-1 | CTCACGAATGCAGATGCTC | tatcttgtggaaaggacgaaacaccGCTCACGAATGCAGATGCTCgtttaagagctatgctggaaacagc |
| Cenpk | Cenpk-3 | GGTAACATTAACTGAGTCA | tatcttgtggaaaggacgaaacaccGGGTAACATTAACTGAGTCAgtttaagagctatgctggaaacagc |
| Cenpk | Cenpk-4 | AACTGACAACTAAGATCCG | tatcttgtggaaaggacgaaacaccGAACTGACAACTAAGATCCGgtttaagagctatgctggaaacagc |
| Cenpm | Cenpm-1 | GTGTGCTTCCTTGTCACAG | tatcttgtggaaaggacgaaacaccGGTGTGCTTCCTTGTCACAGgtttaagagctatgctggaaacagc |
| Cenpm | Cenpm-2 | ACACAATCAGGTCAATTCG | tatcttgtggaaaggacgaaacaccGACACAATCAGGTCAATTCGgtttaagagctatgctggaaacagc |
| Cenpm | Cenpm-5 | TGACTGTGCGTCCGAGCTG | tatcttgtggaaaggacgaaacaccGTGACTGTGCGTCCGAGCTGgtttaagagctatgctggaaacagc |
| Cenpq | Cenpq-1 | GTGTCAAAAATGCTGAAAA | tatcttgtggaaaggacgaaacaccGGTGTCAAAAATGCTGAAAAgtttaagagctatgctggaaacagc |
| Cenpq | Cenpq-2 | GGGTTACCTTTTCTTCAAC | tatcttgtggaaaggacgaaacaccGGGGTTACCTTTTCTTCAACgtttaagagctatgctggaaacagc |
| Cenpq | Cenpq-4 | ACGTATGTTCATCTAAAAC | tatcttgtggaaaggacgaaacaccGACGTATGTTCATCTAAAACgtttaagagctatgctggaaacagc |
| Cep55 | Cep55-2 | AAGAATGTTTATTACCTCC | tatcttgtggaaaggacgaaacaccGAAGAATGTTTATTACCTCCgtttaagagctatgctggaaacagc |
| Cep55 | Cep55-3 | TTTCGTCGTGGCGGACAGC | tatcttgtggaaaggacgaaacaccGTTTCGTCGTGGCGGACAGCgtttaagagctatgctggaaacagc |
| Cep55 | Cep55-5 | GCTAAAAGCCCTTTAACGT | tatcttgtggaaaggacgaaacaccGGCTAAAAGCCCTTTAACGTgtttaagagctatgctggaaacagc |
| Cfap36 | Cfap36-2 | AAAAAATGAGTAATTCCCA | tatcttgtggaaaggacgaaacaccGAAAAAATGAGTAATTCCCAgtttaagagctatgctggaaacagc |
| Cfap36 | Cfap36-3 | CAAGCCATCCGAATAATCC | tatcttgtggaaaggacgaaacaccGCAAGCCATCCGAATAATCCgtttaagagctatgctggaaacagc |
| Cfap36 | Cfap36-5 | GAAGCGTGCACGTCCCCTC | tatcttgtggaaaggacgaaacaccGGAAGCGTGCACGTCCCCTCgtttaagagctatgctggaaacagc |
| Cfap44 | Cfap44-1 | GCTGACTCATATACCGACG | tatcttgtggaaaggacgaaacaccGGCTGACTCATATACCGACGgtttaagagctatgctggaaacagc |
| Cfap44 | Cfap44-2 | GCGAATCGCGCTTACGTGC | tatcttgtggaaaggacgaaacaccGGCGAATCGCGCTTACGTGCgtttaagagctatgctggaaacagc |
| Cfap44 | Cfap44-5 | TGATCGGGGTGCCCAAATG | tatcttgtggaaaggacgaaacaccGTGATCGGGGTGCCCAAATGgtttaagagctatgctggaaacagc |
| Chaf1b | Chaf1b-3 | TCGCAATATATTGACCCAA | tatcttgtggaaaggacgaaacaccGTCGCAATATATTGACCCAAgtttaagagctatgctggaaacagc |
| Chaf1b | Chaf1b-4 | ACGAACCTGTCACAGCTCA | tatcttgtggaaaggacgaaacaccGACGAACCTGTCACAGCTCAgtttaagagctatgctggaaacagc |
| Chaf1b | Chaf1b-5 | GGCACGGAAGATGGGCGAT | tatcttgtggaaaggacgaaacaccGGGCACGGAAGATGGGCGATgtttaagagctatgctggaaacagc |
| Chek2 | Chek2-1 | GGATTCTCCAATCTAGGTA | tatcttgtggaaaggacgaaacaccGGGATTCTCCAATCTAGGTAgtttaagagctatgctggaaacagc |
| Chek2 | Chek2-3 | ATAAGCTCGGTATTTACGA | tatcttgtggaaaggacgaaacaccGATAAGCTCGGTATTTACGAgtttaagagctatgctggaaacagc |
| Chek2 | Chek2-5 | AAAGATCATTAGCAAGCGG | tatcttgtggaaaggacgaaacaccGAAAGATCATTAGCAAGCGGgtttaagagctatgctggaaacagc |
| Chmp4b | Chmp4b-2 | TTGGTGTTGGCGTTCTCTA | tatcttgtggaaaggacgaaacaccGTTGGTGTTGGCGTTCTCTAgtttaagagctatgctggaaacagc |
| Chmp4b | Chmp4b-3 | CGGAGGTGCTCAAGAACAT | tatcttgtggaaaggacgaaacaccGCGGAGGTGCTCAAGAACATgtttaagagctatgctggaaacagc |
| Chmp4b | Chmp4b-4 | TATGCCGCCAAGGCCATGA | tatcttgtggaaaggacgaaacaccGTATGCCGCCAAGGCCATGAgtttaagagctatgctggaaacagc |
| Chrna1 | Chrna1-1 | TGGTGGCCATTAACCCGGT | tatcttgtggaaaggacgaaacaccGTGGTGGCCATTAACCCGGTgtttaagagctatgctggaaacagc |
| Chrna1 | Chrna1-3 | GGTCGATTACAACTTGAAA | tatcttgtggaaaggacgaaacaccGGGTCGATTACAACTTGAAAgtttaagagctatgctggaaacagc |
| Chrna1 | Chrna1-5 | AGATTGTACAAGTCACCGT | tatcttgtggaaaggacgaaacaccGAGATTGTACAAGTCACCGTgtttaagagctatgctggaaacagc |
| Ciao2b | Ciao2b-2 | ATGGAGAGGCCAATAAGCG | tatcttgtggaaaggacgaaacaccGATGGAGAGGCCAATAAGCGgtttaagagctatgctggaaacagc |

| | | | |
|---|---|---|---|
| Ciao2b | Ciao2b-3 | CTCCGGGTCATTAATGGAG | tatcttgtggaaaggacgaaacaccGCTCCGGGTCATTAATGGAGgtttaagagctatgctggaaacagc |
| Ciao2b | Ciao2b-4 | CGGATATCGAAGATCTCCC | tatcttgtggaaaggacgaaacaccGCGGATATCGAAGATCTCCCgtttaagagctatgctggaaacagc |
| Ciao3 | Ciao3-1 | ATGACTGCCTGGCGTGCAG | tatcttgtggaaaggacgaaacaccGATGACTGCCTGGCGTGCAGgtttaagagctatgctggaaacagc |
| Ciao3 | Ciao3-2 | GCTCTGGACTGCGGCGATA | tatcttgtggaaaggacgaaacaccGGCTCTGGACTGCGGCGATAgtttaagagctatgctggaaacagc |
| Ciao3 | Ciao3-4 | CCCCTACATCAGTACAGCC | tatcttgtggaaaggacgaaacaccGCCCCTACATCAGTACAGCCgtttaagagctatgctggaaacagc |
| Cip2a | Cip2a-2 | TTGCCAATAGTCCTAGACA | tatcttgtggaaaggacgaaacaccGTTGCCAATAGTCCTAGACAgtttaagagctatgctggaaacagc |
| Cip2a | Cip2a-3 | AGCGAAGTAGAGCCGCAGT | tatcttgtggaaaggacgaaacaccGAGCGAAGTAGAGCCGCAGTgtttaagagctatgctggaaacagc |
| Cip2a | Cip2a-5 | AACTGCTGCTCTATAAGCG | tatcttgtggaaaggacgaaacaccGAACTGCTGCTCTATAAGCGgtttaagagctatgctggaaacagc |
| Clec14a | Clec14a-1 | GGAGGTACCTCCCGGAACA | tatcttgtggaaaggacgaaacaccGGGAGGTACCTCCCGGAACAgtttaagagctatgctggaaacagc |
| Clec14a | Clec14a-4 | AGGAGGCCTGCAGCCTAAG | tatcttgtggaaaggacgaaacaccGAGGAGGCCTGCAGCCTAAGgtttaagagctatgctggaaacagc |
| Clec14a | Clec14a-5 | CCTTCTCTTGAAGGTAGCG | tatcttgtggaaaggacgaaacaccGCCTTCTCTTGAAGGTAGCGgtttaagagctatgctggaaacagc |
| Clec1a | Clec1a-1 | TGTGTGCTTATCTTCAGCA | tatcttgtggaaaggacgaaacaccGTGTGTGCTTATCTTCAGCAgtttaagagctatgctggaaacagc |
| Clec1a | Clec1a-2 | GTGAACTTTACAACAAAAG | tatcttgtggaaaggacgaaacaccGGTGAACTTTACAACAAAAGgtttaagagctatgctggaaacagc |
| Clec1a | Clec1a-4 | TTGCTGGATATTAGACAGC | tatcttgtggaaaggacgaaacaccGTTGCTGGATATTAGACAGCgtttaagagctatgctggaaacagc |
| Clnk | Clnk-1 | AGTGCAAAAGACTTCAGTA | tatcttgtggaaaggacgaaacaccGAGTGCAAAAGACTTCAGTAgtttaagagctatgctggaaacagc |
| Clnk | Clnk-2 | TTTTAATGTCTTACCGAGG | tatcttgtggaaaggacgaaacaccGTTTTAATGTCTTACCGAGGgtttaagagctatgctggaaacagc |
| Clnk | Clnk-5 | TCCGATTCCTGGATAGGTC | tatcttgtggaaaggacgaaacaccGTCCGATTCCTGGATAGGTCgtttaagagctatgctggaaacagc |
| Clvs2 | Clvs2-2 | GCTTCTAAAGAAAGTAAGA | tatcttgtggaaaggacgaaacaccGGCTTCTAAAGAAAGTAAGAgtttaagagctatgctggaaacagc |
| Clvs2 | Clvs2-4 | CTCCCATAATGGTCCAGAT | tatcttgtggaaaggacgaaacaccGCTCCCATAATGGTCCAGATgtttaagagctatgctggaaacagc |
| Clvs2 | Clvs2-5 | TCATCACTAGGCCGGACAT | tatcttgtggaaaggacgaaacaccGTCATCACTAGGCCGGACATgtttaagagctatgctggaaacagc |
| Cmc1 | Cmc1-1 | TGCATTCTTCATAGAAGGC | tatcttgtggaaaggacgaaacaccGTGCATTCTTCATAGAAGGCgtttaagagctatgctggaaacagc |
| Cmc1 | Cmc1-3 | CCAGAGTCCTTGCAACATC | tatcttgtggaaaggacgaaacaccGCCAGAGTCCTTGCAACATCgtttaagagctatgctggaaacagc |
| Cmc1 | Cmc1-4 | GGTGCTCCGAACAAGTTGA | tatcttgtggaaaggacgaaacaccGGGTGCTCCGAACAAGTTGAgtttaagagctatgctggaaacagc |
| Cmtm3 | Cmtm3-2 | TGATGAATGAAAGACCCTG | tatcttgtggaaaggacgaaacaccGTGATGAATGAAAGACCCTGgtttaagagctatgctggaaacagc |
| Cmtm3 | Cmtm3-3 | GACACCACGTAGCAGATGA | tatcttgtggaaaggacgaaacaccGGACACCACGTAGCAGATGAgtttaagagctatgctggaaacagc |
| Cmtm3 | Cmtm3-4 | AGCTGAATGACAAATGGCA | tatcttgtggaaaggacgaaacaccGAGCTGAATGACAAATGGCAgtttaagagctatgctggaaacagc |
| Cnep1r1 | Cnep1r1-2 | TCATCAGTGCTTCTTATAG | tatcttgtggaaaggacgaaacaccGTCATCAGTGCTTCTTATAGgtttaagagctatgctggaaacagc |
| Cnep1r1 | Cnep1r1-4 | GCTAATAGTGAAAAACGGA | tatcttgtggaaaggacgaaacaccGGCTAATAGTGAAAAACGGAgtttaagagctatgctggaaacagc |
| Cnep1r1 | Cnep1r1-5 | TTAGCTGTATAACTCTAAT | tatcttgtggaaaggacgaaacaccGTTAGCTGTATAACTCTAATgtttaagagctatgctggaaacagc |
| Cnih1 | Cnih1-1 | TAAATATTACCTCCAAATA | tatcttgtggaaaggacgaaacaccGTAAATATTACCTCCAAATAgtttaagagctatgctggaaacagc |
| Cnih1 | Cnih1-4 | GGTATTGCACTGGTCTATA | tatcttgtggaaaggacgaaacaccGGGTATTGCACTGGTCTATAgtttaagagctatgctggaaacagc |
| Cnih1 | Cnih1-5 | TCTATAGGGTTCTTGTAGT | tatcttgtggaaaggacgaaacaccGTCTATAGGGTTCTTGTAGTgtttaagagctatgctggaaacagc |
| Cnot3 | Cnot3-1 | ACCAAAGCGTATAGCAAGG | tatcttgtggaaaggacgaaacaccGACCAAAGCGTATAGCAAGGgtttaagagctatgctggaaacagc |
| Cnot3 | Cnot3-2 | TCATCCTTGATCTTGCGGA | tatcttgtggaaaggacgaaacaccGTCATCCTTGATCTTGCGGAgtttaagagctatgctggaaacagc |
| Cnot3 | Cnot3-3 | TACTACGTTGACTCATCCC | tatcttgtggaaaggacgaaacaccGTACTACGTTGACTCATCCCgtttaagagctatgctggaaacagc |
| Cnot4 | Cnot4-2 | ATATACGTTACATAAGCAC | tatcttgtggaaaggacgaaacaccGATATACGTTACATAAGCACgtttaagagctatgctggaaacagc |
| Cnot4 | Cnot4-3 | AATTCGCACTGATGAGAAT | tatcttgtggaaaggacgaaacaccGAATTCGCACTGATGAGAATgtttaagagctatgctggaaacagc |
| Cnot4 | Cnot4-4 | AATTCGATGCCAACAAAAT | tatcttgtggaaaggacgaaacaccGAATTCGATGCCAACAAAATgtttaagagctatgctggaaacagc |
| Cnot8 | Cnot8-1 | AGCTACAGCTATATCGCTA | tatcttgtggaaaggacgaaacaccGAGCTACAGCTATATCGCTAgtttaagagctatgctggaaacagc |
| Cnot8 | Cnot8-2 | TCGGCCGTACAACAACACC | tatcttgtggaaaggacgaaacaccGTCGGCCGTACAACAACACCgtttaagagctatgctggaaacagc |
| Cnot8 | Cnot8-3 | TTCTTAAAATCATCCAGCT | tatcttgtggaaaggacgaaacaccGTTCTTAAAATCATCCAGCTgtttaagagctatgctggaaacagc |
| Cntrob | Cntrob-2 | GATGTGGCGCTCGCAATGC | tatcttgtggaaaggacgaaacaccGGATGTGGCGCTCGCAATGCgtttaagagctatgctggaaacagc |
| Cntrob | Cntrob-3 | GCTACACTAGCCTTCGGCC | tatcttgtggaaaggacgaaacaccGGCTACACTAGCCTTCGGCCgtttaagagctatgctggaaacagc |
| Cntrob | Cntrob-4 | CGATAGGCAGTATCACGGG | tatcttgtggaaaggacgaaacaccGCGATAGGCAGTATCACGGGgtttaagagctatgctggaaacagc |
| Cop1 | Cop1-1 | TGTCTGGCAGCGCGAGCGC | tatcttgtggaaaggacgaaacaccGTGTCTGGCAGCGCGAGCGCgtttaagagctatgctggaaacagc |
| Cop1 | Cop1-2 | TCCCTCCAACGCCGGCGCT | tatcttgtggaaaggacgaaacaccGTCCCTCCAACGCCGGCGCTgtttaagagctatgctggaaacagc |
| Cop1 | Cop1-4 | GGGCTCCTCAACTCCTACG | tatcttgtggaaaggacgaaacaccGGGGCTCCTCAACTCCTACGgtttaagagctatgctggaaacagc |
| Copb2 | Copb2-3 | TGACCGGCTTGTTAAAATA | tatcttgtggaaaggacgaaacaccGTGACCGGCTTGTTAAAATAgtttaagagctatgctggaaacagc |
| Copb2 | Copb2-4 | AGCACGCTGAATTACGGAA | tatcttgtggaaaggacgaaacaccGAGCACGCTGAATTACGGAAgtttaagagctatgctggaaacagc |
| Copb2 | Copb2-5 | TTGTTGCAGCTTGGTCGGG | tatcttgtggaaaggacgaaacaccGTTGTTGCAGCTTGGTCGGGgtttaagagctatgctggaaacagc |
| Cops4 | Cops4-3 | ATTGCAAGGCTATACCTGG | tatcttgtggaaaggacgaaacaccGATTGCAAGGCTATACCTGGgtttaagagctatgctggaaacagc |
| Cops4 | Cops4-4 | GACGATCCCGTCCAGGCCG | tatcttgtggaaaggacgaaacaccGGACGATCCCGTCCAGGCCGgtttaagagctatgctggaaacagc |
| Cops4 | Cops4-5 | GTTCTGAAGTAAGGACGCT | tatcttgtggaaaggacgaaacaccGGTTCTGAAGTAAGGACGCTgtttaagagctatgctggaaacagc |
| Cops5 | Cops5-2 | CCACCCTGGTTATGGCTGC | tatcttgtggaaaggacgaaacaccGCCACCCTGGTTATGGCTGCgtttaagagctatgctggaaacagc |
| Cops5 | Cops5-4 | TCCAAGTTGCCTCCTGACC | tatcttgtggaaaggacgaaacaccGTCCAAGTTGCCTCCTGACCgtttaagagctatgctggaaacagc |
| Cops5 | Cops5-5 | GCGGCGAAACCCTGGACTA | tatcttgtggaaaggacgaaacaccGGCGGCGAAACCCTGGACTAgtttaagagctatgctggaaacagc |
| Cops6 | Cops6-2 | GAATATTATTACACCAAGG | tatcttgtggaaaggacgaaacaccGGAATATTATTACACCAAGGgtttaagagctatgctggaaacagc |
| Cops6 | Cops6-3 | CTGGGTTGGTATACCACAG | tatcttgtggaaaggacgaaacaccGCTGGGTTGGTATACCACAGgtttaagagctatgctggaaacagc |
| Cops6 | Cops6-4 | CTATACCTGCTTATGGACG | tatcttgtggaaaggacgaaacaccGCTATACCTGCTTATGGACGgtttaagagctatgctggaaacagc |
| Cops8 | Cops8-1 | TACTTACAGACTTTATAGC | tatcttgtggaaaggacgaaacaccGTACTTACAGACTTTATAGCgtttaagagctatgctggaaacagc |
| Cops8 | Cops8-2 | GATGGTTGTATAGATCCCA | tatcttgtggaaaggacgaaacaccGGATGGTTGTATAGATCCCAgtttaagagctatgctggaaacagc |
| Cops8 | Cops8-5 | GGTATAAGCTTGAGAGACC | tatcttgtggaaaggacgaaacaccGGGTATAAGCTTGAGAGACCgtttaagagctatgctggaaacagc |
| Cox7a2 | Cox7a2-3 | CCCCGCCTTTCAGATGAAC | tatcttgtggaaaggacgaaacaccGCCCCGCCTTTCAGATGAACgtttaagagctatgctggaaacagc |
| Cox7a2 | Cox7a2-4 | TTTTAACCAGGAGGATAAT | tatcttgtggaaaggacgaaacaccGTTTTAACCAGGAGGATAATgtttaagagctatgctggaaacagc |
| Cox7a2 | Cox7a2-5 | GAGAAACAAAAGCTGTTTC | tatcttgtggaaaggacgaaacaccGGAGAAACAAAAGCTGTTTCgtttaagagctatgctggaaacagc |
| Cpsf2 | Cpsf2-3 | CTTGTTTAAGCAAACGTCC | tatcttgtggaaaggacgaaacaccGCTTGTTTAAGCAAACGTCCgtttaagagctatgctggaaacagc |
| Cpsf2 | Cpsf2-4 | GGAAACGCTTCGGGGCGAC | tatcttgtggaaaggacgaaacaccGGGAAACGCTTCGGGGCGACgtttaagagctatgctggaaacagc |
| Cpsf2 | Cpsf2-5 | AGCGAGTAAACGCCCAACC | tatcttgtggaaaggacgaaacaccGAGCGAGTAAACGCCCAACCgtttaagagctatgctggaaacagc |
| Cpsf4 | Cpsf4-1 | GTCCATTCCGCCACATCAG | tatcttgtggaaaggacgaaacaccGGTCCATTCCGCCACATCAGgtttaagagctatgctggaaacagc |
| Cpsf4 | Cpsf4-3 | TTTTACTCCAAGTTCGGTA | tatcttgtggaaaggacgaaacaccGTTTTACTCCAAGTTCGGTAgtttaagagctatgctggaaacagc |
| Cpsf4 | Cpsf4-4 | GTGTGCCGATGCCTACACA | tatcttgtggaaaggacgaaacaccGGTGTGCCGATGCCTACACAgtttaagagctatgctggaaacagc |
| Cr2 | Cr2-1 | GACGGATTTCTATAAACCA | tatcttgtggaaaggacgaaacaccGGACGGATTTCTATAAACCAgtttaagagctatgctggaaacagc |
| Cr2 | Cr2-4 | TATGACTTCCATTACGAAC | tatcttgtggaaaggacgaaacaccGTATGACTTCCATTACGAACgtttaagagctatgctggaaacagc |
| Cr2 | Cr2-5 | GTGTCATCCCGACATGCAA | tatcttgtggaaaggacgaaacaccGGTGTCATCCCGACATGCAAgtttaagagctatgctggaaacagc |
| Cramp1 | Cramp1-2 | CTACCCGGTGTAGCCCGTG | tatcttgtggaaaggacgaaacaccGCTACCCGGTGTAGCCCGTGgtttaagagctatgctggaaacagc |
| Cramp1 | Cramp1-4 | GGTGCGGTAGTAAAAATGC | tatcttgtggaaaggacgaaacaccGGGTGCGGTAGTAAAAATGCgtttaagagctatgctggaaacagc |

| | | | |
|---|---|---|---|
| Cramp1 | Cramp1-5 | CGGGCCGCGAGGAAAAGGT | tatcttgtggaaaggacgaaacaccGCGGGCCGCGAGGAAAAGGTgtttaagagctatgctggaaacagc |
| Crebrf | Crebrf-1 | GGATGCTTTAACTTCAAAC | tatcttgtggaaaggacgaaacaccGGGATGCTTTAACTTCAAACgtttaagagctatgctggaaacagc |
| Crebrf | Crebrf-4 | AGGACTAAGTAATAGCTCC | tatcttgtggaaaggacgaaacaccGAGGACTAAGTAATAGCTCCgtttaagagctatgctggaaacagc |
| Crebrf | Crebrf-5 | CAGGACTATCCTGCGTCAG | tatcttgtggaaaggacgaaacaccGCAGGACTATCCTGCGTCAGgtttaagagctatgctggaaacagc |
| Cript | Cript-1 | TTTAATATATAGAGAGTGG | tatcttgtggaaaggacgaaacaccGTTTAATATATAGAGAGTGGgtttaagagctatgctggaaacagc |
| Cript | Cript-2 | TTTGACTTCGAAAAAAGCA | tatcttgtggaaaggacgaaacaccGTTTGACTTCGAAAAAAGCAgtttaagagctatgctggaaacagc |
| Cript | Cript-4 | CTTTTACAAATTCTGCAAG | tatcttgtggaaaggacgaaacaccGCTTTTACAAATTCTGCAAGgtttaagagctatgctggaaacagc |
| Csn1s2b | Csn1s2b-1 | ACTTCTTGAGAGTTATCCA | tatcttgtggaaaggacgaaacaccGACTTCTTGAGAGTTATCCAgtttaagagctatgctggaaacagc |
| Csn1s2b | Csn1s2b-2 | TAGAACTTTAAGCAGAATA | tatcttgtggaaaggacgaaacaccGTAGAACTTTAAGCAGAATAgtttaagagctatgctggaaacagc |
| Csn1s2b | Csn1s2b-5 | CAATACAACCAGAAAATGA | tatcttgtggaaaggacgaaacaccGCAATACAACCAGAAAATGAgtttaagagctatgctggaaacagc |
| Csn3 | Csn3-1 | ATCAACTTACGTTTGAATC | tatcttgtggaaaggacgaaacaccGATCAACTTACGTTTGAATCgtttaagagctatgctggaaacagc |
| Csn3 | Csn3-2 | ATTGAAGTTCAGGACGGAG | tatcttgtggaaaggacgaaacaccGATTGAAGTTCAGGACGGAGgtttaagagctatgctggaaacagc |
| Csn3 | Csn3-4 | TGTATGGGCTAGCAGTAGC | tatcttgtggaaaggacgaaacaccGTGTATGGGCTAGCAGTAGCgtttaagagctatgctggaaacagc |
| Csnk1a1 | Csnk1a1-1 | CGGGTGATTACCTCGCCAT | tatcttgtggaaaggacgaaacaccGCGGGTGATTACCTCGCCATgtttaagagctatgctggaaacagc |
| Csnk1a1 | Csnk1a1-2 | TGCTCTCGTACAGCAACTG | tatcttgtggaaaggacgaaacaccGTGCTCTCGTACAGCAACTGgtttaagagctatgctggaaacagc |
| Csnk1a1 | Csnk1a1-4 | GTACTTATGTTAGCTGACC | tatcttgtggaaaggacgaaacaccGGTACTTATGTTAGCTGACCgtttaagagctatgctggaaacagc |
| Cstf1 | Cstf1-2 | GCTCTGACACGGTTGCCCC | tatcttgtggaaaggacgaaacaccGGCTCTGACACGGTTGCCCCgtttaagagctatgctggaaacagc |
| Cstf1 | Cstf1-4 | CGTAAAGAGTTCGGATCAC | tatcttgtggaaaggacgaaacaccGCGTAAAGAGTTCGGATCACgtttaagagctatgctggaaacagc |
| Cstf1 | Cstf1-5 | GTATAATCCCTTGAGCCCG | tatcttgtggaaaggacgaaacaccGGTATAATCCCTTGAGCCCGgtttaagagctatgctggaaacagc |
| Ctc1 | Ctc1-2 | TGGTCGAATAACCCGCCTG | tatcttgtggaaaggacgaaacaccGTGGTCGAATAACCCGCCTGgtttaagagctatgctggaaacagc |
| Ctc1 | Ctc1-3 | TTGAAGCCGAACAGTGCCA | tatcttgtggaaaggacgaaacaccGTTGAAGCCGAACAGTGCCAgtttaagagctatgctggaaacagc |
| Ctc1 | Ctc1-5 | GCTGTACGCATGCCGCATG | tatcttgtggaaaggacgaaacaccGGCTGTACGCATGCCGCATGgtttaagagctatgctggaaacagc |
| Cyp2a22 | Cyp2a22-1 | TGGACCCCAATTCCCCACG | tatcttgtggaaaggacgaaacaccGTGGACCCCAATTCCCCACGgtttaagagctatgctggaaacagc |
| Cyp2a22 | Cyp2a22-2 | TGTGGGTGAAGCAGCAAAT | tatcttgtggaaaggacgaaacaccGTGTGGGTGAAGCAGCAAATgtttaagagctatgctggaaacagc |
| Cyp2a22 | Cyp2a22-4 | TTCTCCATAGCCACACTGA | tatcttgtggaaaggacgaaacaccGTTCTCCATAGCCACACTGAgtttaagagctatgctggaaacagc |
| Cyp2b9 | Cyp2b9-1 | ACTATGGGCAATGTAGTCG | tatcttgtggaaaggacgaaacaccGACTATGGGCAATGTAGTCGgtttaagagctatgctggaaacagc |
| Cyp2b9 | Cyp2b9-2 | CAAGCGCCACCCTCCACTA | tatcttgtggaaaggacgaaacaccGCAAGCGCCACCCTCCACTAgtttaagagctatgctggaaacagc |
| Cyp2b9 | Cyp2b9-3 | TGTGGCATACCTGTGACAT | tatcttgtggaaaggacgaaacaccGTGTGGCATACCTGTGACATgtttaagagctatgctggaaacagc |
| Cyp2c55 | Cyp2c55-1 | GTTTCCAATAATTGGGAAA | tatcttgtggaaaggacgaaacaccGGTTTCCAATAATTGGGAAAgtttaagagctatgctggaaacagc |
| Cyp2c55 | Cyp2c55-2 | CCTCATCAATTATCTCCCA | tatcttgtggaaaggacgaaacaccGCCTCATCAATTATCTCCCAgtttaagagctatgctggaaacagc |
| Cyp2c55 | Cyp2c55-3 | CAATAGTAAACTCCGAATG | tatcttgtggaaaggacgaaacaccGCAATAGTAAACTCCGAATGgtttaagagctatgctggaaacagc |
| Cyp2g1 | Cyp2g1-2 | TAGTGTAGGCATTTCACCA | tatcttgtggaaaggacgaaacaccGTAGTGTAGGCATTTCACCAgtttaagagctatgctggaaacagc |
| Cyp2g1 | Cyp2g1-3 | ATGCTACTTACCATAACCT | tatcttgtggaaaggacgaaacaccGATGCTACTTACCATAACCTgtttaagagctatgctggaaacagc |
| Cyp2g1 | Cyp2g1-5 | ACCACGGAACAGATGACGT | tatcttgtggaaaggacgaaacaccGACCACGGAACAGATGACGTgtttaagagctatgctggaaacagc |
| Cyp2j11 | Cyp2j11-2 | GAGCAGTGTATACAAGAAG | tatcttgtggaaaggacgaaacaccGGAGCAGTGTATACAAGAAGgtttaagagctatgctggaaacagc |
| Cyp2j11 | Cyp2j11-4 | TAATCAAAGAAGCTATCAC | tatcttgtggaaaggacgaaacaccGTAATCAAAGAAGCTATCACgtttaagagctatgctggaaacagc |
| Cyp2j11 | Cyp2j11-5 | TGACACATCTAAATCAAAC | tatcttgtggaaaggacgaaacaccGTGACACATCTAAATCAAACgtttaagagctatgctggaaacagc |
| Cyp7b1 | Cyp7b1-1 | GTATGTCAGATACTAAGTA | tatcttgtggaaaggacgaaacaccGGTATGTCAGATACTAAGTAgtttaagagctatgctggaaacagc |
| Cyp7b1 | Cyp7b1-2 | GAGCCTATCTACTTCTACA | tatcttgtggaaaggacgaaacaccGGAGCCTATCTACTTCTACAgtttaagagctatgctggaaacagc |
| Cyp7b1 | Cyp7b1-4 | TATCAAGGGTGGTTCACGA | tatcttgtggaaaggacgaaacaccGTATCAAGGGTGGTTCACGAgtttaagagctatgctggaaacagc |
| Cysltr2 | Cysltr2-3 | AAGTCCCCAAATATCCAAT | tatcttgtggaaaggacgaaacaccGAAGTCCCCAAATATCCAATgtttaagagctatgctggaaacagc |
| Cysltr2 | Cysltr2-4 | TTATTTCAGAGGTTCCAAT | tatcttgtggaaaggacgaaacaccGTTATTTCAGAGGTTCCAATgtttaagagctatgctggaaacagc |
| Cysltr2 | Cysltr2-5 | ATGGAAAAGCCATTTCCCA | tatcttgtggaaaggacgaaacaccGATGGAAAAGCCATTTCCCAgtttaagagctatgctggaaacagc |
| Dbr1 | Dbr1-1 | TTCCAGGCGGTGCGCAACG | tatcttgtggaaaggacgaaacaccGTTCCAGGCGGTGCGCAACGgtttaagagctatgctggaaacagc |
| Dbr1 | Dbr1-2 | GCGCTACCCAGCCACCATA | tatcttgtggaaaggacgaaacaccGGCGCTACCCAGCCACCATAgtttaagagctatgctggaaacagc |
| Dbr1 | Dbr1-4 | GGTAGATGAATTATAAGGG | tatcttgtggaaaggacgaaacaccGGGTAGATGAATTATAAGGGgtttaagagctatgctggaaacagc |
| Dclre1c | Dclre1c-2 | CAGGGAAACATACACGACT | tatcttgtggaaaggacgaaacaccGCAGGGAAACATACACGACTgtttaagagctatgctggaaacagc |
| Dclre1c | Dclre1c-4 | TTATTCACCAACCTAAGCG | tatcttgtggaaaggacgaaacaccGTTATTCACCAACCTAAGCGgtttaagagctatgctggaaacagc |
| Dclre1c | Dclre1c-5 | GTCCGTTGTGAGATGGTGC | tatcttgtggaaaggacgaaacaccGGTCCGTTGTGAGATGGTGCgtttaagagctatgctggaaacagc |
| Dctn6 | Dctn6-1 | GTCTTGGCTGCAGCCTACG | tatcttgtggaaaggacgaaacaccGGTCTTGGCTGCAGCCTACGgtttaagagctatgctggaaacagc |
| Dctn6 | Dctn6-2 | ATTCCCAAGCCATGAAAAT | tatcttgtggaaaggacgaaacaccGATTCCCAAGCCATGAAAATgtttaagagctatgctggaaacagc |
| Dctn6 | Dctn6-3 | CCAACAACGTATTTGAAGT | tatcttgtggaaaggacgaaacaccGCCAACAACGTATTTGAAGTgtttaagagctatgctggaaacagc |
| Ddb1 | Ddb1-1 | GACCGCATTGGTCGCCCCT | tatcttgtggaaaggacgaaacaccGGACCGCATTGGTCGCCCCTgtttaagagctatgctggaaacagc |
| Ddb1 | Ddb1-3 | AAGGCCATCGTAGAGTCGA | tatcttgtggaaaggacgaaacaccGAAGGCCATCGTAGAGTCGAgtttaagagctatgctggaaacagc |
| Ddb1 | Ddb1-4 | CCGTTCCGGATGATCCGCA | tatcttgtggaaaggacgaaacaccGCCGTTCCGGATGATCCGCAgtttaagagctatgctggaaacagc |
| Ddx19a | Ddx19a-2 | ATGAGCTGGCATTACAAAC | tatcttgtggaaaggacgaaacaccGATGAGCTGGCATTACAAACgtttaagagctatgctggaaacagc |
| Ddx19a | Ddx19a-4 | CACGGTTCAGCAAGCATCA | tatcttgtggaaaggacgaaacaccGCACGGTTCAGCAAGCATCAgtttaagagctatgctggaaacagc |
| Ddx19a | Ddx19a-5 | TCCAAGGAGTCTACGCCAT | tatcttgtggaaaggacgaaacaccGTCCAAGGAGTCTACGCCATgtttaagagctatgctggaaacagc |
| Ddx43 | Ddx43-2 | TCACCCCGAAGCAGAGCGG | tatcttgtggaaaggacgaaacaccGTCACCCCGAAGCAGAGCGGgtttaagagctatgctggaaacagc |
| Ddx43 | Ddx43-3 | ACAACGAACACTAGGATAC | tatcttgtggaaaggacgaaacaccGACAACGAACACTAGGATACgtttaagagctatgctggaaacagc |
| Ddx43 | Ddx43-5 | GTTAGTTCTTACGCCCACC | tatcttgtggaaaggacgaaacaccGGTTAGTTCTTACGCCCACCgtttaagagctatgctggaaacagc |
| Ddx47 | Ddx47-1 | AGCTTGTGACCAGTTGGGA | tatcttgtggaaaggacgaaacaccGAGCTTGTGACCAGTTGGGAgtttaagagctatgctggaaacagc |
| Ddx47 | Ddx47-2 | CAAAGGAATAGCTTCGATC | tatcttgtggaaaggacgaaacaccGCAAAGGAATAGCTTCGATCgtttaagagctatgctggaaacagc |
| Ddx47 | Ddx47-3 | CCAGGGCGAACAGTCGCTG | tatcttgtggaaaggacgaaacaccGCCAGGGCGAACAGTCGCTGgtttaagagctatgctggaaacagc |
| Ddx55 | Ddx55-1 | GCGGGACCGATACCTGCAC | tatcttgtggaaaggacgaaacaccGGCGGGACCGATACCTGCACgtttaagagctatgctggaaacagc |
| Ddx55 | Ddx55-2 | ATGGCCAGCTCGCGAGTCG | tatcttgtggaaaggacgaaacaccGATGGCCAGCTCGCGAGTCGgtttaagagctatgctggaaacagc |
| Ddx55 | Ddx55-4 | GATTCGGACAGGGTTCCGG | tatcttgtggaaaggacgaaacaccGGATTCGGACAGGGTTCCGGgtttaagagctatgctggaaacagc |
| Def6 | Def6-1 | GACAAGTACCCGCTGATCA | tatcttgtggaaaggacgaaacaccGGACAAGTACCCGCTGATCAgtttaagagctatgctggaaacagc |
| Def6 | Def6-3 | AGGTGCTTCCGGACCGCGA | tatcttgtggaaaggacgaaacaccGAGGTGCTTCCGGACCGCGAgtttaagagctatgctggaaacagc |
| Def6 | Def6-5 | AGCCGGATCGCAGTCTGGA | tatcttgtggaaaggacgaaacaccGAGCCGGATCGCAGTCTGGAgtttaagagctatgctggaaacagc |
| Defb14 | Defb14-1 | TCTGCAGAAAAATTTTCGG | tatcttgtggaaaggacgaaacaccGTCTGCAGAAAAATTTTCGGgtttaagagctatgctggaaacagc |
| Defb14 | Defb14-2 | CTGCAGAATAAGAGGTGGC | tatcttgtggaaaggacgaaacaccGCTGCAGAATAAGAGGTGGCgtttaagagctatgctggaaacagc |
| Defb14 | Defb14-4 | TATTTGTTCTTCCTTTCCA | tatcttgtggaaaggacgaaacaccGTATTTGTTCTTCCTTTCCAgtttaagagctatgctggaaacagc |
| Defb7 | Defb7-1 | AACGAGCTTGCTATCGGGA | tatcttgtggaaaggacgaaacaccGAACGAGCTTGCTATCGGGAgtttaagagctatgctggaaacagc |
| Defb7 | Defb7-2 | AATGCCTGCAACGGTGCAT | tatcttgtggaaaggacgaaacaccGAATGCCTGCAACGGTGCATgtttaagagctatgctggaaacagc |
| Defb7 | Defb7-3 | AGTTCCAATCTTATGAAAA | tatcttgtggaaaggacgaaacaccGAGTTCCAATCTTATGAAAAgtttaagagctatgctggaaacagc |

| | | | |
|---|---|---|---|
| Dek | Dek-1 | GAACAGCGAACTCGTGAAG | tatcttgtggaaaggacgaaacaccGGAACAGCGAACTCGTGAAGgtttaagagctatgctggaaacagc |
| Dek | Dek-2 | AGTACCCAGTATAAAAAGA | tatcttgtggaaaggacgaaacaccGAGTACCCAGTATAAAAAGAgtttaagagctatgctggaaacagc |
| Dek | Dek-5 | CACCTTGTGTCACTGTAAA | tatcttgtggaaaggacgaaacaccGCACCTTGTGTCACTGTAAAgtttaagagctatgctggaaacagc |
| Desi2 | Desi2-1 | TTTCAGAGTTTGCTTATGG | tatcttgtggaaaggacgaaacaccGTTTCAGAGTTTGCTTATGGgtttaagagctatgctggaaacagc |
| Desi2 | Desi2-3 | GCTGTTGTTCTAGGAAGTA | tatcttgtggaaaggacgaaacaccGGCTGTTGTTCTAGGAAGTAgtttaagagctatgctggaaacagc |
| Desi2 | Desi2-4 | CCTCGCTGGATCAACCGGC | tatcttgtggaaaggacgaaacaccGCCTCGCTGGATCAACCGGCgtttaagagctatgctggaaacagc |
| Dhps | Dhps-1 | AAGGCTACGACTTCAACCG | tatcttgtggaaaggacgaaacaccGAAGGCTACGACTTCAACCGgtttaagagctatgctggaaacagc |
| Dhps | Dhps-3 | CACCATGTTGTGCTGCACG | tatcttgtggaaaggacgaaacaccGCACCATGTTGTGCTGCACGgtttaagagctatgctggaaacagc |
| Dhps | Dhps-5 | CCGGGAGAGTGGGATCAAC | tatcttgtggaaaggacgaaacaccGCCGGGAGAGTGGGATCAACgtttaagagctatgctggaaacagc |
| Dhx37 | Dhx37-1 | CTTCCTAAGCCGACGGCAA | tatcttgtggaaaggacgaaacaccGCTTCCTAAGCCGACGGCAAgtttaagagctatgctggaaacagc |
| Dhx37 | Dhx37-3 | CTACTGCGGCCACTCGGCG | tatcttgtggaaaggacgaaacaccGCTACTGCGGCCACTCGGCGgtttaagagctatgctggaaacagc |
| Dhx37 | Dhx37-4 | AAGCTCCCGATCCTCGCCG | tatcttgtggaaaggacgaaacaccGAAGCTCCCGATCCTCGCCGgtttaagagctatgctggaaacagc |
| Dmpk | Dmpk-2 | CGGTTCCGTATTTAGGTAG | tatcttgtggaaaggacgaaacaccGCGGTTCCGTATTTAGGTAGgtttaagagctatgctggaaacagc |
| Dmpk | Dmpk-3 | CATGGAATACTACGTGGGC | tatcttgtggaaaggacgaaacaccGCATGGAATACTACGTGGGCgtttaagagctatgctggaaacagc |
| Dmpk | Dmpk-4 | ACGTAGCCCAGCCGGTGCA | tatcttgtggaaaggacgaaacaccGACGTAGCCCAGCCGGTGCAgtttaagagctatgctggaaacagc |
| Dnaaf6b | Dnaaf6b-1 | GATTTTGATTTTGAACAGG | tatcttgtggaaaggacgaaacaccGGATTTTGATTTTGAACAGGgtttaagagctatgctggaaacagc |
| Dnaaf6b | Dnaaf6b-4 | CTAAAGCAAAAGAGTCCAA | tatcttgtggaaaggacgaaacaccGCTAAAGCAAAAGAGTCCAAgtttaagagctatgctggaaacagc |
| Dnaaf6b | Dnaaf6b-5 | ACATATACCTAGGATTGAC | tatcttgtggaaaggacgaaacaccGACATATACCTAGGATTGACgtttaagagctatgctggaaacagc |
| Dnm2 | Dnm2-1 | ATCAGGAATTGTCACCCGG | tatcttgtggaaaggacgaaacaccGATCAGGAATTGTCACCCGGgtttaagagctatgctggaaacagc |
| Dnm2 | Dnm2-3 | TTTCAGGCCTACGGACCAT | tatcttgtggaaaggacgaaacaccGTTTCAGGCCTACGGACCATgtttaagagctatgctggaaacagc |
| Dnm2 | Dnm2-5 | TGAAAGATACGATTGATGC | tatcttgtggaaaggacgaaacaccGTGAAAGATACGATTGATGCgtttaagagctatgctggaaacagc |
| Dohh | Dohh-1 | CTATCTCGTGGATCAGCCG | tatcttgtggaaaggacgaaacaccGCTATCTCGTGGATCAGCCGgtttaagagctatgctggaaacagc |
| Dohh | Dohh-3 | CGGGTTACCAATCGCCCCC | tatcttgtggaaaggacgaaacaccGCGGGTTACCAATCGCCCCCgtttaagagctatgctggaaacagc |
| Dohh | Dohh-4 | CGTAGAATACTGCTTCAGG | tatcttgtggaaaggacgaaacaccGCGTAGAATACTGCTTCAGGgtttaagagctatgctggaaacagc |
| Dtl | Dtl-1 | TTACCTAGCTTTCGAGTGC | tatcttgtggaaaggacgaaacaccGTTACCTAGCTTTCGAGTGCgtttaagagctatgctggaaacagc |
| Dtl | Dtl-3 | TTAAGTTCACCAGGGACCC | tatcttgtggaaaggacgaaacaccGTTAAGTTCACCAGGGACCCgtttaagagctatgctggaaacagc |
| Dtl | Dtl-5 | ACACGTCTTATGGAGAAAC | tatcttgtggaaaggacgaaacaccGACACGTCTTATGGAGAAACgtttaagagctatgctggaaacagc |
| Dut | Dut-1 | CGGCTCTCGGAGCACGCCA | tatcttgtggaaaggacgaaacaccGCGGCTCTCGGAGCACGCCAgtttaagagctatgctggaaacagc |
| Dut | Dut-4 | TCACGATGGCTTTCTCCAT | tatcttgtggaaaggacgaaacaccGTCACGATGGCTTTCTCCATgtttaagagctatgctggaaacagc |
| Dut | Dut-5 | CTCTTCCATAGCACCCAGA | tatcttgtggaaaggacgaaacaccGCTCTTCCATAGCACCCAGAgtttaagagctatgctggaaacagc |
| E130311K13Rik | E130311K13Rik-3 | CTTTCCATGAAGATATGAA | tatcttgtggaaaggacgaaacaccGCTTTCCATGAAGATATGAAgtttaagagctatgctggaaacagc |
| E130311K13Rik | E130311K13Rik-4 | ACTACGTGGCCGATTACAG | tatcttgtggaaaggacgaaacaccGACTACGTGGCCGATTACAGgtttaagagctatgctggaaacagc |
| E130311K13Rik | E130311K13Rik-5 | GTTGCTGATAAGGAGCGTC | tatcttgtggaaaggacgaaacaccGGTTGCTGATAAGGAGCGTCgtttaagagctatgctggaaacagc |
| Ebna1bp2 | Ebna1bp2-2 | GCTGAATTCAGACGGGATC | tatcttgtggaaaggacgaaacaccGGCTGAATTCAGACGGGATCgtttaagagctatgctggaaacagc |
| Ebna1bp2 | Ebna1bp2-4 | TGTAATGTCAGCTACCGGC | tatcttgtggaaaggacgaaacaccGTGTAATGTCAGCTACCGGCgtttaagagctatgctggaaacagc |
| Ebna1bp2 | Ebna1bp2-5 | GCTCCAAGTCCCTACGAAG | tatcttgtggaaaggacgaaacaccGGCTCCAAGTCCCTACGAAGgtttaagagctatgctggaaacagc |
| Ecd | Ecd-3 | CATTTCGTGAAAGTGACCT | tatcttgtggaaaggacgaaacaccGCATTTCGTGAAAGTGACCTgtttaagagctatgctggaaacagc |
| Ecd | Ecd-4 | TAGGTCAATGGGGTCCCGT | tatcttgtggaaaggacgaaacaccGTAGGTCAATGGGGTCCCGTgtttaagagctatgctggaaacagc |
| Ecd | Ecd-5 | GACCAGTATGGGTGCGAAC | tatcttgtggaaaggacgaaacaccGGACCAGTATGGGTGCGAACgtttaagagctatgctggaaacagc |
| Ect2 | Ect2-2 | TTCAGATGGACGCTCGTGC | tatcttgtggaaaggacgaaacaccGTTCAGATGGACGCTCGTGCgtttaagagctatgctggaaacagc |
| Ect2 | Ect2-4 | CGTTGGTTCATCATATGGG | tatcttgtggaaaggacgaaacaccGCGTTGGTTCATCATATGGGgtttaagagctatgctggaaacagc |
| Ect2 | Ect2-5 | GTACAATACAGCGGCCGGC | tatcttgtggaaaggacgaaacaccGGTACAATACAGCGGCCGGCgtttaagagctatgctggaaacagc |
| Eed | Eed-2 | TCCAGCAACTGCTAATAGA | tatcttgtggaaaggacgaaacaccGTCCAGCAACTGCTAATAGAgtttaagagctatgctggaaacagc |
| Eed | Eed-3 | CCGTATCTCCCCCTGTGAA | tatcttgtggaaaggacgaaacaccGCCGTATCTCCCCCTGTGAAgtttaagagctatgctggaaacagc |
| Eed | Eed-4 | TACGCCAAATGCACCAGGA | tatcttgtggaaaggacgaaacaccGTACGCCAAATGCACCAGGAgtttaagagctatgctggaaacagc |
| Eef2 | Eef2-1 | CGTGTCACCGATGGAGCTC | tatcttgtggaaaggacgaaacaccGCGTGTCACCGATGGAGCTCgtttaagagctatgctggaaacagc |
| Eef2 | Eef2-4 | GGTACTTTGATCCGGCCAA | tatcttgtggaaaggacgaaacaccGGGTACTTTGATCCGGCCAAgtttaagagctatgctggaaacagc |
| Eef2 | Eef2-5 | TGGCGGCCTCATCGTCGGG | tatcttgtggaaaggacgaaacaccGTGGCGGCCTCATCGTCGGGgtttaagagctatgctggaaacagc |
| Efnb3 | Efnb3-1 | CATCAGACGGGACCCGGGA | tatcttgtggaaaggacgaaacaccGCATCAGACGGGACCCGGGAgtttaagagctatgctggaaacagc |
| Efnb3 | Efnb3-2 | ACCAATTATGTAGTAATCG | tatcttgtggaaaggacgaaacaccGACCAATTATGTAGTAATCGgtttaagagctatgctggaaacagc |
| Efnb3 | Efnb3-5 | CTGGAACTCGGCGAATAAG | tatcttgtggaaaggacgaaacaccGCTGGAACTCGGCGAATAAGgtttaagagctatgctggaaacagc |
| Ehd3 | Ehd3-2 | CCTCGAATGGTTTGCTGAG | tatcttgtggaaaggacgaaacaccGCCTCGAATGGTTTGCTGAGgtttaagagctatgctggaaacagc |
| Ehd3 | Ehd3-4 | GTATACTCGCATCAGCTGC | tatcttgtggaaaggacgaaacaccGGTATACTCGCATCAGCTGCgtttaagagctatgctggaaacagc |
| Ehd3 | Ehd3-5 | AACACCCCCGAGGTGATCC | tatcttgtggaaaggacgaaacaccGAACACCCCCGAGGTGATCCgtttaagagctatgctggaaacagc |
| Eif2b2 | Eif2b2-1 | GTGGTGGTCCGTGATAAGC | tatcttgtggaaaggacgaaacaccGGTGGTGGTCCGTGATAAGCgtttaagagctatgctggaaacagc |
| Eif2b2 | Eif2b2-2 | CGCACCCGCGTTGTTCCAG | tatcttgtggaaaggacgaaacaccGCGCACCCGCGTTGTTCCAGgtttaagagctatgctggaaacagc |
| Eif2b2 | Eif2b2-4 | CGCACCATATTTCCCACGG | tatcttgtggaaaggacgaaacaccGCGCACCATATTTCCCACGGgtttaagagctatgctggaaacagc |
| Eif3a | Eif3a-1 | TACTATGGCCCGTTCCAGT | tatcttgtggaaaggacgaaacaccGTACTATGGCCCGTTCCAGTgtttaagagctatgctggaaacagc |
| Eif3a | Eif3a-3 | GAGCAATATCAGTTCGCTC | tatcttgtggaaaggacgaaacaccGGAGCAATATCAGTTCGCTCgtttaagagctatgctggaaacagc |
| Eif3a | Eif3a-5 | TTGGGCGATATCATGGTAA | tatcttgtggaaaggacgaaacaccGTTGGGCGATATCATGGTAAgtttaagagctatgctggaaacagc |
| Eif3f | Eif3f-3 | GATCGAGGCCAAAATGACG | tatcttgtggaaaggacgaaacaccGGATCGAGGCCAAAATGACGgtttaagagctatgctggaaacagc |
| Eif3f | Eif3f-4 | AGCTACGAACGCCGCAACG | tatcttgtggaaaggacgaaacaccGAGCTACGAACGCCGCAACGgtttaagagctatgctggaaacagc |
| Eif3f | Eif3f-5 | GCTGTTGCAGGTATGCCAC | tatcttgtggaaaggacgaaacaccGGCTGTTGCAGGTATGCCACgtttaagagctatgctggaaacagc |
| Eif3g | Eif3g-1 | GGGGCAACGGGTGGTCCAA | tatcttgtggaaaggacgaaacaccGGGGGCAACGGGTGGTCCAAgtttaagagctatgctggaaacagc |
| Eif3g | Eif3g-2 | TCGCCCTTGCAAATGCGGC | tatcttgtggaaaggacgaaacaccGTCGCCCTTGCAAATGCGGCgtttaagagctatgctggaaacagc |
| Eif3g | Eif3g-5 | TTCCGGGTCTCAATCCTGA | tatcttgtggaaaggacgaaacaccGTTCCGGGTCTCAATCCTGAgtttaagagctatgctggaaacagc |
| Eif3i | Eif3i-1 | TCATCGCAGGCCACGAGAG | tatcttgtggaaaggacgaaacaccGTCATCGCAGGCCACGAGAGgtttaagagctatgctggaaacagc |
| Eif3i | Eif3i-2 | GCCGTCTGGGGCCCCCTCG | tatcttgtggaaaggacgaaacaccGGCCGTCTGGGGCCCCCTCGgtttaagagctatgctggaaacagc |
| Eif3i | Eif3i-3 | TGCGGGATCCAAGCCAGAT | tatcttgtggaaaggacgaaacaccGTGCGGGATCCAAGCCAGATgtttaagagctatgctggaaacagc |
| Eif3m | Eif3m-3 | AGTACACTGTGTACCTCAC | tatcttgtggaaaggacgaaacaccGAGTACACTGTGTACCTCACgtttaagagctatgctggaaacagc |
| Eif3m | Eif3m-4 | ACAACTGTAGTCTCAGAGA | tatcttgtggaaaggacgaaacaccGACAACTGTAGTCTCAGAGAgtttaagagctatgctggaaacagc |
| Eif3m | Eif3m-5 | GGTTCCAGGATCAACAGGA | tatcttgtggaaaggacgaaacaccGGGTTCCAGGATCAACAGGAgtttaagagctatgctggaaacagc |
| Eif5a | Eif5a-1 | TATCCCTGCTCCAGGACAG | tatcttgtggaaaggacgaaacaccGTATCCCTGCTCCAGGACAGgtttaagagctatgctggaaacagc |
| Eif5a | Eif5a-2 | CAGGGATAGGTACCCATCC | tatcttgtggaaaggacgaaacaccGCAGGGATAGGTACCCATCCgtttaagagctatgctggaaacagc |
| Eif5a | Eif5a-4 | GGTGATTCTAACCTTGGCA | tatcttgtggaaaggacgaaacaccGGGTGATTCTAACCTTGGCAgtttaagagctatgctggaaacagc |
| Elmo2 | Elmo2-2 | AGCGCGTTAATCAGAGCAA | tatcttgtggaaaggacgaaacaccGAGCGCGTTAATCAGAGCAAgtttaagagctatgctggaaacagc |

| | | | |
|---|---|---|---|
| Elmo2 | Elmo2-3 | ATAGCGGAGGAGATCACCG | tatcttgtggaaaggacgaaacaccGATAGCGGAGGAGATCACCGgtttaagagctatgctggaaacagc |
| Elmo2 | Elmo2-5 | GCGGAGGGTGTAGTACTCC | tatcttgtggaaaggacgaaacaccGGCGGAGGGTGTAGTACTCCgtttaagagctatgctggaaacagc |
| Elmod1 | Elmod1-1 | CGAAGCAGTGGTGTGAGAT | tatcttgtggaaaggacgaaacaccGCGAAGCAGTGGTGTGAGATgtttaagagctatgctggaaacagc |
| Elmod1 | Elmod1-4 | CTTCCAGAACCATGAAAAT | tatcttgtggaaaggacgaaacaccGCTTCCAGAACCATGAAAATgtttaagagctatgctggaaacagc |
| Elmod1 | Elmod1-5 | TTTGCTACGGCACCAAACC | tatcttgtggaaaggacgaaacaccGTTTGCTACGGCACCAAACCgtttaagagctatgctggaaacagc |
| Epn2 | Epn2-1 | ACTCCAGGAAGAACGCCTC | tatcttgtggaaaggacgaaacaccGACTCCAGGAAGAACGCCTCgtttaagagctatgctggaaacagc |
| Epn2 | Epn2-3 | GAGTAGCTGGTAGAAAGGT | tatcttgtggaaaggacgaaacaccGGAGTAGCTGGTAGAAAGGTgtttaagagctatgctggaaacagc |
| Epn2 | Epn2-4 | CTCTCGAACGTTAATACCC | tatcttgtggaaaggacgaaacaccGCTCTCGAACGTTAATACCCgtttaagagctatgctggaaacagc |
| Ercc2 | Ercc2-1 | CCGCTGATAGGCCACAATC | tatcttgtggaaaggacgaaacaccGCCGCTGATAGGCCACAATCgtttaagagctatgctggaaacagc |
| Ercc2 | Ercc2-2 | AAGCTGCCGTTCTTAGGAC | tatcttgtggaaaggacgaaacaccGAAGCTGCCGTTCTTAGGACgtttaagagctatgctggaaacagc |
| Ercc2 | Ercc2-5 | AAGTGCTCAGCCGTACGGA | tatcttgtggaaaggacgaaacaccGAAGTGCTCAGCCGTACGGAgtttaagagctatgctggaaacagc |
| Erich3 | Erich3-1 | GAACCTTTGTGTCTACAAA | tatcttgtggaaaggacgaaacaccGGAACCTTTGTGTCTACAAAgtttaagagctatgctggaaacagc |
| Erich3 | Erich3-2 | CGCTTGGAGATAAACTGAA | tatcttgtggaaaggacgaaacaccGCGCTTGGAGATAAACTGAAgtttaagagctatgctggaaacagc |
| Erich3 | Erich3-5 | AGGACAACGCTGTTAAATA | tatcttgtggaaaggacgaaacaccGAGGACAACGCTGTTAAATAgtttaagagctatgctggaaacagc |
| Erich5 | Erich5-3 | TGGCAAATGATGAGACCCC | tatcttgtggaaaggacgaaacaccGTGGCAAATGATGAGACCCCgtttaagagctatgctggaaacagc |
| Erich5 | Erich5-4 | ACATATCATCTTTGCCTCC | tatcttgtggaaaggacgaaacaccGACATATCATCTTTGCCTCCgtttaagagctatgctggaaacagc |
| Erich5 | Erich5-5 | AGCATCTCCTTTTAAAGAC | tatcttgtggaaaggacgaaacaccGAGCATCTCCTTTTAAAGACgtttaagagctatgctggaaacagc |
| Esrrg | Esrrg-1 | TGGCGTCGGAAGACCCACC | tatcttgtggaaaggacgaaacaccGTGGCGTCGGAAGACCCACCgtttaagagctatgctggaaacagc |
| Esrrg | Esrrg-3 | CATCATACAGTTTCCGGAC | tatcttgtggaaaggacgaaacaccGCATCATACAGTTTCCGGACgtttaagagctatgctggaaacagc |
| Esrrg | Esrrg-5 | ACATGATGCAACCCCATAG | tatcttgtggaaaggacgaaacaccGACATGATGCAACCCCATAGgtttaagagctatgctggaaacagc |
| Etfrf1 | Etfrf1-1 | AGGAGCAGACTATTTTAAA | tatcttgtggaaaggacgaaacaccGAGGAGCAGACTATTTTAAAgtttaagagctatgctggaaacagc |
| Etfrf1 | Etfrf1-3 | TCAAAGAACTTATCGCACG | tatcttgtggaaaggacgaaacaccGTCAAAGAACTTATCGCACGgtttaagagctatgctggaaacagc |
| Etfrf1 | Etfrf1-4 | CGAGGAGAATTTGTAATGA | tatcttgtggaaaggacgaaacaccGCGAGGAGAATTTGTAATGAgtttaagagctatgctggaaacagc |
| Exosc4 | Exosc4-1 | GCGTGTTTGCGCAGGCCGA | tatcttgtggaaaggacgaaacaccGGCGTGTTTGCGCAGGCCGAgtttaagagctatgctggaaacagc |
| Exosc4 | Exosc4-3 | GTGGTCTACGGGCCGCACG | tatcttgtggaaaggacgaaacaccGGTGGTCTACGGGCCGCACGgtttaagagctatgctggaaacagc |
| Exosc4 | Exosc4-5 | GATGTCGATCTGAGAGCGG | tatcttgtggaaaggacgaaacaccGGATGTCGATCTGAGAGCGGgtttaagagctatgctggaaacagc |
| Exosc8 | Exosc8-1 | ATCGTTACATCAAGCTCAC | tatcttgtggaaaggacgaaacaccGATCGTTACATCAAGCTCACgtttaagagctatgctggaaacagc |
| Exosc8 | Exosc8-2 | CCGACATATCCTCTATCGG | tatcttgtggaaaggacgaaacaccGCCGACATATCCTCTATCGGgtttaagagctatgctggaaacagc |
| Exosc8 | Exosc8-4 | CCGCTGTACTGATTGAACC | tatcttgtggaaaggacgaaacaccGCCGCTGTACTGATTGAACCgtttaagagctatgctggaaacagc |
| Faap100 | Faap100-2 | TGCGTGAGTATTGCCTCCC | tatcttgtggaaaggacgaaacaccGTGCGTGAGTATTGCCTCCCgtttaagagctatgctggaaacagc |
| Faap100 | Faap100-3 | AAGATCCTCCACCACCTAG | tatcttgtggaaaggacgaaacaccGAAGATCCTCCACCACCTAGgtttaagagctatgctggaaacagc |
| Faap100 | Faap100-5 | TAGACGAACTCGCGTCCGG | tatcttgtggaaaggacgaaacaccGTAGACGAACTCGCGTCCGGgtttaagagctatgctggaaacagc |
| Faf2 | Faf2-3 | GGTCAGTGACCCGGCTGCG | tatcttgtggaaaggacgaaacaccGGGTCAGTGACCCGGCTGCGgtttaagagctatgctggaaacagc |
| Faf2 | Faf2-4 | ACGTTCCCTGGTAGAAGAC | tatcttgtggaaaggacgaaacaccGACGTTCCCTGGTAGAAGACgtttaagagctatgctggaaacagc |
| Faf2 | Faf2-5 | GGCACTTAACGATGCCAAG | tatcttgtggaaaggacgaaacaccGGGCACTTAACGATGCCAAGgtttaagagctatgctggaaacagc |
| Fam133b | Fam133b-2 | GTTCTAAGGGAGGAAGTCA | tatcttgtggaaaggacgaaacaccGGTTCTAAGGGAGGAAGTCAgtttaagagctatgctggaaacagc |
| Fam133b | Fam133b-4 | TCTCCTGAGAGCTCGGGGT | tatcttgtggaaaggacgaaacaccGTCTCCTGAGAGCTCGGGGTgtttaagagctatgctggaaacagc |
| Fam133b | Fam133b-5 | GGCTTCAGACAGCAAGGTG | tatcttgtggaaaggacgaaacaccGGGCTTCAGACAGCAAGGTGgtttaagagctatgctggaaacagc |
| Fam166c | Fam166c-2 | CCATAAGAAGAGCCAAAGG | tatcttgtggaaaggacgaaacaccGCCATAAGAAGAGCCAAAGGgtttaagagctatgctggaaacagc |
| Fam166c | Fam166c-3 | GCTATGCGCTCCTCAGTGG | tatcttgtggaaaggacgaaacaccGGCTATGCGCTCCTCAGTGGgtttaagagctatgctggaaacagc |
| Fam166c | Fam166c-4 | GACTGATAACTCACAGAAC | tatcttgtggaaaggacgaaacaccGGACTGATAACTCACAGAACgtttaagagctatgctggaaacagc |
| Fam183b | Fam183b-2 | AGACCGAAGCCCAAGAAAT | tatcttgtggaaaggacgaaacaccGAGACCGAAGCCCAAGAAATgtttaagagctatgctggaaacagc |
| Fam183b | Fam183b-4 | TCTAGTTCACACAATTACC | tatcttgtggaaaggacgaaacaccGTCTAGTTCACACAATTACCgtttaagagctatgctggaaacagc |
| Fam183b | Fam183b-6 | ACTCTTTCTCAAAGAGCTA | tatcttgtggaaaggacgaaacaccGACTCTTTCTCAAAGAGCTAgtttaagagctatgctggaaacagc |
| Fam220a | Fam220a-1 | CATGGAAAGTCACGGGCCA | tatcttgtggaaaggacgaaacaccGCATGGAAAGTCACGGGCCAgtttaagagctatgctggaaacagc |
| Fam220a | Fam220a-2 | TTTGAAGCCACCATGATGC | tatcttgtggaaaggacgaaacaccGTTTGAAGCCACCATGATGCgtttaagagctatgctggaaacagc |
| Fam220a | Fam220a-3 | CAGAGCCACAGCCTTGCCG | tatcttgtggaaaggacgaaacaccGCAGAGCCACAGCCTTGCCGgtttaagagctatgctggaaacagc |
| Fam50a | Fam50a-1 | AGGCTGGAGATCTTCCGCT | tatcttgtggaaaggacgaaacaccGAGGCTGGAGATCTTCCGCTgtttaagagctatgctggaaacagc |
| Fam50a | Fam50a-3 | GAAGCTTGTGTCCACATCA | tatcttgtggaaaggacgaaacaccGGAAGCTTGTGTCCACATCAgtttaagagctatgctggaaacagc |
| Fam50a | Fam50a-5 | GCCTACCTTAACTGTGCGC | tatcttgtggaaaggacgaaacaccGGCCTACCTTAACTGTGCGCgtttaagagctatgctggaaacagc |
| Fanca | Fanca-1 | CGGTAGGCATCGTAGATCG | tatcttgtggaaaggacgaaacaccGCGGTAGGCATCGTAGATCGgtttaagagctatgctggaaacagc |
| Fanca | Fanca-2 | ACTCTCGGAGCAAGCAGCG | tatcttgtggaaaggacgaaacaccGACTCTCGGAGCAAGCAGCGgtttaagagctatgctggaaacagc |
| Fanca | Fanca-5 | CACAGCGTCGTGAGCCTAC | tatcttgtggaaaggacgaaacaccGCACAGCGTCGTGAGCCTACgtttaagagctatgctggaaacagc |
| Fancb | Fancb-1 | TCACCGATCCACTCAATAG | tatcttgtggaaaggacgaaacaccGTCACCGATCCACTCAATAGgtttaagagctatgctggaaacagc |
| Fancb | Fancb-3 | CACGTGCATGTCTGGGCGG | tatcttgtggaaaggacgaaacaccGCACGTGCATGTCTGGGCGGgtttaagagctatgctggaaacagc |
| Fancb | Fancb-5 | GAACAACTTAGCTTAGTAC | tatcttgtggaaaggacgaaacaccGGAACAACTTAGCTTAGTACgtttaagagctatgctggaaacagc |
| Fancc | Fancc-1 | GGATGTACTCACGACAAGA | tatcttgtggaaaggacgaaacaccGGGATGTACTCACGACAAGAgtttaagagctatgctggaaacagc |
| Fancc | Fancc-2 | GTGAGTAGAGCCTCTACCA | tatcttgtggaaaggacgaaacaccGGTGAGTAGAGCCTCTACCAgtttaagagctatgctggaaacagc |
| Fancc | Fancc-4 | TGGACTGAGCACTCAAAGT | tatcttgtggaaaggacgaaacaccGTGGACTGAGCACTCAAAGTgtttaagagctatgctggaaacagc |
| Fancd2 | Fancd2-1 | GGGAAGCCTGAATTCGTGA | tatcttgtggaaaggacgaaacaccGGGGAAGCCTGAATTCGTGAgtttaagagctatgctggaaacagc |
| Fancd2 | Fancd2-2 | ATTGAACGCATTGAGTCCG | tatcttgtggaaaggacgaaacaccGATTGAACGCATTGAGTCCGgtttaagagctatgctggaaacagc |
| Fancd2 | Fancd2-4 | GTGAAAGCGCTCTATGGAC | tatcttgtggaaaggacgaaacaccGGTGAAAGCGCTCTATGGACgtttaagagctatgctggaaacagc |
| Fancg | Fancg-1 | CCGCCGTCCGTGTTAGTCC | tatcttgtggaaaggacgaaacaccGCCGCCGTCCGTGTTAGTCCgtttaagagctatgctggaaacagc |
| Fancg | Fancg-3 | GCTACTACTGCTAAAATCC | tatcttgtggaaaggacgaaacaccGGCTACTACTGCTAAAATCCgtttaagagctatgctggaaacagc |
| Fancg | Fancg-4 | TGCTGTTACAAACCCTAAA | tatcttgtggaaaggacgaaacaccGTGCTGTTACAAACCCTAAAgtttaagagctatgctggaaacagc |
| Fanci | Fanci-1 | ATCTCCCGATTCAACCAAC | tatcttgtggaaaggacgaaacaccGATCTCCCGATTCAACCAACgtttaagagctatgctggaaacagc |
| Fanci | Fanci-2 | TATTCCCGAGCTAGCCAGC | tatcttgtggaaaggacgaaacaccGTATTCCCGAGCTAGCCAGCgtttaagagctatgctggaaacagc |
| Fanci | Fanci-4 | TACGTCGTAAAGCATGAGC | tatcttgtggaaaggacgaaacaccGTACGTCGTAAAGCATGAGCgtttaagagctatgctggaaacagc |
| Fancm | Fancm-3 | GTTGCAGAACAGCGCGCTC | tatcttgtggaaaggacgaaacaccGGTTGCAGAACAGCGCGCTCgtttaagagctatgctggaaacagc |
| Fancm | Fancm-4 | AAGACCTTCATTGCCGCCG | tatcttgtggaaaggacgaaacaccGAAGACCTTCATTGCCGCCGgtttaagagctatgctggaaacagc |
| Fancm | Fancm-5 | GAACTGGCCGCAAACGGCA | tatcttgtggaaaggacgaaacaccGGAACTGGCCGCAAACGGCAgtttaagagctatgctggaaacagc |
| Farsa | Farsa-3 | TCACCACTCGGAACACCCG | tatcttgtggaaaggacgaaacaccGTCACCACTCGGAACACCCGgtttaagagctatgctggaaacagc |
| Farsa | Farsa-4 | AGCACAAGCGTGTCTAAGC | tatcttgtggaaaggacgaaacaccGAGCACAAGCGTGTCTAAGCgtttaagagctatgctggaaacagc |
| Farsa | Farsa-5 | GTCCCTCCAAGAGCCACTA | tatcttgtggaaaggacgaaacaccGGTCCCTCCAAGAGCCACTAgtttaagagctatgctggaaacagc |
| Fbxw16 | Fbxw16-2 | TCAAATTTCCATTATGCCA | tatcttgtggaaaggacgaaacaccGTCAAATTTCCATTATGCCAgtttaagagctatgctggaaacagc |
| Fbxw16 | Fbxw16-3 | CAATATAGGATTCCTGTAC | tatcttgtggaaaggacgaaacaccGCAATATAGGATTCCTGTACgtttaagagctatgctggaaacagc |

| | | | |
|---|---|---|---|
| Fbxw16 | Fbxw16-4 | TATGGTGACTTCCACGAAC | tatcttgtggaaaggacgaaacaccGTATGGTGACTTCCACGAACgtttaagagctatgctggaaacagc |
| Fbxw21 | Fbxw21-1 | AAGTTTATCACATTTGATA | tatcttgtggaaaggacgaaacaccGAAGTTTATCACATTTGATAgtttaagagctatgctggaaacagc |
| Fbxw21 | Fbxw21-3 | TGGTAATATGGTATAAGTC | tatcttgtggaaaggacgaaacaccGTGGTAATATGGTATAAGTCgtttaagagctatgctggaaacagc |
| Fbxw21 | Fbxw21-5 | ATTGCAGTCACTGTAGATA | tatcttgtggaaaggacgaaacaccGATTGCAGTCACTGTAGATAgtttaagagctatgctggaaacagc |
| Fez1 | Fez1-1 | ACATCTCCTCATCACCTGG | tatcttgtggaaaggacgaaacaccGACATCTCCTCATCACCTGGgtttaagagctatgctggaaacagc |
| Fez1 | Fez1-2 | CTTCTACAGATTCATGAGA | tatcttgtggaaaggacgaaacaccGCTTCTACAGATTCATGAGAgtttaagagctatgctggaaacagc |
| Fez1 | Fez1-4 | AACTCTCCAGATCCTGAAG | tatcttgtggaaaggacgaaacaccGAACTCTCCAGATCCTGAAGgtttaagagctatgctggaaacagc |
| Foxd3 | Foxd3-1 | CGTGATGAGCGCTCACCGG | tatcttgtggaaaggacgaaacaccGCGTGATGAGCGCTCACCGGgtttaagagctatgctggaaacagc |
| Foxd3 | Foxd3-3 | GCGAGGACGCGGTGACAGG | tatcttgtggaaaggacgaaacaccGGCGAGGACGCGGTGACAGGgtttaagagctatgctggaaacagc |
| Foxd3 | Foxd3-4 | AACCGTTTTCCGTACTACC | tatcttgtggaaaggacgaaacaccGAACCGTTTTCCGTACTACCgtttaagagctatgctggaaacagc |
| Foxh1 | Foxh1-2 | TCTAATCGGTGCTTCCATA | tatcttgtggaaaggacgaaacaccGTCTAATCGGTGCTTCCATAgtttaagagctatgctggaaacagc |
| Foxh1 | Foxh1-3 | TGATTTCAGATTATCCGTC | tatcttgtggaaaggacgaaacaccGTGATTTCAGATTATCCGTCgtttaagagctatgctggaaacagc |
| Foxh1 | Foxh1-4 | TCTTTCGAAGACCCCCAAG | tatcttgtggaaaggacgaaacaccGTCTTTCGAAGACCCCCAAGgtttaagagctatgctggaaacagc |
| Foxn2 | Foxn2-1 | TTCTAACAAACCTCAGACT | tatcttgtggaaaggacgaaacaccGTTCTAACAAACCTCAGACTgtttaagagctatgctggaaacagc |
| Foxn2 | Foxn2-2 | AGTCCGCTGTATGACATAG | tatcttgtggaaaggacgaaacaccGAGTCCGCTGTATGACATAGgtttaagagctatgctggaaacagc |
| Foxn2 | Foxn2-5 | TTTAAGGCCTGCATAAGAT | tatcttgtggaaaggacgaaacaccGTTTAAGGCCTGCATAAGATgtttaagagctatgctggaaacagc |
| Fra10ac1 | Fra10ac1-2 | CATAGATTCCTTTGGAATG | tatcttgtggaaaggacgaaacaccGCATAGATTCCTTTGGAATGgtttaagagctatgctggaaacagc |
| Fra10ac1 | Fra10ac1-4 | CGCCGCAGACTTGCTCGAC | tatcttgtggaaaggacgaaacaccGCGCCGCAGACTTGCTCGACgtttaagagctatgctggaaacagc |
| Fra10ac1 | Fra10ac1-5 | CGTTTCAAAAAGAAAGGCA | tatcttgtggaaaggacgaaacaccGCGTTTCAAAAAGAAAGGCAgtttaagagctatgctggaaacagc |
| Fsbp | Fsbp-2 | CCTACGCAGCCTTTACAAA | tatcttgtggaaaggacgaaacaccGCCTACGCAGCCTTTACAAAgtttaagagctatgctggaaacagc |
| Fsbp | Fsbp-3 | GAGGCCGAAGCTGGCAATC | tatcttgtggaaaggacgaaacaccGGAGGCCGAAGCTGGCAATCgtttaagagctatgctggaaacagc |
| Fsbp | Fsbp-4 | ATGTGCTATAGTGCAAACT | tatcttgtggaaaggacgaaacaccGATGTGCTATAGTGCAAACTgtttaagagctatgctggaaacagc |
| Fshr | Fshr-2 | TTCAAGCGGTATGTTGACC | tatcttgtggaaaggacgaaacaccGTTCAAGCGGTATGTTGACCgtttaagagctatgctggaaacagc |
| Fshr | Fshr-3 | TCTTACAGATATCGGAGAC | tatcttgtggaaaggacgaaacaccGTCTTACAGATATCGGAGACgtttaagagctatgctggaaacagc |
| Fshr | Fshr-5 | CGTTCCGGGGGAGGTCGGG | tatcttgtggaaaggacgaaacaccGCGTTCCGGGGGAGGTCGGGgtttaagagctatgctggaaacagc |
| Fus | Fus-1 | ATGGTTCCACTGGAGGCTA | tatcttgtggaaaggacgaaacaccGATGGTTCCACTGGAGGCTAgtttaagagctatgctggaaacagc |
| Fus | Fus-3 | GTTGTCCATAACCCTGAGG | tatcttgtggaaaggacgaaacaccGGTTGTCCATAACCCTGAGGgtttaagagctatgctggaaacagc |
| Fus | Fus-4 | AAGATCAGTCCTCTATGAG | tatcttgtggaaaggacgaaacaccGAAGATCAGTCCTCTATGAGgtttaagagctatgctggaaacagc |
| G2e3 | G2e3-1 | GATAACTTCGTTTACACGT | tatcttgtggaaaggacgaaacaccGGATAACTTCGTTTACACGTgtttaagagctatgctggaaacagc |
| G2e3 | G2e3-2 | CGCAAAATTGTCAGTGAAC | tatcttgtggaaaggacgaaacaccGCGCAAAATTGTCAGTGAACgtttaagagctatgctggaaacagc |
| G2e3 | G2e3-4 | AGACTACAACGAACCTAAC | tatcttgtggaaaggacgaaacaccGAGACTACAACGAACCTAACgtttaagagctatgctggaaacagc |
| Gab3 | Gab3-1 | TTAGTTCTTTTACCCCACT | tatcttgtggaaaggacgaaacaccGTTAGTTCTTTTACCCCACTgtttaagagctatgctggaaacagc |
| Gab3 | Gab3-2 | TTTGACCAGCTATCACATC | tatcttgtggaaaggacgaaacaccGTTTGACCAGCTATCACATCgtttaagagctatgctggaaacagc |
| Gab3 | Gab3-4 | GGTAGTATTCTAAGACATC | tatcttgtggaaaggacgaaacaccGGGTAGTATTCTAAGACATCgtttaagagctatgctggaaacagc |
| Gabpb1 | Gabpb1-1 | GTGTCAGGACTCTCCGGGT | tatcttgtggaaaggacgaaacaccGGTGTCAGGACTCTCCGGGTgtttaagagctatgctggaaacagc |
| Gabpb1 | Gabpb1-2 | ACAGAACATAATCATCAAG | tatcttgtggaaaggacgaaacaccGACAGAACATAATCATCAAGgtttaagagctatgctggaaacagc |
| Gabpb1 | Gabpb1-5 | AGCTCCTTTTACTACAGAC | tatcttgtggaaaggacgaaacaccGAGCTCCTTTTACTACAGACgtttaagagctatgctggaaacagc |
| Gak | Gak-1 | TTGAACTGTTACGCCCAGG | tatcttgtggaaaggacgaaacaccGTTGAACTGTTACGCCCAGGgtttaagagctatgctggaaacagc |
| Gak | Gak-3 | CCGGTATATCCTTGGAGAA | tatcttgtggaaaggacgaaacaccGCCGGTATATCCTTGGAGAAgtttaagagctatgctggaaacagc |
| Gak | Gak-4 | GAACTTACGAATAAGGTCA | tatcttgtggaaaggacgaaacaccGGAACTTACGAATAAGGTCAgtttaagagctatgctggaaacagc |
| Gar1 | Gar1-2 | AAACAGGGGCGTTGAAGTA | tatcttgtggaaaggacgaaacaccGAAACAGGGGCGTTGAAGTAgtttaagagctatgctggaaacagc |
| Gar1 | Gar1-3 | TACAGACGACACGTTCTGG | tatcttgtggaaaggacgaaacaccGTACAGACGACACGTTCTGGgtttaagagctatgctggaaacagc |
| Gar1 | Gar1-4 | TTAATAAATTTCAAGATCA | tatcttgtggaaaggacgaaacaccGTTAATAAATTTCAAGATCAgtttaagagctatgctggaaacagc |
| Gje1 | Gje1-1 | CCAGAATGCGAACACCTCC | tatcttgtggaaaggacgaaacaccGCCAGAATGCGAACACCTCCgtttaagagctatgctggaaacagc |
| Gje1 | Gje1-2 | GTCTTGTTAAGAATCAGCC | tatcttgtggaaaggacgaaacaccGGTCTTGTTAAGAATCAGCCgtttaagagctatgctggaaacagc |
| Gje1 | Gje1-4 | GAACTGATTGTAACAGAAC | tatcttgtggaaaggacgaaacaccGGAACTGATTGTAACAGAACgtttaagagctatgctggaaacagc |
| Glmn | Glmn-1 | AGATGCAAAACACCGAAAA | tatcttgtggaaaggacgaaacaccGAGATGCAAAACACCGAAAAgtttaagagctatgctggaaacagc |
| Glmn | Glmn-2 | TCGAACAGTCAGAAGACGT | tatcttgtggaaaggacgaaacaccGTCGAACAGTCAGAAGACGTgtttaagagctatgctggaaacagc |
| Glmn | Glmn-4 | CACCCCATAGACTTAATGA | tatcttgtggaaaggacgaaacaccGCACCCCATAGACTTAATGAgtttaagagctatgctggaaacagc |
| Glrx | Glrx-1 | TTTGAAAGGCAGTTGACTG | tatcttgtggaaaggacgaaacaccGTTTGAAAGGCAGTTGACTGgtttaagagctatgctggaaacagc |
| Glrx | Glrx-3 | ATCGCACTGGTGTTGTTAG | tatcttgtggaaaggacgaaacaccGATCGCACTGGTGTTGTTAGgtttaagagctatgctggaaacagc |
| Glrx | Glrx-5 | ATTATTTACAACAGCTCAC | tatcttgtggaaaggacgaaacaccGATTATTTACAACAGCTCACgtttaagagctatgctggaaacagc |
| Gm4847 | Gm4847-1 | ATATTTAAACATGGAGTAC | tatcttgtggaaaggacgaaacaccGATATTTAAACATGGAGTACgtttaagagctatgctggaaacagc |
| Gm4847 | Gm4847-2 | TGAGTCATCACTCAAGAAA | tatcttgtggaaaggacgaaacaccGTGAGTCATCACTCAAGAAAgtttaagagctatgctggaaacagc |
| Gm4847 | Gm4847-4 | TCGGGGTGTTTGTACTCCC | tatcttgtggaaaggacgaaacaccGTCGGGGTGTTTGTACTCCCgtttaagagctatgctggaaacagc |
| Gm5150 | Gm5150-1 | GCCGCTGGAAAAGAGTTGA | tatcttgtggaaaggacgaaacaccGGCCGCTGGAAAAGAGTTGAgtttaagagctatgctggaaacagc |
| Gm5150 | Gm5150-2 | ATGTCACTTTTGCTGATGC | tatcttgtggaaaggacgaaacaccGATGTCACTTTTGCTGATGCgtttaagagctatgctggaaacagc |
| Gm5150 | Gm5150-3 | ACTTGATATATTCTTACAC | tatcttgtggaaaggacgaaacaccGACTTGATATATTCTTACACgtttaagagctatgctggaaacagc |
| Gm527 | Gm527-1 | GTCTGATCTCAACATAGAG | tatcttgtggaaaggacgaaacaccGGTCTGATCTCAACATAGAGgtttaagagctatgctggaaacagc |
| Gm527 | Gm527-2 | AAAATACCTGCACCATAGA | tatcttgtggaaaggacgaaacaccGAAAATACCTGCACCATAGAgtttaagagctatgctggaaacagc |
| Gm527 | Gm527-5 | AGGGAGTTACTGCTCTAAC | tatcttgtggaaaggacgaaacaccGAGGGAGTTACTGCTCTAACgtttaagagctatgctggaaacagc |
| Gm5916 | Gm5916-1 | TAAGCCTGGATGCAGAACC | tatcttgtggaaaggacgaaacaccGTAAGCCTGGATGCAGAACCgtttaagagctatgctggaaacagc |
| Gm5916 | Gm5916-2 | AGAGTGGACATTGTTCGGC | tatcttgtggaaaggacgaaacaccGAGAGTGGACATTGTTCGGCgtttaagagctatgctggaaacagc |
| Gm5916 | Gm5916-3 | TCTGCAACTCTTTTAAGAG | tatcttgtggaaaggacgaaacaccGTCTGCAACTCTTTTAAGAGgtttaagagctatgctggaaacagc |
| Gm8882 | Gm8882-1 | AGACGCACAGGGTCCCCCT | tatcttgtggaaaggacgaaacaccGAGACGCACAGGGTCCCCCTgtttaagagctatgctggaaacagc |
| Gm8882 | Gm8882-2 | CACCATGGACCTCCCCCAC | tatcttgtggaaaggacgaaacaccGCACCATGGACCTCCCCCACgtttaagagctatgctggaaacagc |
| Gm8882 | Gm8882-3 | TCTTTACTTGAATAGAGAT | tatcttgtggaaaggacgaaacaccGTCTTTACTTGAATAGAGATgtttaagagctatgctggaaacagc |
| Gna13 | Gna13-2 | ATCACGTTGCTGTAGATGG | tatcttgtggaaaggacgaaacaccGATCACGTTGCTGTAGATGGgtttaagagctatgctggaaacagc |
| Gna13 | Gna13-3 | TTTGATACCCGCGCCCCCA | tatcttgtggaaaggacgaaacaccGTTTGATACCCGCGCCCCCAgtttaagagctatgctggaaacagc |
| Gna13 | Gna13-5 | TTGGATAAACTTGGAGTAC | tatcttgtggaaaggacgaaacaccGTTGGATAAACTTGGAGTACgtttaagagctatgctggaaacagc |
| Gnb1 | Gnb1-1 | CAAACAATATTGATCCAGT | tatcttgtggaaaggacgaaacaccGCAAACAATATTGATCCAGTgtttaagagctatgctggaaacagc |
| Gnb1 | Gnb1-3 | CCAGAAGGAGCGTATGCGC | tatcttgtggaaaggacgaaacaccGCCAGAAGGAGCGTATGCGCgtttaagagctatgctggaaacagc |
| Gnb1 | Gnb1-5 | TTGTCAGGTACTCACCATG | tatcttgtggaaaggacgaaacaccGTTGTCAGGTACTCACCATGgtttaagagctatgctggaaacagc |
| Gngt2 | Gngt2-2 | GAAATCAAGGATTATGTAG | tatcttgtggaaaggacgaaacaccGGAAATCAAGGATTATGTAGgtttaagagctatgctggaaacagc |
| Gngt2 | Gngt2-3 | AAGAGGGTCTGTCCCTGCT | tatcttgtggaaaggacgaaacaccGAAGAGGGTCTGTCCCTGCTgtttaagagctatgctggaaacagc |
| Gngt2 | Gngt2-4 | TGGGATGCCTTTGAGAAGA | tatcttgtggaaaggacgaaacaccGTGGGATGCCTTTGAGAAGAgtttaagagctatgctggaaacagc |

| | | | |
|---|---|---|---|
| Gnl2 | Gnl2-1 | AATACTCGTGTGATCAAGC | tatcttgtggaaaggacgaaacaccGAATACTCGTGTGATCAAGCgtttaagagctatgctggaaacagc |
| Gnl2 | Gnl2-3 | AACTTACGGTTGCCCAAGT | tatcttgtggaaaggacgaaacaccGAACTTACGGTTGCCCAAGTgtttaagagctatgctggaaacagc |
| Gnl2 | Gnl2-4 | TCCAAGAAAGTTTGCAACG | tatcttgtggaaaggacgaaacaccGTCCAAGAAAGTTTGCAACGgtttaagagctatgctggaaacagc |
| Golga4 | Golga4-1 | CTGTTTCGAAGTCCCATAA | tatcttgtggaaaggacgaaacaccGCTGTTTCGAAGTCCCATAAgtttaagagctatgctggaaacagc |
| Golga4 | Golga4-4 | GCGCCGCACGCGAGCAGGC | tatcttgtggaaaggacgaaacaccGGCGCCGCACGCGAGCAGGCgtttaagagctatgctggaaacagc |
| Golga4 | Golga4-5 | GACACGGTGTCGAGACTGG | tatcttgtggaaaggacgaaacaccGGACACGGTGTCGAGACTGGgtttaagagctatgctggaaacagc |
| Gpatch11 | Gpatch11-1 | GCGGGAAAGACGTGACATC | tatcttgtggaaaggacgaaacaccGGCGGGAAAGACGTGACATCgtttaagagctatgctggaaacagc |
| Gpatch11 | Gpatch11-4 | TTTTGACATTGAGAGGAAT | tatcttgtggaaaggacgaaacaccGTTTTGACATTGAGAGGAATgtttaagagctatgctggaaacagc |
| Gpatch11 | Gpatch11-5 | TCATCATTAAAACGGAAAG | tatcttgtggaaaggacgaaacaccGTCATCATTAAAACGGAAAGgtttaagagctatgctggaaacagc |
| Gpn2 | Gpn2-1 | AGTGAGTTCCTGCGCGCTC | tatcttgtggaaaggacgaaacaccGAGTGAGTTCCTGCGCGCTCgtttaagagctatgctggaaacagc |
| Gpn2 | Gpn2-2 | AGGCAGTCCGTCGTTAGCG | tatcttgtggaaaggacgaaacaccGAGGCAGTCCGTCGTTAGCGgtttaagagctatgctggaaacagc |
| Gpn2 | Gpn2-3 | TACGAGTGTGCGGTGGACG | tatcttgtggaaaggacgaaacaccGTACGAGTGTGCGGTGGACGgtttaagagctatgctggaaacagc |
| Gps1 | Gps1-1 | TTGAAGGTCCTCTGCACGA | tatcttgtggaaaggacgaaacaccGTTGAAGGTCCTCTGCACGAgtttaagagctatgctggaaacagc |
| Gps1 | Gps1-3 | GGCGCGGCCACGATGACCT | tatcttgtggaaaggacgaaacaccGGGCGCGGCCACGATGACCTgtttaagagctatgctggaaacagc |
| Gps1 | Gps1-4 | GTGATCACTACCTGGACTG | tatcttgtggaaaggacgaaacaccGGTGATCACTACCTGGACTGgtttaagagctatgctggaaacagc |
| Gria4 | Gria4-3 | TTTTGGACTCTATGACAAG | tatcttgtggaaaggacgaaacaccGTTTTGGACTCTATGACAAGgtttaagagctatgctggaaacagc |
| Gria4 | Gria4-4 | AAACACCCCTCTAGAATAC | tatcttgtggaaaggacgaaacaccGAAACACCCCTCTAGAATACgtttaagagctatgctggaaacagc |
| Gria4 | Gria4-5 | ATGAGGTACCAAATTGAAA | tatcttgtggaaaggacgaaacaccGATGAGGTACCAAATTGAAAgtttaagagctatgctggaaacagc |
| Grid1 | Grid1-1 | TTGGCGGATGCACAGCCCG | tatcttgtggaaaggacgaaacaccGTTGGCGGATGCACAGCCCGgtttaagagctatgctggaaacagc |
| Grid1 | Grid1-3 | GAGGAGCTGAATCGCTACC | tatcttgtggaaaggacgaaacaccGGAGGAGCTGAATCGCTACCgtttaagagctatgctggaaacagc |
| Grid1 | Grid1-5 | GTAACGCTTGGGTTGTCCA | tatcttgtggaaaggacgaaacaccGGTAACGCTTGGGTTGTCCAgtttaagagctatgctggaaacagc |
| Gtf2a1 | Gtf2a1-3 | TGAAATCCCTCCAGGAAGT | tatcttgtggaaaggacgaaacaccGTGAAATCCCTCCAGGAAGTgtttaagagctatgctggaaacagc |
| Gtf2a1 | Gtf2a1-4 | GGTTATACTGCTTACCTGC | tatcttgtggaaaggacgaaacaccGGGTTATACTGCTTACCTGCgtttaagagctatgctggaaacagc |
| Gtf2a1 | Gtf2a1-5 | GAACCGGAGTCACTCCTGC | tatcttgtggaaaggacgaaacaccGGAACCGGAGTCACTCCTGCgtttaagagctatgctggaaacagc |
| Gtf2f1 | Gtf2f1-3 | TTCAACCGCAAGCTTAGGG | tatcttgtggaaaggacgaaacaccGTTCAACCGCAAGCTTAGGGgtttaagagctatgctggaaacagc |
| Gtf2f1 | Gtf2f1-4 | AGATGCCAGAGTCCGGTGC | tatcttgtggaaaggacgaaacaccGAGATGCCAGAGTCCGGTGCgtttaagagctatgctggaaacagc |
| Gtf2f1 | Gtf2f1-5 | TAAAGTCAACTTTGCTACC | tatcttgtggaaaggacgaaacaccGTAAAGTCAACTTTGCTACCgtttaagagctatgctggaaacagc |
| Gtf3c3 | Gtf3c3-1 | AGCGTGCCAATAGACATCA | tatcttgtggaaaggacgaaacaccGAGCGTGCCAATAGACATCAgtttaagagctatgctggaaacagc |
| Gtf3c3 | Gtf3c3-2 | GGAGAACGTTTTATGCAGT | tatcttgtggaaaggacgaaacaccGGGAGAACGTTTTATGCAGTgtttaagagctatgctggaaacagc |
| Gtf3c3 | Gtf3c3-3 | TACTAATGTCCGTTACCTG | tatcttgtggaaaggacgaaacaccGTACTAATGTCCGTTACCTGgtttaagagctatgctggaaacagc |
| H2ax | H2ax-1 | CCGCGTACACCGGCTGCTG | tatcttgtggaaaggacgaaacaccGCCGCGTACACCGGCTGCTGgtttaagagctatgctggaaacagc |
| H2ax | H2ax-3 | GCAGGCGCGCCGGTGTACC | tatcttgtggaaaggacgaaacaccGGCAGGCGCGCCGGTGTACCgtttaagagctatgctggaaacagc |
| H2ax | H2ax-4 | GTACTCGAGCACCGCTGCC | tatcttgtggaaaggacgaaacaccGGTACTCGAGCACCGCTGCCgtttaagagctatgctggaaacagc |
| Hbb-y | Hbb-y-1 | GACAACCTCAAGTCTGCCT | tatcttgtggaaaggacgaaacaccGGACAACCTCAAGTCTGCCTgtttaagagctatgctggaaacagc |
| Hbb-y | Hbb-y-2 | AGGTTGTCTAGATTCTTAA | tatcttgtggaaaggacgaaacaccGAGGTTGTCTAGATTCTTAAgtttaagagctatgctggaaacagc |
| Hbb-y | Hbb-y-3 | ACCCAAGGGTCAAAGCCCA | tatcttgtggaaaggacgaaacaccGACCCAAGGGTCAAAGCCCAgtttaagagctatgctggaaacagc |
| Hdac11 | Hdac11-1 | AGGTGATCAACTTCCTGAA | tatcttgtggaaaggacgaaacaccGAGGTGATCAACTTCCTGAAgtttaagagctatgctggaaacagc |
| Hdac11 | Hdac11-3 | ACCGGGGGGATCTCAGTGA | tatcttgtggaaaggacgaaacaccGACCGGGGGGATCTCAGTGAgtttaagagctatgctggaaacagc |
| Hdac11 | Hdac11-5 | CAGTTCCTGTTTGAACGCG | tatcttgtggaaaggacgaaacaccGCAGTTCCTGTTTGAACGCGgtttaagagctatgctggaaacagc |
| Hdlbp | Hdlbp-1 | GGAGGCGCAAACATTAAAA | tatcttgtggaaaggacgaaacaccGGGAGGCGCAAACATTAAAAgtttaagagctatgctggaaacagc |
| Hdlbp | Hdlbp-2 | GTCTCCCTCAATTCGGATC | tatcttgtggaaaggacgaaacaccGGTCTCCCTCAATTCGGATCgtttaagagctatgctggaaacagc |
| Hdlbp | Hdlbp-5 | GGCCGGATCTTGTTACTCC | tatcttgtggaaaggacgaaacaccGGGCCGGATCTTGTTACTCCgtttaagagctatgctggaaacagc |
| Helq | Helq-1 | CCCGACCGCTTGGTAGCAC | tatcttgtggaaaggacgaaacaccGCCCGACCGCTTGGTAGCACgtttaagagctatgctggaaacagc |
| Helq | Helq-3 | CTATGCGCTTTCTCGATAG | tatcttgtggaaaggacgaaacaccGCTATGCGCTTTCTCGATAGgtttaagagctatgctggaaacagc |
| Helq | Helq-4 | TCATCAACGTGAGCTAAGA | tatcttgtggaaaggacgaaacaccGTCATCAACGTGAGCTAAGAgtttaagagctatgctggaaacagc |
| Hgfac | Hgfac-2 | ATGACCGAGACCGGGCCTG | tatcttgtggaaaggacgaaacaccGATGACCGAGACCGGGCCTGgtttaagagctatgctggaaacagc |
| Hgfac | Hgfac-3 | CGCTATGAGTATTTTGAGG | tatcttgtggaaaggacgaaacaccGCGCTATGAGTATTTTGAGGgtttaagagctatgctggaaacagc |
| Hgfac | Hgfac-4 | CCAGCATAGCCCAACGGGC | tatcttgtggaaaggacgaaacaccGCCAGCATAGCCCAACGGGCgtttaagagctatgctggaaacagc |
| Hltf | Hltf-1 | TTGGACTACGCTATTACAC | tatcttgtggaaaggacgaaacaccGTTGGACTACGCTATTACACgtttaagagctatgctggaaacagc |
| Hltf | Hltf-2 | GGACCAAGCTATAGTAGGC | tatcttgtggaaaggacgaaacaccGGGACCAAGCTATAGTAGGCgtttaagagctatgctggaaacagc |
| Hltf | Hltf-4 | AAATAGAGTCATCCACGTG | tatcttgtggaaaggacgaaacaccGAAATAGAGTCATCCACGTGgtttaagagctatgctggaaacagc |
| Hmgcs1 | Hmgcs1-2 | GCATTGGGCGGCTAGAAGT | tatcttgtggaaaggacgaaacaccGGCATTGGGCGGCTAGAAGTgtttaagagctatgctggaaacagc |
| Hmgcs1 | Hmgcs1-3 | GCGTTTGGCCCAATTAGCA | tatcttgtggaaaggacgaaacaccGGCGTTTGGCCCAATTAGCAgtttaagagctatgctggaaacagc |
| Hmgcs1 | Hmgcs1-4 | AGAGCTTTCCGTCGACCAC | tatcttgtggaaaggacgaaacaccGAGAGCTTTCCGTCGACCACgtttaagagctatgctggaaacagc |
| Hmgn5 | Hmgn5-2 | AAGGCCTAAACGAGAAACC | tatcttgtggaaaggacgaaacaccGAAGGCCTAAACGAGAAACCgtttaagagctatgctggaaacagc |
| Hmgn5 | Hmgn5-3 | CAGCTTCCATTTTAGGAAT | tatcttgtggaaaggacgaaacaccGCAGCTTCCATTTTAGGAATgtttaagagctatgctggaaacagc |
| Hmgn5 | Hmgn5-4 | TCTTTAACATCTTCTGGCT | tatcttgtggaaaggacgaaacaccGTCTTTAACATCTTCTGGCTgtttaagagctatgctggaaacagc |
| Hormad1 | Hormad1-1 | GGAAATATTCCTCTCAAAT | tatcttgtggaaaggacgaaacaccGGGAAATATTCCTCTCAAATgtttaagagctatgctggaaacagc |
| Hormad1 | Hormad1-4 | ACCAAAAATGGACCAATCA | tatcttgtggaaaggacgaaacaccGACCAAAAATGGACCAATCAgtttaagagctatgctggaaacagc |
| Hormad1 | Hormad1-5 | GACAAACATCATTAGGTAA | tatcttgtggaaaggacgaaacaccGGACAAACATCATTAGGTAAgtttaagagctatgctggaaacagc |
| Hrob | Hrob-2 | TCGTTACGGAAGGAGCCTA | tatcttgtggaaaggacgaaacaccGTCGTTACGGAAGGAGCCTAgtttaagagctatgctggaaacagc |
| Hrob | Hrob-4 | GCACCTCAGTATCTGCCAG | tatcttgtggaaaggacgaaacaccGGCACCTCAGTATCTGCCAGgtttaagagctatgctggaaacagc |
| Hrob | Hrob-5 | CGTTGAGGTCCAGAAAAAC | tatcttgtggaaaggacgaaacaccGCGTTGAGGTCCAGAAAAACgtttaagagctatgctggaaacagc |
| Hsd3b2 | Hsd3b2-1 | ATTATGGCCCTTAAACATA | tatcttgtggaaaggacgaaacaccGATTATGGCCCTTAAACATAgtttaagagctatgctggaaacagc |
| Hsd3b2 | Hsd3b2-2 | GCAGGGCCCAACTCTTACA | tatcttgtggaaaggacgaaacaccGGCAGGGCCCAACTCTTACAgtttaagagctatgctggaaacagc |
| Hsd3b2 | Hsd3b2-3 | GGCACACTGGCTTGGATAC | tatcttgtggaaaggacgaaacaccGGGCACACTGGCTTGGATACgtttaagagctatgctggaaacagc |
| Hus1 | Hus1-1 | CTTACGTCACAGTCTGGGA | tatcttgtggaaaggacgaaacaccGCTTACGTCACAGTCTGGGAgtttaagagctatgctggaaacagc |
| Hus1 | Hus1-2 | TCTACAGACACGGTAAGAC | tatcttgtggaaaggacgaaacaccGTCTACAGACACGGTAAGACgtttaagagctatgctggaaacagc |
| Hus1 | Hus1-5 | TTCTCCGGGCTGATGCGGA | tatcttgtggaaaggacgaaacaccGTTCTCCGGGCTGATGCGGAgtttaagagctatgctggaaacagc |
| Hykk | Hykk-1 | GTCCTCAAAATAAGCAATA | tatcttgtggaaaggacgaaacaccGGTCCTCAAAATAAGCAATAgtttaagagctatgctggaaacagc |
| Hykk | Hykk-4 | CTCTAGGGCTTTATCCAAT | tatcttgtggaaaggacgaaacaccGCTCTAGGGCTTTATCCAATgtttaagagctatgctggaaacagc |
| Hykk | Hykk-5 | CCATGTATTTCTCTAGAAG | tatcttgtggaaaggacgaaacaccGCCATGTATTTCTCTAGAAGgtttaagagctatgctggaaacagc |
| Ifit1bl1 | Ifit1bl1-1 | GAGCAAGGTTGTTTCTGCA | tatcttgtggaaaggacgaaacaccGGAGCAAGGTTGTTTCTGCAgtttaagagctatgctggaaacagc |
| Ifit1bl1 | Ifit1bl1-2 | TTTTACCGAAGGAAAGGAT | tatcttgtggaaaggacgaaacaccGTTTTACCGAAGGAAAGGATgtttaagagctatgctggaaacagc |
| Ifit1bl1 | Ifit1bl1-5 | TCAATGTGATCCAGGCGAT | tatcttgtggaaaggacgaaacaccGTCAATGTGATCCAGGCGATgtttaagagctatgctggaaacagc |
| Ifit1bl2 | Ifit1bl2-1 | ATGAAAAAAGCTACAAGCA | tatcttgtggaaaggacgaaacaccGATGAAAAAAGCTACAAGCAgtttaagagctatgctggaaacagc |

| | | | |
|---|---|---|---|
| Ifit1bl2 | Ifit1bl2-3 | TCTCTACAACCCTTAAAGA | tatcttgtggaaaggacgaaacaccGTCTCTACAACCCTTAAAGAgtttaagagctatgctggaaacagc |
| Ifit1bl2 | Ifit1bl2-5 | TGTGAGCCACACTTCCGCA | tatcttgtggaaaggacgaaacaccGTGTGAGCCACACTTCCGCAgtttaagagctatgctggaaacagc |
| Ifitm6 | Ifitm6-1 | ATCTTGGCGGTTGAAGCAT | tatcttgtggaaaggacgaaacaccGATCTTGGCGGTTGAAGCATgtttaagagctatgctggaaacagc |
| Ifitm6 | Ifitm6-3 | GATGTAGGCAATGAAACCC | tatcttgtggaaaggacgaaacaccGGATGTAGGCAATGAAACCCgtttaagagctatgctggaaacagc |
| Ifitm6 | Ifitm6-4 | TTAACACAGTGTTCATGAA | tatcttgtggaaaggacgaaacaccGTTAACACAGTGTTCATGAAgtttaagagctatgctggaaacagc |
| Il15 | Il15-2 | ATGTGTACTCACTTGAATA | tatcttgtggaaaggacgaaacaccGATGTGTACTCACTTGAATAgtttaagagctatgctggaaacagc |
| Il15 | Il15-4 | TATCTTACATCTATCCAGT | tatcttgtggaaaggacgaaacaccGTATCTTACATCTATCCAGTgtttaagagctatgctggaaacagc |
| Il15 | Il15-5 | GTAGGTCTCCCTAAAACAG | tatcttgtggaaaggacgaaacaccGGTAGGTCTCCCTAAAACAGgtttaagagctatgctggaaacagc |
| Il6st | Il6st-3 | TCTGTACGGCTCATATGGA | tatcttgtggaaaggacgaaacaccGTCTGTACGGCTCATATGGAgtttaagagctatgctggaaacagc |
| Il6st | Il6st-4 | TGCTAGAGACGCGACATAG | tatcttgtggaaaggacgaaacaccGTGCTAGAGACGCGACATAGgtttaagagctatgctggaaacagc |
| Il6st | Il6st-5 | GCAGTACGCACCTGTGACG | tatcttgtggaaaggacgaaacaccGGCAGTACGCACCTGTGACGgtttaagagctatgctggaaacagc |
| Imp3 | Imp3-1 | TACAACCAGCTGAGCCGAG | tatcttgtggaaaggacgaaacaccGTACAACCAGCTGAGCCGAGgtttaagagctatgctggaaacagc |
| Imp3 | Imp3-2 | GAATGGGTCGCGTTCCGGC | tatcttgtggaaaggacgaaacaccGGAATGGGTCGCGTTCCGGCgtttaagagctatgctggaaacagc |
| Imp3 | Imp3-3 | CCATTCCGCGTGCGCGCCT | tatcttgtggaaaggacgaaacaccGCCATTCCGCGTGCGCGCCTgtttaagagctatgctggaaacagc |
| Ints3 | Ints3-2 | AGCCAATGATGGCCCACCG | tatcttgtggaaaggacgaaacaccGAGCCAATGATGGCCCACCGgtttaagagctatgctggaaacagc |
| Ints3 | Ints3-3 | CTTACCCGTTCCCGAAGCA | tatcttgtggaaaggacgaaacaccGCTTACCCGTTCCCGAAGCAgtttaagagctatgctggaaacagc |
| Ints3 | Ints3-4 | GTGGTCCACAATAAGTCGG | tatcttgtggaaaggacgaaacaccGGTGGTCCACAATAAGTCGGgtttaagagctatgctggaaacagc |
| Ints7 | Ints7-1 | ATCCACAGTAACGACCCCG | tatcttgtggaaaggacgaaacaccGATCCACAGTAACGACCCCGgtttaagagctatgctggaaacagc |
| Ints7 | Ints7-2 | TACCGGAGAGTGATGGCTC | tatcttgtggaaaggacgaaacaccGTACCGGAGAGTGATGGCTCgtttaagagctatgctggaaacagc |
| Ints7 | Ints7-4 | TTCAGTGTAGTTCCCGGTA | tatcttgtggaaaggacgaaacaccGTTCAGTGTAGTTCCCGGTAgtttaagagctatgctggaaacagc |
| Jak1 | Jak1-2 | CCGTCTTGGGGTCCCGAAT | tatcttgtggaaaggacgaaacaccGCCGTCTTGGGGTCCCGAATgtttaagagctatgctggaaacagc |
| Jak1 | Jak1-3 | CCGGCTCCACTACCGCATG | tatcttgtggaaaggacgaaacaccGCCGGCTCCACTACCGCATGgtttaagagctatgctggaaacagc |
| Jak1 | Jak1-4 | CCGAACCGAATCATCACTG | tatcttgtggaaaggacgaaacaccGCCGAACCGAATCATCACTGgtttaagagctatgctggaaacagc |
| Kctd14 | Kctd14-1 | AGTTCTACACTACTACCAT | tatcttgtggaaaggacgaaacaccGAGTTCTACACTACTACCATgtttaagagctatgctggaaacagc |
| Kctd14 | Kctd14-4 | CAGATCTTCGGTGAGCAAG | tatcttgtggaaaggacgaaacaccGCAGATCTTCGGTGAGCAAGgtttaagagctatgctggaaacagc |
| Kctd14 | Kctd14-5 | GTGGCGATGCGGTCGTCAA | tatcttgtggaaaggacgaaacaccGGTGGCGATGCGGTCGTCAAgtttaagagctatgctggaaacagc |
| Kctd18 | Kctd18-1 | TAATTTGGGTACTTGTAGA | tatcttgtggaaaggacgaaacaccGTAATTTGGGTACTTGTAGAgtttaagagctatgctggaaacagc |
| Kctd18 | Kctd18-3 | CAGCATAGAATCCTTAAAG | tatcttgtggaaaggacgaaacaccGCAGCATAGAATCCTTAAAGgtttaagagctatgctggaaacagc |
| Kctd18 | Kctd18-5 | GATATCCTTCGGCTGAATG | tatcttgtggaaaggacgaaacaccGGATATCCTTCGGCTGAATGgtttaagagctatgctggaaacagc |
| Khdrbs1 | Khdrbs1-1 | GATACCCCACGACCACGAG | tatcttgtggaaaggacgaaacaccGGATACCCCACGACCACGAGgtttaagagctatgctggaaacagc |
| Khdrbs1 | Khdrbs1-2 | AGAGCATAAGCTTCACACG | tatcttgtggaaaggacgaaacaccGAGAGCATAAGCTTCACACGgtttaagagctatgctggaaacagc |
| Khdrbs1 | Khdrbs1-3 | AAATATGCCCATTTAAATA | tatcttgtggaaaggacgaaacaccGAAATATGCCCATTTAAATAgtttaagagctatgctggaaacagc |
| Kif18b | Kif18b-2 | TACGTGAGACTTGCGGCCC | tatcttgtggaaaggacgaaacaccGTACGTGAGACTTGCGGCCCgtttaagagctatgctggaaacagc |
| Kif18b | Kif18b-3 | CTCATCTTGGCTACTCGAA | tatcttgtggaaaggacgaaacaccGCTCATCTTGGCTACTCGAAgtttaagagctatgctggaaacagc |
| Kif18b | Kif18b-5 | TTTGCCTATGGCGCCACCG | tatcttgtggaaaggacgaaacaccGTTTGCCTATGGCGCCACCGgtttaagagctatgctggaaacagc |
| Klra9 | Klra9-1 | GATTACATCAAAAGAGAAC | tatcttgtggaaaggacgaaacaccGGATTACATCAAAAGAGAACgtttaagagctatgctggaaacagc |
| Klra9 | Klra9-2 | CAAAGTGATTTCAACTTAA | tatcttgtggaaaggacgaaacaccGCAAAGTGATTTCAACTTAAgtttaagagctatgctggaaacagc |
| Klra9 | Klra9-4 | TTAAGAGTGTTCAGTATCC | tatcttgtggaaaggacgaaacaccGTTAAGAGTGTTCAGTATCCgtttaagagctatgctggaaacagc |
| Kmt2c | Kmt2c-1 | CCGAATCGATGCGTGAGCT | tatcttgtggaaaggacgaaacaccGCCGAATCGATGCGTGAGCTgtttaagagctatgctggaaacagc |
| Kmt2c | Kmt2c-2 | GACCCCGGTCCAAACAGGT | tatcttgtggaaaggacgaaacaccGGACCCCGGTCCAAACAGGTgtttaagagctatgctggaaacagc |
| Kmt2c | Kmt2c-5 | CCGCAGACAAAAGACCTCG | tatcttgtggaaaggacgaaacaccGCCGCAGACAAAAGACCTCGgtttaagagctatgctggaaacagc |
| Knl1 | Knl1-2 | GCACGTGGTCAGGTTGAAA | tatcttgtggaaaggacgaaacaccGGCACGTGGTCAGGTTGAAAgtttaagagctatgctggaaacagc |
| Knl1 | Knl1-3 | CGCCCAGTGATATAACTCC | tatcttgtggaaaggacgaaacaccGCGCCCAGTGATATAACTCCgtttaagagctatgctggaaacagc |
| Knl1 | Knl1-5 | ACAATCGATGAAAACCGTA | tatcttgtggaaaggacgaaacaccGACAATCGATGAAAACCGTAgtttaagagctatgctggaaacagc |
| Kpna2 | Kpna2-3 | AGGGTTCTTGTTTCGACAA | tatcttgtggaaaggacgaaacaccGAGGGTTCTTGTTTCGACAAgtttaagagctatgctggaaacagc |
| Kpna2 | Kpna2-4 | TATTATCAACTTACTGCCA | tatcttgtggaaaggacgaaacaccGTATTATCAACTTACTGCCAgtttaagagctatgctggaaacagc |
| Kpna2 | Kpna2-5 | CGCTTTCTGTCCCAGGTGA | tatcttgtggaaaggacgaaacaccGCGCTTTCTGTCCCAGGTGAgtttaagagctatgctggaaacagc |
| Krcc1 | Krcc1-1 | GCATGATCTTGGGTAGGAA | tatcttgtggaaaggacgaaacaccGGCATGATCTTGGGTAGGAAgtttaagagctatgctggaaacagc |
| Krcc1 | Krcc1-2 | AGTTCACAGACAGAAGACC | tatcttgtggaaaggacgaaacaccGAGTTCACAGACAGAAGACCgtttaagagctatgctggaaacagc |
| Krcc1 | Krcc1-5 | ATTGTATTCTTGCTGACTA | tatcttgtggaaaggacgaaacaccGATTGTATTCTTGCTGACTAgtttaagagctatgctggaaacagc |
| Krt10 | Krt10-1 | AAATACTACAAAACCATCG | tatcttgtggaaaggacgaaacaccGAAATACTACAAAACCATCGgtttaagagctatgctggaaacagc |
| Krt10 | Krt10-4 | GCTTTGGGGGCTCATCAGG | tatcttgtggaaaggacgaaacaccGGCTTTGGGGGCTCATCAGGgtttaagagctatgctggaaacagc |
| Krt10 | Krt10-5 | CGGTGGCTCTTTTAGCCGT | tatcttgtggaaaggacgaaacaccGCGGTGGCTCTTTTAGCCGTgtttaagagctatgctggaaacagc |
| L3mbtl2 | L3mbtl2-1 | GTACCCCTCTATGACCAGT | tatcttgtggaaaggacgaaacaccGGTACCCCTCTATGACCAGTgtttaagagctatgctggaaacagc |
| L3mbtl2 | L3mbtl2-4 | TCACGTACCCGCATGGCCG | tatcttgtggaaaggacgaaacaccGTCACGTACCCGCATGGCCGgtttaagagctatgctggaaacagc |
| L3mbtl2 | L3mbtl2-5 | AGTAATCGGGGGTCGCCTC | tatcttgtggaaaggacgaaacaccGAGTAATCGGGGGTCGCCTCgtttaagagctatgctggaaacagc |
| Lars2 | Lars2-1 | CGTGCAATGGTGTCGCTGA | tatcttgtggaaaggacgaaacaccGCGTGCAATGGTGTCGCTGAgtttaagagctatgctggaaacagc |
| Lars2 | Lars2-3 | ATCTATGGCATCTCCCACG | tatcttgtggaaaggacgaaacaccGATCTATGGCATCTCCCACGgtttaagagctatgctggaaacagc |
| Lars2 | Lars2-5 | CCTGACTCGGAAAGCCCGT | tatcttgtggaaaggacgaaacaccGCCTGACTCGGAAAGCCCGTgtttaagagctatgctggaaacagc |
| Ldlr | Ldlr-1 | CAGTCGACATCCCCGTCGC | tatcttgtggaaaggacgaaacaccGCAGTCGACATCCCCGTCGCgtttaagagctatgctggaaacagc |
| Ldlr | Ldlr-4 | CCGGTTGGTGGACCTCCAC | tatcttgtggaaaggacgaaacaccGCCGGTTGGTGGACCTCCACgtttaagagctatgctggaaacagc |
| Ldlr | Ldlr-5 | CATCTTCCGGACCTCGTGG | tatcttgtggaaaggacgaaacaccGCATCTTCCGGACCTCGTGGgtttaagagctatgctggaaacagc |
| Lifr | Lifr-1 | ATCTGACGATGCATGCAAA | tatcttgtggaaaggacgaaacaccGATCTGACGATGCATGCAAAgtttaagagctatgctggaaacagc |
| Lifr | Lifr-3 | CGAATGCTCACCGAGTGAG | tatcttgtggaaaggacgaaacaccGCGAATGCTCACCGAGTGAGgtttaagagctatgctggaaacagc |
| Lifr | Lifr-5 | GACGATGTGTACGGAACGG | tatcttgtggaaaggacgaaacaccGGACGATGTGTACGGAACGGgtttaagagctatgctggaaacagc |
| Lig4 | Lig4-1 | TTAAACCAGAGTACGTCAG | tatcttgtggaaaggacgaaacaccGTTAAACCAGAGTACGTCAGgtttaagagctatgctggaaacagc |
| Lig4 | Lig4-2 | GCGCGAGACTCTTAGGAAG | tatcttgtggaaaggacgaaacaccGGCGCGAGACTCTTAGGAAGgtttaagagctatgctggaaacagc |
| Lig4 | Lig4-3 | ACCGTTTCTGGAGAAGTAC | tatcttgtggaaaggacgaaacaccGACCGTTTCTGGAGAAGTACgtttaagagctatgctggaaacagc |
| Lin52 | Lin52-3 | TCGGTGGAGAGCTTGTGAT | tatcttgtggaaaggacgaaacaccGTCGGTGGAGAGCTTGTGATgtttaagagctatgctggaaacagc |
| Lin52 | Lin52-4 | TCTCAGCCATCCATTTCGG | tatcttgtggaaaggacgaaacaccGTCTCAGCCATCCATTTCGGgtttaagagctatgctggaaacagc |
| Lin52 | Lin52-5 | ATGACATTGATATGTTGAA | tatcttgtggaaaggacgaaacaccGATGACATTGATATGTTGAAgtttaagagctatgctggaaacagc |
| Lrrd1 | Lrrd1-3 | GCCTATCATGTTTCCAGTA | tatcttgtggaaaggacgaaacaccGGCCTATCATGTTTCCAGTAgtttaagagctatgctggaaacagc |
| Lrrd1 | Lrrd1-4 | CATCTAGCTGACAATAAAC | tatcttgtggaaaggacgaaacaccGCATCTAGCTGACAATAAACgtttaagagctatgctggaaacagc |
| Lrrd1 | Lrrd1-5 | AGATTCCAAATTTACGCAG | tatcttgtggaaaggacgaaacaccGAGATTCCAAATTTACGCAGgtttaagagctatgctggaaacagc |
| Lxn | Lxn-2 | TCACATTCGAAGGAGAAAT | tatcttgtggaaaggacgaaacaccGTCACATTCGAAGGAGAAATgtttaagagctatgctggaaacagc |
| Lxn | Lxn-3 | AAGTTCTTTACCCCCAAAT | tatcttgtggaaaggacgaaacaccGAAGTTCTTTACCCCCAAATgtttaagagctatgctggaaacagc |

| | | | |
|---|---|---|---|
| Lxn | Lxn-5 | CTGCTGGACCGTCTGCACC | tatcttgtggaaaggacgaaacaccGCTGCTGGACCGTCTGCACCgtttaagagctatgctggaaacagc |
| Ly6k | Ly6k-1 | TTGACAGCACTTATAAAAT | tatcttgtggaaaggacgaaacaccGTTGACAGCACTTATAAAATgtttaagagctatgctggaaacagc |
| Ly6k | Ly6k-3 | GCTAGGCGGGCTGACTACG | tatcttgtggaaaggacgaaacaccGGCTAGGCGGGCTGACTACGgtttaagagctatgctggaaacagc |
| Ly6k | Ly6k-5 | CACTGGGACGGATTCGAGC | tatcttgtggaaaggacgaaacaccGCACTGGGACGGATTCGAGCgtttaagagctatgctggaaacagc |
| Lysmd3 | Lysmd3-3 | GCTCTTGAAAATACGGAAC | tatcttgtggaaaggacgaaacaccGGCTCTTGAAAATACGGAACgtttaagagctatgctggaaacagc |
| Lysmd3 | Lysmd3-4 | TTTGTTATCAGGTTCAAAT | tatcttgtggaaaggacgaaacaccGTTTGTTATCAGGTTCAAATgtttaagagctatgctggaaacagc |
| Lysmd3 | Lysmd3-5 | CCGTATTACGGAGCAGACT | tatcttgtggaaaggacgaaacaccGCCGTATTACGGAGCAGACTgtttaagagctatgctggaaacagc |
| Maea | Maea-2 | ACAGCTGGGCAACTACTCA | tatcttgtggaaaggacgaaacaccGACAGCTGGGCAACTACTCAgtttaagagctatgctggaaacagc |
| Maea | Maea-3 | ATGTGGAAGCGGAAGCGCA | tatcttgtggaaaggacgaaacaccGATGTGGAAGCGGAAGCGCAgtttaagagctatgctggaaacagc |
| Maea | Maea-5 | GTTTCGATATGATAACTAC | tatcttgtggaaaggacgaaacaccGGTTTCGATATGATAACTACgtttaagagctatgctggaaacagc |
| Marchf6 | Marchf6-2 | GTTCCCTCTCATTTGTGGC | tatcttgtggaaaggacgaaacaccGGTTCCCTCTCATTTGTGGCgtttaagagctatgctggaaacagc |
| Marchf6 | Marchf6-4 | CTTTGTGCGTTCATCAGCC | tatcttgtggaaaggacgaaacaccGCTTTGTGCGTTCATCAGCCgtttaagagctatgctggaaacagc |
| Marchf6 | Marchf6-5 | TGGCACTGCAATACGATAC | tatcttgtggaaaggacgaaacaccGTGGCACTGCAATACGATACgtttaagagctatgctggaaacagc |
| Marf1 | Marf1-1 | GCGATCAGCTTGCTGAATG | tatcttgtggaaaggacgaaacaccGGCGATCAGCTTGCTGAATGgtttaagagctatgctggaaacagc |
| Marf1 | Marf1-2 | GCCGAATGGAGTCAAAGTC | tatcttgtggaaaggacgaaacaccGGCCGAATGGAGTCAAAGTCgtttaagagctatgctggaaacagc |
| Marf1 | Marf1-4 | GAATCTTCAGTGGTAACCG | tatcttgtggaaaggacgaaacaccGGAATCTTCAGTGGTAACCGgtttaagagctatgctggaaacagc |
| Mars1 | Mars1-1 | TGGCGAAGGCGACAATACC | tatcttgtggaaaggacgaaacaccGTGGCGAAGGCGACAATACCgtttaagagctatgctggaaacagc |
| Mars1 | Mars1-2 | GGGGGACATTGTTGACATA | tatcttgtggaaaggacgaaacaccGGGGGGACATTGTTGACATAgtttaagagctatgctggaaacagc |
| Mars1 | Mars1-5 | CCTTAGGCCGGGTAAGGAA | tatcttgtggaaaggacgaaacaccGCCTTAGGCCGGGTAAGGAAgtttaagagctatgctggaaacagc |
| Mbd2 | Mbd2-1 | GAAAGGGACAGGATCCCGC | tatcttgtggaaaggacgaaacaccGGAAAGGGACAGGATCCCGCgtttaagagctatgctggaaacagc |
| Mbd2 | Mbd2-3 | AGGAAGTGATCCGAAAATC | tatcttgtggaaaggacgaaacaccGAGGAAGTGATCCGAAAATCgtttaagagctatgctggaaacagc |
| Mbd2 | Mbd2-5 | TTCAGAAGTAAACCTCAGC | tatcttgtggaaaggacgaaacaccGTTCAGAAGTAAACCTCAGCgtttaagagctatgctggaaacagc |
| Mcl1 | Mcl1-2 | CGTCACCGCGTCGGCCGAA | tatcttgtggaaaggacgaaacaccGCGTCACCGCGTCGGCCGAAgtttaagagctatgctggaaacagc |
| Mcl1 | Mcl1-4 | ACGGCCGGGCGCTTGCCGA | tatcttgtggaaaggacgaaacaccGACGGCCGGGCGCTTGCCGAgtttaagagctatgctggaaacagc |
| Mcl1 | Mcl1-5 | CGAGATGATCTCCAGCGAC | tatcttgtggaaaggacgaaacaccGCGAGATGATCTCCAGCGACgtttaagagctatgctggaaacagc |
| Mcm2 | Mcm2-1 | TACCAACGTATCCGCATCC | tatcttgtggaaaggacgaaacaccGTACCAACGTATCCGCATCCgtttaagagctatgctggaaacagc |
| Mcm2 | Mcm2-2 | CTGATCCGTACCAGTGGCG | tatcttgtggaaaggacgaaacaccGCTGATCCGTACCAGTGGCGgtttaagagctatgctggaaacagc |
| Mcm2 | Mcm2-3 | GGCCACTCGGTGCGCGAGT | tatcttgtggaaaggacgaaacaccGGGCCACTCGGTGCGCGAGTgtttaagagctatgctggaaacagc |
| Mcm6 | Mcm6-1 | ACCTTCTTAAATAGGCACG | tatcttgtggaaaggacgaaacaccGACCTTCTTAAATAGGCACGgtttaagagctatgctggaaacagc |
| Mcm6 | Mcm6-3 | AGACTAACTCCCGAGTCAG | tatcttgtggaaaggacgaaacaccGAGACTAACTCCCGAGTCAGgtttaagagctatgctggaaacagc |
| Mcm6 | Mcm6-5 | GGCTGCTCACCCGAATCAG | tatcttgtggaaaggacgaaacaccGGGCTGCTCACCCGAATCAGgtttaagagctatgctggaaacagc |
| Mcm7 | Mcm7-1 | GAACCACTGTGATGCTACG | tatcttgtggaaaggacgaaacaccGGAACCACTGTGATGCTACGgtttaagagctatgctggaaacagc |
| Mcm7 | Mcm7-2 | AGTGCCAGACCAATCGCTC | tatcttgtggaaaggacgaaacaccGAGTGCCAGACCAATCGCTCgtttaagagctatgctggaaacagc |
| Mcm7 | Mcm7-4 | GGAAATTGTTAACTGTGCG | tatcttgtggaaaggacgaaacaccGGGAAATTGTTAACTGTGCGgtttaagagctatgctggaaacagc |
| Mcm8 | Mcm8-3 | GAAATACATCTCTATACGA | tatcttgtggaaaggacgaaacaccGGAAATACATCTCTATACGAgtttaagagctatgctggaaacagc |
| Mcm8 | Mcm8-4 | GTTCACATTCTATCGTCCG | tatcttgtggaaaggacgaaacaccGGTTCACATTCTATCGTCCGgtttaagagctatgctggaaacagc |
| Mcm8 | Mcm8-5 | GATCTCTCGGATGGCATAG | tatcttgtggaaaggacgaaacaccGGATCTCTCGGATGGCATAGgtttaagagctatgctggaaacagc |
| Me3 | Me3-1 | TCAGGGGTACGGGTCGAGG | tatcttgtggaaaggacgaaacaccGTCAGGGGTACGGGTCGAGGgtttaagagctatgctggaaacagc |
| Me3 | Me3-3 | ACCTGGGCTGCTACGGCAT | tatcttgtggaaaggacgaaacaccGACCTGGGCTGCTACGGCATgtttaagagctatgctggaaacagc |
| Me3 | Me3-5 | AAACACCAGCGTGTCCGTG | tatcttgtggaaaggacgaaacaccGAAACACCAGCGTGTCCGTGgtttaagagctatgctggaaacagc |
| Meaf6 | Meaf6-1 | TAGGAAACACTTGCAAACT | tatcttgtggaaaggacgaaacaccGTAGGAAACACTTGCAAACTgtttaagagctatgctggaaacagc |
| Meaf6 | Meaf6-2 | GCTTTTGAAGGAAGCTACC | tatcttgtggaaaggacgaaacaccGGCTTTTGAAGGAAGCTACCgtttaagagctatgctggaaacagc |
| Meaf6 | Meaf6-4 | CAAAAACGACCGGAGGAAC | tatcttgtggaaaggacgaaacaccGCAAAAACGACCGGAGGAACgtttaagagctatgctggaaacagc |
| Med1 | Med1-1 | CGGACCATCCTCTGAATAC | tatcttgtggaaaggacgaaacaccGCGGACCATCCTCTGAATACgtttaagagctatgctggaaacagc |
| Med1 | Med1-3 | GGAACGGGGGGTCTACCAT | tatcttgtggaaaggacgaaacaccGGGAACGGGGGGTCTACCATgtttaagagctatgctggaaacagc |
| Med1 | Med1-4 | TTATCTCACCCCGCGGAGT | tatcttgtggaaaggacgaaacaccGTTATCTCACCCCGCGGAGTgtttaagagctatgctggaaacagc |
| Med12 | Med12-2 | GCAGAAGCGACGACGCAAC | tatcttgtggaaaggacgaaacaccGGCAGAAGCGACGACGCAACgtttaagagctatgctggaaacagc |
| Med12 | Med12-3 | AGATCACGAGCTTCGCCCT | tatcttgtggaaaggacgaaacaccGAGATCACGAGCTTCGCCCTgtttaagagctatgctggaaacagc |
| Med12 | Med12-5 | TCGGAATCCAATATGCGCT | tatcttgtggaaaggacgaaacaccGTCGGAATCCAATATGCGCTgtttaagagctatgctggaaacagc |
| Med18 | Med18-3 | GCGCTCCATGGACAGGGCA | tatcttgtggaaaggacgaaacaccGGCGCTCCATGGACAGGGCAgtttaagagctatgctggaaacagc |
| Med18 | Med18-4 | GCCAGCCCGTTCGTCCTAA | tatcttgtggaaaggacgaaacaccGGCCAGCCCGTTCGTCCTAAgtttaagagctatgctggaaacagc |
| Med18 | Med18-5 | ACCATCTCGTGATCAAGGA | tatcttgtggaaaggacgaaacaccGACCATCTCGTGATCAAGGAgtttaagagctatgctggaaacagc |
| Med20 | Med20-1 | CCTTGGCTGCCCAGGGTAG | tatcttgtggaaaggacgaaacaccGCCTTGGCTGCCCAGGGTAGgtttaagagctatgctggaaacagc |
| Med20 | Med20-2 | CCAGCAAGATTGAGACCCG | tatcttgtggaaaggacgaaacaccGCCAGCAAGATTGAGACCCGgtttaagagctatgctggaaacagc |
| Med20 | Med20-5 | ACAAGGGCCATACTCCACC | tatcttgtggaaaggacgaaacaccGACAAGGGCCATACTCCACCgtttaagagctatgctggaaacagc |
| Med22 | Med22-2 | CGATGGCTTCATTCACAGA | tatcttgtggaaaggacgaaacaccGCGATGGCTTCATTCACAGAgtttaagagctatgctggaaacagc |
| Med22 | Med22-3 | GGTGAGTCACTGATGAAGC | tatcttgtggaaaggacgaaacaccGGGTGAGTCACTGATGAAGCgtttaagagctatgctggaaacagc |
| Med22 | Med22-4 | CGATGGTGCAGGTTCGAGC | tatcttgtggaaaggacgaaacaccGCGATGGTGCAGGTTCGAGCgtttaagagctatgctggaaacagc |
| Med24 | Med24-1 | TTGATGAATTTCCGGGCGA | tatcttgtggaaaggacgaaacaccGTTGATGAATTTCCGGGCGAgtttaagagctatgctggaaacagc |
| Med24 | Med24-3 | GACTCATTGAGTCCCCCGA | tatcttgtggaaaggacgaaacaccGGACTCATTGAGTCCCCCGAgtttaagagctatgctggaaacagc |
| Med24 | Med24-5 | GGCGTACTTCAGGTACGAC | tatcttgtggaaaggacgaaacaccGGGCGTACTTCAGGTACGACgtttaagagctatgctggaaacagc |
| Med4 | Med4-1 | AGATCTGGAGGTCTTGTCG | tatcttgtggaaaggacgaaacaccGAGATCTGGAGGTCTTGTCGgtttaagagctatgctggaaacagc |
| Med4 | Med4-2 | AGTTGCTAATTCACCGAGA | tatcttgtggaaaggacgaaacaccGAGTTGCTAATTCACCGAGAgtttaagagctatgctggaaacagc |
| Med4 | Med4-4 | GGTTGGGTATGGCCTCCGA | tatcttgtggaaaggacgaaacaccGGGTTGGGTATGGCCTCCGAgtttaagagctatgctggaaacagc |
| Mettl14 | Mettl14-1 | CGATGAGATTGCAGCACCT | tatcttgtggaaaggacgaaacaccGCGATGAGATTGCAGCACCTgtttaagagctatgctggaaacagc |
| Mettl14 | Mettl14-2 | AGACCTCAGAATTTCATCA | tatcttgtggaaaggacgaaacaccGAGACCTCAGAATTTCATCAgtttaagagctatgctggaaacagc |
| Mettl14 | Mettl14-5 | AAATAGCAAAGATGAACAG | tatcttgtggaaaggacgaaacaccGAAATAGCAAAGATGAACAGgtttaagagctatgctggaaacagc |
| Mettl3 | Mettl3-1 | AAGTTTCGCTCTCGAGGTC | tatcttgtggaaaggacgaaacaccGAAGTTTCGCTCTCGAGGTCgtttaagagctatgctggaaacagc |
| Mettl3 | Mettl3-3 | CAGGTGATTACCGTAGAGA | tatcttgtggaaaggacgaaacaccGCAGGTGATTACCGTAGAGAgtttaagagctatgctggaaacagc |
| Mettl3 | Mettl3-4 | CTTAGGGCCGCTAGAGGTA | tatcttgtggaaaggacgaaacaccGCTTAGGGCCGCTAGAGGTAgtttaagagctatgctggaaacagc |
| Mfsd14b | Mfsd14b-2 | AGAGGTATGTCCCAATGGC | tatcttgtggaaaggacgaaacaccGAGAGGTATGTCCCAATGGCgtttaagagctatgctggaaacagc |
| Mfsd14b | Mfsd14b-3 | AATCCTCATTAGTGGGATA | tatcttgtggaaaggacgaaacaccGAATCCTCATTAGTGGGATAgtttaagagctatgctggaaacagc |
| Mfsd14b | Mfsd14b-4 | GGTAAAGAAGACCGTGCCA | tatcttgtggaaaggacgaaacaccGGGTAAAGAAGACCGTGCCAgtttaagagctatgctggaaacagc |
| Milr1 | Milr1-1 | GCAGTGGATTGTACGAGGG | tatcttgtggaaaggacgaaacaccGGCAGTGGATTGTACGAGGGgtttaagagctatgctggaaacagc |
| Milr1 | Milr1-3 | GTACCCCAGAAGAGCGAAT | tatcttgtggaaaggacgaaacaccGGTACCCCAGAAGAGCGAATgtttaagagctatgctggaaacagc |
| Milr1 | Milr1-4 | TCTCCAATGCCAACGAGTC | tatcttgtggaaaggacgaaacaccGTCTCCAATGCCAACGAGTCgtttaagagctatgctggaaacagc |

| | | | |
|---|---|---|---|
| Mme | Mme-1 | GCTCGACTGATTCAGAATA | tatcttgtggaaaggacgaaacaccGGCTCGACTGATTCAGAATAgtttaagagctatgctggaaacagc |
| Mme | Mme-2 | AAATTACTGTATCGGGAAC | tatcttgtggaaaggacgaaacaccGAAATTACTGTATCGGGAACgtttaagagctatgctggaaacagc |
| Mme | Mme-5 | GAGACTACTATGAGTGCAC | tatcttgtggaaaggacgaaacaccGGAGACTACTATGAGTGCACgtttaagagctatgctggaaacagc |
| Mmp20 | Mmp20-1 | GTATAATACTTGTCAAGAT | tatcttgtggaaaggacgaaacaccGGTATAATACTTGTCAAGATgtttaagagctatgctggaaacagc |
| Mmp20 | Mmp20-2 | GGCGGTAGTTAGCCACATC | tatcttgtggaaaggacgaaacaccGGGCGGTAGTTAGCCACATCgtttaagagctatgctggaaacagc |
| Mmp20 | Mmp20-5 | ACCCATTTGATGGGCCTCG | tatcttgtggaaaggacgaaacaccGACCCATTTGATGGGCCTCGgtttaagagctatgctggaaacagc |
| Mre11a | Mre11a-1 | TGCCGTGGATACTAAATAC | tatcttgtggaaaggacgaaacaccGTGCCGTGGATACTAAATACgtttaagagctatgctggaaacagc |
| Mre11a | Mre11a-2 | GAATTGTAATTACCCCCGT | tatcttgtggaaaggacgaaacaccGGAATTGTAATTACCCCCGTgtttaagagctatgctggaaacagc |
| Mre11a | Mre11a-5 | ACCGCCGCACGGTGCGGAG | tatcttgtggaaaggacgaaacaccGACCGCCGCACGGTGCGGAGgtttaagagctatgctggaaacagc |
| Mrgprb1 | Mrgprb1-2 | GTGCTTTGGGTCTTGTCCC | tatcttgtggaaaggacgaaacaccGGTGCTTTGGGTCTTGTCCCgtttaagagctatgctggaaacagc |
| Mrgprb1 | Mrgprb1-4 | CCCAGAAACCATAACACTG | tatcttgtggaaaggacgaaacaccGCCCAGAAACCATAACACTGgtttaagagctatgctggaaacagc |
| Mrgprb1 | Mrgprb1-5 | CCCAACTAAGGCAATGATG | tatcttgtggaaaggacgaaacaccGCCCAACTAAGGCAATGATGgtttaagagctatgctggaaacagc |
| Mrpl1 | Mrpl1-1 | CTTAAAACGCTTATACCCA | tatcttgtggaaaggacgaaacaccGCTTAAAACGCTTATACCCAgtttaagagctatgctggaaacagc |
| Mrpl1 | Mrpl1-4 | TTAACATTAGATATGGCAT | tatcttgtggaaaggacgaaacaccGTTAACATTAGATATGGCATgtttaagagctatgctggaaacagc |
| Mrpl1 | Mrpl1-5 | TTTTAGATCATGGATGACG | tatcttgtggaaaggacgaaacaccGTTTTAGATCATGGATGACGgtttaagagctatgctggaaacagc |
| Mrps21 | Mrps21-3 | CGTCGTCGACTTATGACTT | tatcttgtggaaaggacgaaacaccGCGTCGTCGACTTATGACTTgtttaagagctatgctggaaacagc |
| Mrps21 | Mrps21-4 | ACTTCGGTAAGCCCATCCG | tatcttgtggaaaggacgaaacaccGACTTCGGTAAGCCCATCCGgtttaagagctatgctggaaacagc |
| Mrps21 | Mrps21-5 | CAGAATCCTCACCACGGAT | tatcttgtggaaaggacgaaacaccGCAGAATCCTCACCACGGATgtttaagagctatgctggaaacagc |
| Msh2 | Msh2-1 | AAAGTCGCCGCGGTCGAAG | tatcttgtggaaaggacgaaacaccGAAAGTCGCCGCGGTCGAAGgtttaagagctatgctggaaacagc |
| Msh2 | Msh2-3 | TGGGTCTTGAACACCTCGC | tatcttgtggaaaggacgaaacaccGTGGGTCTTGAACACCTCGCgtttaagagctatgctggaaacagc |
| Msh2 | Msh2-5 | TGGCAAGCCGGTTAAGGTC | tatcttgtggaaaggacgaaacaccGTGGCAAGCCGGTTAAGGTCgtttaagagctatgctggaaacagc |
| Msh6 | Msh6-1 | GCGGTATCCGCCTCGTCGC | tatcttgtggaaaggacgaaacaccGGCGGTATCCGCCTCGTCGCgtttaagagctatgctggaaacagc |
| Msh6 | Msh6-2 | CGGATGAACGTTCCATCAA | tatcttgtggaaaggacgaaacaccGCGGATGAACGTTCCATCAAgtttaagagctatgctggaaacagc |
| Msh6 | Msh6-5 | GTGGCGGTCATCCGAAAAC | tatcttgtggaaaggacgaaacaccGGTGGCGGTCATCCGAAAACgtttaagagctatgctggaaacagc |
| Msrb2 | Msrb2-1 | GCTTCGCTTAGAGAGCGCT | tatcttgtggaaaggacgaaacaccGGCTTCGCTTAGAGAGCGCTgtttaagagctatgctggaaacagc |
| Msrb2 | Msrb2-2 | TCAAATACATTCCACTGAA | tatcttgtggaaaggacgaaacaccGTCAAATACATTCCACTGAAgtttaagagctatgctggaaacagc |
| Msrb2 | Msrb2-3 | TTTGAACAACAAGGAGACA | tatcttgtggaaaggacgaaacaccGTTTGAACAACAAGGAGACAgtttaagagctatgctggaaacagc |
| Msx1 | Msx1-3 | TCGGCCATTTCTCAGTCGG | tatcttgtggaaaggacgaaacaccGTCGGCCATTTCTCAGTCGGgtttaagagctatgctggaaacagc |
| Msx1 | Msx1-4 | TCCGAAGGGGCTCAGGCAG | tatcttgtggaaaggacgaaacaccGTCCGAAGGGGCTCAGGCAGgtttaagagctatgctggaaacagc |
| Msx1 | Msx1-5 | CGGCCACCGCAACCGCCAT | tatcttgtggaaaggacgaaacaccGCGGCCACCGCAACCGCCATgtttaagagctatgctggaaacagc |
| Mtarc2 | Mtarc2-2 | GTGGATCTTATTCGAAGAA | tatcttgtggaaaggacgaaacaccGGTGGATCTTATTCGAAGAAgtttaagagctatgctggaaacagc |
| Mtarc2 | Mtarc2-4 | CTGTGGCAAAGTGCGCGAC | tatcttgtggaaaggacgaaacaccGCTGTGGCAAAGTGCGCGACgtttaagagctatgctggaaacagc |
| Mtarc2 | Mtarc2-5 | AGACCGAGTGCACGGACAT | tatcttgtggaaaggacgaaacaccGAGACCGAGTGCACGGACATgtttaagagctatgctggaaacagc |
| Mterf2 | Mterf2-1 | TGAGGCCAATGCTAAAGTG | tatcttgtggaaaggacgaaacaccGTGAGGCCAATGCTAAAGTGgtttaagagctatgctggaaacagc |
| Mterf2 | Mterf2-2 | TAACACCAAAAGGAAACTC | tatcttgtggaaaggacgaaacaccGTAACACCAAAAGGAAACTCgtttaagagctatgctggaaacagc |
| Mterf2 | Mterf2-3 | CAAATATTTTGAAAGAACT | tatcttgtggaaaggacgaaacaccGCAAATATTTTGAAAGAACTgtttaagagctatgctggaaacagc |
| Myh15 | Myh15-3 | GTAAGCTGACTTCCCAAAA | tatcttgtggaaaggacgaaacaccGGTAAGCTGACTTCCCAAAAgtttaagagctatgctggaaacagc |
| Myh15 | Myh15-4 | CATCATATAGCATCCAACT | tatcttgtggaaaggacgaaacaccGCATCATATAGCATCCAACTgtttaagagctatgctggaaacagc |
| Myh15 | Myh15-5 | TCCATCTCATACTAGGTAC | tatcttgtggaaaggacgaaacaccGTCCATCTCATACTAGGTACgtttaagagctatgctggaaacagc |
| Naa38 | Naa38-3 | TCTGTCATACGGATACGCA | tatcttgtggaaaggacgaaacaccGTCTGTCATACGGATACGCAgtttaagagctatgctggaaacagc |
| Naa38 | Naa38-4 | ACAGTCGCGATCAGTGCAC | tatcttgtggaaaggacgaaacaccGACAGTCGCGATCAGTGCACgtttaagagctatgctggaaacagc |
| Naa38 | Naa38-5 | CGCGACTGTAATGTCATCC | tatcttgtggaaaggacgaaacaccGCGCGACTGTAATGTCATCCgtttaagagctatgctggaaacagc |
| Naca | Naca-10 | CTTTAGGCTATGTCCAAAC | tatcttgtggaaaggacgaaacaccGCTTTAGGCTATGTCCAAACgtttaagagctatgctggaaacagc |
| Naca | Naca-8 | TCTGCTTGGCTTTACTAAC | tatcttgtggaaaggacgaaacaccGTCTGCTTGGCTTTACTAACgtttaagagctatgctggaaacagc |
| Naca | Naca-9 | TTCTCACTTCGACTCTGCT | tatcttgtggaaaggacgaaacaccGTTCTCACTTCGACTCTGCTgtttaagagctatgctggaaacagc |
| Nagk | Nagk-2 | GATTAAGTAGTTTTCACTC | tatcttgtggaaaggacgaaacaccGGATTAAGTAGTTTTCACTCgtttaagagctatgctggaaacagc |
| Nagk | Nagk-4 | GTAGGCTTATCAACCCTGA | tatcttgtggaaaggacgaaacaccGGTAGGCTTATCAACCCTGAgtttaagagctatgctggaaacagc |
| Nagk | Nagk-5 | GCTTGGTGTGCAATCCAGT | tatcttgtggaaaggacgaaacaccGGCTTGGTGTGCAATCCAGTgtttaagagctatgctggaaacagc |
| Nanog | Nanog-2 | CACTCTGCAGGTTAAGACC | tatcttgtggaaaggacgaaacaccGCACTCTGCAGGTTAAGACCgtttaagagctatgctggaaacagc |
| Nanog | Nanog-3 | GACCTGGTTTCAAAACCAA | tatcttgtggaaaggacgaaacaccGGACCTGGTTTCAAAACCAAgtttaagagctatgctggaaacagc |
| Nanog | Nanog-4 | GCTATCTGGTGAACGCATC | tatcttgtggaaaggacgaaacaccGGCTATCTGGTGAACGCATCgtttaagagctatgctggaaacagc |
| Nanos2 | Nanos2-2 | ATATGGTAGGCAGACATCC | tatcttgtggaaaggacgaaacaccGATATGGTAGGCAGACATCCgtttaagagctatgctggaaacagc |
| Nanos2 | Nanos2-4 | ACCAGCTGAAGACGCCTGA | tatcttgtggaaaggacgaaacaccGACCAGCTGAAGACGCCTGAgtttaagagctatgctggaaacagc |
| Nanos2 | Nanos2-5 | ACACATAGTGCCTCAGGAT | tatcttgtggaaaggacgaaacaccGACACATAGTGCCTCAGGATgtttaagagctatgctggaaacagc |
| Nanp | Nanp-2 | TAAGCGTGTAGATTTCCAC | tatcttgtggaaaggacgaaacaccGTAAGCGTGTAGATTTCCACgtttaagagctatgctggaaacagc |
| Nanp | Nanp-3 | CAAAGGAGGTGCTGACAAT | tatcttgtggaaaggacgaaacaccGCAAAGGAGGTGCTGACAATgtttaagagctatgctggaaacagc |
| Nanp | Nanp-4 | CACTGGGAAGAAGCAATCC | tatcttgtggaaaggacgaaacaccGCACTGGGAAGAAGCAATCCgtttaagagctatgctggaaacagc |
| Nbn | Nbn-1 | TTTAACTGCCGACATGACA | tatcttgtggaaaggacgaaacaccGTTTAACTGCCGACATGACAgtttaagagctatgctggaaacagc |
| Nbn | Nbn-3 | ATTCCTACATCAACAACGC | tatcttgtggaaaggacgaaacaccGATTCCTACATCAACAACGCgtttaagagctatgctggaaacagc |
| Nbn | Nbn-4 | CTATTCCTGAAGCGGAGAT | tatcttgtggaaaggacgaaacaccGCTATTCCTGAAGCGGAGATgtttaagagctatgctggaaacagc |
| Ncapg2 | Ncapg2-2 | TGCCTGTGGAACGAACCCG | tatcttgtggaaaggacgaaacaccGTGCCTGTGGAACGAACCCGgtttaagagctatgctggaaacagc |
| Ncapg2 | Ncapg2-3 | CGGCCTGTATCTCGACGCC | tatcttgtggaaaggacgaaacaccGCGGCCTGTATCTCGACGCCgtttaagagctatgctggaaacagc |
| Ncapg2 | Ncapg2-5 | GAATCGCGCAGCTGCCAGG | tatcttgtggaaaggacgaaacaccGGAATCGCGCAGCTGCCAGGgtttaagagctatgctggaaacagc |
| Ndufb11b | Ndufb11b-2 | CGGTCCGCTGCGGCCCGAA | tatcttgtggaaaggacgaaacaccGCGGTCCGCTGCGGCCCGAAgtttaagagctatgctggaaacagc |
| Ndufb11b | Ndufb11b-3 | CGTCGTAACCATGGAAGTG | tatcttgtggaaaggacgaaacaccGCGTCGTAACCATGGAAGTGgtttaagagctatgctggaaacagc |
| Ndufb11b | Ndufb11b-5 | AAGACGGCGTGCATATTCA | tatcttgtggaaaggacgaaacaccGAAGACGGCGTGCATATTCAgtttaagagctatgctggaaacagc |
| Ndufb4 | Ndufb4-1 | ATATTTGCTGATCTTGCAT | tatcttgtggaaaggacgaaacaccGATATTTGCTGATCTTGCATgtttaagagctatgctggaaacagc |
| Ndufb4 | Ndufb4-2 | GCATAGGTCCAGCGAATCA | tatcttgtggaaaggacgaaacaccGGCATAGGTCCAGCGAATCAgtttaagagctatgctggaaacagc |
| Ndufb4 | Ndufb4-5 | CGTGTCTCCGGAGACCAGA | tatcttgtggaaaggacgaaacaccGCGTGTCTCCGGAGACCAGAgtttaagagctatgctggaaacagc |
| Ndufc2 | Ndufc2-1 | GACCCGCGGCTTGTCTACA | tatcttgtggaaaggacgaaacaccGGACCCGCGGCTTGTCTACAgtttaagagctatgctggaaacagc |
| Ndufc2 | Ndufc2-2 | GGCTACTGCACGGGCCTGA | tatcttgtggaaaggacgaaacaccGGGCTACTGCACGGGCCTGAgtttaagagctatgctggaaacagc |
| Ndufc2 | Ndufc2-4 | TGCGACCGGTGATGAGAGC | tatcttgtggaaaggacgaaacaccGTGCGACCGGTGATGAGAGCgtttaagagctatgctggaaacagc |
| Necab1 | Necab1-1 | ACTGCTGACATTGAACTGC | tatcttgtggaaaggacgaaacaccGACTGCTGACATTGAACTGCgtttaagagctatgctggaaacagc |
| Necab1 | Necab1-4 | GTGACAAATTGTTCCAAAT | tatcttgtggaaaggacgaaacaccGGTGACAAATTGTTCCAAATgtttaagagctatgctggaaacagc |
| Necab1 | Necab1-5 | TATGCTTTAAATTCCTCAA | tatcttgtggaaaggacgaaacaccGTATGCTTTAAATTCCTCAAgtttaagagctatgctggaaacagc |
| Nelfcd | Nelfcd-1 | GATCAAGCACTTCGACCCT | tatcttgtggaaaggacgaaacaccGGATCAAGCACTTCGACCCTgtttaagagctatgctggaaacagc |

| | | | |
|---|---|---|---|
| Nelfcd | Nelfcd-2 | TCTGATGCGGGGTACCAAG | tatcttgtggaaaggacgaaacaccGTCTGATGCGGGGTACCAAGgtttaagagctatgctggaaacagc |
| Nelfcd | Nelfcd-4 | GAAGTCCAGCGCTTCGCCC | tatcttgtggaaaggacgaaacaccGGAAGTCCAGCGCTTCGCCCgtttaagagctatgctggaaacagc |
| Nf2 | Nf2-1 | GTGAACTACTTTACAATCC | tatcttgtggaaaggacgaaacaccGGTGAACTACTTTACAATCCgtttaagagctatgctggaaacagc |
| Nf2 | Nf2-2 | CTTGGTATGCGGAGCACCG | tatcttgtggaaaggacgaaacaccGCTTGGTATGCGGAGCACCGgtttaagagctatgctggaaacagc |
| Nf2 | Nf2-4 | GAGCACGGACGCCTCGGGA | tatcttgtggaaaggacgaaacaccGGAGCACGGACGCCTCGGGAgtttaagagctatgctggaaacagc |
| Nle1 | Nle1-1 | AGACGGCAGTGTACGAGTC | tatcttgtggaaaggacgaaacaccGAGACGGCAGTGTACGAGTCgtttaagagctatgctggaaacagc |
| Nle1 | Nle1-2 | CTGGCCACGTAGCGGCACT | tatcttgtggaaaggacgaaacaccGCTGGCCACGTAGCGGCACTgtttaagagctatgctggaaacagc |
| Nle1 | Nle1-3 | TTTCTTAGGACACCGGCAC | tatcttgtggaaaggacgaaacaccGTTTCTTAGGACACCGGCACgtttaagagctatgctggaaacagc |
| Nol10 | Nol10-1 | TGTCATAACACCGGACCCG | tatcttgtggaaaggacgaaacaccGTGTCATAACACCGGACCCGgtttaagagctatgctggaaacagc |
| Nol10 | Nol10-2 | GAGAAGTCTCGGCCGAACT | tatcttgtggaaaggacgaaacaccGGAGAAGTCTCGGCCGAACTgtttaagagctatgctggaaacagc |
| Nol10 | Nol10-5 | AGCGTGAGCCTACCTGAGC | tatcttgtggaaaggacgaaacaccGAGCGTGAGCCTACCTGAGCgtttaagagctatgctggaaacagc |
| Nom1 | Nom1-2 | AGAAGTGCCCGCTTCCGGG | tatcttgtggaaaggacgaaacaccGAGAAGTGCCCGCTTCCGGGgtttaagagctatgctggaaacagc |
| Nom1 | Nom1-4 | CCTCGGCTGAGGAACGAGG | tatcttgtggaaaggacgaaacaccGCCTCGGCTGAGGAACGAGGgtttaagagctatgctggaaacagc |
| Nom1 | Nom1-5 | TACCTCTATTCCGACCGTG | tatcttgtggaaaggacgaaacaccGTACCTCTATTCCGACCGTGgtttaagagctatgctggaaacagc |
| Nop16 | Nop16-2 | GGAAGCCCTATGTGGTAAA | tatcttgtggaaaggacgaaacaccGGGAAGCCCTATGTGGTAAAgtttaagagctatgctggaaacagc |
| Nop16 | Nop16-4 | TCTTTACAGGTAAAGGCCA | tatcttgtggaaaggacgaaacaccGTCTTTACAGGTAAAGGCCAgtttaagagctatgctggaaacagc |
| Nop16 | Nop16-5 | TTCCCACATACGACATGCC | tatcttgtggaaaggacgaaacaccGTTCCCACATACGACATGCCgtttaagagctatgctggaaacagc |
| Nqo1 | Nqo1-1 | ATGGAGCCGCTACCCCCAG | tatcttgtggaaaggacgaaacaccGATGGAGCCGCTACCCCCAGgtttaagagctatgctggaaacagc |
| Nqo1 | Nqo1-2 | AAAGGACCGTTGTCGTACA | tatcttgtggaaaggacgaaacaccGAAAGGACCGTTGTCGTACAgtttaagagctatgctggaaacagc |
| Nqo1 | Nqo1-4 | TGATACCTGAAATATCACC | tatcttgtggaaaggacgaaacaccGTGATACCTGAAATATCACCgtttaagagctatgctggaaacagc |
| Nrf1 | Nrf1-2 | AGACACGTTTGCTTCGGTG | tatcttgtggaaaggacgaaacaccGAGACACGTTTGCTTCGGTGgtttaagagctatgctggaaacagc |
| Nrf1 | Nrf1-3 | ATGAGTACACGACGCGAGT | tatcttgtggaaaggacgaaacaccGATGAGTACACGACGCGAGTgtttaagagctatgctggaaacagc |
| Nrf1 | Nrf1-5 | TCAAGTATTCCACAGGTCG | tatcttgtggaaaggacgaaacaccGTCAAGTATTCCACAGGTCGgtttaagagctatgctggaaacagc |
| Nsf | Nsf-1 | TTCAGACATACCGATAACG | tatcttgtggaaaggacgaaacaccGTTCAGACATACCGATAACGgtttaagagctatgctggaaacagc |
| Nsf | Nsf-2 | ACCAGGGGGTCCGTATAAC | tatcttgtggaaaggacgaaacaccGACCAGGGGGTCCGTATAACgtttaagagctatgctggaaacagc |
| Nsf | Nsf-3 | AAAATGGGGATCGGCGGTC | tatcttgtggaaaggacgaaacaccGAAAATGGGGATCGGCGGTCgtttaagagctatgctggaaacagc |
| Nsun6 | Nsun6-10 | AAGCAGACAAGAAGCAGAA | tatcttgtggaaaggacgaaacaccGAAGCAGACAAGAAGCAGAAgtttaagagctatgctggaaacagc |
| Nsun6 | Nsun6-3 | CTAAAGTATTTCTCGGAAA | tatcttgtggaaaggacgaaacaccGCTAAAGTATTTCTCGGAAAgtttaagagctatgctggaaacagc |
| Nsun6 | Nsun6-4 | CCGTTAATATCAGAATACA | tatcttgtggaaaggacgaaacaccGCCGTTAATATCAGAATACAgtttaagagctatgctggaaacagc |
| Nubp2 | Nubp2-1 | GTCTATTGGGGATGTGAGG | tatcttgtggaaaggacgaaacaccGGTCTATTGGGGATGTGAGGgtttaagagctatgctggaaacagc |
| Nubp2 | Nubp2-3 | AAGCAGTTTGTGTCTGACG | tatcttgtggaaaggacgaaacaccGAAGCAGTTTGTGTCTGACGgtttaagagctatgctggaaacagc |
| Nubp2 | Nubp2-5 | CAGGTGGGAATCCTAGATG | tatcttgtggaaaggacgaaacaccGCAGGTGGGAATCCTAGATGgtttaagagctatgctggaaacagc |
| Nudt21 | Nudt21-1 | GATGAGAAGGACTGTAGAA | tatcttgtggaaaggacgaaacaccGGATGAGAAGGACTGTAGAAgtttaagagctatgctggaaacagc |
| Nudt21 | Nudt21-3 | TCCCTCATGCGCTGAAATC | tatcttgtggaaaggacgaaacaccGTCCCTCATGCGCTGAAATCgtttaagagctatgctggaaacagc |
| Nudt21 | Nudt21-4 | AGAGCTGTCCTTCTCATAG | tatcttgtggaaaggacgaaacaccGAGAGCTGTCCTTCTCATAGgtttaagagctatgctggaaacagc |
| Nuf2 | Nuf2-1 | TTTAGTACCGTTCCGGCTG | tatcttgtggaaaggacgaaacaccGTTTAGTACCGTTCCGGCTGgtttaagagctatgctggaaacagc |
| Nuf2 | Nuf2-2 | TGCTGCATTCTGACCATAG | tatcttgtggaaaggacgaaacaccGTGCTGCATTCTGACCATAGgtttaagagctatgctggaaacagc |
| Nuf2 | Nuf2-3 | AAGTCATTCACCCGGCAAA | tatcttgtggaaaggacgaaacaccGAAGTCATTCACCCGGCAAAgtttaagagctatgctggaaacagc |
| Nufip2 | Nufip2-2 | GTTCAGGCTATATTACTAA | tatcttgtggaaaggacgaaacaccGGTTCAGGCTATATTACTAAgtttaagagctatgctggaaacagc |
| Nufip2 | Nufip2-4 | GAAGCACGTCACCCATAGC | tatcttgtggaaaggacgaaacaccGGAAGCACGTCACCCATAGCgtttaagagctatgctggaaacagc |
| Nufip2 | Nufip2-5 | AATTTGATGATCGGCCTAA | tatcttgtggaaaggacgaaacaccGAATTTGATGATCGGCCTAAgtttaagagctatgctggaaacagc |
| Numa1 | Numa1-2 | GCTAATCTTTCGATCCATC | tatcttgtggaaaggacgaaacaccGGCTAATCTTTCGATCCATCgtttaagagctatgctggaaacagc |
| Numa1 | Numa1-3 | GCATTCAACGACCTGATAG | tatcttgtggaaaggacgaaacaccGGCATTCAACGACCTGATAGgtttaagagctatgctggaaacagc |
| Numa1 | Numa1-4 | TAGGAGACGCCTTGCAGGT | tatcttgtggaaaggacgaaacaccGTAGGAGACGCCTTGCAGGTgtttaagagctatgctggaaacagc |
| Nup85 | Nup85-1 | TTCCTTCAACCAAACAGGT | tatcttgtggaaaggacgaaacaccGTTCCTTCAACCAAACAGGTgtttaagagctatgctggaaacagc |
| Nup85 | Nup85-3 | CAGTCGGACCCAGTCAAGA | tatcttgtggaaaggacgaaacaccGCAGTCGGACCCAGTCAAGAgtttaagagctatgctggaaacagc |
| Nup85 | Nup85-4 | CTTCAGGGCCGGCTCGATG | tatcttgtggaaaggacgaaacaccGCTTCAGGGCCGGCTCGATGgtttaagagctatgctggaaacagc |
| Oaz2 | Oaz2-1 | ACCGGGGATCTTCGACAGT | tatcttgtggaaaggacgaaacaccGACCGGGGATCTTCGACAGTgtttaagagctatgctggaaacagc |
| Oaz2 | Oaz2-2 | AGCTCTAATATATAAGGTA | tatcttgtggaaaggacgaaacaccGAGCTCTAATATATAAGGTAgtttaagagctatgctggaaacagc |
| Oaz2 | Oaz2-3 | TTTTCCATCGTTCACAGGG | tatcttgtggaaaggacgaaacaccGTTTTCCATCGTTCACAGGGgtttaagagctatgctggaaacagc |
| Obsl1 | Obsl1-1 | GAACCGTTGAACACGACAT | tatcttgtggaaaggacgaaacaccGGAACCGTTGAACACGACATgtttaagagctatgctggaaacagc |
| Obsl1 | Obsl1-3 | GTCGCGGTCGCGGTACATG | tatcttgtggaaaggacgaaacaccGGTCGCGGTCGCGGTACATGgtttaagagctatgctggaaacagc |
| Obsl1 | Obsl1-4 | ACACGTACACCCCGGAATC | tatcttgtggaaaggacgaaacaccGACACGTACACCCCGGAATCgtttaagagctatgctggaaacagc |
| Ogn | Ogn-1 | TTAACTATGAGTATGCAAC | tatcttgtggaaaggacgaaacaccGTTAACTATGAGTATGCAACgtttaagagctatgctggaaacagc |
| Ogn | Ogn-2 | GAAGAAACCAAATTTAGCC | tatcttgtggaaaggacgaaacaccGGAAGAAACCAAATTTAGCCgtttaagagctatgctggaaacagc |
| Ogn | Ogn-5 | CGCATGTCTGCAAAATCCT | tatcttgtggaaaggacgaaacaccGCGCATGTCTGCAAAATCCTgtttaagagctatgctggaaacagc |
| Omd | Omd-1 | GATTTCATTATAACCAAGA | tatcttgtggaaaggacgaaacaccGGATTTCATTATAACCAAGAgtttaagagctatgctggaaacagc |
| Omd | Omd-3 | ACATTCACCAGCCCGTCCA | tatcttgtggaaaggacgaaacaccGACATTCACCAGCCCGTCCAgtttaagagctatgctggaaacagc |
| Omd | Omd-5 | ACGTTGAGTTCTATAAGAT | tatcttgtggaaaggacgaaacaccGACGTTGAGTTCTATAAGATgtttaagagctatgctggaaacagc |
| Oosp3 | Oosp3-2 | CCCTAATAATAATTCATCA | tatcttgtggaaaggacgaaacaccGCCCTAATAATAATTCATCAgtttaagagctatgctggaaacagc |
| Oosp3 | Oosp3-3 | CTGGCCTTATGGAAGTTAC | tatcttgtggaaaggacgaaacaccGCTGGCCTTATGGAAGTTACgtttaagagctatgctggaaacagc |
| Oosp3 | Oosp3-4 | ACCTGTATGAAAGTCCCAC | tatcttgtggaaaggacgaaacaccGACCTGTATGAAAGTCCCACgtttaagagctatgctggaaacagc |
| Oxnad1 | Oxnad1-1 | AGAGACACAAACCTCCCGG | tatcttgtggaaaggacgaaacaccGAGAGACACAAACCTCCCGGgtttaagagctatgctggaaacagc |
| Oxnad1 | Oxnad1-3 | GACAGAATAATAGAACTGG | tatcttgtggaaaggacgaaacaccGGACAGAATAATAGAACTGGgtttaagagctatgctggaaacagc |
| Oxnad1 | Oxnad1-5 | ATCAAGATGAGGTTTCGAG | tatcttgtggaaaggacgaaacaccGATCAAGATGAGGTTTCGAGgtttaagagctatgctggaaacagc |
| Paf1 | Paf1-1 | GATGAACTTGGGGTCAAAT | tatcttgtggaaaggacgaaacaccGGATGAACTTGGGGTCAAATgtttaagagctatgctggaaacagc |
| Paf1 | Paf1-3 | CAATGGTGACCCCCAGATC | tatcttgtggaaaggacgaaacaccGCAATGGTGACCCCCAGATCgtttaagagctatgctggaaacagc |
| Paf1 | Paf1-4 | GCATGCAAAGGTGGTACCG | tatcttgtggaaaggacgaaacaccGGCATGCAAAGGTGGTACCGgtttaagagctatgctggaaacagc |
| Paip2b | Paip2b-1 | CCTGAGGCAGGTCTCGTGC | tatcttgtggaaaggacgaaacaccGCCTGAGGCAGGTCTCGTGCgtttaagagctatgctggaaacagc |
| Paip2b | Paip2b-3 | GAAGAGGATTTCAACAGAC | tatcttgtggaaaggacgaaacaccGGAAGAGGATTTCAACAGACgtttaagagctatgctggaaacagc |
| Paip2b | Paip2b-4 | CCACATGTATTCTGCAAAT | tatcttgtggaaaggacgaaacaccGCCACATGTATTCTGCAAATgtttaagagctatgctggaaacagc |
| Pak1ip1 | Pak1ip1-1 | GGGCTCTGGTGCATCACGC | tatcttgtggaaaggacgaaacaccGGGGCTCTGGTGCATCACGCgtttaagagctatgctggaaacagc |
| Pak1ip1 | Pak1ip1-3 | ATCCACCCGTCTGGCAAGT | tatcttgtggaaaggacgaaacaccGATCCACCCGTCTGGCAAGTgtttaagagctatgctggaaacagc |
| Pak1ip1 | Pak1ip1-5 | TAGGACTCTGTCCTTGCAG | tatcttgtggaaaggacgaaacaccGTAGGACTCTGTCCTTGCAGgtttaagagctatgctggaaacagc |
| Pals2 | Pals2-3 | TGAATACCGAGAATACGGA | tatcttgtggaaaggacgaaacaccGTGAATACCGAGAATACGGAgtttaagagctatgctggaaacagc |
| Pals2 | Pals2-4 | GAGGAATGATAGATCGACA | tatcttgtggaaaggacgaaacaccGGAGGAATGATAGATCGACAgtttaagagctatgctggaaacagc |

| | | | |
|---|---|---|---|
| Pals2 | Pals2-5 | CAGGACTAAAATTTTCCAA | tatcttgtggaaaggacgaaacaccGCAGGACTAAAATTTTCCAAgtttaagagctatgctggaaacagc |
| Pam16 | Pam16-2 | AGCCCCGAGGAGGTCCAGA | tatcttgtggaaaggacgaaacaccGAGCCCCGAGGAGGTCCAGAgtttaagagctatgctggaaacagc |
| Pam16 | Pam16-3 | GGAGACGTTGAGAATCTGC | tatcttgtggaaaggacgaaacaccGGGAGACGTTGAGAATCTGCgtttaagagctatgctggaaacagc |
| Pam16 | Pam16-4 | CAGCTGCATCCAATCTCTC | tatcttgtggaaaggacgaaacaccGCAGCTGCATCCAATCTCTCgtttaagagctatgctggaaacagc |
| Parp4 | Parp4-1 | GTAATAGACTTCGTAGCCC | tatcttgtggaaaggacgaaacaccGGTAATAGACTTCGTAGCCCgtttaagagctatgctggaaacagc |
| Parp4 | Parp4-2 | GCTTCGAGGCATTCATCAA | tatcttgtggaaaggacgaaacaccGGCTTCGAGGCATTCATCAAgtttaagagctatgctggaaacagc |
| Parp4 | Parp4-3 | ATGAGGACTTTAGCCCGAG | tatcttgtggaaaggacgaaacaccGATGAGGACTTTAGCCCGAGgtttaagagctatgctggaaacagc |
| Pate6 | Pate6-2 | GGATCATACACGTTTCACC | tatcttgtggaaaggacgaaacaccGGGATCATACACGTTTCACCgtttaagagctatgctggaaacagc |
| Pate6 | Pate6-3 | AGACATGCACTACAAAACC | tatcttgtggaaaggacgaaacaccGAGACATGCACTACAAAACCgtttaagagctatgctggaaacagc |
| Pate6 | Pate6-5 | GTGTCAGTCTTACAAAAAT | tatcttgtggaaaggacgaaacaccGGTGTCAGTCTTACAAAAATgtttaagagctatgctggaaacagc |
| Pcdhb22 | Pcdhb22-1 | GCGTTCAGAACTATAAGAT | tatcttgtggaaaggacgaaacaccGGCGTTCAGAACTATAAGATgtttaagagctatgctggaaacagc |
| Pcdhb22 | Pcdhb22-2 | CCATCCGCAACCGAGGTGA | tatcttgtggaaaggacgaaacaccGCCATCCGCAACCGAGGTGAgtttaagagctatgctggaaacagc |
| Pcdhb22 | Pcdhb22-5 | GGATTCGAGACCGAGACTC | tatcttgtggaaaggacgaaacaccGGGATTCGAGACCGAGACTCgtttaagagctatgctggaaacagc |
| Pcgf3 | Pcgf3-2 | CCCGATATACTGTAATGGG | tatcttgtggaaaggacgaaacaccGCCCGATATACTGTAATGGGgtttaagagctatgctggaaacagc |
| Pcgf3 | Pcgf3-3 | ATTCTTAATAGCGGAAATG | tatcttgtggaaaggacgaaacaccGATTCTTAATAGCGGAAATGgtttaagagctatgctggaaacagc |
| Pcgf3 | Pcgf3-4 | AGGGAATTCTATCACAAAC | tatcttgtggaaaggacgaaacaccGAGGGAATTCTATCACAAACgtttaagagctatgctggaaacagc |
| Pcid2 | Pcid2-2 | ATTATTGGCAAATATCCGA | tatcttgtggaaaggacgaaacaccGATTATTGGCAAATATCCGAgtttaagagctatgctggaaacagc |
| Pcid2 | Pcid2-3 | GTGCTACTGCATACATGAC | tatcttgtggaaaggacgaaacaccGGTGCTACTGCATACATGACgtttaagagctatgctggaaacagc |
| Pcid2 | Pcid2-4 | ATATTGGACTATCACCGTC | tatcttgtggaaaggacgaaacaccGATATTGGACTATCACCGTCgtttaagagctatgctggaaacagc |
| Pdap1 | Pdap1-1 | ACTAAGAAGGTCACGCAAC | tatcttgtggaaaggacgaaacaccGACTAAGAAGGTCACGCAACgtttaagagctatgctggaaacagc |
| Pdap1 | Pdap1-4 | AAAAGCGAAAAGGCGTGGA | tatcttgtggaaaggacgaaacaccGAAAAGCGAAAAGGCGTGGAgtttaagagctatgctggaaacagc |
| Pdap1 | Pdap1-5 | GCTTCAGGTGACCCAAAAA | tatcttgtggaaaggacgaaacaccGGCTTCAGGTGACCCAAAAAgtttaagagctatgctggaaacagc |
| Pdcd2l | Pdcd2l-2 | GAACGGGGAGCCGTCCAGC | tatcttgtggaaaggacgaaacaccGGAACGGGGAGCCGTCCAGCgtttaagagctatgctggaaacagc |
| Pdcd2l | Pdcd2l-3 | CCTGCACGACGAGCGTGAG | tatcttgtggaaaggacgaaacaccGCCTGCACGACGAGCGTGAGgtttaagagctatgctggaaacagc |
| Pdcd2l | Pdcd2l-4 | AGCGGCCACACTGCGGCCC | tatcttgtggaaaggacgaaacaccGAGCGGCCACACTGCGGCCCgtttaagagctatgctggaaacagc |
| Pdcl2 | Pdcl2-2 | GGTTAACCACCAAGCACAT | tatcttgtggaaaggacgaaacaccGGGTTAACCACCAAGCACATgtttaagagctatgctggaaacagc |
| Pdcl2 | Pdcl2-3 | AATTCATCTCTATAGATCA | tatcttgtggaaaggacgaaacaccGAATTCATCTCTATAGATCAgtttaagagctatgctggaaacagc |
| Pdcl2 | Pdcl2-4 | AGAAAAGCGGTTACAGGAA | tatcttgtggaaaggacgaaacaccGAGAAAAGCGGTTACAGGAAgtttaagagctatgctggaaacagc |
| Pelo | Pelo-3 | GCTTACCACACCATCGAAC | tatcttgtggaaaggacgaaacaccGGCTTACCACACCATCGAACgtttaagagctatgctggaaacagc |
| Pelo | Pelo-4 | TCGATGGCCTCCACGCAAA | tatcttgtggaaaggacgaaacaccGTCGATGGCCTCCACGCAAAgtttaagagctatgctggaaacagc |
| Pelo | Pelo-5 | TCGGTCTGGACCTTGCGGA | tatcttgtggaaaggacgaaacaccGTCGGTCTGGACCTTGCGGAgtttaagagctatgctggaaacagc |
| Pex12 | Pex12-1 | GCCTACCCTTTTGTTAACA | tatcttgtggaaaggacgaaacaccGGCCTACCCTTTTGTTAACAgtttaagagctatgctggaaacagc |
| Pex12 | Pex12-2 | ATCGTTTCCAGCGAGAAGA | tatcttgtggaaaggacgaaacaccGATCGTTTCCAGCGAGAAGAgtttaagagctatgctggaaacagc |
| Pex12 | Pex12-5 | AATCAAATCCTGCGCACTA | tatcttgtggaaaggacgaaacaccGAATCAAATCCTGCGCACTAgtttaagagctatgctggaaacagc |
| Pfdn2 | Pfdn2-2 | GAATTGAATGAGCACAGGT | tatcttgtggaaaggacgaaacaccGGAATTGAATGAGCACAGGTgtttaagagctatgctggaaacagc |
| Pfdn2 | Pfdn2-3 | TGTATCGATCACCAGGCTA | tatcttgtggaaaggacgaaacaccGTGTATCGATCACCAGGCTAgtttaagagctatgctggaaacagc |
| Pfdn2 | Pfdn2-4 | ATCGATACACTGAAGGAAG | tatcttgtggaaaggacgaaacaccGATCGATACACTGAAGGAAGgtttaagagctatgctggaaacagc |
| Pfdn5 | Pfdn5-2 | AACAAGAGCAACGAGGGTA | tatcttgtggaaaggacgaaacaccGAACAAGAGCAACGAGGGTAgtttaagagctatgctggaaacagc |
| Pfdn5 | Pfdn5-4 | CCTCTCAGATGTACGTCCC | tatcttgtggaaaggacgaaacaccGCCTCTCAGATGTACGTCCCgtttaagagctatgctggaaacagc |
| Pfdn5 | Pfdn5-5 | ACCGGCTACTACGTGGAGA | tatcttgtggaaaggacgaaacaccGACCGGCTACTACGTGGAGAgtttaagagctatgctggaaacagc |
| Phb | Phb-2 | CAGTCATCACTGGTAGCAA | tatcttgtggaaaggacgaaacaccGCAGTCATCACTGGTAGCAAgtttaagagctatgctggaaacagc |
| Phb | Phb-3 | CGAATCCTCTTCCGGCCGG | tatcttgtggaaaggacgaaacaccGCGAATCCTCTTCCGGCCGGgtttaagagctatgctggaaacagc |
| Phb | Phb-5 | GGATGACGTGTCCCTGGTA | tatcttgtggaaaggacgaaacaccGGGATGACGTGTCCCTGGTAgtttaagagctatgctggaaacagc |
| Phb2 | Phb2-2 | CTATGACCGCCTTCCACTG | tatcttgtggaaaggacgaaacaccGCTATGACCGCCTTCCACTGgtttaagagctatgctggaaacagc |
| Phb2 | Phb2-4 | GCCCGAATGTCATAGATGA | tatcttgtggaaaggacgaaacaccGGCCCGAATGTCATAGATGAgtttaagagctatgctggaaacagc |
| Phb2 | Phb2-5 | ATAGTCCAGCCCTAGACGC | tatcttgtggaaaggacgaaacaccGATAGTCCAGCCCTAGACGCgtttaagagctatgctggaaacagc |
| Pigw | Pigw-1 | GCGGCAATGGTTTGTCCAG | tatcttgtggaaaggacgaaacaccGGCGGCAATGGTTTGTCCAGgtttaagagctatgctggaaacagc |
| Pigw | Pigw-3 | CTGTTAATGACCCGGTAAC | tatcttgtggaaaggacgaaacaccGCTGTTAATGACCCGGTAACgtttaagagctatgctggaaacagc |
| Pigw | Pigw-5 | CAGTCAGAGTGATCACTAG | tatcttgtggaaaggacgaaacaccGCAGTCAGAGTGATCACTAGgtttaagagctatgctggaaacagc |
| Pknox1 | Pknox1-1 | CCAATGGATGCCGACAAGC | tatcttgtggaaaggacgaaacaccGCCAATGGATGCCGACAAGCgtttaagagctatgctggaaacagc |
| Pknox1 | Pknox1-3 | AGCGAGACCTTGTTGAGTG | tatcttgtggaaaggacgaaacaccGAGCGAGACCTTGTTGAGTGgtttaagagctatgctggaaacagc |
| Pknox1 | Pknox1-5 | GCGGAGTGACGACGGTGAC | tatcttgtggaaaggacgaaacaccGGCGGAGTGACGACGGTGACgtttaagagctatgctggaaacagc |
| Pmp2 | Pmp2-3 | AAGGCACTTTCCGTTCTAA | tatcttgtggaaaggacgaaacaccGAAGGCACTTTCCGTTCTAAgtttaagagctatgctggaaacagc |
| Pmp2 | Pmp2-4 | GTGATCATCAGCAAAAAGG | tatcttgtggaaaggacgaaacaccGGTGATCATCAGCAAAAAGGgtttaagagctatgctggaaacagc |
| Pmp2 | Pmp2-5 | AACAGAAAACTGGGAAACT | tatcttgtggaaaggacgaaacaccGAACAGAAAACTGGGAAACTgtttaagagctatgctggaaacagc |
| Pnkp | Pnkp-1 | CGGTCCGTCACTTGGGTCA | tatcttgtggaaaggacgaaacaccGCGGTCCGTCACTTGGGTCAgtttaagagctatgctggaaacagc |
| Pnkp | Pnkp-4 | CGTCTGGGGTGAAGCCCCA | tatcttgtggaaaggacgaaacaccGCGTCTGGGGTGAAGCCCCAgtttaagagctatgctggaaacagc |
| Pnkp | Pnkp-5 | TTCTGGCCGTACCTGAAAT | tatcttgtggaaaggacgaaacaccGTTCTGGCCGTACCTGAAATgtttaagagctatgctggaaacagc |
| Pnma3 | Pnma3-1 | AAGATACAGGATCATCGGC | tatcttgtggaaaggacgaaacaccGAAGATACAGGATCATCGGCgtttaagagctatgctggaaacagc |
| Pnma3 | Pnma3-2 | AATTCATCATCTGAATGAG | tatcttgtggaaaggacgaaacaccGAATTCATCATCTGAATGAGgtttaagagctatgctggaaacagc |
| Pnma3 | Pnma3-4 | ATCGAAGGCCAATAACCCT | tatcttgtggaaaggacgaaacaccGATCGAAGGCCAATAACCCTgtttaagagctatgctggaaacagc |
| Pnn | Pnn-1 | GTAGCTTATTGCTAAGGTA | tatcttgtggaaaggacgaaacaccGGTAGCTTATTGCTAAGGTAgtttaagagctatgctggaaacagc |
| Pnn | Pnn-2 | GGCGTGGATTCTCAGATAG | tatcttgtggaaaggacgaaacaccGGGCGTGGATTCTCAGATAGgtttaagagctatgctggaaacagc |
| Pnn | Pnn-5 | GTGCGTAATGAAGAACAGA | tatcttgtggaaaggacgaaacaccGGTGCGTAATGAAGAACAGAgtttaagagctatgctggaaacagc |
| Pof1b | Pof1b-1 | TAGAACATTTGGAGCGAAT | tatcttgtggaaaggacgaaacaccGTAGAACATTTGGAGCGAATgtttaagagctatgctggaaacagc |
| Pof1b | Pof1b-3 | ATAGTACAAAACCCTGAAC | tatcttgtggaaaggacgaaacaccGATAGTACAAAACCCTGAACgtttaagagctatgctggaaacagc |
| Pof1b | Pof1b-4 | CAAACCAAACTTTGGTAGG | tatcttgtggaaaggacgaaacaccGCAAACCAAACTTTGGTAGGgtttaagagctatgctggaaacagc |
| Pold1 | Pold1-1 | GGCGCTCTAGTAGTCGGAG | tatcttgtggaaaggacgaaacaccGGGCGCTCTAGTAGTCGGAGgtttaagagctatgctggaaacagc |
| Pold1 | Pold1-4 | TCCGAACGTACTTTCCAGC | tatcttgtggaaaggacgaaacaccGTCCGAACGTACTTTCCAGCgtttaagagctatgctggaaacagc |
| Pold1 | Pold1-5 | GCGGCGGGCTGGTGCCATA | tatcttgtggaaaggacgaaacaccGGCGGCGGGCTGGTGCCATAgtttaagagctatgctggaaacagc |
| Pole3 | Pole3-2 | CATTGAAAGAAGCTCTAGA | tatcttgtggaaaggacgaaacaccGCATTGAAAGAAGCTCTAGAgtttaagagctatgctggaaacagc |
| Pole3 | Pole3-3 | CCAATAACTTCGCAATGAA | tatcttgtggaaaggacgaaacaccGCCAATAACTTCGCAATGAAgtttaagagctatgctggaaacagc |
| Pole3 | Pole3-4 | CGTTTTGTATGCCACATCC | tatcttgtggaaaggacgaaacaccGCGTTTTGTATGCCACATCCgtttaagagctatgctggaaacagc |
| Poll | Poll-1 | CTAAAGCCTACAGTGTCCA | tatcttgtggaaaggacgaaacaccGCTAAAGCCTACAGTGTCCAgtttaagagctatgctggaaacagc |
| Poll | Poll-4 | GGTCCTGGGTAAATCCCGC | tatcttgtggaaaggacgaaacaccGGGTCCTGGGTAAATCCCGCgtttaagagctatgctggaaacagc |
| Poll | Poll-5 | GAGCCCACATTATGCCCGC | tatcttgtggaaaggacgaaacaccGGAGCCCACATTATGCCCGCgtttaagagctatgctggaaacagc |

| | | | |
|---|---|---|---|
| Polm | Polm-1 | TTGAGCACCGAGGCCGCTC | tatcttgtggaaaggacgaaacaccGTTGAGCACCGAGGCCGCTCgtttaagagctatgctggaaacagc |
| Polm | Polm-3 | CTGGCATCGACGCTGAGAC | tatcttgtggaaaggacgaaacaccGCTGGCATCGACGCTGAGACgtttaagagctatgctggaaacagc |
| Polm | Polm-5 | ACCTTGCGGAGCCGCGAAT | tatcttgtggaaaggacgaaacaccGACCTTGCGGAGCCGCGAATgtttaagagctatgctggaaacagc |
| Polq | Polq-2 | TACGAGCTATTCTGGAGGT | tatcttgtggaaaggacgaaacaccGTACGAGCTATTCTGGAGGTgtttaagagctatgctggaaacagc |
| Polq | Polq-3 | CGGTGTTTTCCAACCGCCT | tatcttgtggaaaggacgaaacaccGCGGTGTTTTCCAACCGCCTgtttaagagctatgctggaaacagc |
| Polq | Polq-5 | GCTCTCTGCGCGTTGAGTA | tatcttgtggaaaggacgaaacaccGGCTCTCTGCGCGTTGAGTAgtttaagagctatgctggaaacagc |
| Polr1a | Polr1a-2 | GGACGGTCGGTGGTTCGAA | tatcttgtggaaaggacgaaacaccGGGACGGTCGGTGGTTCGAAgtttaagagctatgctggaaacagc |
| Polr1a | Polr1a-3 | CATAGTCGTACCGCCATCC | tatcttgtggaaaggacgaaacaccGCATAGTCGTACCGCCATCCgtttaagagctatgctggaaacagc |
| Polr1a | Polr1a-5 | GTTGACTACGCAGCCCGCT | tatcttgtggaaaggacgaaacaccGGTTGACTACGCAGCCCGCTgtttaagagctatgctggaaacagc |
| Polr1b | Polr1b-1 | GACCTGACGCGGGCGCACG | tatcttgtggaaaggacgaaacaccGGACCTGACGCGGGCGCACGgtttaagagctatgctggaaacagc |
| Polr1b | Polr1b-2 | TTCAACTACGCGGCGCTGG | tatcttgtggaaaggacgaaacaccGTTCAACTACGCGGCGCTGGgtttaagagctatgctggaaacagc |
| Polr1b | Polr1b-3 | CCGAAAGAGCACCTACCGA | tatcttgtggaaaggacgaaacaccGCCGAAAGAGCACCTACCGAgtttaagagctatgctggaaacagc |
| Polr1d | Polr1d-1 | TGTATGGTCCTCCTCGTGC | tatcttgtggaaaggacgaaacaccGTGTATGGTCCTCCTCGTGCgtttaagagctatgctggaaacagc |
| Polr1d | Polr1d-2 | TACACTAGGAAACTGTCTG | tatcttgtggaaaggacgaaacaccGTACACTAGGAAACTGTCTGgtttaagagctatgctggaaacagc |
| Polr1d | Polr1d-4 | ATTTGCGAATTCAGACCCG | tatcttgtggaaaggacgaaacaccGATTTGCGAATTCAGACCCGgtttaagagctatgctggaaacagc |
| Polr2i | Polr2i-1 | AATCGCATTCTGCTGTACG | tatcttgtggaaaggacgaaacaccGAATCGCATTCTGCTGTACGgtttaagagctatgctggaaacagc |
| Polr2i | Polr2i-2 | CGGAATTGCGATTACCAGC | tatcttgtggaaaggacgaaacaccGCGGAATTGCGATTACCAGCgtttaagagctatgctggaaacagc |
| Polr2i | Polr2i-5 | CCGGGGCAACGTGGGGTCC | tatcttgtggaaaggacgaaacaccGCCGGGGCAACGTGGGGTCCgtttaagagctatgctggaaacagc |
| Polr2j | Polr2j-1 | ATGGTGAACAAGCAGGCGT | tatcttgtggaaaggacgaaacaccGATGGTGAACAAGCAGGCGTgtttaagagctatgctggaaacagc |
| Polr2j | Polr2j-4 | CTTGTGCTCCAAGGGGTGA | tatcttgtggaaaggacgaaacaccGCTTGTGCTCCAAGGGGTGAgtttaagagctatgctggaaacagc |
| Polr2j | Polr2j-5 | TGGGGACTGTAGTCTGGGG | tatcttgtggaaaggacgaaacaccGTGGGGACTGTAGTCTGGGGgtttaagagctatgctggaaacagc |
| Polr3b | Polr3b-1 | TCCGCAACGCCTTACCCAT | tatcttgtggaaaggacgaaacaccGTCCGCAACGCCTTACCCATgtttaagagctatgctggaaacagc |
| Polr3b | Polr3b-2 | GCTCGAACTGTGTTCTAAC | tatcttgtggaaaggacgaaacaccGGCTCGAACTGTGTTCTAACgtttaagagctatgctggaaacagc |
| Polr3b | Polr3b-3 | GATCATTGTGGAGGCCGAC | tatcttgtggaaaggacgaaacaccGGATCATTGTGGAGGCCGACgtttaagagctatgctggaaacagc |
| Polr3d | Polr3d-1 | GCAGTTCATCGATGACCCG | tatcttgtggaaaggacgaaacaccGGCAGTTCATCGATGACCCGgtttaagagctatgctggaaacagc |
| Polr3d | Polr3d-2 | CCCATTCTATCTTCGAGCA | tatcttgtggaaaggacgaaacaccGCCCATTCTATCTTCGAGCAgtttaagagctatgctggaaacagc |
| Polr3d | Polr3d-5 | AGGCGACCGGGCGTAAGCG | tatcttgtggaaaggacgaaacaccGAGGCGACCGGGCGTAAGCGgtttaagagctatgctggaaacagc |
| Pop5 | Pop5-2 | ACGATCGCCCGGGTGCACG | tatcttgtggaaaggacgaaacaccGACGATCGCCCGGGTGCACGgtttaagagctatgctggaaacagc |
| Pop5 | Pop5-4 | AAGAGCTGACCACACGAGC | tatcttgtggaaaggacgaaacaccGAAGAGCTGACCACACGAGCgtttaagagctatgctggaaacagc |
| Pop5 | Pop5-5 | TTCTTCAACACATTACACG | tatcttgtggaaaggacgaaacaccGTTCTTCAACACATTACACGgtttaagagctatgctggaaacagc |
| Pou5f1 | Pou5f1-1 | TCGTATGCGGGCGGACATG | tatcttgtggaaaggacgaaacaccGTCGTATGCGGGCGGACATGgtttaagagctatgctggaaacagc |
| Pou5f1 | Pou5f1-2 | TGGGCCTAGTCCCCCAAGT | tatcttgtggaaaggacgaaacaccGTGGGCCTAGTCCCCCAAGTgtttaagagctatgctggaaacagc |
| Pou5f1 | Pou5f1-5 | CATTGAGAACCGTGTGAGG | tatcttgtggaaaggacgaaacaccGCATTGAGAACCGTGTGAGGgtttaagagctatgctggaaacagc |
| Ppan | Ppan-1 | ACTCGTTCGTGTTCACGCG | tatcttgtggaaaggacgaaacaccGACTCGTTCGTGTTCACGCGgtttaagagctatgctggaaacagc |
| Ppan | Ppan-2 | CCGGACATTGCGACCCGCT | tatcttgtggaaaggacgaaacaccGCCGGACATTGCGACCCGCTgtttaagagctatgctggaaacagc |
| Ppan | Ppan-5 | GCGTCAAAGTGGTTCCCGT | tatcttgtggaaaggacgaaacaccGGCGTCAAAGTGGTTCCCGTgtttaagagctatgctggaaacagc |
| Ppp1r2 | Ppp1r2-1 | TGACCTACCTCGCTCCTCA | tatcttgtggaaaggacgaaacaccGTGACCTACCTCGCTCCTCAgtttaagagctatgctggaaacagc |
| Ppp1r2 | Ppp1r2-3 | CATCGTCACCTATCATACT | tatcttgtggaaaggacgaaacaccGCATCGTCACCTATCATACTgtttaagagctatgctggaaacagc |
| Ppp1r2 | Ppp1r2-4 | CGTCACGTTACTTGTGATA | tatcttgtggaaaggacgaaacaccGCGTCACGTTACTTGTGATAgtttaagagctatgctggaaacagc |
| Ppp1r7 | Ppp1r7-3 | AACTGCAAATGTTGGAGCT | tatcttgtggaaaggacgaaacaccGAACTGCAAATGTTGGAGCTgtttaagagctatgctggaaacagc |
| Ppp1r7 | Ppp1r7-4 | GGTCTAACCGGATTCGGGT | tatcttgtggaaaggacgaaacaccGGGTCTAACCGGATTCGGGTgtttaagagctatgctggaaacagc |
| Ppp1r7 | Ppp1r7-5 | ATTACAAAACTTCAGAACC | tatcttgtggaaaggacgaaacaccGATTACAAAACTTCAGAACCgtttaagagctatgctggaaacagc |
| Ppp2ca | Ppp2ca-1 | CATCGAACCTCTTGAACGT | tatcttgtggaaaggacgaaacaccGCATCGAACCTCTTGAACGTgtttaagagctatgctggaaacagc |
| Ppp2ca | Ppp2ca-4 | CGAGCACTCGATCGCCTAC | tatcttgtggaaaggacgaaacaccGCGAGCACTCGATCGCCTACgtttaagagctatgctggaaacagc |
| Ppp2ca | Ppp2ca-5 | GCTGGGGGATATCTCCTCG | tatcttgtggaaaggacgaaacaccGGCTGGGGGATATCTCCTCGgtttaagagctatgctggaaacagc |
| Ppp4c | Ppp4c-2 | GCGACTCTCATGATTGCCC | tatcttgtggaaaggacgaaacaccGGCGACTCTCATGATTGCCCgtttaagagctatgctggaaacagc |
| Ppp4c | Ppp4c-3 | CTGTCAGGATAGCGAACCT | tatcttgtggaaaggacgaaacaccGCTGTCAGGATAGCGAACCTgtttaagagctatgctggaaacagc |
| Ppp4c | Ppp4c-4 | CCTCACCTTAAGAGCCAGC | tatcttgtggaaaggacgaaacaccGCCTCACCTTAAGAGCCAGCgtttaagagctatgctggaaacagc |
| Pramel26 | Pramel26-2 | AAGGTGGCCCTCCATGAAT | tatcttgtggaaaggacgaaacaccGAAGGTGGCCCTCCATGAATgtttaagagctatgctggaaacagc |
| Pramel26 | Pramel26-4 | GGCCACACAGGTATCATGG | tatcttgtggaaaggacgaaacaccGGGCCACACAGGTATCATGGgtttaagagctatgctggaaacagc |
| Pramel26 | Pramel26-5 | GCAATACAGCGTCTACTGA | tatcttgtggaaaggacgaaacaccGGCAATACAGCGTCTACTGAgtttaagagctatgctggaaacagc |
| Pramel32 | Pramel32-1 | TGAGAGAAAACAGTATCGG | tatcttgtggaaaggacgaaacaccGTGAGAGAAAACAGTATCGGgtttaagagctatgctggaaacagc |
| Pramel32 | Pramel32-3 | TCAGGTAGGGATTAAGAAA | tatcttgtggaaaggacgaaacaccGTCAGGTAGGGATTAAGAAAgtttaagagctatgctggaaacagc |
| Pramel32 | Pramel32-4 | AGTTTTGGAGGATATGGGC | tatcttgtggaaaggacgaaacaccGAGTTTTGGAGGATATGGGCgtttaagagctatgctggaaacagc |
| Prelid1 | Prelid1-2 | GACAGTAGCTTCTGGTCGG | tatcttgtggaaaggacgaaacaccGGACAGTAGCTTCTGGTCGGgtttaagagctatgctggaaacagc |
| Prelid1 | Prelid1-3 | ACAGTCGCTCTGCCCAACG | tatcttgtggaaaggacgaaacaccGACAGTCGCTCTGCCCAACGgtttaagagctatgctggaaacagc |
| Prelid1 | Prelid1-5 | ATCAACCATGCCCGGCTGA | tatcttgtggaaaggacgaaacaccGATCAACCATGCCCGGCTGAgtttaagagctatgctggaaacagc |
| Prkdc | Prkdc-3 | ATGAAGAAGTACGTGGCGT | tatcttgtggaaaggacgaaacaccGATGAAGAAGTACGTGGCGTgtttaagagctatgctggaaacagc |
| Prkdc | Prkdc-4 | ATGCGTCTTAGGTGATCGA | tatcttgtggaaaggacgaaacaccGATGCGTCTTAGGTGATCGAgtttaagagctatgctggaaacagc |
| Prkdc | Prkdc-5 | CGAGGAGTGCTACAATGGC | tatcttgtggaaaggacgaaacaccGCGAGGAGTGCTACAATGGCgtttaagagctatgctggaaacagc |
| Prl2c5 | Prl2c5-1 | ACATTATTCTAACGTGCCT | tatcttgtggaaaggacgaaacaccGACATTATTCTAACGTGCCTgtttaagagctatgctggaaacagc |
| Prl2c5 | Prl2c5-2 | AGAAGTATGGCATCTCATG | tatcttgtggaaaggacgaaacaccGAGAAGTATGGCATCTCATGgtttaagagctatgctggaaacagc |
| Prl2c5 | Prl2c5-3 | AAACTCACGCGTATGTGCC | tatcttgtggaaaggacgaaacaccGAAACTCACGCGTATGTGCCgtttaagagctatgctggaaacagc |
| Prl7a2 | Prl7a2-1 | ACCCAGAGTATACTCAACT | tatcttgtggaaaggacgaaacaccGACCCAGAGTATACTCAACTgtttaagagctatgctggaaacagc |
| Prl7a2 | Prl7a2-2 | GCTGATCCTCAGTAGAATG | tatcttgtggaaaggacgaaacaccGGCTGATCCTCAGTAGAATGgtttaagagctatgctggaaacagc |
| Prl7a2 | Prl7a2-5 | TTGCTACTCAAAGGTACAG | tatcttgtggaaaggacgaaacaccGTTGCTACTCAAAGGTACAGgtttaagagctatgctggaaacagc |
| Prl8a1 | Prl8a1-2 | TTCAGGTCCAATATTTGAA | tatcttgtggaaaggacgaaacaccGTTCAGGTCCAATATTTGAAgtttaagagctatgctggaaacagc |
| Prl8a1 | Prl8a1-3 | AGGGTTTGTGATAGTAGAG | tatcttgtggaaaggacgaaacaccGAGGGTTTGTGATAGTAGAGgtttaagagctatgctggaaacagc |
| Prl8a1 | Prl8a1-4 | AACTCACAACTGATTAGGC | tatcttgtggaaaggacgaaacaccGAACTCACAACTGATTAGGCgtttaagagctatgctggaaacagc |
| Prmt1 | Prmt1-3 | TGCATGTTTGCTGCCAAGG | tatcttgtggaaaggacgaaacaccGTGCATGTTTGCTGCCAAGGgtttaagagctatgctggaaacagc |
| Prmt1 | Prmt1-4 | AGATGCCGATTGTGAAACA | tatcttgtggaaaggacgaaacaccGAGATGCCGATTGTGAAACAgtttaagagctatgctggaaacagc |
| Prmt1 | Prmt1-5 | ACCTCGTGGATGCCAAAGT | tatcttgtggaaaggacgaaacaccGACCTCGTGGATGCCAAAGTgtttaagagctatgctggaaacagc |
| Prmt5 | Prmt5-1 | TAAGTATTCCAAGTACTGG | tatcttgtggaaaggacgaaacaccGTAAGTATTCCAAGTACTGGgtttaagagctatgctggaaacagc |
| Prmt5 | Prmt5-2 | TAATCACGTCATTGACCGC | tatcttgtggaaaggacgaaacaccGTAATCACGTCATTGACCGCgtttaagagctatgctggaaacagc |
| Prmt5 | Prmt5-4 | AACCTGCTAAGAATCGGCC | tatcttgtggaaaggacgaaacaccGAACCTGCTAAGAATCGGCCgtttaagagctatgctggaaacagc |
| Prpf19 | Prpf19-3 | ACCCCTGAGATTATCCAGA | tatcttgtggaaaggacgaaacaccGACCCCTGAGATTATCCAGAgtttaagagctatgctggaaacagc |

| | | | |
|---|---|---|---|
| Prpf19 | Prpf19-4 | CGGCAGGTGGCATCCCATG | tatcttgtggaaaggacgaaacaccGCGGCAGGTGGCATCCCATGgtttaagagctatgctggaaacagc |
| Prpf19 | Prpf19-5 | ATTCCTGGGATTCTCGCTC | tatcttgtggaaaggacgaaacaccGATTCCTGGGATTCTCGCTCgtttaagagctatgctggaaacagc |
| Prpf38a | Prpf38a-1 | CGAGTTAGTGCCCTAGAAG | tatcttgtggaaaggacgaaacaccGCGAGTTAGTGCCCTAGAAGgtttaagagctatgctggaaacagc |
| Prpf38a | Prpf38a-4 | GGTAGGTATGTCCGGATGC | tatcttgtggaaaggacgaaacaccGGGTAGGTATGTCCGGATGCgtttaagagctatgctggaaacagc |
| Prpf38a | Prpf38a-5 | TGATTCATAGATTCGCGTC | tatcttgtggaaaggacgaaacaccGTGATTCATAGATTCGCGTCgtttaagagctatgctggaaacagc |
| Prpf8 | Prpf8-3 | CGGGAACGGATCCGTCGCG | tatcttgtggaaaggacgaaacaccGCGGGAACGGATCCGTCGCGgtttaagagctatgctggaaacagc |
| Prpf8 | Prpf8-4 | GGCGAGTGCAACGTCATGC | tatcttgtggaaaggacgaaacaccGGGCGAGTGCAACGTCATGCgtttaagagctatgctggaaacagc |
| Prpf8 | Prpf8-5 | GTACTGTAGTCACCGATCG | tatcttgtggaaaggacgaaacaccGGTACTGTAGTCACCGATCGgtttaagagctatgctggaaacagc |
| Prss48 | Prss48-1 | ACGCCTCACTGGAAATGAC | tatcttgtggaaaggacgaaacaccGACGCCTCACTGGAAATGACgtttaagagctatgctggaaacagc |
| Prss48 | Prss48-2 | GGCAGAATGACCGATGAGA | tatcttgtggaaaggacgaaacaccGGGCAGAATGACCGATGAGAgtttaagagctatgctggaaacagc |
| Prss48 | Prss48-3 | CTTGAGTACTCTCTATCGA | tatcttgtggaaaggacgaaacaccGCTTGAGTACTCTCTATCGAgtttaagagctatgctggaaacagc |
| Prss54 | Prss54-2 | AATGCATTAACTGCAGACG | tatcttgtggaaaggacgaaacaccGAATGCATTAACTGCAGACGgtttaagagctatgctggaaacagc |
| Prss54 | Prss54-3 | GCCATGCATTTCAACGACC | tatcttgtggaaaggacgaaacaccGGCCATGCATTTCAACGACCgtttaagagctatgctggaaacagc |
| Prss54 | Prss54-5 | GTTGTTGGTATATCTAACA | tatcttgtggaaaggacgaaacaccGGTTGTTGGTATATCTAACAgtttaagagctatgctggaaacagc |
| Psip1 | Psip1-1 | GAAAGAAGTTGAATCTAAA | tatcttgtggaaaggacgaaacaccGGAAAGAAGTTGAATCTAAAgtttaagagctatgctggaaacagc |
| Psip1 | Psip1-3 | CTAAGAGAGGACGACCTGC | tatcttgtggaaaggacgaaacaccGCTAAGAGAGGACGACCTGCgtttaagagctatgctggaaacagc |
| Psip1 | Psip1-4 | CTTGCTTTCACATTAGAAG | tatcttgtggaaaggacgaaacaccGCTTGCTTTCACATTAGAAGgtttaagagctatgctggaaacagc |
| Psma4 | Psma4-3 | CGTTCTGACTAACGAGCTA | tatcttgtggaaaggacgaaacaccGCGTTCTGACTAACGAGCTAgtttaagagctatgctggaaacagc |
| Psma4 | Psma4-4 | GGCTCATTGCTCAACGGTA | tatcttgtggaaaggacgaaacaccGGGCTCATTGCTCAACGGTAgtttaagagctatgctggaaacagc |
| Psma4 | Psma4-5 | GCACTGTGTGATATCAAAC | tatcttgtggaaaggacgaaacaccGGCACTGTGTGATATCAAACgtttaagagctatgctggaaacagc |
| Psmd6 | Psmd6-1 | GCTATCCGTGACTTCAAAC | tatcttgtggaaaggacgaaacaccGGCTATCCGTGACTTCAAACgtttaagagctatgctggaaacagc |
| Psmd6 | Psmd6-2 | AGCCACACAATATAAGCCC | tatcttgtggaaaggacgaaacaccGAGCCACACAATATAAGCCCgtttaagagctatgctggaaacagc |
| Psmd6 | Psmd6-3 | AATATCCAGGCGGTGACCC | tatcttgtggaaaggacgaaacaccGAATATCCAGGCGGTGACCCgtttaagagctatgctggaaacagc |
| Psme3ip1 | Psme3ip1-1 | ACTCACAGTAATCTCAACA | tatcttgtggaaaggacgaaacaccGACTCACAGTAATCTCAACAgtttaagagctatgctggaaacagc |
| Psme3ip1 | Psme3ip1-2 | CGGCGAGAAGAAGAACTAG | tatcttgtggaaaggacgaaacaccGCGGCGAGAAGAAGAACTAGgtttaagagctatgctggaaacagc |
| Psme3ip1 | Psme3ip1-4 | CCTCAGAAAACATGGTAAG | tatcttgtggaaaggacgaaacaccGCCTCAGAAAACATGGTAAGgtttaagagctatgctggaaacagc |
| Psmf1 | Psmf1-2 | TCCTTAGACTCATACCGGA | tatcttgtggaaaggacgaaacaccGTCCTTAGACTCATACCGGAgtttaagagctatgctggaaacagc |
| Psmf1 | Psmf1-3 | AGAACTACTGCCAGCTAAG | tatcttgtggaaaggacgaaacaccGAGAACTACTGCCAGCTAAGgtttaagagctatgctggaaacagc |
| Psmf1 | Psmf1-5 | GCTACTATGCCTTGGGCAC | tatcttgtggaaaggacgaaacaccGGCTACTATGCCTTGGGCACgtttaagagctatgctggaaacagc |
| Psmg1 | Psmg1-1 | AAGGCTGCAGGTAGACTCG | tatcttgtggaaaggacgaaacaccGAAGGCTGCAGGTAGACTCGgtttaagagctatgctggaaacagc |
| Psmg1 | Psmg1-2 | TGTTATGAACTCGGGAGTC | tatcttgtggaaaggacgaaacaccGTGTTATGAACTCGGGAGTCgtttaagagctatgctggaaacagc |
| Psmg1 | Psmg1-3 | CCTACTGCAATTATAAACT | tatcttgtggaaaggacgaaacaccGCCTACTGCAATTATAAACTgtttaagagctatgctggaaacagc |
| Psmg2 | Psmg2-3 | ACTAGCTTCTTTGACGGTA | tatcttgtggaaaggacgaaacaccGACTAGCTTCTTTGACGGTAgtttaagagctatgctggaaacagc |
| Psmg2 | Psmg2-4 | CTTGAGAGAACAATGATCC | tatcttgtggaaaggacgaaacaccGCTTGAGAGAACAATGATCCgtttaagagctatgctggaaacagc |
| Psmg2 | Psmg2-5 | ATCGTTGCGGTGATACGAA | tatcttgtggaaaggacgaaacaccGATCGTTGCGGTGATACGAAgtttaagagctatgctggaaacagc |
| Psmg4 | Psmg4-1 | CAGCCGCATGACATGAAAG | tatcttgtggaaaggacgaaacaccGCAGCCGCATGACATGAAAGgtttaagagctatgctggaaacagc |
| Psmg4 | Psmg4-2 | CGGCGTTGCTCCCACCCAC | tatcttgtggaaaggacgaaacaccGCGGCGTTGCTCCCACCCACgtttaagagctatgctggaaacagc |
| Psmg4 | Psmg4-3 | CTGCGAGGTTGCGTAGATG | tatcttgtggaaaggacgaaacaccGCTGCGAGGTTGCGTAGATGgtttaagagctatgctggaaacagc |
| Ptcd3 | Ptcd3-1 | CTACTGCACGATGATCCGA | tatcttgtggaaaggacgaaacaccGCTACTGCACGATGATCCGAgtttaagagctatgctggaaacagc |
| Ptcd3 | Ptcd3-3 | TGAGCCTCATATACCGGTA | tatcttgtggaaaggacgaaacaccGTGAGCCTCATATACCGGTAgtttaagagctatgctggaaacagc |
| Ptcd3 | Ptcd3-5 | TCTTATTCTAGGGATAAAG | tatcttgtggaaaggacgaaacaccGTCTTATTCTAGGGATAAAGgtttaagagctatgctggaaacagc |
| Pten | Pten-1 | CCGCCAAATTTAACTGCAG | tatcttgtggaaaggacgaaacaccGCCGCCAAATTTAACTGCAGgtttaagagctatgctggaaacagc |
| Pten | Pten-2 | GTTTGATAAGTTCTAGCTG | tatcttgtggaaaggacgaaacaccGGTTTGATAAGTTCTAGCTGgtttaagagctatgctggaaacagc |
| Pten | Pten-5 | TATACATAGCGCCTCTGAC | tatcttgtggaaaggacgaaacaccGTATACATAGCGCCTCTGACgtttaagagctatgctggaaacagc |
| Ptpn2 | Ptpn2-1 | ACTCTGGAACCCCAAAATC | tatcttgtggaaaggacgaaacaccGACTCTGGAACCCCAAAATCgtttaagagctatgctggaaacagc |
| Ptpn2 | Ptpn2-4 | CGAACTGTAGAAAAAGAAT | tatcttgtggaaaggacgaaacaccGCGAACTGTAGAAAAAGAATgtttaagagctatgctggaaacagc |
| Ptpn2 | Ptpn2-5 | GCAGCATGTGTTCGGAAGT | tatcttgtggaaaggacgaaacaccGGCAGCATGTGTTCGGAAGTgtttaagagctatgctggaaacagc |
| Puf60 | Puf60-1 | CTTGCCAGTTGTGGGGTCC | tatcttgtggaaaggacgaaacaccGCTTGCCAGTTGTGGGGTCCgtttaagagctatgctggaaacagc |
| Puf60 | Puf60-4 | TTCATGGTAACGGAGTCCC | tatcttgtggaaaggacgaaacaccGTTCATGGTAACGGAGTCCCgtttaagagctatgctggaaacagc |
| Puf60 | Puf60-5 | CCCCTGACGCCCGAGCAGC | tatcttgtggaaaggacgaaacaccGCCCCTGACGCCCGAGCAGCgtttaagagctatgctggaaacagc |
| Rab11fip2 | Rab11fip2-2 | CTGGCAGCATTGTGGATGA | tatcttgtggaaaggacgaaacaccGCTGGCAGCATTGTGGATGAgtttaagagctatgctggaaacagc |
| Rab11fip2 | Rab11fip2-3 | GTTAAATCAGACATCGAAT | tatcttgtggaaaggacgaaacaccGGTTAAATCAGACATCGAATgtttaagagctatgctggaaacagc |
| Rab11fip2 | Rab11fip2-5 | GTGCTCGGAACGATGGCAG | tatcttgtggaaaggacgaaacaccGGTGCTCGGAACGATGGCAGgtttaagagctatgctggaaacagc |
| Rab1a | Rab1a-2 | TACGAACTATAGAGTTAGA | tatcttgtggaaaggacgaaacaccGTACGAACTATAGAGTTAGAgtttaagagctatgctggaaacagc |
| Rab1a | Rab1a-3 | TCACTTCCAGTTATTACAG | tatcttgtggaaaggacgaaacaccGTCACTTCCAGTTATTACAGgtttaagagctatgctggaaacagc |
| Rab1a | Rab1a-4 | TACACAACTATGATGCCAT | tatcttgtggaaaggacgaaacaccGTACACAACTATGATGCCATgtttaagagctatgctggaaacagc |
| Rac1 | Rac1-1 | ATCAAGCTTCGTCCCCACG | tatcttgtggaaaggacgaaacaccGATCAAGCTTCGTCCCCACGgtttaagagctatgctggaaacagc |
| Rac1 | Rac1-3 | ACTGTCTGCGGGTAGGAGA | tatcttgtggaaaggacgaaacaccGACTGTCTGCGGGTAGGAGAgtttaagagctatgctggaaacagc |
| Rac1 | Rac1-4 | ACCAGTGAATCTGGGCCTA | tatcttgtggaaaggacgaaacaccGACCAGTGAATCTGGGCCTAgtttaagagctatgctggaaacagc |
| Rad17 | Rad17-2 | GGGATCCAAGTACAAGAAT | tatcttgtggaaaggacgaaacaccGGGGATCCAAGTACAAGAATgtttaagagctatgctggaaacagc |
| Rad17 | Rad17-3 | TATTAATAACAGGTCCTCC | tatcttgtggaaaggacgaaacaccGTATTAATAACAGGTCCTCCgtttaagagctatgctggaaacagc |
| Rad17 | Rad17-4 | AATTGAAGAGGTCGAAACC | tatcttgtggaaaggacgaaacaccGAATTGAAGAGGTCGAAACCgtttaagagctatgctggaaacagc |
| Rad50 | Rad50-1 | GAAATTGTCAGGTCCTACG | tatcttgtggaaaggacgaaacaccGGAAATTGTCAGGTCCTACGgtttaagagctatgctggaaacagc |
| Rad50 | Rad50-2 | CGAGATCAGATTACGAGTA | tatcttgtggaaaggacgaaacaccGCGAGATCAGATTACGAGTAgtttaagagctatgctggaaacagc |
| Rad50 | Rad50-5 | TTGGTTGGACCCAATGGGG | tatcttgtggaaaggacgaaacaccGTTGGTTGGACCCAATGGGGgtttaagagctatgctggaaacagc |
| Rad51 | Rad51-1 | GAAGCCGGTTACCATACAG | tatcttgtggaaaggacgaaacaccGGAAGCCGGTTACCATACAGgtttaagagctatgctggaaacagc |
| Rad51 | Rad51-4 | GCCATGTACATTGACACCG | tatcttgtggaaaggacgaaacaccGGCCATGTACATTGACACCGgtttaagagctatgctggaaacagc |
| Rad51 | Rad51-5 | GGGCACCTTTAGGCCGGAG | tatcttgtggaaaggacgaaacaccGGGGCACCTTTAGGCCGGAGgtttaagagctatgctggaaacagc |
| Rad51ap1 | Rad51ap1-3 | GACGCCTCGAGTGTTGAAG | tatcttgtggaaaggacgaaacaccGGACGCCTCGAGTGTTGAAGgtttaagagctatgctggaaacagc |
| Rad51ap1 | Rad51ap1-4 | GATTTGGACAAGATCACAG | tatcttgtggaaaggacgaaacaccGGATTTGGACAAGATCACAGgtttaagagctatgctggaaacagc |
| Rad51ap1 | Rad51ap1-5 | AGAGGTCTAGAAGTTGCCC | tatcttgtggaaaggacgaaacaccGAGAGGTCTAGAAGTTGCCCgtttaagagctatgctggaaacagc |
| Rad51d | Rad51d-1 | TCTAGCCTGTAGTAGCTGG | tatcttgtggaaaggacgaaacaccGTCTAGCCTGTAGTAGCTGGgtttaagagctatgctggaaacagc |
| Rad51d | Rad51d-4 | GCTGGCCTCTATACTGGGG | tatcttgtggaaaggacgaaacaccGGCTGGCCTCTATACTGGGGgtttaagagctatgctggaaacagc |
| Rad51d | Rad51d-5 | CCTTCCGATGCCGGTGGAC | tatcttgtggaaaggacgaaacaccGCCTTCCGATGCCGGTGGACgtttaagagctatgctggaaacagc |
| Rad9a | Rad9a-2 | CTTATTGCGCACCCCAGCA | tatcttgtggaaaggacgaaacaccGCTTATTGCGCACCCCAGCAgtttaagagctatgctggaaacagc |
| Rad9a | Rad9a-4 | GACGAGTTCACAGTCCGTA | tatcttgtggaaaggacgaaacaccGGACGAGTTCACAGTCCGTAgtttaagagctatgctggaaacagc |

| | | | |
|---|---|---|---|
| Rad9a | Rad9a-5 | ACCCTTGAAGGACGGGGTA | tatcttgtggaaaggacgaaacaccGACCCTTGAAGGACGGGGTAgtttaagagctatgctggaaacagc |
| Rae1 | Rae1-1 | CGCCCATGATCCTGCAATA | tatcttgtggaaaggacgaaacaccGCGCCCATGATCCTGCAATAgtttaagagctatgctggaaacagc |
| Rae1 | Rae1-3 | CTGTGTGATGACCGGGAGC | tatcttgtggaaaggacgaaacaccGCTGTGTGATGACCGGGAGCgtttaagagctatgctggaaacagc |
| Rae1 | Rae1-4 | CGCTGTTACTGTGCAGATG | tatcttgtggaaaggacgaaacaccGCGCTGTTACTGTGCAGATGgtttaagagctatgctggaaacagc |
| Raf1 | Raf1-1 | CACAAAGAGAGCGGGCACC | tatcttgtggaaaggacgaaacaccGCACAAAGAGAGCGGGCACCgtttaagagctatgctggaaacagc |
| Raf1 | Raf1-2 | CAAACAACCTGAGTCCAAC | tatcttgtggaaaggacgaaacaccGCAAACAACCTGAGTCCAACgtttaagagctatgctggaaacagc |
| Raf1 | Raf1-3 | AAGTACAACATTACCTAGC | tatcttgtggaaaggacgaaacaccGAAGTACAACATTACCTAGCgtttaagagctatgctggaaacagc |
| Ralyl | Ralyl-1 | AATGTTTCTTACCTGTACA | tatcttgtggaaaggacgaaacaccGAATGTTTCTTACCTGTACAgtttaagagctatgctggaaacagc |
| Ralyl | Ralyl-2 | TTTTACAGGCTTGAATCAA | tatcttgtggaaaggacgaaacaccGTTTTACAGGCTTGAATCAAgtttaagagctatgctggaaacagc |
| Ralyl | Ralyl-3 | GTCAAATACGTAACCACTA | tatcttgtggaaaggacgaaacaccGGTCAAATACGTAACCACTAgtttaagagctatgctggaaacagc |
| Ranbp1 | Ranbp1-2 | GCGTGGGTATTCCAGACCC | tatcttgtggaaaggacgaaacaccGGCGTGGGTATTCCAGACCCgtttaagagctatgctggaaacagc |
| Ranbp1 | Ranbp1-4 | ATCCGCCTTCTTATGAGGA | tatcttgtggaaaggacgaaacaccGATCCGCCTTCTTATGAGGAgtttaagagctatgctggaaacagc |
| Ranbp1 | Ranbp1-5 | TCGCTCCTTCCATTCTGGG | tatcttgtggaaaggacgaaacaccGTCGCTCCTTCCATTCTGGGgtttaagagctatgctggaaacagc |
| Rbbp8 | Rbbp8-3 | AGCGCTCTCTATGTACAAA | tatcttgtggaaaggacgaaacaccGAGCGCTCTCTATGTACAAAgtttaagagctatgctggaaacagc |
| Rbbp8 | Rbbp8-4 | TACTATTGAAGGAACCCCA | tatcttgtggaaaggacgaaacaccGTACTATTGAAGGAACCCCAgtttaagagctatgctggaaacagc |
| Rbbp8 | Rbbp8-5 | TAGAGGTAGCTCCAAATAC | tatcttgtggaaaggacgaaacaccGTAGAGGTAGCTCCAAATACgtttaagagctatgctggaaacagc |
| Rbm19 | Rbm19-1 | GAGCTTTATTGACACGACC | tatcttgtggaaaggacgaaacaccGGAGCTTTATTGACACGACCgtttaagagctatgctggaaacagc |
| Rbm19 | Rbm19-2 | CCCATCGAAGCCCCGAGCT | tatcttgtggaaaggacgaaacaccGCCCATCGAAGCCCCGAGCTgtttaagagctatgctggaaacagc |
| Rbm19 | Rbm19-5 | AGCTTTTCTCGGCCTACGG | tatcttgtggaaaggacgaaacaccGAGCTTTTCTCGGCCTACGGgtttaagagctatgctggaaacagc |
| Rbm22 | Rbm22-1 | CGAAACACTGTGAATGGCC | tatcttgtggaaaggacgaaacaccGCGAAACACTGTGAATGGCCgtttaagagctatgctggaaacagc |
| Rbm22 | Rbm22-2 | TGAAGCGCATGCGGACCCC | tatcttgtggaaaggacgaaacaccGTGAAGCGCATGCGGACCCCgtttaagagctatgctggaaacagc |
| Rbm22 | Rbm22-5 | TATGGAATCAACGACCCCG | tatcttgtggaaaggacgaaacaccGTATGGAATCAACGACCCCGgtttaagagctatgctggaaacagc |
| Rbm31y | Rbm31y-1 | GCCAGTAAGATGTTCATCG | tatcttgtggaaaggacgaaacaccGGCCAGTAAGATGTTCATCGgtttaagagctatgctggaaacagc |
| Rbm31y | Rbm31y-3 | GATGAAAACTCCGTTAGAA | tatcttgtggaaaggacgaaacaccGGATGAAAACTCCGTTAGAAgtttaagagctatgctggaaacagc |
| Rbm31y | Rbm31y-4 | AGCCGTTGCGAAGTTAAAA | tatcttgtggaaaggacgaaacaccGAGCCGTTGCGAAGTTAAAAgtttaagagctatgctggaaacagc |
| Rbm39 | Rbm39-2 | ATTTACAAAAGGGAAGTGC | tatcttgtggaaaggacgaaacaccGATTTACAAAAGGGAAGTGCgtttaagagctatgctggaaacagc |
| Rbm39 | Rbm39-3 | GGCTCTTGTCTTTCCTAAA | tatcttgtggaaaggacgaaacaccGGGCTCTTGTCTTTCCTAAAgtttaagagctatgctggaaacagc |
| Rbm39 | Rbm39-5 | AAAGAGCAAAAGTCGAGAG | tatcttgtggaaaggacgaaacaccGAAAGAGCAAAAGTCGAGAGgtttaagagctatgctggaaacagc |
| Rbm41 | Rbm41-2 | AAACCTCTGTGGCAGGCAA | tatcttgtggaaaggacgaaacaccGAAACCTCTGTGGCAGGCAAgtttaagagctatgctggaaacagc |
| Rbm41 | Rbm41-3 | AGCCATGTTTCCGGTGAAA | tatcttgtggaaaggacgaaacaccGAGCCATGTTTCCGGTGAAAgtttaagagctatgctggaaacagc |
| Rbm41 | Rbm41-4 | ATGTACAAGCCCTTTGGGA | tatcttgtggaaaggacgaaacaccGATGTACAAGCCCTTTGGGAgtttaagagctatgctggaaacagc |
| Rbmx2 | Rbmx2-1 | GAAAAGGTGTCTTGGCACT | tatcttgtggaaaggacgaaacaccGGAAAAGGTGTCTTGGCACTgtttaagagctatgctggaaacagc |
| Rbmx2 | Rbmx2-2 | TTCCTTATGAATTGACAGA | tatcttgtggaaaggacgaaacaccGTTCCTTATGAATTGACAGAgtttaagagctatgctggaaacagc |
| Rbmx2 | Rbmx2-3 | CCTGTGCTATGAAGACCAG | tatcttgtggaaaggacgaaacaccGCCTGTGCTATGAAGACCAGgtttaagagctatgctggaaacagc |
| Rcc1 | Rcc1-1 | TGGTGATGCTCACGAACGA | tatcttgtggaaaggacgaaacaccGTGGTGATGCTCACGAACGAgtttaagagctatgctggaaacagc |
| Rcc1 | Rcc1-2 | GATGCGCCCGTGGTAAAGG | tatcttgtggaaaggacgaaacaccGGATGCGCCCGTGGTAAAGGgtttaagagctatgctggaaacagc |
| Rcc1 | Rcc1-5 | GCAATGACGAGGGTGCCCT | tatcttgtggaaaggacgaaacaccGGCAATGACGAGGGTGCCCTgtttaagagctatgctggaaacagc |
| Retreg1 | Retreg1-1 | TCTTGGGAAGTTACATTCC | tatcttgtggaaaggacgaaacaccGTCTTGGGAAGTTACATTCCgtttaagagctatgctggaaacagc |
| Retreg1 | Retreg1-2 | GGTAGCTGAGTATGACCCC | tatcttgtggaaaggacgaaacaccGGGTAGCTGAGTATGACCCCgtttaagagctatgctggaaacagc |
| Retreg1 | Retreg1-4 | CGGAGTGTACCCCTCTGAA | tatcttgtggaaaggacgaaacaccGCGGAGTGTACCCCTCTGAAgtttaagagctatgctggaaacagc |
| Rfwd3 | Rfwd3-1 | TGAACAGGAGCGCATGAAA | tatcttgtggaaaggacgaaacaccGTGAACAGGAGCGCATGAAAgtttaagagctatgctggaaacagc |
| Rfwd3 | Rfwd3-2 | CTAGTGTCCAAAGCTCGCA | tatcttgtggaaaggacgaaacaccGCTAGTGTCCAAAGCTCGCAgtttaagagctatgctggaaacagc |
| Rfwd3 | Rfwd3-3 | GCGTAGCGCTGAGATCCGG | tatcttgtggaaaggacgaaacaccGGCGTAGCGCTGAGATCCGGgtttaagagctatgctggaaacagc |
| Rfx7 | Rfx7-1 | GAGACGTCACTCCCCAAAC | tatcttgtggaaaggacgaaacaccGGAGACGTCACTCCCCAAACgtttaagagctatgctggaaacagc |
| Rfx7 | Rfx7-2 | CACGCCGTTTAGGCACTAG | tatcttgtggaaaggacgaaacaccGCACGCCGTTTAGGCACTAGgtttaagagctatgctggaaacagc |
| Rfx7 | Rfx7-4 | TGATAATATTACCGGAATG | tatcttgtggaaaggacgaaacaccGTGATAATATTACCGGAATGgtttaagagctatgctggaaacagc |
| Rgs1 | Rgs1-1 | TAGGAATACTTACTCTGGT | tatcttgtggaaaggacgaaacaccGTAGGAATACTTACTCTGGTgtttaagagctatgctggaaacagc |
| Rgs1 | Rgs1-4 | AGATCGATGATCCCACATC | tatcttgtggaaaggacgaaacaccGAGATCGATGATCCCACATCgtttaagagctatgctggaaacagc |
| Rgs1 | Rgs1-5 | GGTGTCTTTTGTAGTGGAA | tatcttgtggaaaggacgaaacaccGGGTGTCTTTTGTAGTGGAAgtttaagagctatgctggaaacagc |
| Rhag | Rhag-2 | TGACCTTTCTGAAGAAATA | tatcttgtggaaaggacgaaacaccGTGACCTTTCTGAAGAAATAgtttaagagctatgctggaaacagc |
| Rhag | Rhag-4 | AATGGGGCTTGTTTTCCCC | tatcttgtggaaaggacgaaacaccGAATGGGGCTTGTTTTCCCCgtttaagagctatgctggaaacagc |
| Rhag | Rhag-5 | CAGGTGTGTTATATCGGCC | tatcttgtggaaaggacgaaacaccGCAGGTGTGTTATATCGGCCgtttaagagctatgctggaaacagc |
| Rhbdd3 | Rhbdd3-1 | GGCCTTGGGGTGTCCGGTG | tatcttgtggaaaggacgaaacaccGGGCCTTGGGGTGTCCGGTGgtttaagagctatgctggaaacagc |
| Rhbdd3 | Rhbdd3-3 | CAGGCAAAGGATACAGGCT | tatcttgtggaaaggacgaaacaccGCAGGCAAAGGATACAGGCTgtttaagagctatgctggaaacagc |
| Rhbdd3 | Rhbdd3-4 | GACTGCAGGTGTTACAGGA | tatcttgtggaaaggacgaaacaccGGACTGCAGGTGTTACAGGAgtttaagagctatgctggaaacagc |
| Ric8a | Ric8a-1 | ATTCGGATAGTCTGCAGCC | tatcttgtggaaaggacgaaacaccGATTCGGATAGTCTGCAGCCgtttaagagctatgctggaaacagc |
| Ric8a | Ric8a-4 | GTCCCCAGGTACCGGTAAA | tatcttgtggaaaggacgaaacaccGGTCCCCAGGTACCGGTAAAgtttaagagctatgctggaaacagc |
| Ric8a | Ric8a-5 | CTTCTGCGGCACTGCGTGA | tatcttgtggaaaggacgaaacaccGCTTCTGCGGCACTGCGTGAgtttaagagctatgctggaaacagc |
| Rmi1 | Rmi1-1 | GTAGCATCAGCATCCGAGA | tatcttgtggaaaggacgaaacaccGGTAGCATCAGCATCCGAGAgtttaagagctatgctggaaacagc |
| Rmi1 | Rmi1-2 | ACTTTGGGCACGTTGTGAA | tatcttgtggaaaggacgaaacaccGACTTTGGGCACGTTGTGAAgtttaagagctatgctggaaacagc |
| Rmi1 | Rmi1-3 | TCTGCAGCAAATAATGACG | tatcttgtggaaaggacgaaacaccGTCTGCAGCAAATAATGACGgtttaagagctatgctggaaacagc |
| Rmi2 | Rmi2-2 | GGGCGCGTGGCGGCTGTCG | tatcttgtggaaaggacgaaacaccGGGGCGCGTGGCGGCTGTCGgtttaagagctatgctggaaacagc |
| Rmi2 | Rmi2-3 | CAGGGTACCGTGCTGGCGG | tatcttgtggaaaggacgaaacaccGCAGGGTACCGTGCTGGCGGgtttaagagctatgctggaaacagc |
| Rmi2 | Rmi2-5 | GGAGCGCGTACCACGCGGC | tatcttgtggaaaggacgaaacaccGGGAGCGCGTACCACGCGGCgtttaagagctatgctggaaacagc |
| Rnaseh2b | Rnaseh2b-2 | GGCCAACCCGCACTCAGGT | tatcttgtggaaaggacgaaacaccGGGCCAACCCGCACTCAGGTgtttaagagctatgctggaaacagc |
| Rnaseh2b | Rnaseh2b-3 | TAAATCAATCAGTTCAATC | tatcttgtggaaaggacgaaacaccGTAAATCAATCAGTTCAATCgtttaagagctatgctggaaacagc |
| Rnaseh2b | Rnaseh2b-4 | CTCCTTCACTATCTCCTAA | tatcttgtggaaaggacgaaacaccGCTCCTTCACTATCTCCTAAgtttaagagctatgctggaaacagc |
| Rnasek | Rnasek-1 | GTAGCTGACTTGCTCGTAC | tatcttgtggaaaggacgaaacaccGGTAGCTGACTTGCTCGTACgtttaagagctatgctggaaacagc |
| Rnasek | Rnasek-2 | ACAGGTTGTATATGTTCTG | tatcttgtggaaaggacgaaacaccGACAGGTTGTATATGTTCTGgtttaagagctatgctggaaacagc |
| Rnasek | Rnasek-4 | CTCAATTAACACAGCAGAA | tatcttgtggaaaggacgaaacaccGCTCAATTAACACAGCAGAAgtttaagagctatgctggaaacagc |
| Rnf168 | Rnf168-1 | AGACCCGGCGACGACAGAA | tatcttgtggaaaggacgaaacaccGAGACCCGGCGACGACAGAAgtttaagagctatgctggaaacagc |
| Rnf168 | Rnf168-4 | TTGAGGCGAGAATATGAAG | tatcttgtggaaaggacgaaacaccGTTGAGGCGAGAATATGAAGgtttaagagctatgctggaaacagc |
| Rnf168 | Rnf168-5 | CTGTGTAAACTTACGATAC | tatcttgtggaaaggacgaaacaccGCTGTGTAAACTTACGATACgtttaagagctatgctggaaacagc |
| Rnf8 | Rnf8-1 | TTCAGGTGACCATAGGACG | tatcttgtggaaaggacgaaacaccGTTCAGGTGACCATAGGACGgtttaagagctatgctggaaacagc |
| Rnf8 | Rnf8-2 | GGCTTCGGGAAATCATCAG | tatcttgtggaaaggacgaaacaccGGGCTTCGGGAAATCATCAGgtttaagagctatgctggaaacagc |
| Rnf8 | Rnf8-3 | AGGGTTATTGCATCCGTAA | tatcttgtggaaaggacgaaacaccGAGGGTTATTGCATCCGTAAgtttaagagctatgctggaaacagc |

| | | | |
|---|---|---|---|
| Rpa1 | Rpa1-1 | ACCTGGAGTAATTCCCGGG | tatcttgtggaaaggacgaaacaccGACCTGGAGTAATTCCCGGGgtttaagagctatgctggaaacagc |
| Rpa1 | Rpa1-4 | CAACACCCTGAAAGACGGC | tatcttgtggaaaggacgaaacaccGCAACACCCTGAAAGACGGCgtttaagagctatgctggaaacagc |
| Rpa1 | Rpa1-5 | AGCTGAACACCCTGGTCGA | tatcttgtggaaaggacgaaacaccGAGCTGAACACCCTGGTCGAgtttaagagctatgctggaaacagc |
| Rpa2 | Rpa2-3 | GATATGACAGCTCCGCCCA | tatcttgtggaaaggacgaaacaccGGATATGACAGCTCCGCCCAgtttaagagctatgctggaaacagc |
| Rpa2 | Rpa2-4 | GTTCGTCAGTGGGTTGACA | tatcttgtggaaaggacgaaacaccGGTTCGTCAGTGGGTTGACAgtttaagagctatgctggaaacagc |
| Rpa2 | Rpa2-5 | CTTTCACGTATGTTTCGGG | tatcttgtggaaaggacgaaacaccGCTTTCACGTATGTTTCGGGgtttaagagctatgctggaaacagc |
| Rpain | Rpain-1 | TTACCTCCCCAGGGATGTC | tatcttgtggaaaggacgaaacaccGTTACCTCCCCAGGGATGTCgtttaagagctatgctggaaacagc |
| Rpain | Rpain-3 | CTCCTGAACAAATATCGCC | tatcttgtggaaaggacgaaacaccGCTCCTGAACAAATATCGCCgtttaagagctatgctggaaacagc |
| Rpain | Rpain-7 | TGACCTGAAGCAAGGCCTC | tatcttgtggaaaggacgaaacaccGTGACCTGAAGCAAGGCCTCgtttaagagctatgctggaaacagc |
| Rpf2 | Rpf2-3 | ACATCGGTTTTGTTCCCTC | tatcttgtggaaaggacgaaacaccGACATCGGTTTTGTTCCCTCgtttaagagctatgctggaaacagc |
| Rpf2 | Rpf2-4 | TAGACTAGTAAGTGCCCCG | tatcttgtggaaaggacgaaacaccGTAGACTAGTAAGTGCCCCGgtttaagagctatgctggaaacagc |
| Rpf2 | Rpf2-5 | TTCAATCATATCCAGCACA | tatcttgtggaaaggacgaaacaccGTTCAATCATATCCAGCACAgtttaagagctatgctggaaacagc |
| Rpl7l1 | Rpl7l1-1 | GAAAATGGCACTCACCCCC | tatcttgtggaaaggacgaaacaccGGAAAATGGCACTCACCCCCgtttaagagctatgctggaaacagc |
| Rpl7l1 | Rpl7l1-2 | TGCGTATAACGAAGGCCAA | tatcttgtggaaaggacgaaacaccGTGCGTATAACGAAGGCCAAgtttaagagctatgctggaaacagc |
| Rpl7l1 | Rpl7l1-3 | GTGCGTGTCCAGCGGCTGG | tatcttgtggaaaggacgaaacaccGGTGCGTGTCCAGCGGCTGGgtttaagagctatgctggaaacagc |
| Rps21 | Rps21-1 | GCGTCCATCCAGATGAACG | tatcttgtggaaaggacgaaacaccGGCGTCCATCCAGATGAACGgtttaagagctatgctggaaacagc |
| Rps21 | Rps21-2 | GCCTGTGGTCCTATCAACC | tatcttgtggaaaggacgaaacaccGGCCTGTGGTCCTATCAACCgtttaagagctatgctggaaacagc |
| Rps21 | Rps21-5 | CATCTGCGGGGCCATTCGC | tatcttgtggaaaggacgaaacaccGCATCTGCGGGGCCATTCGCgtttaagagctatgctggaaacagc |
| Rps5 | Rps5-1 | AGCATACCGTCCGGCACTG | tatcttgtggaaaggacgaaacaccGAGCATACCGTCCGGCACTGgtttaagagctatgctggaaacagc |
| Rps5 | Rps5-2 | TGTGCTTTGCGGAAGCGCT | tatcttgtggaaaggacgaaacaccGTGTGCTTTGCGGAAGCGCTgtttaagagctatgctggaaacagc |
| Rps5 | Rps5-5 | CCTTACCTCACCAGTGAGC | tatcttgtggaaaggacgaaacaccGCCTTACCTCACCAGTGAGCgtttaagagctatgctggaaacagc |
| Rptor | Rptor-1 | TACGACCTGCAGACGTGGA | tatcttgtggaaaggacgaaacaccGTACGACCTGCAGACGTGGAgtttaagagctatgctggaaacagc |
| Rptor | Rptor-3 | GGTGCGGTGTCTCACCCCG | tatcttgtggaaaggacgaaacaccGGGTGCGGTGTCTCACCCCGgtttaagagctatgctggaaacagc |
| Rptor | Rptor-4 | GTATGTCGGGGGGAGCCGC | tatcttgtggaaaggacgaaacaccGGTATGTCGGGGGGAGCCGCgtttaagagctatgctggaaacagc |
| Rpusd4 | Rpusd4-2 | CTGCACTCTCCGCTGAACG | tatcttgtggaaaggacgaaacaccGCTGCACTCTCCGCTGAACGgtttaagagctatgctggaaacagc |
| Rpusd4 | Rpusd4-4 | CTATTTAGAACCCGCCAGG | tatcttgtggaaaggacgaaacaccGCTATTTAGAACCCGCCAGGgtttaagagctatgctggaaacagc |
| Rpusd4 | Rpusd4-5 | ACACGCACCGTGATGGCCC | tatcttgtggaaaggacgaaacaccGACACGCACCGTGATGGCCCgtttaagagctatgctggaaacagc |
| Rrm1 | Rrm1-2 | CTAAGTCCGCTGGGGGAAT | tatcttgtggaaaggacgaaacaccGCTAAGTCCGCTGGGGGAATgtttaagagctatgctggaaacagc |
| Rrm1 | Rrm1-3 | CCACTGGTAGCTACATCGC | tatcttgtggaaaggacgaaacaccGCCACTGGTAGCTACATCGCgtttaagagctatgctggaaacagc |
| Rrm1 | Rrm1-5 | GGTAAATAGCAAACGCGCC | tatcttgtggaaaggacgaaacaccGGGTAAATAGCAAACGCGCCgtttaagagctatgctggaaacagc |
| Rrs1 | Rrs1-2 | CTGGCGCAGCACGGTCGGG | tatcttgtggaaaggacgaaacaccGCTGGCGCAGCACGGTCGGGgtttaagagctatgctggaaacagc |
| Rrs1 | Rrs1-3 | CTGCGTGTTGTCCCGCGCC | tatcttgtggaaaggacgaaacaccGCTGCGTGTTGTCCCGCGCCgtttaagagctatgctggaaacagc |
| Rrs1 | Rrs1-5 | GGCGCGCTTGTAGCCCCAA | tatcttgtggaaaggacgaaacaccGGGCGCGCTTGTAGCCCCAAgtttaagagctatgctggaaacagc |
| Rspo3 | Rspo3-1 | ACAGGTTACCCTTGGCTGA | tatcttgtggaaaggacgaaacaccGACAGGTTACCCTTGGCTGAgtttaagagctatgctggaaacagc |
| Rspo3 | Rspo3-2 | GTGGATTTTACTTACACCT | tatcttgtggaaaggacgaaacaccGGTGGATTTTACTTACACCTgtttaagagctatgctggaaacagc |
| Rspo3 | Rspo3-3 | GAGTTCCGTAATATCCACT | tatcttgtggaaaggacgaaacaccGGAGTTCCGTAATATCCACTgtttaagagctatgctggaaacagc |
| Ruvbl1 | Ruvbl1-2 | TACCCACCATCGGGCAGAA | tatcttgtggaaaggacgaaacaccGTACCCACCATCGGGCAGAAgtttaagagctatgctggaaacagc |
| Ruvbl1 | Ruvbl1-4 | GTTTTGCCGTACCCACCCA | tatcttgtggaaaggacgaaacaccGGTTTTGCCGTACCCACCCAgtttaagagctatgctggaaacagc |
| Ruvbl1 | Ruvbl1-5 | AGTACGTCCCTTTGCCAAA | tatcttgtggaaaggacgaaacaccGAGTACGTCCCTTTGCCAAAgtttaagagctatgctggaaacagc |
| S100a2 | S100a2-1 | GGAGCTGCTGAGCTATGTG | tatcttgtggaaaggacgaaacaccGGGAGCTGCTGAGCTATGTGgtttaagagctatgctggaaacagc |
| S100a2 | S100a2-2 | GATAACGTAGATGATGAGA | tatcttgtggaaaggacgaaacaccGGATAACGTAGATGATGAGAgtttaagagctatgctggaaacagc |
| S100a2 | S100a2-3 | GAAATGAACAACCTCGATA | tatcttgtggaaaggacgaaacaccGGAAATGAACAACCTCGATAgtttaagagctatgctggaaacagc |
| S100a7l2 | S100a7l2-3 | GAATTTCCAAATTTTCTCC | tatcttgtggaaaggacgaaacaccGGAATTTCCAAATTTTCTCCgtttaagagctatgctggaaacagc |
| S100a7l2 | S100a7l2-4 | GAGAAGGAATTCAGAAAGT | tatcttgtggaaaggacgaaacaccGGAGAAGGAATTCAGAAAGTgtttaagagctatgctggaaacagc |
| S100a7l2 | S100a7l2-5 | CTGGGAACGACAGTCATAC | tatcttgtggaaaggacgaaacaccGCTGGGAACGACAGTCATACgtttaagagctatgctggaaacagc |
| Sae1 | Sae1-1 | ATTCGCAAAGGTGTACCCA | tatcttgtggaaaggacgaaacaccGATTCGCAAAGGTGTACCCAgtttaagagctatgctggaaacagc |
| Sae1 | Sae1-2 | TGGGTCTGTGGGCCGCAAT | tatcttgtggaaaggacgaaacaccGTGGGTCTGTGGGCCGCAATgtttaagagctatgctggaaacagc |
| Sae1 | Sae1-5 | ATGCCAACAATCAGCACCC | tatcttgtggaaaggacgaaacaccGATGCCAACAATCAGCACCCgtttaagagctatgctggaaacagc |
| Samsn1 | Samsn1-1 | ACCGACTCCCTTAAAATCA | tatcttgtggaaaggacgaaacaccGACCGACTCCCTTAAAATCAgtttaagagctatgctggaaacagc |
| Samsn1 | Samsn1-3 | AATCGGGACAGCTTCCGAC | tatcttgtggaaaggacgaaacaccGAATCGGGACAGCTTCCGACgtttaagagctatgctggaaacagc |
| Samsn1 | Samsn1-5 | TGGGATCACTGTTCCGATA | tatcttgtggaaaggacgaaacaccGTGGGATCACTGTTCCGATAgtttaagagctatgctggaaacagc |
| Sars | Sars-2 | TTGGATAAGCGCCTGCTCC | tatcttgtggaaaggacgaaacaccGTTGGATAAGCGCCTGCTCCgtttaagagctatgctggaaacagc |
| Sars | Sars-3 | TATGGTCCTTAGGGGCCCC | tatcttgtggaaaggacgaaacaccGTATGGTCCTTAGGGGCCCCgtttaagagctatgctggaaacagc |
| Sars | Sars-5 | TTGAGAACCTCCGCGAGAT | tatcttgtggaaaggacgaaacaccGTTGAGAACCTCCGCGAGATgtttaagagctatgctggaaacagc |
| Sart3 | Sart3-2 | TCGAGAACGCTCTGAGTGC | tatcttgtggaaaggacgaaacaccGTCGAGAACGCTCTGAGTGCgtttaagagctatgctggaaacagc |
| Sart3 | Sart3-3 | TCGACTTCGAGATGAAAAT | tatcttgtggaaaggacgaaacaccGTCGACTTCGAGATGAAAATgtttaagagctatgctggaaacagc |
| Sart3 | Sart3-4 | TGGTATTCCGCCAGCCGAG | tatcttgtggaaaggacgaaacaccGTGGTATTCCGCCAGCCGAGgtttaagagctatgctggaaacagc |
| Saxo2 | Saxo2-1 | TGAAAGTCATTCCGGTGAG | tatcttgtggaaaggacgaaacaccGTGAAAGTCATTCCGGTGAGgtttaagagctatgctggaaacagc |
| Saxo2 | Saxo2-4 | AGTTTCCGAATTTCACATC | tatcttgtggaaaggacgaaacaccGAGTTTCCGAATTTCACATCgtttaagagctatgctggaaacagc |
| Saxo2 | Saxo2-5 | ACAGCCTTTGATAAAAGTC | tatcttgtggaaaggacgaaacaccGACAGCCTTTGATAAAAGTCgtttaagagctatgctggaaacagc |
| Scgb1b30 | Scgb1b30-1 | GGTCCCATCGAAAAATTTA | tatcttgtggaaaggacgaaacaccGGGTCCCATCGAAAAATTTAgtttaagagctatgctggaaacagc |
| Scgb1b30 | Scgb1b30-2 | GTCATTATTGTATTGTTTC | tatcttgtggaaaggacgaaacaccGGTCATTATTGTATTGTTTCgtttaagagctatgctggaaacagc |
| Scgb1b30 | Scgb1b30-3 | AATAATGACCCTCTCGTAC | tatcttgtggaaaggacgaaacaccGAATAATGACCCTCTCGTACgtttaagagctatgctggaaacagc |
| Sec13 | Sec13-2 | GCTGGGCCCCTCATGACTA | tatcttgtggaaaggacgaaacaccGGCTGGGCCCCTCATGACTAgtttaagagctatgctggaaacagc |
| Sec13 | Sec13-3 | GGCCTCCTGTTCCTATGAC | tatcttgtggaaaggacgaaacaccGGGCCTCCTGTTCCTATGACgtttaagagctatgctggaaacagc |
| Sec13 | Sec13-5 | ATTTTGACGGACCTATCCG | tatcttgtggaaaggacgaaacaccGATTTTGACGGACCTATCCGgtttaagagctatgctggaaacagc |
| Sectm1b | Sectm1b-1 | CAACTCACCATTAACAATC | tatcttgtggaaaggacgaaacaccGCAACTCACCATTAACAATCgtttaagagctatgctggaaacagc |
| Sectm1b | Sectm1b-3 | TGGTGATCACAGATGCTCA | tatcttgtggaaaggacgaaacaccGTGGTGATCACAGATGCTCAgtttaagagctatgctggaaacagc |
| Sectm1b | Sectm1b-4 | GGCAGCTTCATATTCAAGG | tatcttgtggaaaggacgaaacaccGGGCAGCTTCATATTCAAGGgtttaagagctatgctggaaacagc |
| Seh1l | Seh1l-1 | TAAGAGCGAGAGCGGAGAC | tatcttgtggaaaggacgaaacaccGTAAGAGCGAGAGCGGAGACgtttaagagctatgctggaaacagc |
| Seh1l | Seh1l-2 | CATTGTACAGCTAGCTGGA | tatcttgtggaaaggacgaaacaccGCATTGTACAGCTAGCTGGAgtttaagagctatgctggaaacagc |
| Seh1l | Seh1l-4 | TAGAGTTGTCCTCTTCACC | tatcttgtggaaaggacgaaacaccGTAGAGTTGTCCTCTTCACCgtttaagagctatgctggaaacagc |
| Selenos | Selenos-1 | ACTCTACATTGTCATCCAG | tatcttgtggaaaggacgaaacaccGACTCTACATTGTCATCCAGgtttaagagctatgctggaaacagc |
| Selenos | Selenos-2 | CAGTCGAAGGGAGAGCCTC | tatcttgtggaaaggacgaaacaccGCAGTCGAAGGGAGAGCCTCgtttaagagctatgctggaaacagc |
| Selenos | Selenos-5 | GCCCAAGTTGAAAAACATA | tatcttgtggaaaggacgaaacaccGGCCCAAGTTGAAAAACATAgtttaagagctatgctggaaacagc |
| Semp2l2b | Semp2l2b-1 | GCTTGGAGCCTGATTTACC | tatcttgtggaaaggacgaaacaccGGCTTGGAGCCTGATTTACCgtttaagagctatgctggaaacagc |

| | | | |
|---|---|---|---|
| Semp2l2b | Semp2l2b-4 | GATCCTTCGGTCAACTCTA | tatcttgtggaaaggacgaaacaccGGATCCTTCGGTCAACTCTAgtttaagagctatgctggaaacagc |
| Semp2l2b | Semp2l2b-5 | TCGGGAGCTCTTGATCTCG | tatcttgtggaaaggacgaaacaccGTCGGGAGCTCTTGATCTCGgtttaagagctatgctggaaacagc |
| Serpina3i | Serpina3i-2 | TTTGGCCTGGCAAGGCTGC | tatcttgtggaaaggacgaaacaccGTTTGGCCTGGCAAGGCTGCgtttaagagctatgctggaaacagc |
| Serpina3i | Serpina3i-3 | CTGGATAAGAGCACGTTGA | tatcttgtggaaaggacgaaacaccGCTGGATAAGAGCACGTTGAgtttaagagctatgctggaaacagc |
| Serpina3i | Serpina3i-4 | TTGGAATTAAATGTATCAC | tatcttgtggaaaggacgaaacaccGTTGGAATTAAATGTATCACgtttaagagctatgctggaaacagc |
| Sgo2b | Sgo2b-2 | CCCAAATACATGACAATGA | tatcttgtggaaaggacgaaacaccGCCCAAATACATGACAATGAgtttaagagctatgctggaaacagc |
| Sgo2b | Sgo2b-3 | CACAAATGTCTGTCGAGAA | tatcttgtggaaaggacgaaacaccGCACAAATGTCTGTCGAGAAgtttaagagctatgctggaaacagc |
| Sgo2b | Sgo2b-4 | GCTCCATGCGCTTTACATA | tatcttgtggaaaggacgaaacaccGGCTCCATGCGCTTTACATAgtttaagagctatgctggaaacagc |
| Sh3glb1 | Sh3glb1-2 | GAAATGCCCTTATTAAATG | tatcttgtggaaaggacgaaacaccGGAAATGCCCTTATTAAATGgtttaagagctatgctggaaacagc |
| Sh3glb1 | Sh3glb1-4 | CCGGGTTGTTTATACGACT | tatcttgtggaaaggacgaaacaccGCCGGGTTGTTTATACGACTgtttaagagctatgctggaaacagc |
| Sh3glb1 | Sh3glb1-5 | GAATTTGTTTATGAGAAAC | tatcttgtggaaaggacgaaacaccGGAATTTGTTTATGAGAAACgtttaagagctatgctggaaacagc |
| Shb | Shb-1 | GCCGCCTGGACTACTGCGG | tatcttgtggaaaggacgaaacaccGGCCGCCTGGACTACTGCGGgtttaagagctatgctggaaacagc |
| Shb | Shb-3 | CTTCGAGGACCCGTACAAC | tatcttgtggaaaggacgaaacaccGCTTCGAGGACCCGTACAACgtttaagagctatgctggaaacagc |
| Shb | Shb-4 | CGCTCCTTCTGCGCGCGGT | tatcttgtggaaaggacgaaacaccGCGCTCCTTCTGCGCGCGGTgtttaagagctatgctggaaacagc |
| Shld2 | Shld2-1 | TAACTACAATCGTTGCTAA | tatcttgtggaaaggacgaaacaccGTAACTACAATCGTTGCTAAgtttaagagctatgctggaaacagc |
| Shld2 | Shld2-2 | CCATCCAAGTACCGCTAAC | tatcttgtggaaaggacgaaacaccGCCATCCAAGTACCGCTAACgtttaagagctatgctggaaacagc |
| Shld2 | Shld2-5 | TTTGGAAGTTCCGAACGCA | tatcttgtggaaaggacgaaacaccGTTTGGAAGTTCCGAACGCAgtttaagagctatgctggaaacagc |
| Sin3a | Sin3a-2 | GAGCCCAGTCGTTTACAAG | tatcttgtggaaaggacgaaacaccGGAGCCCAGTCGTTTACAAGgtttaagagctatgctggaaacagc |
| Sin3a | Sin3a-3 | TGCTGGTAACTTTTCGGTA | tatcttgtggaaaggacgaaacaccGTGCTGGTAACTTTTCGGTAgtttaagagctatgctggaaacagc |
| Sin3a | Sin3a-5 | ACCAATCTTGCAACAATCC | tatcttgtggaaaggacgaaacaccGACCAATCTTGCAACAATCCgtttaagagctatgctggaaacagc |
| Sipa1l1 | Sipa1l1-2 | TTGCGCAATTTACGAAAAA | tatcttgtggaaaggacgaaacaccGTTGCGCAATTTACGAAAAAgtttaagagctatgctggaaacagc |
| Sipa1l1 | Sipa1l1-3 | CGTATTTTCGGAATGAGAT | tatcttgtggaaaggacgaaacaccGCGTATTTTCGGAATGAGATgtttaagagctatgctggaaacagc |
| Sipa1l1 | Sipa1l1-5 | AGGTACACCGTGGAACACG | tatcttgtggaaaggacgaaacaccGAGGTACACCGTGGAACACGgtttaagagctatgctggaaacagc |
| Sis | Sis-2 | GAAATTTACCGGGAGTAAA | tatcttgtggaaaggacgaaacaccGGAAATTTACCGGGAGTAAAgtttaagagctatgctggaaacagc |
| Sis | Sis-4 | AATAATATCGTACATCACC | tatcttgtggaaaggacgaaacaccGAATAATATCGTACATCACCgtttaagagctatgctggaaacagc |
| Sis | Sis-5 | TCTCGTCTCGAGTAAAGAT | tatcttgtggaaaggacgaaacaccGTCTCGTCTCGAGTAAAGATgtttaagagctatgctggaaacagc |
| Skint1 | Skint1-1 | CTAAGCCGTTCACTATAAA | tatcttgtggaaaggacgaaacaccGCTAAGCCGTTCACTATAAAgtttaagagctatgctggaaacagc |
| Skint1 | Skint1-2 | TACAGAACTCTTAAAAGAT | tatcttgtggaaaggacgaaacaccGTACAGAACTCTTAAAAGATgtttaagagctatgctggaaacagc |
| Skint1 | Skint1-3 | GTACCTGTAAGTTTATGGC | tatcttgtggaaaggacgaaacaccGGTACCTGTAAGTTTATGGCgtttaagagctatgctggaaacagc |
| Skint3 | Skint3-2 | CCACAAAATGCACAACAAA | tatcttgtggaaaggacgaaacaccGCCACAAAATGCACAACAAAgtttaagagctatgctggaaacagc |
| Skint3 | Skint3-3 | TGGAGAGATAGCAATGGAC | tatcttgtggaaaggacgaaacaccGTGGAGAGATAGCAATGGACgtttaagagctatgctggaaacagc |
| Skint3 | Skint3-5 | AATGTAGCAAGTGACAATC | tatcttgtggaaaggacgaaacaccGAATGTAGCAAGTGACAATCgtttaagagctatgctggaaacagc |
| Slc39a13 | Slc39a13-1 | TTGCCCAGGAGTCCACCTA | tatcttgtggaaaggacgaaacaccGTTGCCCAGGAGTCCACCTAgtttaagagctatgctggaaacagc |
| Slc39a13 | Slc39a13-3 | ACCATGAGAGAGCCCAAGA | tatcttgtggaaaggacgaaacaccGACCATGAGAGAGCCCAAGAgtttaagagctatgctggaaacagc |
| Slc39a13 | Slc39a13-4 | TAATAAAGAAAGCGAGTCC | tatcttgtggaaaggacgaaacaccGTAATAAAGAAAGCGAGTCCgtttaagagctatgctggaaacagc |
| Slco1a1 | Slco1a1-2 | CACGTGTTTATGTAGACAT | tatcttgtggaaaggacgaaacaccGCACGTGTTTATGTAGACATgtttaagagctatgctggaaacagc |
| Slco1a1 | Slco1a1-4 | ATTCGTGAGTTATTTTGGA | tatcttgtggaaaggacgaaacaccGATTCGTGAGTTATTTTGGAgtttaagagctatgctggaaacagc |
| Slco1a1 | Slco1a1-5 | TGTAGTTGGATTTATCACT | tatcttgtggaaaggacgaaacaccGTGTAGTTGGATTTATCACTgtttaagagctatgctggaaacagc |
| Slpi | Slpi-1 | GGGAGGTGCGTCAAAACTC | tatcttgtggaaaggacgaaacaccGGGGAGGTGCGTCAAAACTCgtttaagagctatgctggaaacagc |
| Slpi | Slpi-3 | CCACAATGCCGTACTGACT | tatcttgtggaaaggacgaaacaccGCCACAATGCCGTACTGACTgtttaagagctatgctggaaacagc |
| Slpi | Slpi-5 | CTTTAGATGCTATCAAAAT | tatcttgtggaaaggacgaaacaccGCTTTAGATGCTATCAAAATgtttaagagctatgctggaaacagc |
| Smarcad1 | Smarcad1-2 | AATCCTAAGAAAGCCGTAG | tatcttgtggaaaggacgaaacaccGAATCCTAAGAAAGCCGTAGgtttaagagctatgctggaaacagc |
| Smarcad1 | Smarcad1-3 | TTCCCAATTATTAAAGGGC | tatcttgtggaaaggacgaaacaccGTTCCCAATTATTAAAGGGCgtttaagagctatgctggaaacagc |
| Smarcad1 | Smarcad1-4 | TTACGATGCTCACCGGGAA | tatcttgtggaaaggacgaaacaccGTTACGATGCTCACCGGGAAgtttaagagctatgctggaaacagc |
| Smlr1 | Smlr1-1 | CTTTCAGTTGAAGAAGAAC | tatcttgtggaaaggacgaaacaccGCTTTCAGTTGAAGAAGAACgtttaagagctatgctggaaacagc |
| Smlr1 | Smlr1-2 | ATCAGTTTGATCCTGAAGT | tatcttgtggaaaggacgaaacaccGATCAGTTTGATCCTGAAGTgtttaagagctatgctggaaacagc |
| Smlr1 | Smlr1-3 | CCTCCTTCTTGCCTACTTC | tatcttgtggaaaggacgaaacaccGCCTCCTTCTTGCCTACTTCgtttaagagctatgctggaaacagc |
| Snapc1 | Snapc1-1 | GGAAGCTTTAGCTTTGGCG | tatcttgtggaaaggacgaaacaccGGGAAGCTTTAGCTTTGGCGgtttaagagctatgctggaaacagc |
| Snapc1 | Snapc1-2 | CCAGTCCTTCAAGGCAACT | tatcttgtggaaaggacgaaacaccGCCAGTCCTTCAAGGCAACTgtttaagagctatgctggaaacagc |
| Snapc1 | Snapc1-4 | CTTGGGCATAGCTGTGAAG | tatcttgtggaaaggacgaaacaccGCTTGGGCATAGCTGTGAAGgtttaagagctatgctggaaacagc |
| Snapc4 | Snapc4-1 | CTGAATTGCCCCATCGGTC | tatcttgtggaaaggacgaaacaccGCTGAATTGCCCCATCGGTCgtttaagagctatgctggaaacagc |
| Snapc4 | Snapc4-2 | ATTGGGCATCGCTCCTACC | tatcttgtggaaaggacgaaacaccGATTGGGCATCGCTCCTACCgtttaagagctatgctggaaacagc |
| Snapc4 | Snapc4-4 | TTCCATGAAGTAGACGACT | tatcttgtggaaaggacgaaacaccGTTCCATGAAGTAGACGACTgtttaagagctatgctggaaacagc |
| Snrnp200 | Snrnp200-1 | CCTTTACGCTGGCGTCGGA | tatcttgtggaaaggacgaaacaccGCCTTTACGCTGGCGTCGGAgtttaagagctatgctggaaacagc |
| Snrnp200 | Snrnp200-2 | TGCGAGTCGACCCTGCTAA | tatcttgtggaaaggacgaaacaccGTGCGAGTCGACCCTGCTAAgtttaagagctatgctggaaacagc |
| Snrnp200 | Snrnp200-3 | CGGCCGCAGTATGACACCA | tatcttgtggaaaggacgaaacaccGCGGCCGCAGTATGACACCAgtttaagagctatgctggaaacagc |
| Snrnp35 | Snrnp35-1 | TGGTTCGGGACCTGGTGAC | tatcttgtggaaaggacgaaacaccGTGGTTCGGGACCTGGTGACgtttaagagctatgctggaaacagc |
| Snrnp35 | Snrnp35-2 | TACGCCTTCATTGAGTACA | tatcttgtggaaaggacgaaacaccGTACGCCTTCATTGAGTACAgtttaagagctatgctggaaacagc |
| Snrnp35 | Snrnp35-5 | GCTGGATCCCTCGGCGGCT | tatcttgtggaaaggacgaaacaccGGCTGGATCCCTCGGCGGCTgtttaagagctatgctggaaacagc |
| Snrpb | Snrpb-1 | AATACCTCACCTGTTGGGA | tatcttgtggaaaggacgaaacaccGAATACCTCACCTGTTGGGAgtttaagagctatgctggaaacagc |
| Snrpb | Snrpb-2 | GGGCATAGGAACACCAGCC | tatcttgtggaaaggacgaaacaccGGGGCATAGGAACACCAGCCgtttaagagctatgctggaaacagc |
| Snrpb | Snrpb-4 | TTCTACTGTCATGGAGACC | tatcttgtggaaaggacgaaacaccGTTCTACTGTCATGGAGACCgtttaagagctatgctggaaacagc |
| Snrpc | Snrpc-3 | TTGAAACTTACGGAGACTG | tatcttgtggaaaggacgaaacaccGTTGAAACTTACGGAGACTGgtttaagagctatgctggaaacagc |
| Snrpc | Snrpc-4 | TAGCGGGCCCTCCTCGGCC | tatcttgtggaaaggacgaaacaccGTAGCGGGCCCTCCTCGGCCgtttaagagctatgctggaaacagc |
| Snrpc | Snrpc-5 | TGATGCCTGCCCCCCACAT | tatcttgtggaaaggacgaaacaccGTGATGCCTGCCCCCCACATgtttaagagctatgctggaaacagc |
| Sox2 | Sox2-1 | GGATTATAAATACCGGCCG | tatcttgtggaaaggacgaaacaccGGGATTATAAATACCGGCCGgtttaagagctatgctggaaacagc |
| Sox2 | Sox2-4 | GCGGGCGTGAACCAGCGCA | tatcttgtggaaaggacgaaacaccGGCGGGCGTGAACCAGCGCAgtttaagagctatgctggaaacagc |
| Sox2 | Sox2-5 | GGCCTCAACGCTCACGGCG | tatcttgtggaaaggacgaaacaccGGGCCTCAACGCTCACGGCGgtttaagagctatgctggaaacagc |
| Spata2 | Spata2-2 | ATCTGCTCACACTCGACCT | tatcttgtggaaaggacgaaacaccGATCTGCTCACACTCGACCTgtttaagagctatgctggaaacagc |
| Spata2 | Spata2-3 | TGTACACAGTCCCCAACTC | tatcttgtggaaaggacgaaacaccGTGTACACAGTCCCCAACTCgtttaagagctatgctggaaacagc |
| Spata2 | Spata2-4 | CTTGACATAGTAAACAAAA | tatcttgtggaaaggacgaaacaccGCTTGACATAGTAAACAAAAgtttaagagctatgctggaaacagc |
| Spata22 | Spata22-2 | TACTTCTGAGATTGGGCGT | tatcttgtggaaaggacgaaacaccGTACTTCTGAGATTGGGCGTgtttaagagctatgctggaaacagc |
| Spata22 | Spata22-4 | GTTGAGCTCCCAAATTCAC | tatcttgtggaaaggacgaaacaccGGTTGAGCTCCCAAATTCACgtttaagagctatgctggaaacagc |
| Spata22 | Spata22-5 | TTGATAAACATGTTTGTCG | tatcttgtggaaaggacgaaacaccGTTGATAAACATGTTTGTCGgtttaagagctatgctggaaacagc |
| Spc24 | Spc24-1 | TGCAGAACCTACACTGCAG | tatcttgtggaaaggacgaaacaccGTGCAGAACCTACACTGCAGgtttaagagctatgctggaaacagc |
| Spc24 | Spc24-4 | TGTCCCATTGCAGAGACTC | tatcttgtggaaaggacgaaacaccGTGTCCCATTGCAGAGACTCgtttaagagctatgctggaaacagc |

| | | | |
|---|---|---|---|
| Spc24 | Spc24-5 | GATGCAGGACGGCGCCTAC | tatcttgtggaaaggacgaaacaccGGATGCAGGACGGCGCCTACgtttaagagctatgctggaaacagc |
| Spcs2 | Spcs2-1 | CGAACCTATTTTGTGATGA | tatcttgtggaaaggacgaaacaccGCGAACCTATTTTGTGATGAgtttaagagctatgctggaaacagc |
| Spcs2 | Spcs2-3 | TCCCAAATCAAAGCCACGA | tatcttgtggaaaggacgaaacaccGTCCCAAATCAAAGCCACGAgtttaagagctatgctggaaacagc |
| Spcs2 | Spcs2-4 | GCCTGTAAAAATTGATAAG | tatcttgtggaaaggacgaaacaccGGCCTGTAAAAATTGATAAGgtttaagagctatgctggaaacagc |
| Spdl1 | Spdl1-2 | CAGGTGGAGGATCGAAGGG | tatcttgtggaaaggacgaaacaccGCAGGTGGAGGATCGAAGGGgtttaagagctatgctggaaacagc |
| Spdl1 | Spdl1-4 | AATTGGATGAAGCAAGGCT | tatcttgtggaaaggacgaaacaccGAATTGGATGAAGCAAGGCTgtttaagagctatgctggaaacagc |
| Spdl1 | Spdl1-5 | TTCATGACATTTATCCAGT | tatcttgtggaaaggacgaaacaccGTTCATGACATTTATCCAGTgtttaagagctatgctggaaacagc |
| Spinkl | Spinkl-1 | TTCTATCATCTGCACAAAC | tatcttgtggaaaggacgaaacaccGTTCTATCATCTGCACAAACgtttaagagctatgctggaaacagc |
| Spinkl | Spinkl-3 | TCAGATTTACTCTTATACA | tatcttgtggaaaggacgaaacaccGTCAGATTTACTCTTATACAgtttaagagctatgctggaaacagc |
| Spinkl | Spinkl-4 | CCAGAGAATGTTATTAAAG | tatcttgtggaaaggacgaaacaccGCCAGAGAATGTTATTAAAGgtttaagagctatgctggaaacagc |
| Spo11 | Spo11-2 | ACACTGAGGTATCATCCGA | tatcttgtggaaaggacgaaacaccGACACTGAGGTATCATCCGAgtttaagagctatgctggaaacagc |
| Spo11 | Spo11-4 | ATACAGGAAATGTCATCGA | tatcttgtggaaaggacgaaacaccGATACAGGAAATGTCATCGAgtttaagagctatgctggaaacagc |
| Spo11 | Spo11-5 | CTACAGGTACACTGGACTC | tatcttgtggaaaggacgaaacaccGCTACAGGTACACTGGACTCgtttaagagctatgctggaaacagc |
| Srp19 | Srp19-1 | ATCCGTCTCCCCTCAGCTA | tatcttgtggaaaggacgaaacaccGATCCGTCTCCCCTCAGCTAgtttaagagctatgctggaaacagc |
| Srp19 | Srp19-4 | GTCCGGGTACAGCTCAAAC | tatcttgtggaaaggacgaaacaccGGTCCGGGTACAGCTCAAACgtttaagagctatgctggaaacagc |
| Srp19 | Srp19-5 | TTTAAAAAGCTTACGTGAC | tatcttgtggaaaggacgaaacaccGTTTAAAAAGCTTACGTGACgtttaagagctatgctggaaacagc |
| Srsf1 | Srsf1-2 | TCGAACTCAACGAAGGCGA | tatcttgtggaaaggacgaaacaccGTCGAACTCAACGAAGGCGAgtttaagagctatgctggaaacagc |
| Srsf1 | Srsf1-3 | ATGCGGTGTACGGTCGCGA | tatcttgtggaaaggacgaaacaccGATGCGGTGTACGGTCGCGAgtttaagagctatgctggaaacagc |
| Srsf1 | Srsf1-4 | TACGACGGCTACCGGCTGC | tatcttgtggaaaggacgaaacaccGTACGACGGCTACCGGCTGCgtttaagagctatgctggaaacagc |
| Srsf2 | Srsf2-2 | TTGTCGTGGAACCGGACGA | tatcttgtggaaaggacgaaacaccGTTGTCGTGGAACCGGACGAgtttaagagctatgctggaaacagc |
| Srsf2 | Srsf2-3 | GCTACACCAAGGAGTCCCG | tatcttgtggaaaggacgaaacaccGGCTACACCAAGGAGTCCCGgtttaagagctatgctggaaacagc |
| Srsf2 | Srsf2-4 | GGCGACGTGTACATTCCGC | tatcttgtggaaaggacgaaacaccGGGCGACGTGTACATTCCGCgtttaagagctatgctggaaacagc |
| Stat3 | Stat3-1 | CCAGTTTACCACGAAAGTC | tatcttgtggaaaggacgaaacaccGCCAGTTTACCACGAAAGTCgtttaagagctatgctggaaacagc |
| Stat3 | Stat3-4 | ATCGCTTACTCTCCGCATC | tatcttgtggaaaggacgaaacaccGATCGCTTACTCTCCGCATCgtttaagagctatgctggaaacagc |
| Stat3 | Stat3-5 | TCGAAGGTTGTGCTGATAG | tatcttgtggaaaggacgaaacaccGTCGAAGGTTGTGCTGATAGgtttaagagctatgctggaaacagc |
| Stn1 | Stn1-3 | ACGCAGTTTATAACCCCAG | tatcttgtggaaaggacgaaacaccGACGCAGTTTATAACCCCAGgtttaagagctatgctggaaacagc |
| Stn1 | Stn1-4 | AGACTTTCTACAGCTACGG | tatcttgtggaaaggacgaaacaccGAGACTTTCTACAGCTACGGgtttaagagctatgctggaaacagc |
| Stn1 | Stn1-5 | AACGGGCATCCAATAAGGC | tatcttgtggaaaggacgaaacaccGAACGGGCATCCAATAAGGCgtttaagagctatgctggaaacagc |
| Strada | Strada-1 | AGGTAGACCTTCCCATCCG | tatcttgtggaaaggacgaaacaccGAGGTAGACCTTCCCATCCGgtttaagagctatgctggaaacagc |
| Strada | Strada-4 | TCATTATCTGCGATGAAGG | tatcttgtggaaaggacgaaacaccGTCATTATCTGCGATGAAGGgtttaagagctatgctggaaacagc |
| Strada | Strada-5 | ATACGGTACTATATTGGGA | tatcttgtggaaaggacgaaacaccGATACGGTACTATATTGGGAgtttaagagctatgctggaaacagc |
| Stxbp6 | Stxbp6-1 | ATTGGCAGTTAATAAACTC | tatcttgtggaaaggacgaaacaccGATTGGCAGTTAATAAACTCgtttaagagctatgctggaaacagc |
| Stxbp6 | Stxbp6-2 | CTAGCAACCCACTGGTCAA | tatcttgtggaaaggacgaaacaccGCTAGCAACCCACTGGTCAAgtttaagagctatgctggaaacagc |
| Stxbp6 | Stxbp6-3 | ATTCGTCCGAAGATCCCAA | tatcttgtggaaaggacgaaacaccGATTCGTCCGAAGATCCCAAgtttaagagctatgctggaaacagc |
| Sult1e1 | Sult1e1-1 | TTCTGATTTCAGTTCCGTA | tatcttgtggaaaggacgaaacaccGTTCTGATTTCAGTTCCGTAgtttaagagctatgctggaaacagc |
| Sult1e1 | Sult1e1-2 | AGAAATTTATGCAAGGGCA | tatcttgtggaaaggacgaaacaccGAGAAATTTATGCAAGGGCAgtttaagagctatgctggaaacagc |
| Sult1e1 | Sult1e1-5 | TTGCTACATATCCTAAATC | tatcttgtggaaaggacgaaacaccGTTGCTACATATCCTAAATCgtttaagagctatgctggaaacagc |
| Sult2a6 | Sult2a6-1 | GTTTGAACACATCCATGGC | tatcttgtggaaaggacgaaacaccGGTTTGAACACATCCATGGCgtttaagagctatgctggaaacagc |
| Sult2a6 | Sult2a6-2 | ATTTTGAACAATTCCTGAA | tatcttgtggaaaggacgaaacaccGATTTTGAACAATTCCTGAAgtttaagagctatgctggaaacagc |
| Sult2a6 | Sult2a6-5 | TAATTGGGAACGTTCACCC | tatcttgtggaaaggacgaaacaccGTAATTGGGAACGTTCACCCgtttaagagctatgctggaaacagc |
| Supt20 | Supt20-2 | GACAACCATAAGTGGACCC | tatcttgtggaaaggacgaaacaccGGACAACCATAAGTGGACCCgtttaagagctatgctggaaacagc |
| Supt20 | Supt20-4 | CAATAGGCTGCTCTATAAC | tatcttgtggaaaggacgaaacaccGCAATAGGCTGCTCTATAACgtttaagagctatgctggaaacagc |
| Supt20 | Supt20-5 | GAACACTCGCCCAATGAAA | tatcttgtggaaaggacgaaacaccGGAACACTCGCCCAATGAAAgtttaagagctatgctggaaacagc |
| Surf6 | Surf6-1 | CTGATCACGAAGCTCGTCC | tatcttgtggaaaggacgaaacaccGCTGATCACGAAGCTCGTCCgtttaagagctatgctggaaacagc |
| Surf6 | Surf6-3 | CACTCTCGAGAAAAGACAG | tatcttgtggaaaggacgaaacaccGCACTCTCGAGAAAAGACAGgtttaagagctatgctggaaacagc |
| Surf6 | Surf6-4 | AAAAGATCCAGCTGGCCCG | tatcttgtggaaaggacgaaacaccGAAAAGATCCAGCTGGCCCGgtttaagagctatgctggaaacagc |
| Susd6 | Susd6-1 | GTCTTTACAAGGCCGGGGG | tatcttgtggaaaggacgaaacaccGGTCTTTACAAGGCCGGGGGgtttaagagctatgctggaaacagc |
| Susd6 | Susd6-3 | AATACCTGACCTGCAAAAA | tatcttgtggaaaggacgaaacaccGAATACCTGACCTGCAAAAAgtttaagagctatgctggaaacagc |
| Susd6 | Susd6-4 | GACAGCTAACCTCCATGGC | tatcttgtggaaaggacgaaacaccGGACAGCTAACCTCCATGGCgtttaagagctatgctggaaacagc |
| Suz12 | Suz12-1 | CGGCTTCGGGCGGCAAATC | tatcttgtggaaaggacgaaacaccGCGGCTTCGGGCGGCAAATCgtttaagagctatgctggaaacagc |
| Suz12 | Suz12-3 | CGAGTAGGACTTCACCATA | tatcttgtggaaaggacgaaacaccGCGAGTAGGACTTCACCATAgtttaagagctatgctggaaacagc |
| Suz12 | Suz12-5 | CGTTTACAGCTTTTAGATG | tatcttgtggaaaggacgaaacaccGCGTTTACAGCTTTTAGATGgtttaagagctatgctggaaacagc |
| Sypl | Sypl-2 | GGTTACACGAACCTCTACC | tatcttgtggaaaggacgaaacaccGGGTTACACGAACCTCTACCgtttaagagctatgctggaaacagc |
| Sypl | Sypl-3 | TTTGCGACTATCCCGGTAG | tatcttgtggaaaggacgaaacaccGTTTGCGACTATCCCGGTAGgtttaagagctatgctggaaacagc |
| Sypl | Sypl-4 | AAACGGTTACTTACGATCA | tatcttgtggaaaggacgaaacaccGAAACGGTTACTTACGATCAgtttaagagctatgctggaaacagc |
| Szrd1 | Szrd1-1 | AAGTCCCTAGCACAGCGGG | tatcttgtggaaaggacgaaacaccGAAGTCCCTAGCACAGCGGGgtttaagagctatgctggaaacagc |
| Szrd1 | Szrd1-2 | TGACCACACCGTTGCTGGT | tatcttgtggaaaggacgaaacaccGTGACCACACCGTTGCTGGTgtttaagagctatgctggaaacagc |
| Szrd1 | Szrd1-3 | ACAGATCCGCATCCTCAAG | tatcttgtggaaaggacgaaacaccGACAGATCCGCATCCTCAAGgtttaagagctatgctggaaacagc |
| Tada3 | Tada3-3 | TCGGTGCTCCCCCCAAACA | tatcttgtggaaaggacgaaacaccGTCGGTGCTCCCCCCAAACAgtttaagagctatgctggaaacagc |
| Tada3 | Tada3-4 | TAGGTCGCGACCACGAGCT | tatcttgtggaaaggacgaaacaccGTAGGTCGCGACCACGAGCTgtttaagagctatgctggaaacagc |
| Tada3 | Tada3-5 | TGGCCCGCTCCGAGGACGA | tatcttgtggaaaggacgaaacaccGTGGCCCGCTCCGAGGACGAgtttaagagctatgctggaaacagc |
| Taf13 | Taf13-1 | CAGTGCGCTGTATGATGTA | tatcttgtggaaaggacgaaacaccGCAGTGCGCTGTATGATGTAgtttaagagctatgctggaaacagc |
| Taf13 | Taf13-4 | AGTGATGAACTCTATGACA | tatcttgtggaaaggacgaaacaccGAGTGATGAACTCTATGACAgtttaagagctatgctggaaacagc |
| Taf13 | Taf13-5 | CTCACAAGGCAATGTCGAT | tatcttgtggaaaggacgaaacaccGCTCACAAGGCAATGTCGATgtttaagagctatgctggaaacagc |
| Taf1b | Taf1b-1 | GATCAACTCCATCAACAGG | tatcttgtggaaaggacgaaacaccGGATCAACTCCATCAACAGGgtttaagagctatgctggaaacagc |
| Taf1b | Taf1b-2 | AGTTCTGGGCTTACTCCGA | tatcttgtggaaaggacgaaacaccGAGTTCTGGGCTTACTCCGAgtttaagagctatgctggaaacagc |
| Taf1b | Taf1b-5 | CCGTTATGTCTGGGAAACG | tatcttgtggaaaggacgaaacaccGCCGTTATGTCTGGGAAACGgtttaagagctatgctggaaacagc |
| Taf5l | Taf5l-1 | AGCTGTTCCGTGTTATAGA | tatcttgtggaaaggacgaaacaccGAGCTGTTCCGTGTTATAGAgtttaagagctatgctggaaacagc |
| Taf5l | Taf5l-2 | GTATCAAGCGGGTCAAGGA | tatcttgtggaaaggacgaaacaccGGTATCAAGCGGGTCAAGGAgtttaagagctatgctggaaacagc |
| Taf5l | Taf5l-4 | CACGGCAAGCACTTTGCAC | tatcttgtggaaaggacgaaacaccGCACGGCAAGCACTTTGCACgtttaagagctatgctggaaacagc |
| Tas2r131 | Tas2r131-1 | TGCTAGGGACACAAGAAAT | tatcttgtggaaaggacgaaacaccGTGCTAGGGACACAAGAAATgtttaagagctatgctggaaacagc |
| Tas2r131 | Tas2r131-3 | GAATAAGTAGAATATGCTC | tatcttgtggaaaggacgaaacaccGGAATAAGTAGAATATGCTCgtttaagagctatgctggaaacagc |
| Tas2r131 | Tas2r131-4 | AACTGATCAACTATCAACA | tatcttgtggaaaggacgaaacaccGAACTGATCAACTATCAACAgtttaagagctatgctggaaacagc |
| Tbl3 | Tbl3-1 | GGATCAGAGCATACGCATC | tatcttgtggaaaggacgaaacaccGGGATCAGAGCATACGCATCgtttaagagctatgctggaaacagc |
| Tbl3 | Tbl3-2 | GTTGGAAGCCACGACGATG | tatcttgtggaaaggacgaaacaccGGTTGGAAGCCACGACGATGgtttaagagctatgctggaaacagc |
| Tbl3 | Tbl3-5 | GCTGGCTGTTACGAGTACC | tatcttgtggaaaggacgaaacaccGGCTGGCTGTTACGAGTACCgtttaagagctatgctggaaacagc |

| | | | |
|---|---|---|---|
| Tbx19 | Tbx19-2 | CTGCATAAATATGAACCTC | tatcttgtggaaaggacgaaacaccGCTGCATAAATATGAACCTCgtttaagagctatgctggaaacagc |
| Tbx19 | Tbx19-3 | CAACAAGCTCAATGGAGGT | tatcttgtggaaaggacgaaacaccGCAACAAGCTCAATGGAGGTgtttaagagctatgctggaaacagc |
| Tbx19 | Tbx19-5 | GGTCCGGACGAAGTCCAGT | tatcttgtggaaaggacgaaacaccGGGTCCGGACGAAGTCCAGTgtttaagagctatgctggaaacagc |
| Tceal7 | Tceal7-1 | AACCCTGCGGAGCACTTGA | tatcttgtggaaaggacgaaacaccGAACCCTGCGGAGCACTTGAgtttaagagctatgctggaaacagc |
| Tceal7 | Tceal7-3 | TAGACTATAGGCATTTTAA | tatcttgtggaaaggacgaaacaccGTAGACTATAGGCATTTTAAgtttaagagctatgctggaaacagc |
| Tceal7 | Tceal7-4 | TTAAAGGTGAAGAAATGAC | tatcttgtggaaaggacgaaacaccGTTAAAGGTGAAGAAATGACgtttaagagctatgctggaaacagc |
| Tceanc | Tceanc-1 | AATGCACTCGCAAGAGCTC | tatcttgtggaaaggacgaaacaccGAATGCACTCGCAAGAGCTCgtttaagagctatgctggaaacagc |
| Tceanc | Tceanc-2 | CTCAAATGGAAATTAATGA | tatcttgtggaaaggacgaaacaccGCTCAAATGGAAATTAATGAgtttaagagctatgctggaaacagc |
| Tceanc | Tceanc-3 | TCTAGCCATAGTGAGATCA | tatcttgtggaaaggacgaaacaccGTCTAGCCATAGTGAGATCAgtttaagagctatgctggaaacagc |
| Tcerg1 | Tcerg1-1 | CACCAAATGCCGTGATGCG | tatcttgtggaaaggacgaaacaccGCACCAAATGCCGTGATGCGgtttaagagctatgctggaaacagc |
| Tcerg1 | Tcerg1-2 | ATGGGCCCACCTCACCTAC | tatcttgtggaaaggacgaaacaccGATGGGCCCACCTCACCTACgtttaagagctatgctggaaacagc |
| Tcerg1 | Tcerg1-5 | GTAGATGTCGGAGTTGACG | tatcttgtggaaaggacgaaacaccGGTAGATGTCGGAGTTGACGgtttaagagctatgctggaaacagc |
| Teddm1a | Teddm1a-1 | TCATTGAATATCACAGCTC | tatcttgtggaaaggacgaaacaccGTCATTGAATATCACAGCTCgtttaagagctatgctggaaacagc |
| Teddm1a | Teddm1a-2 | CCTACCTTTCCAAGAATAA | tatcttgtggaaaggacgaaacaccGCCTACCTTTCCAAGAATAAgtttaagagctatgctggaaacagc |
| Teddm1a | Teddm1a-4 | ATCTCATAAACGATCAAGA | tatcttgtggaaaggacgaaacaccGATCTCATAAACGATCAAGAgtttaagagctatgctggaaacagc |
| Telo2 | Telo2-2 | CGGCGGCCACTCCTCATAC | tatcttgtggaaaggacgaaacaccGCGGCGGCCACTCCTCATACgtttaagagctatgctggaaacagc |
| Telo2 | Telo2-4 | CCAAGAACTCGAGATACGA | tatcttgtggaaaggacgaaacaccGCCAAGAACTCGAGATACGAgtttaagagctatgctggaaacagc |
| Telo2 | Telo2-5 | CTCTGAAACGATACCTCGG | tatcttgtggaaaggacgaaacaccGCTCTGAAACGATACCTCGGgtttaagagctatgctggaaacagc |
| Terb1 | Terb1-2 | TTAGCATCATACTTGCCAT | tatcttgtggaaaggacgaaacaccGTTAGCATCATACTTGCCATgtttaagagctatgctggaaacagc |
| Terb1 | Terb1-3 | ATATAGGTCGAATTACCTC | tatcttgtggaaaggacgaaacaccGATATAGGTCGAATTACCTCgtttaagagctatgctggaaacagc |
| Terb1 | Terb1-4 | GTGTCAACAATCCTCAAAA | tatcttgtggaaaggacgaaacaccGGTGTCAACAATCCTCAAAAgtttaagagctatgctggaaacagc |
| Tex29 | Tex29-2 | GAACACCTGTACATAAACT | tatcttgtggaaaggacgaaacaccGGAACACCTGTACATAAACTgtttaagagctatgctggaaacagc |
| Tex29 | Tex29-4 | CATTGCCGTCATTTATAGG | tatcttgtggaaaggacgaaacaccGCATTGCCGTCATTTATAGGgtttaagagctatgctggaaacagc |
| Tex29 | Tex29-5 | GTGGGGGAGGTTCTCGAAT | tatcttgtggaaaggacgaaacaccGGTGGGGGAGGTTCTCGAATgtttaagagctatgctggaaacagc |
| Tfpi | Tfpi-1 | GTGAACGATTCGTGTACGG | tatcttgtggaaaggacgaaacaccGGTGAACGATTCGTGTACGGgtttaagagctatgctggaaacagc |
| Tfpi | Tfpi-3 | GTAAGAAAACATGCATACC | tatcttgtggaaaggacgaaacaccGGTAAGAAAACATGCATACCgtttaagagctatgctggaaacagc |
| Tfpi | Tfpi-4 | TATATACGGGGGATGTGAA | tatcttgtggaaaggacgaaacaccGTATATACGGGGGATGTGAAgtttaagagctatgctggaaacagc |
| Tgif2 | Tgif2-1 | CTCTTTCTAGATATGTAAC | tatcttgtggaaaggacgaaacaccGCTCTTTCTAGATATGTAACgtttaagagctatgctggaaacagc |
| Tgif2 | Tgif2-3 | AAGGAGAGCATGTTAGTGG | tatcttgtggaaaggacgaaacaccGAAGGAGAGCATGTTAGTGGgtttaagagctatgctggaaacagc |
| Tgif2 | Tgif2-4 | GCAGCACCCTTCCCACAAG | tatcttgtggaaaggacgaaacaccGGCAGCACCCTTCCCACAAGgtttaagagctatgctggaaacagc |
| Thg1l | Thg1l-1 | TACGTAACTGGAGGCAAAC | tatcttgtggaaaggacgaaacaccGTACGTAACTGGAGGCAAACgtttaagagctatgctggaaacagc |
| Thg1l | Thg1l-2 | AGGACATTGTGATTGCGTA | tatcttgtggaaaggacgaaacaccGAGGACATTGTGATTGCGTAgtttaagagctatgctggaaacagc |
| Thg1l | Thg1l-5 | GGACGGCCGGAATTTCCAC | tatcttgtggaaaggacgaaacaccGGGACGGCCGGAATTTCCACgtttaagagctatgctggaaacagc |
| Thoc6 | Thoc6-3 | ATTTCGGGTACTTCCAGAC | tatcttgtggaaaggacgaaacaccGATTTCGGGTACTTCCAGACgtttaagagctatgctggaaacagc |
| Thoc6 | Thoc6-4 | CGATGGAGAGGTCAAGGGT | tatcttgtggaaaggacgaaacaccGCGATGGAGAGGTCAAGGGTgtttaagagctatgctggaaacagc |
| Thoc6 | Thoc6-5 | TGCCAGAAACTTCCCGCAG | tatcttgtggaaaggacgaaacaccGTGCCAGAAACTTCCCGCAGgtttaagagctatgctggaaacagc |
| Thumpd2 | Thumpd2-2 | AATGTCAGAAAGGTCATCA | tatcttgtggaaaggacgaaacaccGAATGTCAGAAAGGTCATCAgtttaagagctatgctggaaacagc |
| Thumpd2 | Thumpd2-4 | ACCCACTAAAACGGAAAGC | tatcttgtggaaaggacgaaacaccGACCCACTAAAACGGAAAGCgtttaagagctatgctggaaacagc |
| Thumpd2 | Thumpd2-5 | ATAGAGGGCAAATCTGAAC | tatcttgtggaaaggacgaaacaccGATAGAGGGCAAATCTGAACgtttaagagctatgctggaaacagc |
| Timmdc1 | Timmdc1-2 | GTTATGATAAATCTCCGCT | tatcttgtggaaaggacgaaacaccGGTTATGATAAATCTCCGCTgtttaagagctatgctggaaacagc |
| Timmdc1 | Timmdc1-3 | GAAATTGACTATATCTACA | tatcttgtggaaaggacgaaacaccGGAAATTGACTATATCTACAgtttaagagctatgctggaaacagc |
| Timmdc1 | Timmdc1-5 | TCCGGATTCCAGGGAACCG | tatcttgtggaaaggacgaaacaccGTCCGGATTCCAGGGAACCGgtttaagagctatgctggaaacagc |
| Tiparp | Tiparp-2 | CAGTGGCAGATTCTACAAC | tatcttgtggaaaggacgaaacaccGCAGTGGCAGATTCTACAACgtttaagagctatgctggaaacagc |
| Tiparp | Tiparp-3 | TAACTTTATCTACGCAGTC | tatcttgtggaaaggacgaaacaccGTAACTTTATCTACGCAGTCgtttaagagctatgctggaaacagc |
| Tiparp | Tiparp-4 | TCAGTACTCAACTTATCAC | tatcttgtggaaaggacgaaacaccGTCAGTACTCAACTTATCACgtttaagagctatgctggaaacagc |
| Tipin | Tipin-2 | GGTCTGTTTACTAGGATCA | tatcttgtggaaaggacgaaacaccGGGTCTGTTTACTAGGATCAgtttaagagctatgctggaaacagc |
| Tipin | Tipin-3 | ACCTAAGCTAGATGCTACG | tatcttgtggaaaggacgaaacaccGACCTAAGCTAGATGCTACGgtttaagagctatgctggaaacagc |
| Tipin | Tipin-5 | AGTCGAATTCGTTTTAAAC | tatcttgtggaaaggacgaaacaccGAGTCGAATTCGTTTTAAACgtttaagagctatgctggaaacagc |
| Tiprl | Tiprl-1 | ATGCAATTCATCTTCAAAC | tatcttgtggaaaggacgaaacaccGATGCAATTCATCTTCAAACgtttaagagctatgctggaaacagc |
| Tiprl | Tiprl-2 | AGTTATTAAACCATATGAC | tatcttgtggaaaggacgaaacaccGAGTTATTAAACCATATGACgtttaagagctatgctggaaacagc |
| Tiprl | Tiprl-3 | ACGTTCTAAGGATCCAGCA | tatcttgtggaaaggacgaaacaccGACGTTCTAAGGATCCAGCAgtttaagagctatgctggaaacagc |
| Tlr6 | Tlr6-1 | GATAACTGAGAGAATCGAC | tatcttgtggaaaggacgaaacaccGGATAACTGAGAGAATCGACgtttaagagctatgctggaaacagc |
| Tlr6 | Tlr6-2 | ATTGAGGTACTCCACCGGT | tatcttgtggaaaggacgaaacaccGATTGAGGTACTCCACCGGTgtttaagagctatgctggaaacagc |
| Tlr6 | Tlr6-5 | TATGGTAGACTATTCAAAC | tatcttgtggaaaggacgaaacaccGTATGGTAGACTATTCAAACgtttaagagctatgctggaaacagc |
| Tmem123 | Tmem123-1 | CCTCCAATATTAACATAAC | tatcttgtggaaaggacgaaacaccGCCTCCAATATTAACATAACgtttaagagctatgctggaaacagc |
| Tmem123 | Tmem123-3 | AGGCAGTAGGCGTCACATG | tatcttgtggaaaggacgaaacaccGAGGCAGTAGGCGTCACATGgtttaagagctatgctggaaacagc |
| Tmem123 | Tmem123-5 | GTTTATTGACAACTGTAAC | tatcttgtggaaaggacgaaacaccGGTTTATTGACAACTGTAACgtttaagagctatgctggaaacagc |
| Tmem225 | Tmem225-1 | TTGATAGTAGGACTCATCA | tatcttgtggaaaggacgaaacaccGTTGATAGTAGGACTCATCAgtttaagagctatgctggaaacagc |
| Tmem225 | Tmem225-4 | GTTACCTGCGAAGAAACTC | tatcttgtggaaaggacgaaacaccGGTTACCTGCGAAGAAACTCgtttaagagctatgctggaaacagc |
| Tmem225 | Tmem225-5 | AATGATCGCATAGAACAAA | tatcttgtggaaaggacgaaacaccGAATGATCGCATAGAACAAAgtttaagagctatgctggaaacagc |
| Tmem258 | Tmem258-1 | CGGCATGTTCTTCACTGCC | tatcttgtggaaaggacgaaacaccGCGGCATGTTCTTCACTGCCgtttaagagctatgctggaaacagc |
| Tmem258 | Tmem258-3 | GCAGTCCTACCTAGTTACG | tatcttgtggaaaggacgaaacaccGGCAGTCCTACCTAGTTACGgtttaagagctatgctggaaacagc |
| Tmem258 | Tmem258-4 | TAAATATCGCGTGTGTACT | tatcttgtggaaaggacgaaacaccGTAAATATCGCGTGTGTACTgtttaagagctatgctggaaacagc |
| Tmem41a | Tmem41a-2 | GCCACTTTATCAGGAAAGT | tatcttgtggaaaggacgaaacaccGGCCACTTTATCAGGAAAGTgtttaagagctatgctggaaacagc |
| Tmem41a | Tmem41a-3 | AGTCTTTTCGGCAAACAGC | tatcttgtggaaaggacgaaacaccGAGTCTTTTCGGCAAACAGCgtttaagagctatgctggaaacagc |
| Tmem41a | Tmem41a-5 | TGAAGTTCTTCGAGAGTAC | tatcttgtggaaaggacgaaacaccGTGAAGTTCTTCGAGAGTACgtttaagagctatgctggaaacagc |
| Tmx1 | Tmx1-2 | CTTTTGTAGTTATGCTCCC | tatcttgtggaaaggacgaaacaccGCTTTTGTAGTTATGCTCCCgtttaagagctatgctggaaacagc |
| Tmx1 | Tmx1-3 | GGGAAAGTTTTGCCGAGTG | tatcttgtggaaaggacgaaacaccGGGGAAAGTTTTGCCGAGTGgtttaagagctatgctggaaacagc |
| Tmx1 | Tmx1-5 | CTTCGTAAGTGATAAAGAA | tatcttgtggaaaggacgaaacaccGCTTCGTAAGTGATAAAGAAgtttaagagctatgctggaaacagc |
| Tomm6 | Tomm6-2 | AAATTGAGGATCAAGTTCC | tatcttgtggaaaggacgaaacaccGAAATTGAGGATCAAGTTCCgtttaagagctatgctggaaacagc |
| Tomm6 | Tomm6-4 | GGCGAAGCGGAAGACACCG | tatcttgtggaaaggacgaaacaccGGGCGAAGCGGAAGACACCGgtttaagagctatgctggaaacagc |
| Tomm6 | Tomm6-5 | ACGTGGGAGACTGGCTCCG | tatcttgtggaaaggacgaaacaccGACGTGGGAGACTGGCTCCGgtttaagagctatgctggaaacagc |
| Tonsl | Tonsl-1 | ATCCGTGACTCTCCGTACC | tatcttgtggaaaggacgaaacaccGATCCGTGACTCTCCGTACCgtttaagagctatgctggaaacagc |
| Tonsl | Tonsl-2 | TGCATGACGCCCTCAACTG | tatcttgtggaaaggacgaaacaccGTGCATGACGCCCTCAACTGgtttaagagctatgctggaaacagc |
| Tonsl | Tonsl-3 | GGAACCGGCGCAATGACAT | tatcttgtggaaaggacgaaacaccGGGAACCGGCGCAATGACATgtttaagagctatgctggaaacagc |
| Top2a | Top2a-1 | CGGCACGTACCGTCTACCA | tatcttgtggaaaggacgaaacaccGCGGCACGTACCGTCTACCAgtttaagagctatgctggaaacagc |

| | | | |
|---|---|---|---|
| Top2a | Top2a-3 | GCACACTTATCCTGACTGA | tatcttgtggaaaggacgaaacaccGGCACACTTATCCTGACTGAgtttaagagctatgctggaaacagc |
| Top2a | Top2a-4 | TGAACAGGTCAATCCCCGG | tatcttgtggaaaggacgaaacaccGTGAACAGGTCAATCCCCGGgtttaagagctatgctggaaacagc |
| Top3a | Top3a-2 | TGGCCCTACTCCCCGAAAC | tatcttgtggaaaggacgaaacaccGTGGCCCTACTCCCCGAAACgtttaagagctatgctggaaacagc |
| Top3a | Top3a-4 | GTCCAACGGGCGTATGAGG | tatcttgtggaaaggacgaaacaccGGTCCAACGGGCGTATGAGGgtttaagagctatgctggaaacagc |
| Top3a | Top3a-5 | AGCAAGTCGGCGATCCCCT | tatcttgtggaaaggacgaaacaccGAGCAAGTCGGCGATCCCCTgtttaagagctatgctggaaacagc |
| Topbp1 | Topbp1-2 | GAAATGACGACTCCACGAT | tatcttgtggaaaggacgaaacaccGGAAATGACGACTCCACGATgtttaagagctatgctggaaacagc |
| Topbp1 | Topbp1-3 | ACTCTACAGTTCGTGAAGA | tatcttgtggaaaggacgaaacaccGACTCTACAGTTCGTGAAGAgtttaagagctatgctggaaacagc |
| Topbp1 | Topbp1-4 | GTTGTTACAAACACGTGGC | tatcttgtggaaaggacgaaacaccGGTTGTTACAAACACGTGGCgtttaagagctatgctggaaacagc |
| Triap1 | Triap1-1 | TTGCTGAGAAGTTCCTCAA | tatcttgtggaaaggacgaaacaccGTTGCTGAGAAGTTCCTCAAgtttaagagctatgctggaaacagc |
| Triap1 | Triap1-3 | TTTGAAGAGGTCGGTGCAC | tatcttgtggaaaggacgaaacaccGTTTGAAGAGGTCGGTGCACgtttaagagctatgctggaaacagc |
| Triap1 | Triap1-5 | ACTCACCTGCACGCACTGC | tatcttgtggaaaggacgaaacaccGACTCACCTGCACGCACTGCgtttaagagctatgctggaaacagc |
| Trim17 | Trim17-2 | GACATAAAATATCTACGGG | tatcttgtggaaaggacgaaacaccGGACATAAAATATCTACGGGgtttaagagctatgctggaaacagc |
| Trim17 | Trim17-3 | GAAGAACGGAGGATCCTCC | tatcttgtggaaaggacgaaacaccGGAAGAACGGAGGATCCTCCgtttaagagctatgctggaaacagc |
| Trim17 | Trim17-4 | GAGCGGCTTTGGTGGTCCA | tatcttgtggaaaggacgaaacaccGGAGCGGCTTTGGTGGTCCAgtttaagagctatgctggaaacagc |
| Trim28 | Trim28-1 | AGCCGCGTCGTCCCCTGCG | tatcttgtggaaaggacgaaacaccGAGCCGCGTCGTCCCCTGCGgtttaagagctatgctggaaacagc |
| Trim28 | Trim28-4 | TTGTCCATTAGGATCCGCC | tatcttgtggaaaggacgaaacaccGTTGTCCATTAGGATCCGCCgtttaagagctatgctggaaacagc |
| Trim28 | Trim28-5 | GTCCTGGTACGAACTCCAC | tatcttgtggaaaggacgaaacaccGGTCCTGGTACGAACTCCACgtttaagagctatgctggaaacagc |
| Trnt1 | Trnt1-1 | TCCAGTCCCTTTTCACCGA | tatcttgtggaaaggacgaaacaccGTCCAGTCCCTTTTCACCGAgtttaagagctatgctggaaacagc |
| Trnt1 | Trnt1-2 | ATCATGAATTAAGAATAGC | tatcttgtggaaaggacgaaacaccGATCATGAATTAAGAATAGCgtttaagagctatgctggaaacagc |
| Trnt1 | Trnt1-3 | GTGAGGGATTTACTGAATG | tatcttgtggaaaggacgaaacaccGGTGAGGGATTTACTGAATGgtttaagagctatgctggaaacagc |
| Trp53 | Trp53-2 | AATAAGCTATTCTGCCAGC | tatcttgtggaaaggacgaaacaccGAATAAGCTATTCTGCCAGCgtttaagagctatgctggaaacagc |
| Trp53 | Trp53-3 | CGCGGACACGGCTCCCAGC | tatcttgtggaaaggacgaaacaccGCGCGGACACGGCTCCCAGCgtttaagagctatgctggaaacagc |
| Trp53 | Trp53-5 | CCCGGATAAGATGCTGGGG | tatcttgtggaaaggacgaaacaccGCCCGGATAAGATGCTGGGGgtttaagagctatgctggaaacagc |
| Trp53bp1 | Trp53bp1-1 | GAACCTGTCAGACCCGATC | tatcttgtggaaaggacgaaacaccGGAACCTGTCAGACCCGATCgtttaagagctatgctggaaacagc |
| Trp53bp1 | Trp53bp1-4 | GCGTGACTAACCTTACAAG | tatcttgtggaaaggacgaaacaccGGCGTGACTAACCTTACAAGgtttaagagctatgctggaaacagc |
| Trp53bp1 | Trp53bp1-5 | CCCGGTAAAGGTATCCATG | tatcttgtggaaaggacgaaacaccGCCCGGTAAAGGTATCCATGgtttaagagctatgctggaaacagc |
| Tsen34 | Tsen34-1 | CCCCGCCAGAACTCGCGCC | tatcttgtggaaaggacgaaacaccGCCCCGCCAGAACTCGCGCCgtttaagagctatgctggaaacagc |
| Tsen34 | Tsen34-2 | CACCGCGCCTATCTCGGCC | tatcttgtggaaaggacgaaacaccGCACCGCGCCTATCTCGGCCgtttaagagctatgctggaaacagc |
| Tsen34 | Tsen34-5 | TCAGCCCTGCTTATCCAGC | tatcttgtggaaaggacgaaacaccGTCAGCCCTGCTTATCCAGCgtttaagagctatgctggaaacagc |
| Ttc27 | Ttc27-1 | CATATCTCAACGTCCACCA | tatcttgtggaaaggacgaaacaccGCATATCTCAACGTCCACCAgtttaagagctatgctggaaacagc |
| Ttc27 | Ttc27-3 | ATTAACGTACCAGACTCCA | tatcttgtggaaaggacgaaacaccGATTAACGTACCAGACTCCAgtttaagagctatgctggaaacagc |
| Ttc27 | Ttc27-4 | CCAGTTCATAAGCTAACGG | tatcttgtggaaaggacgaaacaccGCCAGTTCATAAGCTAACGGgtttaagagctatgctggaaacagc |
| Ttc4 | Ttc4-2 | ACCTGAAAGCCATCATAAG | tatcttgtggaaaggacgaaacaccGACCTGAAAGCCATCATAAGgtttaagagctatgctggaaacagc |
| Ttc4 | Ttc4-3 | TGCTGTCCTGTATACCAAC | tatcttgtggaaaggacgaaacaccGTGCTGTCCTGTATACCAACgtttaagagctatgctggaaacagc |
| Ttc4 | Ttc4-5 | CAAGTCAGGAAACTCCTCG | tatcttgtggaaaggacgaaacaccGCAAGTCAGGAAACTCCTCGgtttaagagctatgctggaaacagc |
| Ttf2 | Ttf2-1 | AGTGGGGACCCCGGCCTAC | tatcttgtggaaaggacgaaacaccGAGTGGGGACCCCGGCCTACgtttaagagctatgctggaaacagc |
| Ttf2 | Ttf2-4 | ACGTCCCATGGCGGCAGGT | tatcttgtggaaaggacgaaacaccGACGTCCCATGGCGGCAGGTgtttaagagctatgctggaaacagc |
| Ttf2 | Ttf2-5 | TGTGCCGGACGAACACATG | tatcttgtggaaaggacgaaacaccGTGTGCCGGACGAACACATGgtttaagagctatgctggaaacagc |
| Tti1 | Tti1-1 | CAGGTCCCCATAATAACCC | tatcttgtggaaaggacgaaacaccGCAGGTCCCCATAATAACCCgtttaagagctatgctggaaacagc |
| Tti1 | Tti1-4 | ACCTTCAAGAGGGGTCCCG | tatcttgtggaaaggacgaaacaccGACCTTCAAGAGGGGTCCCGgtttaagagctatgctggaaacagc |
| Tti1 | Tti1-5 | TCAGCGCAGTTGAGATCCC | tatcttgtggaaaggacgaaacaccGTCAGCGCAGTTGAGATCCCgtttaagagctatgctggaaacagc |
| Tufm | Tufm-1 | TTGTGTCGCGGCCTAGCTG | tatcttgtggaaaggacgaaacaccGTTGTGTCGCGGCCTAGCTGgtttaagagctatgctggaaacagc |
| Tufm | Tufm-2 | CCATGGTCCACATGTCCGA | tatcttgtggaaaggacgaaacaccGCCATGGTCCACATGTCCGAgtttaagagctatgctggaaacagc |
| Tufm | Tufm-5 | GAGCCGAGCCTACGATGAC | tatcttgtggaaaggacgaaacaccGGAGCCGAGCCTACGATGACgtttaagagctatgctggaaacagc |
| Tut1 | Tut1-1 | TCGGAGTTGCACCAAATCC | tatcttgtggaaaggacgaaacaccGTCGGAGTTGCACCAAATCCgtttaagagctatgctggaaacagc |
| Tut1 | Tut1-3 | GGACTTCGAGTCCGGCCAA | tatcttgtggaaaggacgaaacaccGGGACTTCGAGTCCGGCCAAgtttaagagctatgctggaaacagc |
| Tut1 | Tut1-5 | CAAAAGTACCGGTTACTGA | tatcttgtggaaaggacgaaacaccGCAAAAGTACCGGTTACTGAgtttaagagctatgctggaaacagc |
| Txnl4a | Txnl4a-1 | ATTCGTTTCGGACACGACT | tatcttgtggaaaggacgaaacaccGATTCGTTTCGGACACGACTgtttaagagctatgctggaaacagc |
| Txnl4a | Txnl4a-3 | CTGTACAGCATCGCCGAAA | tatcttgtggaaaggacgaaacaccGCTGTACAGCATCGCCGAAAgtttaagagctatgctggaaacagc |
| Txnl4a | Txnl4a-5 | TCATGATTGATTTGGGCAC | tatcttgtggaaaggacgaaacaccGTCATGATTGATTTGGGCACgtttaagagctatgctggaaacagc |
| Uba3 | Uba3-1 | GCTTGTATTGAGTGCACTC | tatcttgtggaaaggacgaaacaccGGCTTGTATTGAGTGCACTCgtttaagagctatgctggaaacagc |
| Uba3 | Uba3-2 | TCTCGCTATGATAGAGTCC | tatcttgtggaaaggacgaaacaccGTCTCGCTATGATAGAGTCCgtttaagagctatgctggaaacagc |
| Uba3 | Uba3-4 | AGAGTTCCTAACTGCAACG | tatcttgtggaaaggacgaaacaccGAGAGTTCCTAACTGCAACGgtttaagagctatgctggaaacagc |
| Ube2cbp | Ube2cbp-1 | GGCCTATTAGCAAAGGGGT | tatcttgtggaaaggacgaaacaccGGGCCTATTAGCAAAGGGGTgtttaagagctatgctggaaacagc |
| Ube2cbp | Ube2cbp-3 | TACTGCAGCCCGCCACACG | tatcttgtggaaaggacgaaacaccGTACTGCAGCCCGCCACACGgtttaagagctatgctggaaacagc |
| Ube2cbp | Ube2cbp-4 | GTACAAGTCTGACCCCTGC | tatcttgtggaaaggacgaaacaccGGTACAAGTCTGACCCCTGCgtttaagagctatgctggaaacagc |
| Ube2m | Ube2m-3 | CCACAGGAGCCGAACCCTG | tatcttgtggaaaggacgaaacaccGCCACAGGAGCCGAACCCTGgtttaagagctatgctggaaacagc |
| Ube2m | Ube2m-5 | AATTATGGAGTTTATCGTA | tatcttgtggaaaggacgaaacaccGAATTATGGAGTTTATCGTAgtttaagagctatgctggaaacagc |
| Ube2m | Ube2m-9 | GCGCAGCTCCGGATTCAGA | tatcttgtggaaaggacgaaacaccGGCGCAGCTCCGGATTCAGAgtttaagagctatgctggaaacagc |
| Ube2t | Ube2t-1 | GAGAAAGGTGTTTTCACAC | tatcttgtggaaaggacgaaacaccGGAGAAAGGTGTTTTCACACgtttaagagctatgctggaaacagc |
| Ube2t | Ube2t-2 | TACCCATTTGAACCTCCAC | tatcttgtggaaaggacgaaacaccGTACCCATTTGAACCTCCACgtttaagagctatgctggaaacagc |
| Ube2t | Ube2t-3 | GAGTTAGAAATCGGACCTG | tatcttgtggaaaggacgaaacaccGGAGTTAGAAATCGGACCTGgtttaagagctatgctggaaacagc |
| Ube2w | Ube2w-1 | TACCTGAGGAGAGTCAAAA | tatcttgtggaaaggacgaaacaccGTACCTGAGGAGAGTCAAAAgtttaagagctatgctggaaacagc |
| Ube2w | Ube2w-2 | TAGACATGGAAGGTGCACC | tatcttgtggaaaggacgaaacaccGTAGACATGGAAGGTGCACCgtttaagagctatgctggaaacagc |
| Ube2w | Ube2w-4 | CATTTAAAGTCATTCCAGG | tatcttgtggaaaggacgaaacaccGCATTTAAAGTCATTCCAGGgtttaagagctatgctggaaacagc |
| Ubp1 | Ubp1-1 | ACACATTACATATTGGAAG | tatcttgtggaaaggacgaaacaccGACACATTACATATTGGAAGgtttaagagctatgctggaaacagc |
| Ubp1 | Ubp1-2 | CACTTACTTACTTGAACCA | tatcttgtggaaaggacgaaacaccGCACTTACTTACTTGAACCAgtttaagagctatgctggaaacagc |
| Ubp1 | Ubp1-3 | CATCCGAATTTCATAAGAC | tatcttgtggaaaggacgaaacaccGCATCCGAATTTCATAAGACgtttaagagctatgctggaaacagc |
| Ubr4 | Ubr4-1 | GGGGTACTCTCGGTGATAC | tatcttgtggaaaggacgaaacaccGGGGGTACTCTCGGTGATACgtttaagagctatgctggaaacagc |
| Ubr4 | Ubr4-2 | GCGATGGTCGTCTCGCCTC | tatcttgtggaaaggacgaaacaccGGCGATGGTCGTCTCGCCTCgtttaagagctatgctggaaacagc |
| Ubr4 | Ubr4-4 | ACGAAGTCTGCGGTGACGA | tatcttgtggaaaggacgaaacaccGACGAAGTCTGCGGTGACGAgtttaagagctatgctggaaacagc |
| Ubr7 | Ubr7-1 | AATGAGGCGTGCGCCGTCC | tatcttgtggaaaggacgaaacaccGAATGAGGCGTGCGCCGTCCgtttaagagctatgctggaaacagc |
| Ubr7 | Ubr7-2 | AAGCTAGACAAATCCCTGC | tatcttgtggaaaggacgaaacaccGAAGCTAGACAAATCCCTGCgtttaagagctatgctggaaacagc |
| Ubr7 | Ubr7-4 | AATGTTGCTTACCCTGCCA | tatcttgtggaaaggacgaaacaccGAATGTTGCTTACCCTGCCAgtttaagagctatgctggaaacagc |
| Ufsp2 | Ufsp2-2 | GGTCGATATCTCCAGCATG | tatcttgtggaaaggacgaaacaccGGGTCGATATCTCCAGCATGgtttaagagctatgctggaaacagc |
| Ufsp2 | Ufsp2-4 | TAACGGTTTTATATCCATC | tatcttgtggaaaggacgaaacaccGTAACGGTTTTATATCCATCgtttaagagctatgctggaaacagc |

| | | | |
|---|---|---|---|
| Ufsp2 | Ufsp2-5 | CAGCTCCAAGCCTATCGAA | tatcttgtggaaaggacgaaacaccGCAGCTCCAAGCCTATCGAAgtttaagagctatgctggaaacagc |
| Ulk2 | Ulk2-1 | GGTCTAATACCAGCCCCAT | tatcttgtggaaaggacgaaacaccGGGTCTAATACCAGCCCCATgtttaagagctatgctggaaacagc |
| Ulk2 | Ulk2-2 | GTCTGGCCGTAGTCCCCAT | tatcttgtggaaaggacgaaacaccGGTCTGGCCGTAGTCCCCATgtttaagagctatgctggaaacagc |
| Ulk2 | Ulk2-3 | GTTGGGAGGACCCAACGGT | tatcttgtggaaaggacgaaacaccGGTTGGGAGGACCCAACGGTgtttaagagctatgctggaaacagc |
| Usp1 | Usp1-1 | GTGTGTTCAGCAGTCGCCT | tatcttgtggaaaggacgaaacaccGGTGTGTTCAGCAGTCGCCTgtttaagagctatgctggaaacagc |
| Usp1 | Usp1-2 | CTGGCTGAATTATCCGGCA | tatcttgtggaaaggacgaaacaccGCTGGCTGAATTATCCGGCAgtttaagagctatgctggaaacagc |
| Usp1 | Usp1-4 | GCTATCGGAGACACACTAG | tatcttgtggaaaggacgaaacaccGGCTATCGGAGACACACTAGgtttaagagctatgctggaaacagc |
| Usp5 | Usp5-1 | CCGATTAGTGGAGAACTCA | tatcttgtggaaaggacgaaacaccGCCGATTAGTGGAGAACTCAgtttaagagctatgctggaaacagc |
| Usp5 | Usp5-2 | TCCGGATGCCCGTGTAACC | tatcttgtggaaaggacgaaacaccGTCCGGATGCCCGTGTAACCgtttaagagctatgctggaaacagc |
| Usp5 | Usp5-3 | GCGCTACTTCGATGGCAGT | tatcttgtggaaaggacgaaacaccGGCGCTACTTCGATGGCAGTgtttaagagctatgctggaaacagc |
| Vhl | Vhl-1 | ACAAAGGCAGCACGACGCG | tatcttgtggaaaggacgaaacaccGACAAAGGCAGCACGACGCGgtttaagagctatgctggaaacagc |
| Vhl | Vhl-3 | GTAAGATCGGGTAGGGCTG | tatcttgtggaaaggacgaaacaccGGTAAGATCGGGTAGGGCTGgtttaagagctatgctggaaacagc |
| Vhl | Vhl-5 | AGTGTATACCCTGAAAGAG | tatcttgtggaaaggacgaaacaccGAGTGTATACCCTGAAAGAGgtttaagagctatgctggaaacagc |
| Vmn1r195 | Vmn1r195-1 | ATGTGCTTACATTTACCAT | tatcttgtggaaaggacgaaacaccGATGTGCTTACATTTACCATgtttaagagctatgctggaaacagc |
| Vmn1r195 | Vmn1r195-3 | GGTCTGAACTTTCTCAAAA | tatcttgtggaaaggacgaaacaccGGGTCTGAACTTTCTCAAAAgtttaagagctatgctggaaacagc |
| Vmn1r195 | Vmn1r195-4 | GTTCAGACCACAAACAATA | tatcttgtggaaaggacgaaacaccGGTTCAGACCACAAACAATAgtttaagagctatgctggaaacagc |
| Vmn1r223 | Vmn1r223-1 | TAATACACTGTTCATTCCA | tatcttgtggaaaggacgaaacaccGTAATACACTGTTCATTCCAgtttaagagctatgctggaaacagc |
| Vmn1r223 | Vmn1r223-2 | TATCTTCATGGCTAAAGCT | tatcttgtggaaaggacgaaacaccGTATCTTCATGGCTAAAGCTgtttaagagctatgctggaaacagc |
| Vmn1r223 | Vmn1r223-3 | TCATTTCACTTATTGGACT | tatcttgtggaaaggacgaaacaccGTCATTTCACTTATTGGACTgtttaagagctatgctggaaacagc |
| Vmn1r224 | Vmn1r224-1 | TAGTAATTGAATAGGTAGC | tatcttgtggaaaggacgaaacaccGTAGTAATTGAATAGGTAGCgtttaagagctatgctggaaacagc |
| Vmn1r224 | Vmn1r224-3 | CTTTGGTGGATTCTCTACA | tatcttgtggaaaggacgaaacaccGCTTTGGTGGATTCTCTACAgtttaagagctatgctggaaacagc |
| Vmn1r224 | Vmn1r224-5 | AAATCATCATGAACTACAG | tatcttgtggaaaggacgaaacaccGAAATCATCATGAACTACAGgtttaagagctatgctggaaacagc |
| Vmn1r236 | Vmn1r236-1 | ATTCCTATAGACAACTCAC | tatcttgtggaaaggacgaaacaccGATTCCTATAGACAACTCACgtttaagagctatgctggaaacagc |
| Vmn1r236 | Vmn1r236-2 | TCATCTCTCTGACTATAAT | tatcttgtggaaaggacgaaacaccGTCATCTCTCTGACTATAATgtttaagagctatgctggaaacagc |
| Vmn1r236 | Vmn1r236-5 | GTGGCATAAAGGATATCAC | tatcttgtggaaaggacgaaacaccGGTGGCATAAAGGATATCACgtttaagagctatgctggaaacagc |
| Vmn1r31 | Vmn1r31-1 | ACCGGTTCTTCTATGTTCA | tatcttgtggaaaggacgaaacaccGACCGGTTCTTCTATGTTCAgtttaagagctatgctggaaacagc |
| Vmn1r31 | Vmn1r31-2 | GAAAATGATGTCAAATGTA | tatcttgtggaaaggacgaaacaccGGAAAATGATGTCAAATGTAgtttaagagctatgctggaaacagc |
| Vmn1r31 | Vmn1r31-3 | AAATATGTCTGTAAGCAAC | tatcttgtggaaaggacgaaacaccGAAATATGTCTGTAAGCAACgtttaagagctatgctggaaacagc |
| Vmn1r4 | Vmn1r4-1 | GGCAGGAAGTCAGGTCCGT | tatcttgtggaaaggacgaaacaccGGGCAGGAAGTCAGGTCCGTgtttaagagctatgctggaaacagc |
| Vmn1r4 | Vmn1r4-4 | GACAGCCTGGAACACGCTC | tatcttgtggaaaggacgaaacaccGGACAGCCTGGAACACGCTCgtttaagagctatgctggaaacagc |
| Vmn1r4 | Vmn1r4-5 | ACATAATATTCTATGTTGG | tatcttgtggaaaggacgaaacaccGACATAATATTCTATGTTGGgtttaagagctatgctggaaacagc |
| Vmn1r72 | Vmn1r72-1 | CTGTAGAGGATAATGATCA | tatcttgtggaaaggacgaaacaccGCTGTAGAGGATAATGATCAgtttaagagctatgctggaaacagc |
| Vmn1r72 | Vmn1r72-2 | ACTGCTCATGGTTTTCATC | tatcttgtggaaaggacgaaacaccGACTGCTCATGGTTTTCATCgtttaagagctatgctggaaacagc |
| Vmn1r72 | Vmn1r72-4 | GGTCAAGTGCTCTATAATC | tatcttgtggaaaggacgaaacaccGGGTCAAGTGCTCTATAATCgtttaagagctatgctggaaacagc |
| Vmn1r87 | Vmn1r87-3 | CAGTTCTAAACATTCCAAG | tatcttgtggaaaggacgaaacaccGCAGTTCTAAACATTCCAAGgtttaagagctatgctggaaacagc |
| Vmn1r87 | Vmn1r87-4 | TAGTAGCAAGATAGTACAG | tatcttgtggaaaggacgaaacaccGTAGTAGCAAGATAGTACAGgtttaagagctatgctggaaacagc |
| Vmn1r87 | Vmn1r87-5 | CATGAGATCCTTTGCAGTC | tatcttgtggaaaggacgaaacaccGCATGAGATCCTTTGCAGTCgtttaagagctatgctggaaacagc |
| Vmn2r89 | Vmn2r89-2 | TGATATTCAGCATCGTTGG | tatcttgtggaaaggacgaaacaccGTGATATTCAGCATCGTTGGgtttaagagctatgctggaaacagc |
| Vmn2r89 | Vmn2r89-3 | CATGTAACATAGGCCTTAC | tatcttgtggaaaggacgaaacaccGCATGTAACATAGGCCTTACgtttaagagctatgctggaaacagc |
| Vmn2r89 | Vmn2r89-6 | GTGGTGTGCAAAAGTGATA | tatcttgtggaaaggacgaaacaccGGTGGTGTGCAAAAGTGATAgtttaagagctatgctggaaacagc |
| Vps54 | Vps54-1 | TAGGATCATTTAATGCTGC | tatcttgtggaaaggacgaaacaccGTAGGATCATTTAATGCTGCgtttaagagctatgctggaaacagc |
| Vps54 | Vps54-3 | TTCTGAACGTAGAGAGATC | tatcttgtggaaaggacgaaacaccGTTCTGAACGTAGAGAGATCgtttaagagctatgctggaaacagc |
| Vps54 | Vps54-5 | CTTTAACAGATCAATCCAC | tatcttgtggaaaggacgaaacaccGCTTTAACAGATCAATCCACgtttaagagctatgctggaaacagc |
| Vrk2 | Vrk2-1 | ATTTCAACATCACTCCTCC | tatcttgtggaaaggacgaaacaccGATTTCAACATCACTCCTCCgtttaagagctatgctggaaacagc |
| Vrk2 | Vrk2-2 | CCTACAGATATTGTCCCAA | tatcttgtggaaaggacgaaacaccGCCTACAGATATTGTCCCAAgtttaagagctatgctggaaacagc |
| Vrk2 | Vrk2-4 | AATTAACCGTCCTACAACT | tatcttgtggaaaggacgaaacaccGAATTAACCGTCCTACAACTgtttaagagctatgctggaaacagc |
| Wbp2nl | Wbp2nl-1 | TTTGTACCACTAAAGAGGT | tatcttgtggaaaggacgaaacaccGTTTGTACCACTAAAGAGGTgtttaagagctatgctggaaacagc |
| Wbp2nl | Wbp2nl-2 | TGTAGTTTGCACCAAAGAT | tatcttgtggaaaggacgaaacaccGTGTAGTTTGCACCAAAGATgtttaagagctatgctggaaacagc |
| Wbp2nl | Wbp2nl-4 | GATGACATAAATTCCTAGA | tatcttgtggaaaggacgaaacaccGGATGACATAAATTCCTAGAgtttaagagctatgctggaaacagc |
| Wdr24 | Wdr24-1 | GTTGCGGGATGGCCGACCT | tatcttgtggaaaggacgaaacaccGGTTGCGGGATGGCCGACCTgtttaagagctatgctggaaacagc |
| Wdr24 | Wdr24-3 | ATCTTGTCACGTCCACCCG | tatcttgtggaaaggacgaaacaccGATCTTGTCACGTCCACCCGgtttaagagctatgctggaaacagc |
| Wdr24 | Wdr24-4 | GGGTCATGCGGATGACGCC | tatcttgtggaaaggacgaaacaccGGGGTCATGCGGATGACGCCgtttaagagctatgctggaaacagc |
| Wdr45b | Wdr45b-1 | TCACCTTTAGGGTTGTAGC | tatcttgtggaaaggacgaaacaccGTCACCTTTAGGGTTGTAGCgtttaagagctatgctggaaacagc |
| Wdr45b | Wdr45b-2 | GGTTTCAAAGACGTGCAAC | tatcttgtggaaaggacgaaacaccGGGTTTCAAAGACGTGCAACgtttaagagctatgctggaaacagc |
| Wdr45b | Wdr45b-3 | AACGTGAACACCTTAATCA | tatcttgtggaaaggacgaaacaccGAACGTGAACACCTTAATCAgtttaagagctatgctggaaacagc |
| Wdr48 | Wdr48-2 | ACCTTATGCGTCCTGAGTG | tatcttgtggaaaggacgaaacaccGACCTTATGCGTCCTGAGTGgtttaagagctatgctggaaacagc |
| Wdr48 | Wdr48-4 | AAGAGCTGGTAGCGTCAGC | tatcttgtggaaaggacgaaacaccGAAGAGCTGGTAGCGTCAGCgtttaagagctatgctggaaacagc |
| Wdr48 | Wdr48-5 | AGCTTCATGAGTTTCGCGC | tatcttgtggaaaggacgaaacaccGAGCTTCATGAGTTTCGCGCgtttaagagctatgctggaaacagc |
| Wdr55 | Wdr55-1 | CAGCAGATCTCGAGTCGGA | tatcttgtggaaaggacgaaacaccGCAGCAGATCTCGAGTCGGAgtttaagagctatgctggaaacagc |
| Wdr55 | Wdr55-2 | CACATCTTAGATGTCGAAC | tatcttgtggaaaggacgaaacaccGCACATCTTAGATGTCGAACgtttaagagctatgctggaaacagc |
| Wdr55 | Wdr55-4 | TGGTTACGGGGGACGACAC | tatcttgtggaaaggacgaaacaccGTGGTTACGGGGGACGACACgtttaagagctatgctggaaacagc |
| Wdr61 | Wdr61-2 | ATTCCGTCTATGGCTCCGC | tatcttgtggaaaggacgaaacaccGATTCCGTCTATGGCTCCGCgtttaagagctatgctggaaacagc |
| Wdr61 | Wdr61-3 | ATGGGAAATACCTGGCCAG | tatcttgtggaaaggacgaaacaccGATGGGAAATACCTGGCCAGgtttaagagctatgctggaaacagc |
| Wdr61 | Wdr61-4 | CCTTACATATGCAATACTA | tatcttgtggaaaggacgaaacaccGCCTTACATATGCAATACTAgtttaagagctatgctggaaacagc |
| Wdr77 | Wdr77-2 | GGGGACAAAGGTATCCTAG | tatcttgtggaaaggacgaaacaccGGGGGACAAAGGTATCCTAGgtttaagagctatgctggaaacagc |
| Wdr77 | Wdr77-3 | CTGGCACACAAGCTGTCAG | tatcttgtggaaaggacgaaacaccGCTGGCACACAAGCTGTCAGgtttaagagctatgctggaaacagc |
| Wdr77 | Wdr77-4 | AGCATCAAAATTTGGGACC | tatcttgtggaaaggacgaaacaccGAGCATCAAAATTTGGGACCgtttaagagctatgctggaaacagc |
| Wee1 | Wee1-1 | AGCGGCCACAGTACGGGGG | tatcttgtggaaaggacgaaacaccGAGCGGCCACAGTACGGGGGgtttaagagctatgctggaaacagc |
| Wee1 | Wee1-4 | TAGTCGGGCTGCGGCGAGC | tatcttgtggaaaggacgaaacaccGTAGTCGGGCTGCGGCGAGCgtttaagagctatgctggaaacagc |
| Wee1 | Wee1-5 | AGTGTGCGGCGTGTCGAAC | tatcttgtggaaaggacgaaacaccGAGTGTGCGGCGTGTCGAACgtttaagagctatgctggaaacagc |
| Wtap | Wtap-2 | CGGCTAGCAACCAAAGAGC | tatcttgtggaaaggacgaaacaccGCGGCTAGCAACCAAAGAGCgtttaagagctatgctggaaacagc |
| Wtap | Wtap-3 | GGAGAACATTCTTGTCATG | tatcttgtggaaaggacgaaacaccGGGAGAACATTCTTGTCATGgtttaagagctatgctggaaacagc |
| Wtap | Wtap-4 | ACTCAGCAAACGATGTGAC | tatcttgtggaaaggacgaaacaccGACTCAGCAAACGATGTGACgtttaagagctatgctggaaacagc |
| Xab2 | Xab2-1 | TACACGCTTGTGCCACTCG | tatcttgtggaaaggacgaaacaccGTACACGCTTGTGCCACTCGgtttaagagctatgctggaaacagc |
| Xab2 | Xab2-2 | GGCGTAGCTGTCGAACACC | tatcttgtggaaaggacgaaacaccGGGCGTAGCTGTCGAACACCgtttaagagctatgctggaaacagc |
| Xab2 | Xab2-5 | CGCTACATCGAGTTCAAAC | tatcttgtggaaaggacgaaacaccGCGCTACATCGAGTTCAAACgtttaagagctatgctggaaacagc |

| | | | |
|---|---|---|---|
| Xcr1 | Xcr1-2 | CAGGACTTTGTTTCGCACA | tatcttgtggaaaggacgaaacaccGCAGGACTTTGTTTCGCACAgtttaagagctatgctggaaacagc |
| Xcr1 | Xcr1-4 | ACCAGCACACGGCAGCGGA | tatcttgtggaaaggacgaaacaccGACCAGCACACGGCAGCGGAgtttaagagctatgctggaaacagc |
| Xcr1 | Xcr1-5 | CACTACAGACAGGTATCGG | tatcttgtggaaaggacgaaacaccGCACTACAGACAGGTATCGGgtttaagagctatgctggaaacagc |
| Xpo5 | Xpo5-2 | CGGCCCTGAACACTCTAGC | tatcttgtggaaaggacgaaacaccGCGGCCCTGAACACTCTAGCgtttaagagctatgctggaaacagc |
| Xpo5 | Xpo5-4 | AATCGGGAATATTCACAGC | tatcttgtggaaaggacgaaacaccGAATCGGGAATATTCACAGCgtttaagagctatgctggaaacagc |
| Xpo5 | Xpo5-5 | GTCTAGTGTTCGAAACATC | tatcttgtggaaaggacgaaacaccGGTCTAGTGTTCGAAACATCgtttaagagctatgctggaaacagc |
| Xrcc4 | Xrcc4-1 | GGAGAGTACCAAACCTGAA | tatcttgtggaaaggacgaaacaccGGGAGAGTACCAAACCTGAAgtttaagagctatgctggaaacagc |
| Xrcc4 | Xrcc4-2 | TGATACAAATCAGCCTCCA | tatcttgtggaaaggacgaaacaccGTGATACAAATCAGCCTCCAgtttaagagctatgctggaaacagc |
| Xrcc4 | Xrcc4-3 | TACTGACGGCCATTCAGCC | tatcttgtggaaaggacgaaacaccGTACTGACGGCCATTCAGCCgtttaagagctatgctggaaacagc |
| Xrcc5 | Xrcc5-2 | GACGCCCTGATTGTGTGCA | tatcttgtggaaaggacgaaacaccGGACGCCCTGATTGTGTGCAgtttaagagctatgctggaaacagc |
| Xrcc5 | Xrcc5-4 | GACGCAAGAACGCTAAAGA | tatcttgtggaaaggacgaaacaccGGACGCAAGAACGCTAAAGAgtttaagagctatgctggaaacagc |
| Xrcc5 | Xrcc5-5 | AATGATATCACTTCCGTAG | tatcttgtggaaaggacgaaacaccGAATGATATCACTTCCGTAGgtttaagagctatgctggaaacagc |
| Xrcc6 | Xrcc6-1 | CTAGCACTCGCTTAGCGCC | tatcttgtggaaaggacgaaacaccGCTAGCACTCGCTTAGCGCCgtttaagagctatgctggaaacagc |
| Xrcc6 | Xrcc6-2 | CCGAGACACGGTTGGCCAT | tatcttgtggaaaggacgaaacaccGCCGAGACACGGTTGGCCATgtttaagagctatgctggaaacagc |
| Xrcc6 | Xrcc6-4 | CGGATACATCAAAGCCTCC | tatcttgtggaaaggacgaaacaccGCGGATACATCAAAGCCTCCgtttaagagctatgctggaaacagc |
| Xrn2 | Xrn2-1 | GAATCAGCAGCGTTCAAGG | tatcttgtggaaaggacgaaacaccGGAATCAGCAGCGTTCAAGGgtttaagagctatgctggaaacagc |
| Xrn2 | Xrn2-3 | TCGTTTAAATAATGACCCT | tatcttgtggaaaggacgaaacaccGTCGTTTAAATAATGACCCTgtttaagagctatgctggaaacagc |
| Xrn2 | Xrn2-5 | GCTCCTCTCCACATCAAAT | tatcttgtggaaaggacgaaacaccGGCTCCTCTCCACATCAAATgtttaagagctatgctggaaacagc |
| Yes1 | Yes1-3 | GATGGTATTTTGGCAAAAT | tatcttgtggaaaggacgaaacaccGGATGGTATTTTGGCAAAATgtttaagagctatgctggaaacagc |
| Yes1 | Yes1-4 | TTGAGAGGGAGTAAGCACC | tatcttgtggaaaggacgaaacaccGTTGAGAGGGAGTAAGCACCgtttaagagctatgctggaaacagc |
| Yes1 | Yes1-5 | AGCTGGTGAAGCACTACAC | tatcttgtggaaaggacgaaacaccGAGCTGGTGAAGCACTACACgtttaagagctatgctggaaacagc |
| Yipf1 | Yipf1-1 | GGTGGCCAAAAGATACACT | tatcttgtggaaaggacgaaacaccGGGTGGCCAAAAGATACACTgtttaagagctatgctggaaacagc |
| Yipf1 | Yipf1-4 | ATGGATTAGGAAGTTAGAA | tatcttgtggaaaggacgaaacaccGATGGATTAGGAAGTTAGAAgtttaagagctatgctggaaacagc |
| Yipf1 | Yipf1-5 | CTGGAATTCGGGCACATAA | tatcttgtggaaaggacgaaacaccGCTGGAATTCGGGCACATAAgtttaagagctatgctggaaacagc |
| Zbtb11 | Zbtb11-1 | CCGCGCACGACGTAGCAGG | tatcttgtggaaaggacgaaacaccGCCGCGCACGACGTAGCAGGgtttaagagctatgctggaaacagc |
| Zbtb11 | Zbtb11-3 | CAAATGGGCCGCCTCGATG | tatcttgtggaaaggacgaaacaccGCAAATGGGCCGCCTCGATGgtttaagagctatgctggaaacagc |
| Zbtb11 | Zbtb11-4 | CGTCACATTGCTCATCGAG | tatcttgtggaaaggacgaaacaccGCGTCACATTGCTCATCGAGgtttaagagctatgctggaaacagc |
| Zbtb16 | Zbtb16-2 | ATCTCGAAGCATTCCAGCG | tatcttgtggaaaggacgaaacaccGATCTCGAAGCATTCCAGCGgtttaagagctatgctggaaacagc |
| Zbtb16 | Zbtb16-3 | ATTTTAGAGATCGAATACC | tatcttgtggaaaggacgaaacaccGATTTTAGAGATCGAATACCgtttaagagctatgctggaaacagc |
| Zbtb16 | Zbtb16-4 | CAGGCTAGCACCGTCCGGT | tatcttgtggaaaggacgaaacaccGCAGGCTAGCACCGTCCGGTgtttaagagctatgctggaaacagc |
| Zbtb8os | Zbtb8os-1 | GCATGGGGAGACACTCTGG | tatcttgtggaaaggacgaaacaccGGCATGGGGAGACACTCTGGgtttaagagctatgctggaaacagc |
| Zbtb8os | Zbtb8os-3 | TTGGTTACATGACAGATAC | tatcttgtggaaaggacgaaacaccGTTGGTTACATGACAGATACgtttaagagctatgctggaaacagc |
| Zbtb8os | Zbtb8os-4 | GTGGAGCCCCTCCGCACCG | tatcttgtggaaaggacgaaacaccGGTGGAGCCCCTCCGCACCGgtttaagagctatgctggaaacagc |
| Zc3h10 | Zc3h10-1 | TCTACGACCTTCCGGAAAG | tatcttgtggaaaggacgaaacaccGTCTACGACCTTCCGGAAAGgtttaagagctatgctggaaacagc |
| Zc3h10 | Zc3h10-4 | CCGTGGATGAAACGACAGT | tatcttgtggaaaggacgaaacaccGCCGTGGATGAAACGACAGTgtttaagagctatgctggaaacagc |
| Zc3h10 | Zc3h10-5 | TGAGGAACGTCTGCAAACG | tatcttgtggaaaggacgaaacaccGTGAGGAACGTCTGCAAACGgtttaagagctatgctggaaacagc |
| Zc3h13 | Zc3h13-1 | ACATGACCGGCGTCACGAA | tatcttgtggaaaggacgaaacaccGACATGACCGGCGTCACGAAgtttaagagctatgctggaaacagc |
| Zc3h13 | Zc3h13-3 | GAAATTAGAAACGATCGGA | tatcttgtggaaaggacgaaacaccGGAAATTAGAAACGATCGGAgtttaagagctatgctggaaacagc |
| Zc3h13 | Zc3h13-5 | GCTCACCTATTCGCCATCA | tatcttgtggaaaggacgaaacaccGGCTCACCTATTCGCCATCAgtttaagagctatgctggaaacagc |
| Zfat | Zfat-2 | GACCGAGACCGCCACATGC | tatcttgtggaaaggacgaaacaccGGACCGAGACCGCCACATGCgtttaagagctatgctggaaacagc |
| Zfat | Zfat-3 | CGAGCTCCGCCTCATGACA | tatcttgtggaaaggacgaaacaccGCGAGCTCCGCCTCATGACAgtttaagagctatgctggaaacagc |
| Zfat | Zfat-5 | GTACCCTCTCCGCGGCACC | tatcttgtggaaaggacgaaacaccGGTACCCTCTCCGCGGCACCgtttaagagctatgctggaaacagc |
| Zfp131 | Zfp131-2 | TCTGACCCCATAAGCAAAC | tatcttgtggaaaggacgaaacaccGTCTGACCCCATAAGCAAACgtttaagagctatgctggaaacagc |
| Zfp131 | Zfp131-4 | CACCATTTTAAGGCCCACA | tatcttgtggaaaggacgaaacaccGCACCATTTTAAGGCCCACAgtttaagagctatgctggaaacagc |
| Zfp131 | Zfp131-5 | CGACCCACCGTCGACAATC | tatcttgtggaaaggacgaaacaccGCGACCCACCGTCGACAATCgtttaagagctatgctggaaacagc |
| Zfp160 | Zfp160-1 | CTGACAAAATTCTTGCTAG | tatcttgtggaaaggacgaaacaccGCTGACAAAATTCTTGCTAGgtttaagagctatgctggaaacagc |
| Zfp160 | Zfp160-2 | CATCATATTCATGAACAAC | tatcttgtggaaaggacgaaacaccGCATCATATTCATGAACAACgtttaagagctatgctggaaacagc |
| Zfp160 | Zfp160-5 | TCAAACCTAAATACCCATC | tatcttgtggaaaggacgaaacaccGTCAAACCTAAATACCCATCgtttaagagctatgctggaaacagc |
| Zfp35 | Zfp35-1 | AAAGCTTTGCTTGGAGTAC | tatcttgtggaaaggacgaaacaccGAAAGCTTTGCTTGGAGTACgtttaagagctatgctggaaacagc |
| Zfp35 | Zfp35-2 | ATACACCAACTACAAGGAT | tatcttgtggaaaggacgaaacaccGATACACCAACTACAAGGATgtttaagagctatgctggaaacagc |
| Zfp35 | Zfp35-4 | ACTAAAGCTTTTGGTACAC | tatcttgtggaaaggacgaaacaccGACTAAAGCTTTTGGTACACgtttaagagctatgctggaaacagc |
| Zfp459 | Zfp459-1 | TGGTGTATAAAAAGTGATG | tatcttgtggaaaggacgaaacaccGTGGTGTATAAAAAGTGATGgtttaagagctatgctggaaacagc |
| Zfp459 | Zfp459-3 | ATTAGTGGGCTTGCTGTAA | tatcttgtggaaaggacgaaacaccGATTAGTGGGCTTGCTGTAAgtttaagagctatgctggaaacagc |
| Zfp459 | Zfp459-4 | AAAAGCAGTGTTCCCAGGT | tatcttgtggaaaggacgaaacaccGAAAAGCAGTGTTCCCAGGTgtttaagagctatgctggaaacagc |
| Zfp472 | Zfp472-2 | TTTCACATTCTGTTATAGC | tatcttgtggaaaggacgaaacaccGTTTCACATTCTGTTATAGCgtttaagagctatgctggaaacagc |
| Zfp472 | Zfp472-3 | AGGGCTTCTGTCTCATGGG | tatcttgtggaaaggacgaaacaccGAGGGCTTCTGTCTCATGGGgtttaagagctatgctggaaacagc |
| Zfp472 | Zfp472-5 | CAGGTTCTGCGATCAATGA | tatcttgtggaaaggacgaaacaccGCAGGTTCTGCGATCAATGAgtttaagagctatgctggaaacagc |
| Zfp53 | Zfp53-1 | TCTCCATTAATCATGTCAC | tatcttgtggaaaggacgaaacaccGTCTCCATTAATCATGTCACgtttaagagctatgctggaaacagc |
| Zfp53 | Zfp53-2 | CTCTGATTTTCAGTAAGGT | tatcttgtggaaaggacgaaacaccGCTCTGATTTTCAGTAAGGTgtttaagagctatgctggaaacagc |
| Zfp53 | Zfp53-5 | TGTACAGTTAAGCTTGAAG | tatcttgtggaaaggacgaaacaccGTGTACAGTTAAGCTTGAAGgtttaagagctatgctggaaacagc |
| Zfp654 | Zfp654-1 | AAAGTGCAAATTTTGCCAA | tatcttgtggaaaggacgaaacaccGAAAGTGCAAATTTTGCCAAgtttaagagctatgctggaaacagc |
| Zfp654 | Zfp654-3 | GCATTCTCTGGTATAGCAA | tatcttgtggaaaggacgaaacaccGGCATTCTCTGGTATAGCAAgtttaagagctatgctggaaacagc |
| Zfp654 | Zfp654-5 | AATTCAAATACTAAGGGCT | tatcttgtggaaaggacgaaacaccGAATTCAAATACTAAGGGCTgtttaagagctatgctggaaacagc |
| Zfp708 | Zfp708-1 | AGCTTTTACTAATTATTCA | tatcttgtggaaaggacgaaacaccGAGCTTTTACTAATTATTCAgtttaagagctatgctggaaacagc |
| Zfp708 | Zfp708-2 | TGATGCTTAACAAGATAAT | tatcttgtggaaaggacgaaacaccGTGATGCTTAACAAGATAATgtttaagagctatgctggaaacagc |
| Zfp708 | Zfp708-6 | AGATCTGTGGAGTGTGAAG | tatcttgtggaaaggacgaaacaccGAGATCTGTGGAGTGTGAAGgtttaagagctatgctggaaacagc |
| Zfp948 | Zfp948-2 | ACAGTGCAAACAAAGAACA | tatcttgtggaaaggacgaaacaccGACAGTGCAAACAAAGAACAgtttaagagctatgctggaaacagc |
| Zfp948 | Zfp948-4 | ATAGAGCTAAGAAACATAC | tatcttgtggaaaggacgaaacaccGATAGAGCTAAGAAACATACgtttaagagctatgctggaaacagc |
| Zfp948 | Zfp948-5 | GCAAGTCTTTACCACAATC | tatcttgtggaaaggacgaaacaccGGCAAGTCTTTACCACAATCgtttaagagctatgctggaaacagc |
| Zfp959 | Zfp959-2 | TACATCAAAATACTGACCG | tatcttgtggaaaggacgaaacaccGTACATCAAAATACTGACCGgtttaagagctatgctggaaacagc |
| Zfp959 | Zfp959-3 | GGGTTAAGCTTTTACTACT | tatcttgtggaaaggacgaaacaccGGGGTTAAGCTTTTACTACTgtttaagagctatgctggaaacagc |
| Zfp959 | Zfp959-5 | AAGCATGACTATCATCAAT | tatcttgtggaaaggacgaaacaccGAAGCATGACTATCATCAATgtttaagagctatgctggaaacagc |
| Zfy2 | Zfy2-1 | TGTTACACCATGATGAATC | tatcttgtggaaaggacgaaacaccGTGTTACACCATGATGAATCgtttaagagctatgctggaaacagc |
| Zfy2 | Zfy2-4 | GATGATTTCTCTTTCTACT | tatcttgtggaaaggacgaaacaccGGATGATTTCTCTTTCTACTgtttaagagctatgctggaaacagc |
| Zfy2 | Zfy2-5 | AATGAGGTTGATGTAATAC | tatcttgtggaaaggacgaaacaccGAATGAGGTTGATGTAATACgtttaagagctatgctggaaacagc |
| Zmym5 | Zmym5-1 | ATCTGTAATAAATTAGGAG | tatcttgtggaaaggacgaaacaccGATCTGTAATAAATTAGGAGgtttaagagctatgctggaaacagc |

| | | | |
|---|---|---|---|
| Zmym5 | Zmym5-4 | AAAAGTAGTACCAAAAATG | tatcttgtggaaaggacgaaacaccGAAAAGTAGTACCAAAAATGgtttaagagctatgctggaaacagc |
| Zmym5 | Zmym5-5 | GGTCCCCATTCAATAAAGT | tatcttgtggaaaggacgaaacaccGGGTCCCCATTCAATAAAGTgtttaagagctatgctggaaacagc |
| Zpr1 | Zpr1-1 | GAAGTCGCGTCGTGCCCTA | tatcttgtggaaaggacgaaacaccGGAAGTCGCGTCGTGCCCTAgtttaagagctatgctggaaacagc |
| Zpr1 | Zpr1-2 | ACGATGATTTCTCTAAAGA | tatcttgtggaaaggacgaaacaccGACGATGATTTCTCTAAAGAgtttaagagctatgctggaaacagc |
| Zpr1 | Zpr1-3 | ACCACAAGGATCCCCGAGC | tatcttgtggaaaggacgaaacaccGACCACAAGGATCCCCGAGCgtttaagagctatgctggaaacagc |
| Zswim4 | Zswim4-1 | GCAAACCAGATGAGCGGTC | tatcttgtggaaaggacgaaacaccGGCAAACCAGATGAGCGGTCgtttaagagctatgctggaaacagc |
| Zswim4 | Zswim4-2 | TTCGAGCACGGGACTCCAA | tatcttgtggaaaggacgaaacaccGTTCGAGCACGGGACTCCAAgtttaagagctatgctggaaacagc |
| Zswim4 | Zswim4-3 | TGGATCGAGTGCTACAAGT | tatcttgtggaaaggacgaaacaccGTGGATCGAGTGCTACAAGTgtttaagagctatgctggaaacagc |
| Actr8 | Actr8-1 | GATGTCAAACGGTACAAGG | tatcttgtggaaaggacgaaacaccGGATGTCAAACGGTACAAGGgtttaagagctatgctggaaacagc |
| Actr8 | Actr8-4 | CTCATCGGAACACACGGTA | tatcttgtggaaaggacgaaacaccGCTCATCGGAACACACGGTAgtttaagagctatgctggaaacagc |
| Actr8 | Actr8-5 | GACACGTCAGAGCCCCCAT | tatcttgtggaaaggacgaaacaccGGACACGTCAGAGCCCCCATgtttaagagctatgctggaaacagc |
| Aplf | Aplf-2 | TTCCGCCAACATCCAAGCT | tatcttgtggaaaggacgaaacaccGTTCCGCCAACATCCAAGCTgtttaagagctatgctggaaacagc |
| Aplf | Aplf-3 | TATCAGGTAAGTTAACCAC | tatcttgtggaaaggacgaaacaccGTATCAGGTAAGTTAACCACgtttaagagctatgctggaaacagc |
| Aplf | Aplf-4 | AGCGTACTTGTCAAGTAAC | tatcttgtggaaaggacgaaacaccGAGCGTACTTGTCAAGTAACgtttaagagctatgctggaaacagc |
| Aptx | Aptx-1 | TAGGTTTACAAAGACGACC | tatcttgtggaaaggacgaaacaccGTAGGTTTACAAAGACGACCgtttaagagctatgctggaaacagc |
| Aptx | Aptx-3 | ATGACGCCCGAGTCAATGC | tatcttgtggaaaggacgaaacaccGATGACGCCCGAGTCAATGCgtttaagagctatgctggaaacagc |
| Aptx | Aptx-5 | TTGAAAGCAGAGTGTAACA | tatcttgtggaaaggacgaaacaccGTTGAAAGCAGAGTGTAACAgtttaagagctatgctggaaacagc |
| Ascc3 | Ascc3-2 | GACATAGCGGTCCTTACAG | tatcttgtggaaaggacgaaacaccGGACATAGCGGTCCTTACAGgtttaagagctatgctggaaacagc |
| Ascc3 | Ascc3-3 | TGCAGATATTTGGCCGAGC | tatcttgtggaaaggacgaaacaccGTGCAGATATTTGGCCGAGCgtttaagagctatgctggaaacagc |
| Ascc3 | Ascc3-4 | ACTGATGCCGTATGCTAGC | tatcttgtggaaaggacgaaacaccGACTGATGCCGTATGCTAGCgtttaagagctatgctggaaacagc |
| Aste1 | Aste1-2 | ATCCGAGAAGTGTTCATAC | tatcttgtggaaaggacgaaacaccGATCCGAGAAGTGTTCATACgtttaagagctatgctggaaacagc |
| Aste1 | Aste1-6 | GTTGAGATATAACCCCATC | tatcttgtggaaaggacgaaacaccGGTTGAGATATAACCCCATCgtttaagagctatgctggaaacagc |
| Aste1 | Aste1-7 | AGGTTGTACAACGGAAGCC | tatcttgtggaaaggacgaaacaccGAGGTTGTACAACGGAAGCCgtttaagagctatgctggaaacagc |
| Atr | Atr-3 | TGAATCGGCTTTACAGGTC | tatcttgtggaaaggacgaaacaccGTGAATCGGCTTTACAGGTCgtttaagagctatgctggaaacagc |
| Atr | Atr-4 | TCCAAGTTACTACTACTGG | tatcttgtggaaaggacgaaacaccGTCCAAGTTACTACTACTGGgtttaagagctatgctggaaacagc |
| Atr | Atr-5 | TCCTTTAACACAACTAGCC | tatcttgtggaaaggacgaaacaccGTCCTTTAACACAACTAGCCgtttaagagctatgctggaaacagc |
| Atrip | Atrip-1 | CCGTCTCCCACGTCAGGTA | tatcttgtggaaaggacgaaacaccGCCGTCTCCCACGTCAGGTAgtttaagagctatgctggaaacagc |
| Atrip | Atrip-2 | TGGTAACCCATCTAATCGG | tatcttgtggaaaggacgaaacaccGTGGTAACCCATCTAATCGGgtttaagagctatgctggaaacagc |
| Atrip | Atrip-4 | TTTACTGCGGACGACCTAG | tatcttgtggaaaggacgaaacaccGTTTACTGCGGACGACCTAGgtttaagagctatgctggaaacagc |
| Babam2 | Babam2-1 | ACATAGTTTTACCTTGAGA | tatcttgtggaaaggacgaaacaccGACATAGTTTTACCTTGAGAgtttaagagctatgctggaaacagc |
| Babam2 | Babam2-2 | AGGATGTAAATGAAGACCC | tatcttgtggaaaggacgaaacaccGAGGATGTAAATGAAGACCCgtttaagagctatgctggaaacagc |
| Babam2 | Babam2-5 | AATGAGACATCCTCCTCCG | tatcttgtggaaaggacgaaacaccGAATGAGACATCCTCCTCCGgtttaagagctatgctggaaacagc |
| Blm | Blm-1 | TTTGAGCGGTTGATACCAC | tatcttgtggaaaggacgaaacaccGTTTGAGCGGTTGATACCACgtttaagagctatgctggaaacagc |
| Blm | Blm-2 | TACCTGGAACATTTCAACG | tatcttgtggaaaggacgaaacaccGTACCTGGAACATTTCAACGgtttaagagctatgctggaaacagc |
| Blm | Blm-3 | GATTTAACGAAGGAATCGG | tatcttgtggaaaggacgaaacaccGGATTTAACGAAGGAATCGGgtttaagagctatgctggaaacagc |
| Brca2 | Brca2-2 | TACCCAAAGTCTCGTCAAG | tatcttgtggaaaggacgaaacaccGTACCCAAAGTCTCGTCAAGgtttaagagctatgctggaaacagc |
| Brca2 | Brca2-3 | AAGTACCGCTTCTAAGGAG | tatcttgtggaaaggacgaaacaccGAAGTACCGCTTCTAAGGAGgtttaagagctatgctggaaacagc |
| Brca2 | Brca2-4 | TGTACCCAATATGCGAGAG | tatcttgtggaaaggacgaaacaccGTGTACCCAATATGCGAGAGgtttaagagctatgctggaaacagc |
| Brd9 | Brd9-1 | GAGCGATCGTCGTAGTAGC | tatcttgtggaaaggacgaaacaccGGAGCGATCGTCGTAGTAGCgtttaagagctatgctggaaacagc |
| Brd9 | Brd9-2 | CGAGAGCACACCTATCCAG | tatcttgtggaaaggacgaaacaccGCGAGAGCACACCTATCCAGgtttaagagctatgctggaaacagc |
| Brd9 | Brd9-3 | GAGCAATTGCATCCGTAAC | tatcttgtggaaaggacgaaacaccGGAGCAATTGCATCCGTAACgtttaagagctatgctggaaacagc |
| Brip1 | Brip1-1 | CAACTCAAGTCGTGCTCAA | tatcttgtggaaaggacgaaacaccGCAACTCAAGTCGTGCTCAAgtttaagagctatgctggaaacagc |
| Brip1 | Brip1-4 | ACTTTCCTATCGGAGACTC | tatcttgtggaaaggacgaaacaccGACTTTCCTATCGGAGACTCgtttaagagctatgctggaaacagc |
| Brip1 | Brip1-5 | AGCTAGTCGTGCATCTACA | tatcttgtggaaaggacgaaacaccGAGCTAGTCGTGCATCTACAgtttaagagctatgctggaaacagc |
| Ccar2 | Ccar2-2 | GCAACGATAGAACTCCTCC | tatcttgtggaaaggacgaaacaccGGCAACGATAGAACTCCTCCgtttaagagctatgctggaaacagc |
| Ccar2 | Ccar2-4 | AACGCAAACAGCGGGCTGG | tatcttgtggaaaggacgaaacaccGAACGCAAACAGCGGGCTGGgtttaagagctatgctggaaacagc |
| Ccar2 | Ccar2-5 | GCCGCATACAACCCAGGCC | tatcttgtggaaaggacgaaacaccGGCCGCATACAACCCAGGCCgtttaagagctatgctggaaacagc |
| Cct8 | Cct8-1 | GTGCTGATTAAGACTGCCG | tatcttgtggaaaggacgaaacaccGGTGCTGATTAAGACTGCCGgtttaagagctatgctggaaacagc |
| Cct8 | Cct8-2 | CCGTACTGCTTACTCATGA | tatcttgtggaaaggacgaaacaccGCCGTACTGCTTACTCATGAgtttaagagctatgctggaaacagc |
| Cct8 | Cct8-3 | ATTGGCCTGTCAGTATCAG | tatcttgtggaaaggacgaaacaccGATTGGCCTGTCAGTATCAGgtttaagagctatgctggaaacagc |
| Chaf1a | Chaf1a-2 | ATCAAGCCAGTACCGAGTG | tatcttgtggaaaggacgaaacaccGATCAAGCCAGTACCGAGTGgtttaagagctatgctggaaacagc |
| Chaf1a | Chaf1a-3 | ACCCCCAGAACATCGGGGT | tatcttgtggaaaggacgaaacaccGACCCCCAGAACATCGGGGTgtttaagagctatgctggaaacagc |
| Chaf1a | Chaf1a-4 | ATACGACGAGCCAAGGAGG | tatcttgtggaaaggacgaaacaccGATACGACGAGCCAAGGAGGgtttaagagctatgctggaaacagc |
| Cnot7 | Cnot7-1 | TAGAGCTACTAACAACATC | tatcttgtggaaaggacgaaacaccGTAGAGCTACTAACAACATCgtttaagagctatgctggaaacagc |
| Cnot7 | Cnot7-4 | TACATTACACCGCAACAGT | tatcttgtggaaaggacgaaacaccGTACATTACACCGCAACAGTgtttaagagctatgctggaaacagc |
| Cnot7 | Cnot7-5 | TGGGTCTTGCAACAACGCC | tatcttgtggaaaggacgaaacaccGTGGGTCTTGCAACAACGCCgtttaagagctatgctggaaacagc |
| Dhx15 | Dhx15-1 | CACACCATAACGCTCCAGA | tatcttgtggaaaggacgaaacaccGCACACCATAACGCTCCAGAgtttaagagctatgctggaaacagc |
| Dhx15 | Dhx15-2 | GGGTCTGGTAAAACAACAC | tatcttgtggaaaggacgaaacaccGGGGTCTGGTAAAACAACACgtttaagagctatgctggaaacagc |
| Dhx15 | Dhx15-5 | GTGACCGAGAACGTGATAG | tatcttgtggaaaggacgaaacaccGGTGACCGAGAACGTGATAGgtttaagagctatgctggaaacagc |
| Dhx36 | Dhx36-1 | GCAGTTCCTTTATATACGC | tatcttgtggaaaggacgaaacaccGGCAGTTCCTTTATATACGCgtttaagagctatgctggaaacagc |
| Dhx36 | Dhx36-3 | TGGATAACTACATCGAAAG | tatcttgtggaaaggacgaaacaccGTGGATAACTACATCGAAAGgtttaagagctatgctggaaacagc |
| Dhx36 | Dhx36-5 | ACCTTAAGGGTCGCGAGAT | tatcttgtggaaaggacgaaacaccGACCTTAAGGGTCGCGAGATgtttaagagctatgctggaaacagc |
| Dmap1 | Dmap1-2 | GAAGCGCAGATCAAATCGG | tatcttgtggaaaggacgaaacaccGGAAGCGCAGATCAAATCGGgtttaagagctatgctggaaacagc |
| Dmap1 | Dmap1-3 | CATGATGACGCATGGACTA | tatcttgtggaaaggacgaaacaccGCATGATGACGCATGGACTAgtttaagagctatgctggaaacagc |
| Dmap1 | Dmap1-4 | TTCGAGCTGGGTTAGTAAA | tatcttgtggaaaggacgaaacaccGTTCGAGCTGGGTTAGTAAAgtttaagagctatgctggaaacagc |
| Dna2 | Dna2-1 | GGAGTGGATAACCGGTACC | tatcttgtggaaaggacgaaacaccGGGAGTGGATAACCGGTACCgtttaagagctatgctggaaacagc |
| Dna2 | Dna2-2 | AACTTCGTGCTCTCGGTCC | tatcttgtggaaaggacgaaacaccGAACTTCGTGCTCTCGGTCCgtttaagagctatgctggaaacagc |
| Dna2 | Dna2-5 | TGGTGTCTCATGTTAACGC | tatcttgtggaaaggacgaaacaccGTGGTGTCTCATGTTAACGCgtttaagagctatgctggaaacagc |
| Dntt | Dntt-1 | AACTCAGGGTCAGACGTCC | tatcttgtggaaaggacgaaacaccGAACTCAGGGTCAGACGTCCgtttaagagctatgctggaaacagc |
| Dntt | Dntt-3 | CATCGAATGCATGGGAGCT | tatcttgtggaaaggacgaaacaccGCATCGAATGCATGGGAGCTgtttaagagctatgctggaaacagc |
| Dntt | Dntt-5 | GGACGCTCGCATGAATGCC | tatcttgtggaaaggacgaaacaccGGGACGCTCGCATGAATGCCgtttaagagctatgctggaaacagc |
| Drosha | Drosha-2 | CAAGCACTACGACGACCAC | tatcttgtggaaaggacgaaacaccGCAAGCACTACGACGACCACgtttaagagctatgctggaaacagc |
| Drosha | Drosha-3 | CCGCTCAGACTATGATCGG | tatcttgtggaaaggacgaaacaccGCCGCTCAGACTATGATCGGgtttaagagctatgctggaaacagc |
| Drosha | Drosha-5 | TGTTGGTCATCGGACGGCA | tatcttgtggaaaggacgaaacaccGTGTTGGTCATCGGACGGCAgtttaagagctatgctggaaacagc |
| Eme1 | Eme1-2 | GGATGCTGGCGACCGGGAC | tatcttgtggaaaggacgaaacaccGGGATGCTGGCGACCGGGACgtttaagagctatgctggaaacagc |
| Eme1 | Eme1-3 | TTCAGACGGTCCAGAGCGG | tatcttgtggaaaggacgaaacaccGTTCAGACGGTCCAGAGCGGgtttaagagctatgctggaaacagc |

| | | | |
|---|---|---|---|
| Eme1 | Eme1-5 | TGTTCTTCGCCCTCTGAGG | tatcttgtggaaaggacgaaacaccGTGTTCTTCGCCCTCTGAGGgtttaagagctatgctggaaacagc |
| Endov | Endov-2 | CGTAGGTCGTTCTTGTGGA | tatcttgtggaaaggacgaaacaccGCGTAGGTCGTTCTTGTGGAgtttaagagctatgctggaaacagc |
| Endov | Endov-4 | GTGGGCCACAGAATAAGCC | tatcttgtggaaaggacgaaacaccGGTGGGCCACAGAATAAGCCgtttaagagctatgctggaaacagc |
| Endov | Endov-5 | ATCCCAGAACCTATACGCC | tatcttgtggaaaggacgaaacaccGATCCCAGAACCTATACGCCgtttaagagctatgctggaaacagc |
| Ep400 | Ep400-1 | TATAAGGCCTTCTGACGAC | tatcttgtggaaaggacgaaacaccGTATAAGGCCTTCTGACGACgtttaagagctatgctggaaacagc |
| Ep400 | Ep400-3 | CCAGTGAGATTTCGGGCGT | tatcttgtggaaaggacgaaacaccGCCAGTGAGATTTCGGGCGTgtttaagagctatgctggaaacagc |
| Ep400 | Ep400-4 | GCATCGGAACATGTAGGGC | tatcttgtggaaaggacgaaacaccGGCATCGGAACATGTAGGGCgtttaagagctatgctggaaacagc |
| Ercc1 | Ercc1-3 | GTGCCCTGGGAATTCGGTG | tatcttgtggaaaggacgaaacaccGGTGCCCTGGGAATTCGGTGgtttaagagctatgctggaaacagc |
| Ercc1 | Ercc1-4 | TTCATGGATGTAGTCTGGA | tatcttgtggaaaggacgaaacaccGTTCATGGATGTAGTCTGGAgtttaagagctatgctggaaacagc |
| Ercc1 | Ercc1-5 | AAGGGCGAAGTTCTTCCCC | tatcttgtggaaaggacgaaacaccGAAGGGCGAAGTTCTTCCCCgtttaagagctatgctggaaacagc |
| Ercc4 | Ercc4-2 | TTCATTTGTCACTCGTCGA | tatcttgtggaaaggacgaaacaccGTTCATTTGTCACTCGTCGAgtttaagagctatgctggaaacagc |
| Ercc4 | Ercc4-4 | GCAACAAGCCGAATACTCG | tatcttgtggaaaggacgaaacaccGGCAACAAGCCGAATACTCGgtttaagagctatgctggaaacagc |
| Ercc4 | Ercc4-5 | TCCGCCAGAAAAACAAGCG | tatcttgtggaaaggacgaaacaccGTCCGCCAGAAAAACAAGCGgtttaagagctatgctggaaacagc |
| Ewsr1 | Ewsr1-1 | CTGGAGGATTTTCCGGACC | tatcttgtggaaaggacgaaacaccGCTGGAGGATTTTCCGGACCgtttaagagctatgctggaaacagc |
| Ewsr1 | Ewsr1-2 | TCCTCAGGTTCATTCCGAC | tatcttgtggaaaggacgaaacaccGTCCTCAGGTTCATTCCGACgtttaagagctatgctggaaacagc |
| Ewsr1 | Ewsr1-3 | CTAGTTACCCCCCTCAGAC | tatcttgtggaaaggacgaaacaccGCTAGTTACCCCCCTCAGACgtttaagagctatgctggaaacagc |
| Exd2 | Exd2-1 | GTGGTGACGGTATCTCAGG | tatcttgtggaaaggacgaaacaccGGTGGTGACGGTATCTCAGGgtttaagagctatgctggaaacagc |
| Exd2 | Exd2-2 | CGCTACCTAGCCATGAAGC | tatcttgtggaaaggacgaaacaccGCGCTACCTAGCCATGAAGCgtttaagagctatgctggaaacagc |
| Exd2 | Exd2-4 | AAGTACGGACCAAATCAAC | tatcttgtggaaaggacgaaacaccGAAGTACGGACCAAATCAACgtttaagagctatgctggaaacagc |
| Exo1 | Exo1-2 | GCTAAGCCGATCCCACGAA | tatcttgtggaaaggacgaaacaccGGCTAAGCCGATCCCACGAAgtttaagagctatgctggaaacagc |
| Exo1 | Exo1-3 | ACCCCGATATCGTGAAGGT | tatcttgtggaaaggacgaaacaccGACCCCGATATCGTGAAGGTgtttaagagctatgctggaaacagc |
| Exo1 | Exo1-4 | CGTGTTCGACCCCATCCAA | tatcttgtggaaaggacgaaacaccGCGTGTTCGACCCCATCCAAgtttaagagctatgctggaaacagc |
| Exo5 | Exo5-2 | GCATCTTCTTTTGTGGCGA | tatcttgtggaaaggacgaaacaccGGCATCTTCTTTTGTGGCGAgtttaagagctatgctggaaacagc |
| Exo5 | Exo5-3 | TCTCAGGTGTCAACGAACC | tatcttgtggaaaggacgaaacaccGTCTCAGGTGTCAACGAACCgtttaagagctatgctggaaacagc |
| Exo5 | Exo5-5 | CTGTTTCACTTAGCAAGCC | tatcttgtggaaaggacgaaacaccGCTGTTTCACTTAGCAAGCCgtttaagagctatgctggaaacagc |
| Faap24 | Faap24-1 | ACCAAGTGGATAATGAGGC | tatcttgtggaaaggacgaaacaccGACCAAGTGGATAATGAGGCgtttaagagctatgctggaaacagc |
| Faap24 | Faap24-3 | AGTTTACTGTGCTAGACCT | tatcttgtggaaaggacgaaacaccGAGTTTACTGTGCTAGACCTgtttaagagctatgctggaaacagc |
| Faap24 | Faap24-4 | AATGCAAGATTTGCTAGAC | tatcttgtggaaaggacgaaacaccGAATGCAAGATTTGCTAGACgtttaagagctatgctggaaacagc |
| Fancf | Fancf-2 | TTGTGGACGCGCGGGCCGC | tatcttgtggaaaggacgaaacaccGTTGTGGACGCGCGGGCCGCgtttaagagctatgctggaaacagc |
| Fancf | Fancf-4 | CGCGAAGCGTCGGTACACG | tatcttgtggaaaggacgaaacaccGCGCGAAGCGTCGGTACACGgtttaagagctatgctggaaacagc |
| Fancf | Fancf-5 | GGACGCCGAAAAGGTGCGC | tatcttgtggaaaggacgaaacaccGGGACGCCGAAAAGGTGCGCgtttaagagctatgctggaaacagc |
| Fancl | Fancl-3 | ACATACTTATCCCAACCAA | tatcttgtggaaaggacgaaacaccGACATACTTATCCCAACCAAgtttaagagctatgctggaaacagc |
| Fancl | Fancl-4 | GAAAGTCCACAACACAATC | tatcttgtggaaaggacgaaacaccGGAAAGTCCACAACACAATCgtttaagagctatgctggaaacagc |
| Fancl | Fancl-5 | TGACTATAAACATCTACCA | tatcttgtggaaaggacgaaacaccGTGACTATAAACATCTACCAgtttaagagctatgctggaaacagc |
| Fen1 | Fen1-3 | GCGCGCCTCTCACTGCGCT | tatcttgtggaaaggacgaaacaccGGCGCGCCTCTCACTGCGCTgtttaagagctatgctggaaacagc |
| Fen1 | Fen1-4 | TACCGTACCATCCGCATGA | tatcttgtggaaaggacgaaacaccGTACCGTACCATCCGCATGAgtttaagagctatgctggaaacagc |
| Fen1 | Fen1-5 | GATTGCTGTTCGTCAGGGT | tatcttgtggaaaggacgaaacaccGGATTGCTGTTCGTCAGGGTgtttaagagctatgctggaaacagc |
| Fip1l1 | Fip1l1-1 | GCAGTGCTAATCCTCCATC | tatcttgtggaaaggacgaaacaccGGCAGTGCTAATCCTCCATCgtttaagagctatgctggaaacagc |
| Fip1l1 | Fip1l1-2 | GAAAATGGCGTGGCAAAAC | tatcttgtggaaaggacgaaacaccGGAAAATGGCGTGGCAAAACgtttaagagctatgctggaaacagc |
| Fip1l1 | Fip1l1-4 | ATAAACCATGGCGGAAACC | tatcttgtggaaaggacgaaacaccGATAAACCATGGCGGAAACCgtttaagagctatgctggaaacagc |
| Helb | Helb-1 | AGCGTGTAAGCCGGAAGAT | tatcttgtggaaaggacgaaacaccGAGCGTGTAAGCCGGAAGATgtttaagagctatgctggaaacagc |
| Helb | Helb-4 | ATATGCTCCGACAAGCCCG | tatcttgtggaaaggacgaaacaccGATATGCTCCGACAAGCCCGgtttaagagctatgctggaaacagc |
| Helb | Helb-5 | ACGGACGTTTCCCACTCAA | tatcttgtggaaaggacgaaacaccGACGGACGTTTCCCACTCAAgtttaagagctatgctggaaacagc |
| Hnrnpa1 | Hnrnpa1-3 | TTTAGGATTCTCAGCGACC | tatcttgtggaaaggacgaaacaccGTTTAGGATTCTCAGCGACCgtttaagagctatgctggaaacagc |
| Hnrnpa1 | Hnrnpa1-5 | ATCTCTTTTAAGGTCGCAG | tatcttgtggaaaggacgaaacaccGATCTCTTTTAAGGTCGCAGgtttaagagctatgctggaaacagc |
| Hspa1l | Hspa1l-1 | CTTCATTGATCACTTGAAA | tatcttgtggaaaggacgaaacaccGCTTCATTGATCACTTGAAAgtttaagagctatgctggaaacagc |
| Hspa1l | Hspa1l-3 | TCAAACCGTGCTCTAGTGA | tatcttgtggaaaggacgaaacaccGTCAAACCGTGCTCTAGTGAgtttaagagctatgctggaaacagc |
| Hspa1l | Hspa1l-4 | CAGGACTACTTTAATGGAC | tatcttgtggaaaggacgaaacaccGCAGGACTACTTTAATGGACgtttaagagctatgctggaaacagc |
| Huwe1 | Huwe1-1 | AACAGGGTCGTTGGTACCG | tatcttgtggaaaggacgaaacaccGAACAGGGTCGTTGGTACCGgtttaagagctatgctggaaacagc |
| Huwe1 | Huwe1-3 | ATCGGCCGATTGAATCCCC | tatcttgtggaaaggacgaaacaccGATCGGCCGATTGAATCCCCgtttaagagctatgctggaaacagc |
| Huwe1 | Huwe1-5 | CGATGTATACCAGCAGTAC | tatcttgtggaaaggacgaaacaccGCGATGTATACCAGCAGTACgtttaagagctatgctggaaacagc |
| Ing3 | Ing3-1 | AGCTGGAAGCCGATAATGC | tatcttgtggaaaggacgaaacaccGAGCTGGAAGCCGATAATGCgtttaagagctatgctggaaacagc |
| Ing3 | Ing3-3 | GAATTTCTTCTCAGGGATA | tatcttgtggaaaggacgaaacaccGGAATTTCTTCTCAGGGATAgtttaagagctatgctggaaacagc |
| Ing3 | Ing3-5 | ACCAATGTTATAGGAGGCC | tatcttgtggaaaggacgaaacaccGACCAATGTTATAGGAGGCCgtttaagagctatgctggaaacagc |
| Insl6 | Insl6-1 | GTTGTACGCTCTTAACATA | tatcttgtggaaaggacgaaacaccGGTTGTACGCTCTTAACATAgtttaagagctatgctggaaacagc |
| Insl6 | Insl6-2 | GATAGTCGGGCAGTGACTG | tatcttgtggaaaggacgaaacaccGGATAGTCGGGCAGTGACTGgtttaagagctatgctggaaacagc |
| Insl6 | Insl6-3 | GGGTTTGTGAATCTTCTCC | tatcttgtggaaaggacgaaacaccGGGGTTTGTGAATCTTCTCCgtttaagagctatgctggaaacagc |
| Kat7 | Kat7-1 | CAGCTGCGGTATAAGGAAA | tatcttgtggaaaggacgaaacaccGCAGCTGCGGTATAAGGAAAgtttaagagctatgctggaaacagc |
| Kat7 | Kat7-3 | ACATGAAGTGTCCTACGCC | tatcttgtggaaaggacgaaacaccGACATGAAGTGTCCTACGCCgtttaagagctatgctggaaacagc |
| Kat7 | Kat7-4 | CTCTCATCATGAGACACGT | tatcttgtggaaaggacgaaacaccGCTCTCATCATGAGACACGTgtttaagagctatgctggaaacagc |
| Kat8 | Kat8-1 | GTTCGCCACGCGCCGGAGT | tatcttgtggaaaggacgaaacaccGGTTCGCCACGCGCCGGAGTgtttaagagctatgctggaaacagc |
| Kat8 | Kat8-2 | CCGGCGGCCGGATAGCACC | tatcttgtggaaaggacgaaacaccGCCGGCGGCCGGATAGCACCgtttaagagctatgctggaaacagc |
| Kat8 | Kat8-4 | TTCGAGTGATCTTGCGCTC | tatcttgtggaaaggacgaaacaccGTTCGAGTGATCTTGCGCTCgtttaagagctatgctggaaacagc |
| Lig3 | Lig3-1 | CGGAAACAAATCTCCTCGG | tatcttgtggaaaggacgaaacaccGCGGAAACAAATCTCCTCGGgtttaagagctatgctggaaacagc |
| Lig3 | Lig3-4 | ATGGCCCGGGACCTAGAAC | tatcttgtggaaaggacgaaacaccGATGGCCCGGGACCTAGAACgtttaagagctatgctggaaacagc |
| Lig3 | Lig3-5 | TCGCGCAACCTGCAGGATG | tatcttgtggaaaggacgaaacaccGTCGCGCAACCTGCAGGATGgtttaagagctatgctggaaacagc |
| Mad2l2 | Mad2l2-1 | GTGCGCGAGGTCTACCCGG | tatcttgtggaaaggacgaaacaccGGTGCGCGAGGTCTACCCGGgtttaagagctatgctggaaacagc |
| Mad2l2 | Mad2l2-2 | GACGCAGTGGAGTGTGTCC | tatcttgtggaaaggacgaaacaccGGACGCAGTGGAGTGTGTCCgtttaagagctatgctggaaacagc |
| Mad2l2 | Mad2l2-4 | CTGTGCTTGCAGAACGATG | tatcttgtggaaaggacgaaacaccGCTGTGCTTGCAGAACGATGgtttaagagctatgctggaaacagc |
| Mcm10 | Mcm10-1 | GATGCTAAACGTTTGCCCC | tatcttgtggaaaggacgaaacaccGGATGCTAAACGTTTGCCCCgtttaagagctatgctggaaacagc |
| Mcm10 | Mcm10-3 | ACACCCGCACAGCCACTAC | tatcttgtggaaaggacgaaacaccGACACCCGCACAGCCACTACgtttaagagctatgctggaaacagc |
| Mcm10 | Mcm10-4 | CGCTTGGCTTTGGGTAAAG | tatcttgtggaaaggacgaaacaccGCGCTTGGCTTTGGGTAAAGgtttaagagctatgctggaaacagc |
| Mdc1 | Mdc1-2 | CTATAGTTAGAAGACTCCG | tatcttgtggaaaggacgaaacaccGCTATAGTTAGAAGACTCCGgtttaagagctatgctggaaacagc |
| Mdc1 | Mdc1-3 | AAGCTAACGGAACGACAGC | tatcttgtggaaaggacgaaacaccGAAGCTAACGGAACGACAGCgtttaagagctatgctggaaacagc |
| Mdc1 | Mdc1-5 | GTATCACCACTTGCTCCAC | tatcttgtggaaaggacgaaacaccGGTATCACCACTTGCTCCACgtttaagagctatgctggaaacagc |
| Mlh1 | Mlh1-2 | GAAGTTCACTTTCTGCACG | tatcttgtggaaaggacgaaacaccGGAAGTTCACTTTCTGCACGgtttaagagctatgctggaaacagc |

| | | | |
|---|---|---|---|
| Mlh1 | Mlh1-3 | CTGCCCAATGCCACAACCG | tatcttgtggaaaggacgaaacaccGCTGCCCAATGCCACAACCGgtttaagagctatgctggaaacagc |
| Mlh1 | Mlh1-4 | TGATGGGAAATGTGCGTAC | tatcttgtggaaaggacgaaacaccGTGATGGGAAATGTGCGTACgtttaagagctatgctggaaacagc |
| Mms19 | Mms19-2 | TACTCGCTCACTAGCAGTC | tatcttgtggaaaggacgaaacaccGTACTCGCTCACTAGCAGTCgtttaagagctatgctggaaacagc |
| Mms19 | Mms19-3 | GATCCCCGTAATCTCCTGC | tatcttgtggaaaggacgaaacaccGGATCCCCGTAATCTCCTGCgtttaagagctatgctggaaacagc |
| Mms19 | Mms19-4 | TGGCGGTCCACCTGTAGCA | tatcttgtggaaaggacgaaacaccGTGGCGGTCCACCTGTAGCAgtttaagagctatgctggaaacagc |
| Mms22l | Mms22l-2 | CGCTGTTTCTGTAACCCAT | tatcttgtggaaaggacgaaacaccGCGCTGTTTCTGTAACCCATgtttaagagctatgctggaaacagc |
| Mms22l | Mms22l-3 | ACGCTCCTACAGTAACACT | tatcttgtggaaaggacgaaacaccGACGCTCCTACAGTAACACTgtttaagagctatgctggaaacagc |
| Mms22l | Mms22l-4 | TATTTAGATACCCACTGGC | tatcttgtggaaaggacgaaacaccGTATTTAGATACCCACTGGCgtttaagagctatgctggaaacagc |
| Msh5 | Msh5-1 | ATTGCCAGCCTCATCGGGA | tatcttgtggaaaggacgaaacaccGATTGCCAGCCTCATCGGGAgtttaagagctatgctggaaacagc |
| Msh5 | Msh5-4 | GGCTACTTTGTACACCGAG | tatcttgtggaaaggacgaaacaccGGGCTACTTTGTACACCGAGgtttaagagctatgctggaaacagc |
| Msh5 | Msh5-5 | GTCATAGTAAGCAATGCCC | tatcttgtggaaaggacgaaacaccGGTCATAGTAAGCAATGCCCgtttaagagctatgctggaaacagc |
| Mus81 | Mus81-1 | CGCTAGGCCTTAGCTCCAG | tatcttgtggaaaggacgaaacaccGCGCTAGGCCTTAGCTCCAGgtttaagagctatgctggaaacagc |
| Mus81 | Mus81-2 | CATTATAGTGGCACCACCG | tatcttgtggaaaggacgaaacaccGCATTATAGTGGCACCACCGgtttaagagctatgctggaaacagc |
| Mus81 | Mus81-5 | ACGCGTTTCGTGTTTCAAA | tatcttgtggaaaggacgaaacaccGACGCGTTTCGTGTTTCAAAgtttaagagctatgctggaaacagc |
| Palb2 | Palb2-1 | TGACAATCTGACATACGAC | tatcttgtggaaaggacgaaacaccGTGACAATCTGACATACGACgtttaagagctatgctggaaacagc |
| Palb2 | Palb2-4 | GTATATTAATTCCGCCATC | tatcttgtggaaaggacgaaacaccGGTATATTAATTCCGCCATCgtttaagagctatgctggaaacagc |
| Palb2 | Palb2-5 | CCCTTTTCGATGCTTACCA | tatcttgtggaaaggacgaaacaccGCCCTTTTCGATGCTTACCAgtttaagagctatgctggaaacagc |
| Parg | Parg-1 | TCCCACGTACCAAACGAGG | tatcttgtggaaaggacgaaacaccGTCCCACGTACCAAACGAGGgtttaagagctatgctggaaacagc |
| Parg | Parg-3 | CGGCACCAACATCTGACAA | tatcttgtggaaaggacgaaacaccGCGGCACCAACATCTGACAAgtttaagagctatgctggaaacagc |
| Parg | Parg-5 | GCAAATGGATCCCACACCG | tatcttgtggaaaggacgaaacaccGGCAAATGGATCCCACACCGgtttaagagctatgctggaaacagc |
| Parp1 | Parp1-1 | GGGACTTTCCCATCGAACA | tatcttgtggaaaggacgaaacaccGGGGACTTTCCCATCGAACAgtttaagagctatgctggaaacagc |
| Parp1 | Parp1-4 | GTCGCTCTTAAAGACCAGC | tatcttgtggaaaggacgaaacaccGGTCGCTCTTAAAGACCAGCgtttaagagctatgctggaaacagc |
| Parp1 | Parp1-5 | TCACACACAACTCGAACGT | tatcttgtggaaaggacgaaacaccGTCACACACAACTCGAACGTgtttaagagctatgctggaaacagc |
| Parp2 | Parp2-1 | AAATACGACATGTTACAGA | tatcttgtggaaaggacgaaacaccGAAATACGACATGTTACAGAgtttaagagctatgctggaaacagc |
| Parp2 | Parp2-3 | CTGACAGTGGCGCAAATCA | tatcttgtggaaaggacgaaacaccGCTGACAGTGGCGCAAATCAgtttaagagctatgctggaaacagc |
| Parp2 | Parp2-5 | AACAAGCGCTCGCCCATGC | tatcttgtggaaaggacgaaacaccGAACAAGCGCTCGCCCATGCgtttaagagctatgctggaaacagc |
| Paxip1 | Paxip1-3 | ATAGATGATCAGGCGTGGA | tatcttgtggaaaggacgaaacaccGATAGATGATCAGGCGTGGAgtttaagagctatgctggaaacagc |
| Paxip1 | Paxip1-4 | TAAGATTGTCACCCCGGAC | tatcttgtggaaaggacgaaacaccGTAAGATTGTCACCCCGGACgtttaagagctatgctggaaacagc |
| Paxip1 | Paxip1-5 | CCCCCGTGGAACGTAACCA | tatcttgtggaaaggacgaaacaccGCCCCCGTGGAACGTAACCAgtttaagagctatgctggaaacagc |
| Paxx | Paxx-2 | AACTCACCTGAACCGGGAA | tatcttgtggaaaggacgaaacaccGAACTCACCTGAACCGGGAAgtttaagagctatgctggaaacagc |
| Paxx | Paxx-3 | AAGCCCGATTTGGCCTAAG | tatcttgtggaaaggacgaaacaccGAAGCCCGATTTGGCCTAAGgtttaagagctatgctggaaacagc |
| Paxx | Paxx-4 | AGGCTGTCAGGCGAGAAGC | tatcttgtggaaaggacgaaacaccGAGGCTGTCAGGCGAGAAGCgtttaagagctatgctggaaacagc |
| Pms2 | Pms2-2 | GCAAGCGTTGGGACTCGAC | tatcttgtggaaaggacgaaacaccGGCAAGCGTTGGGACTCGACgtttaagagctatgctggaaacagc |
| Pms2 | Pms2-3 | CACTAATCAGCTCGGACAG | tatcttgtggaaaggacgaaacaccGCACTAATCAGCTCGGACAGgtttaagagctatgctggaaacagc |
| Pms2 | Pms2-5 | AACGTTAAGGACGACAAAT | tatcttgtggaaaggacgaaacaccGAACGTTAAGGACGACAAATgtttaagagctatgctggaaacagc |
| Pola1 | Pola1-3 | TAACGTTTACCATTTCACG | tatcttgtggaaaggacgaaacaccGTAACGTTTACCATTTCACGgtttaagagctatgctggaaacagc |
| Pola1 | Pola1-4 | TCTTTGGTATCAATACAGG | tatcttgtggaaaggacgaaacaccGTCTTTGGTATCAATACAGGgtttaagagctatgctggaaacagc |
| Pola1 | Pola1-5 | GAAGAGCAGTATTCGAAAC | tatcttgtggaaaggacgaaacaccGGAAGAGCAGTATTCGAAACgtttaagagctatgctggaaacagc |
| Pola2 | Pola2-1 | CGCGCCATAGGAGTGCTCC | tatcttgtggaaaggacgaaacaccGCGCGCCATAGGAGTGCTCCgtttaagagctatgctggaaacagc |
| Pola2 | Pola2-2 | CAAGTCCGTGATTCTCGAG | tatcttgtggaaaggacgaaacaccGCAAGTCCGTGATTCTCGAGgtttaagagctatgctggaaacagc |
| Pola2 | Pola2-5 | ACCGTCAAGTTTCTCCCCG | tatcttgtggaaaggacgaaacaccGACCGTCAAGTTTCTCCCCGgtttaagagctatgctggaaacagc |
| Pold2 | Pold2-1 | TGCTACTGGTATCCGGACT | tatcttgtggaaaggacgaaacaccGTGCTACTGGTATCCGGACTgtttaagagctatgctggaaacagc |
| Pold2 | Pold2-4 | ATAAATATGGGCGTACTGC | tatcttgtggaaaggacgaaacaccGATAAATATGGGCGTACTGCgtttaagagctatgctggaaacagc |
| Pold2 | Pold2-5 | TGCGCTCTCCCAGTCGGAA | tatcttgtggaaaggacgaaacaccGTGCGCTCTCCCAGTCGGAAgtttaagagctatgctggaaacagc |
| Pold3 | Pold3-1 | TCTGTCACTGAACCAAAGC | tatcttgtggaaaggacgaaacaccGTCTGTCACTGAACCAAAGCgtttaagagctatgctggaaacagc |
| Pold3 | Pold3-4 | CTGTATGGCACTAAACCTA | tatcttgtggaaaggacgaaacaccGCTGTATGGCACTAAACCTAgtttaagagctatgctggaaacagc |
| Pold3 | Pold3-5 | AGCTATGCTAAAGGACAGT | tatcttgtggaaaggacgaaacaccGAGCTATGCTAAAGGACAGTgtttaagagctatgctggaaacagc |
| Pold4 | Pold4-1 | CTCCAGCTCTGTTTCCTCC | tatcttgtggaaaggacgaaacaccGCTCCAGCTCTGTTTCCTCCgtttaagagctatgctggaaacagc |
| Pold4 | Pold4-2 | CTGCTGAGGCAGTTTGACC | tatcttgtggaaaggacgaaacaccGCTGCTGAGGCAGTTTGACCgtttaagagctatgctggaaacagc |
| Pold4 | Pold4-5 | CCTGATGGCAGGTATCACA | tatcttgtggaaaggacgaaacaccGCCTGATGGCAGGTATCACAgtttaagagctatgctggaaacagc |
| Pole | Pole-2 | CTCGTCGGGTGATTTCCAC | tatcttgtggaaaggacgaaacaccGCTCGTCGGGTGATTTCCACgtttaagagctatgctggaaacagc |
| Pole | Pole-3 | TATACCATCCTAAACCGGC | tatcttgtggaaaggacgaaacaccGTATACCATCCTAAACCGGCgtttaagagctatgctggaaacagc |
| Pole | Pole-5 | ATTGGACACGGACGGAATA | tatcttgtggaaaggacgaaacaccGATTGGACACGGACGGAATAgtttaagagctatgctggaaacagc |
| Pole2 | Pole2-2 | CACAGTGGCTTATACACCG | tatcttgtggaaaggacgaaacaccGCACAGTGGCTTATACACCGgtttaagagctatgctggaaacagc |
| Pole2 | Pole2-3 | CCTACTGGAACAGTACAAC | tatcttgtggaaaggacgaaacaccGCCTACTGGAACAGTACAACgtttaagagctatgctggaaacagc |
| Pole2 | Pole2-5 | ATCGAACGATCTGTCGTGG | tatcttgtggaaaggacgaaacaccGATCGAACGATCTGTCGTGGgtttaagagctatgctggaaacagc |
| Polh | Polh-1 | ACACGATCCAGTTAAACCC | tatcttgtggaaaggacgaaacaccGACACGATCCAGTTAAACCCgtttaagagctatgctggaaacagc |
| Polh | Polh-4 | ATATAAGCCTCATCGATGC | tatcttgtggaaaggacgaaacaccGATATAAGCCTCATCGATGCgtttaagagctatgctggaaacagc |
| Polh | Polh-5 | GATCCTTATACTTAGGTAC | tatcttgtggaaaggacgaaacaccGGATCCTTATACTTAGGTACgtttaagagctatgctggaaacagc |
| Polk | Polk-1 | GGTCTAGGTTCAACAGACC | tatcttgtggaaaggacgaaacaccGGGTCTAGGTTCAACAGACCgtttaagagctatgctggaaacagc |
| Polk | Polk-2 | TATTTTCTTCACATCGCGC | tatcttgtggaaaggacgaaacaccGTATTTTCTTCACATCGCGCgtttaagagctatgctggaaacagc |
| Polk | Polk-4 | GCCTCTTAGCAATAAATCC | tatcttgtggaaaggacgaaacaccGGCCTCTTAGCAATAAATCCgtttaagagctatgctggaaacagc |
| Poln | Poln-1 | ATACTCTACGACCTTACAA | tatcttgtggaaaggacgaaacaccGATACTCTACGACCTTACAAgtttaagagctatgctggaaacagc |
| Poln | Poln-2 | GTGGTGACATTGATGTACA | tatcttgtggaaaggacgaaacaccGGTGGTGACATTGATGTACAgtttaagagctatgctggaaacagc |
| Poln | Poln-5 | CTGGGTTATTAATGACTCG | tatcttgtggaaaggacgaaacaccGCTGGGTTATTAATGACTCGgtttaagagctatgctggaaacagc |
| Rad1 | Rad1-1 | GCGTGTTCTCTGAAATGAA | tatcttgtggaaaggacgaaacaccGGCGTGTTCTCTGAAATGAAgtttaagagctatgctggaaacagc |
| Rad1 | Rad1-2 | AACGGAATCAAGGTTACAG | tatcttgtggaaaggacgaaacaccGAACGGAATCAAGGTTACAGgtttaagagctatgctggaaacagc |
| Rad1 | Rad1-5 | CGCTTCGGATGTGTTACCA | tatcttgtggaaaggacgaaacaccGCGCTTCGGATGTGTTACCAgtttaagagctatgctggaaacagc |
| Rad51c | Rad51c-1 | AGTTTTATGGTTGATAGAG | tatcttgtggaaaggacgaaacaccGAGTTTTATGGTTGATAGAGgtttaagagctatgctggaaacagc |
| Rad51c | Rad51c-3 | TTTCGTGGCAGTATGCAAT | tatcttgtggaaaggacgaaacaccGTTTCGTGGCAGTATGCAATgtttaagagctatgctggaaacagc |
| Rad51c | Rad51c-5 | GAAGTTGGGATATCTAAAG | tatcttgtggaaaggacgaaacaccGGAAGTTGGGATATCTAAAGgtttaagagctatgctggaaacagc |
| Rad52 | Rad52-3 | GGTTCCTATCATGAGGACG | tatcttgtggaaaggacgaaacaccGGGTTCCTATCATGAGGACGgtttaagagctatgctggaaacagc |
| Rad52 | Rad52-4 | CTAAATAAGCTTCCACGAC | tatcttgtggaaaggacgaaacaccGCTAAATAAGCTTCCACGACgtttaagagctatgctggaaacagc |
| Rad52 | Rad52-5 | TAGTTAAATCCACATCAAG | tatcttgtggaaaggacgaaacaccGTAGTTAAATCCACATCAAGgtttaagagctatgctggaaacagc |
| Rbbp6 | Rbbp6-4 | TCAGGAGGATTCCTATTGG | tatcttgtggaaaggacgaaacaccGTCAGGAGGATTCCTATTGGgtttaagagctatgctggaaacagc |
| Rbbp6 | Rbbp6-5 | TAGACTTGACACCTCCAAT | tatcttgtggaaaggacgaaacaccGTAGACTTGACACCTCCAATgtttaagagctatgctggaaacagc |

| | | | |
|---|---|---|---|
| Rbbp6 | Rbbp6-7 | TACTACAAAGGGTACGCGG | tatcttgtggaaaggacgaaacaccGTACTACAAAGGGTACGCGGgtttaagagctatgctggaaacagc |
| Rfc1 | Rfc1-3 | AACTTGTTTTAGGCCCGAA | tatcttgtggaaaggacgaaacaccGAACTTGTTTTAGGCCCGAAgtttaagagctatgctggaaacagc |
| Rfc1 | Rfc1-4 | AATAGTCGAGCACTGACGT | tatcttgtggaaaggacgaaacaccGAATAGTCGAGCACTGACGTgtttaagagctatgctggaaacagc |
| Rfc1 | Rfc1-5 | TTTACCGGAGTTATTGACC | tatcttgtggaaaggacgaaacaccGTTTACCGGAGTTATTGACCgtttaagagctatgctggaaacagc |
| Rfc2 | Rfc2-1 | TTGAAGCTCAACGAAATAG | tatcttgtggaaaggacgaaacaccGTTGAAGCTCAACGAAATAGgtttaagagctatgctggaaacagc |
| Rfc2 | Rfc2-2 | ACTCAATGCCTCAAATGAC | tatcttgtggaaaggacgaaacaccGACTCAATGCCTCAAATGACgtttaagagctatgctggaaacagc |
| Rfc2 | Rfc2-3 | TTGTGCCGGCCCTTGGGAA | tatcttgtggaaaggacgaaacaccGTTGTGCCGGCCCTTGGGAAgtttaagagctatgctggaaacagc |
| Rfc3 | Rfc3-1 | GATACCCCCGATCCGAAGC | tatcttgtggaaaggacgaaacaccGGATACCCCCGATCCGAAGCgtttaagagctatgctggaaacagc |
| Rfc3 | Rfc3-4 | GAACTTTATGGTATCGGGG | tatcttgtggaaaggacgaaacaccGGAACTTTATGGTATCGGGGgtttaagagctatgctggaaacagc |
| Rfc3 | Rfc3-5 | TGATGGTCCATATACTAAG | tatcttgtggaaaggacgaaacaccGTGATGGTCCATATACTAAGgtttaagagctatgctggaaacagc |
| Rfc4 | Rfc4-2 | TTTTGAACATCTAGAAGTG | tatcttgtggaaaggacgaaacaccGTTTTGAACATCTAGAAGTGgtttaagagctatgctggaaacagc |
| Rfc4 | Rfc4-3 | GCAGCTTTAAGGCGTACCA | tatcttgtggaaaggacgaaacaccGGCAGCTTTAAGGCGTACCAgtttaagagctatgctggaaacagc |
| Rfc4 | Rfc4-4 | CAATCTTAAAGGGAGGACA | tatcttgtggaaaggacgaaacaccGCAATCTTAAAGGGAGGACAgtttaagagctatgctggaaacagc |
| Rfc5 | Rfc5-2 | GGACAGGTAGTTACAGATG | tatcttgtggaaaggacgaaacaccGGGACAGGTAGTTACAGATGgtttaagagctatgctggaaacagc |
| Rfc5 | Rfc5-3 | CTGTGAACTTCTCAATCAC | tatcttgtggaaaggacgaaacaccGCTGTGAACTTCTCAATCACgtttaagagctatgctggaaacagc |
| Rfc5 | Rfc5-5 | GAATGCTTCTGACGACCGA | tatcttgtggaaaggacgaaacaccGGAATGCTTCTGACGACCGAgtttaagagctatgctggaaacagc |
| Rif1 | Rif1-1 | TCCTTTAAAGTCGTATGAC | tatcttgtggaaaggacgaaacaccGTCCTTTAAAGTCGTATGACgtttaagagctatgctggaaacagc |
| Rif1 | Rif1-2 | AACCGTGTACAGCCGAGAA | tatcttgtggaaaggacgaaacaccGAACCGTGTACAGCCGAGAAgtttaagagctatgctggaaacagc |
| Rif1 | Rif1-4 | TGATTAGCGGGTGTTCCAG | tatcttgtggaaaggacgaaacaccGTGATTAGCGGGTGTTCCAGgtttaagagctatgctggaaacagc |
| Rnaseh1 | Rnaseh1-1 | AGCAGGAAACCGGTCCACC | tatcttgtggaaaggacgaaacaccGAGCAGGAAACCGGTCCACCgtttaagagctatgctggaaacagc |
| Rnaseh1 | Rnaseh1-4 | CAGTCGTTGTCTACACGGA | tatcttgtggaaaggacgaaacaccGCAGTCGTTGTCTACACGGAgtttaagagctatgctggaaacagc |
| Rnaseh1 | Rnaseh1-5 | CGTGTTTACTTACAAGGGG | tatcttgtggaaaggacgaaacaccGCGTGTTTACTTACAAGGGGgtttaagagctatgctggaaacagc |
| Rnf138 | Rnf138-1 | AAGACGCCGGTGCGGACCG | tatcttgtggaaaggacgaaacaccGAAGACGCCGGTGCGGACCGgtttaagagctatgctggaaacagc |
| Rnf138 | Rnf138-4 | GAAAGAGCATGTCCGGAAC | tatcttgtggaaaggacgaaacaccGGAAAGAGCATGTCCGGAACgtttaagagctatgctggaaacagc |
| Rnf138 | Rnf138-5 | GATGCTGTTCAAAAAAGGT | tatcttgtggaaaggacgaaacaccGGATGCTGTTCAAAAAAGGTgtttaagagctatgctggaaacagc |
| Rpa3 | Rpa3-1 | CTGGGATTGTAGAAGTAGT | tatcttgtggaaaggacgaaacaccGCTGGGATTGTAGAAGTAGTgtttaagagctatgctggaaacagc |
| Rpa3 | Rpa3-2 | TTCAATGTTTAGCTTGACG | tatcttgtggaaaggacgaaacaccGTTCAATGTTTAGCTTGACGgtttaagagctatgctggaaacagc |
| Rpa3 | Rpa3-3 | AATGGAACCATTGAATTGA | tatcttgtggaaaggacgaaacaccGAATGGAACCATTGAATTGAgtttaagagctatgctggaaacagc |
| Ruvbl2 | Ruvbl2-2 | CTCGATCATCTTGGTACCC | tatcttgtggaaaggacgaaacaccGCTCGATCATCTTGGTACCCgtttaagagctatgctggaaacagc |
| Ruvbl2 | Ruvbl2-3 | ATGGAGACCATCTACGACC | tatcttgtggaaaggacgaaacaccGATGGAGACCATCTACGACCgtttaagagctatgctggaaacagc |
| Ruvbl2 | Ruvbl2-5 | CCGGACACCAATGGAACGG | tatcttgtggaaaggacgaaacaccGCCGGACACCAATGGAACGGgtttaagagctatgctggaaacagc |
| Sem1 | Sem1-1 | ACATGTCTGGGAGGATAAT | tatcttgtggaaaggacgaaacaccGACATGTCTGGGAGGATAATgtttaagagctatgctggaaacagc |
| Sem1 | Sem1-2 | AGATGAAGATGCACATGTC | tatcttgtggaaaggacgaaacaccGAGATGAAGATGCACATGTCgtttaagagctatgctggaaacagc |
| Setmar | Setmar-2 | TTCGATGCAAGCACATCCA | tatcttgtggaaaggacgaaacaccGTTCGATGCAAGCACATCCAgtttaagagctatgctggaaacagc |
| Setmar | Setmar-3 | TTCGGATCCCACATCCCTA | tatcttgtggaaaggacgaaacaccGTTCGGATCCCACATCCCTAgtttaagagctatgctggaaacagc |
| Setmar | Setmar-5 | GGCATGCGCTGCAGAAATA | tatcttgtggaaaggacgaaacaccGGGCATGCGCTGCAGAAATAgtttaagagctatgctggaaacagc |
| Sgo1 | Sgo1-1 | TGTAAAGCCAGTCCAAGAG | tatcttgtggaaaggacgaaacaccGTGTAAAGCCAGTCCAAGAGgtttaagagctatgctggaaacagc |
| Sgo1 | Sgo1-2 | ATCTTTCCTTAAATGGCGA | tatcttgtggaaaggacgaaacaccGATCTTTCCTTAAATGGCGAgtttaagagctatgctggaaacagc |
| Sgo1 | Sgo1-4 | AAAGGACGTCCCTCAGACG | tatcttgtggaaaggacgaaacaccGAAAGGACGTCCCTCAGACGgtttaagagctatgctggaaacagc |
| Shld1 | Shld1-2 | CTTCTTCGACTACTGATCC | tatcttgtggaaaggacgaaacaccGCTTCTTCGACTACTGATCCgtttaagagctatgctggaaacagc |
| Shld1 | Shld1-4 | GCTTCATAGAATCTATCCA | tatcttgtggaaaggacgaaacaccGGCTTCATAGAATCTATCCAgtttaagagctatgctggaaacagc |
| Shld1 | Shld1-5 | AGACAATCTCTGAGCTCGA | tatcttgtggaaaggacgaaacaccGAGACAATCTCTGAGCTCGAgtttaagagctatgctggaaacagc |
| Slc19a1 | Slc19a1-2 | GCCATGCGCTGGTAGCGGG | tatcttgtggaaaggacgaaacaccGGCCATGCGCTGGTAGCGGGgtttaagagctatgctggaaacagc |
| Slc19a1 | Slc19a1-3 | TCACGGGCGGCAGTACTGC | tatcttgtggaaaggacgaaacaccGTCACGGGCGGCAGTACTGCgtttaagagctatgctggaaacagc |
| Slc19a1 | Slc19a1-4 | ACCGCAGTACGCTTGCGCG | tatcttgtggaaaggacgaaacaccGACCGCAGTACGCTTGCGCGgtttaagagctatgctggaaacagc |
| Slx4 | Slx4-2 | GCGTTCCCTGGCCCTAAGG | tatcttgtggaaaggacgaaacaccGGCGTTCCCTGGCCCTAAGGgtttaagagctatgctggaaacagc |
| Slx4 | Slx4-3 | GACAAACCCCGTCAGCGGA | tatcttgtggaaaggacgaaacaccGGACAAACCCCGTCAGCGGAgtttaagagctatgctggaaacagc |
| Slx4 | Slx4-4 | TCGGGGCGTACCCACAGGT | tatcttgtggaaaggacgaaacaccGTCGGGGCGTACCCACAGGTgtttaagagctatgctggaaacagc |
| Smarcal1 | Smarcal1-2 | ATCGCTTCCGGGTAAAGAT | tatcttgtggaaaggacgaaacaccGATCGCTTCCGGGTAAAGATgtttaagagctatgctggaaacagc |
| Smarcal1 | Smarcal1-3 | GACATCGGCTATTCGGAGG | tatcttgtggaaaggacgaaacaccGGACATCGGCTATTCGGAGGgtttaagagctatgctggaaacagc |
| Smarcal1 | Smarcal1-4 | GAAGCGGACCTTTCCGGAG | tatcttgtggaaaggacgaaacaccGGAAGCGGACCTTTCCGGAGgtttaagagctatgctggaaacagc |
| Smc1a | Smc1a-1 | GACCGCTATCAGCGCCTGA | tatcttgtggaaaggacgaaacaccGGACCGCTATCAGCGCCTGAgtttaagagctatgctggaaacagc |
| Smc1a | Smc1a-2 | GATGAGGTAGTACGAGCAC | tatcttgtggaaaggacgaaacaccGGATGAGGTAGTACGAGCACgtttaagagctatgctggaaacagc |
| Smc1a | Smc1a-5 | GATGCACGAATTGACCGAC | tatcttgtggaaaggacgaaacaccGGATGCACGAATTGACCGACgtttaagagctatgctggaaacagc |
| Ssbp1 | Ssbp1-1 | TTACTTGGACGAGTAGGTC | tatcttgtggaaaggacgaaacaccGTTACTTGGACGAGTAGGTCgtttaagagctatgctggaaacagc |
| Ssbp1 | Ssbp1-2 | TTCCACCTGTCTCATGACA | tatcttgtggaaaggacgaaacaccGTTCCACCTGTCTCATGACAgtttaagagctatgctggaaacagc |
| Ssbp1 | Ssbp1-3 | AAATGAGATGTGGCGATCA | tatcttgtggaaaggacgaaacaccGAAATGAGATGTGGCGATCAgtttaagagctatgctggaaacagc |
| Ten1 | Ten1-2 | GCACGCTCCCTCATGACCC | tatcttgtggaaaggacgaaacaccGGCACGCTCCCTCATGACCCgtttaagagctatgctggaaacagc |
| Ten1 | Ten1-3 | CAAGCAGTTGACACTGATC | tatcttgtggaaaggacgaaacaccGCAAGCAGTTGACACTGATCgtttaagagctatgctggaaacagc |
| Ten1 | Ten1-4 | CAAGATGGCTTTCTCTAAC | tatcttgtggaaaggacgaaacaccGCAAGATGGCTTTCTCTAACgtttaagagctatgctggaaacagc |
| Terf1 | Terf1-2 | ACGACGAAAGTCCTCGGAG | tatcttgtggaaaggacgaaacaccGACGACGAAAGTCCTCGGAGgtttaagagctatgctggaaacagc |
| Terf1 | Terf1-3 | CGTACTCGTGACAGCGCCG | tatcttgtggaaaggacgaaacaccGCGTACTCGTGACAGCGCCGgtttaagagctatgctggaaacagc |
| Terf1 | Terf1-5 | GATGAGCGTATTACACCCT | tatcttgtggaaaggacgaaacaccGGATGAGCGTATTACACCCTgtttaagagctatgctggaaacagc |
| Timeless | Timeless-1 | ATCCGATACCTGAGGCACG | tatcttgtggaaaggacgaaacaccGATCCGATACCTGAGGCACGgtttaagagctatgctggaaacagc |
| Timeless | Timeless-2 | ATCCTCACGCAGCATCGCC | tatcttgtggaaaggacgaaacaccGATCCTCACGCAGCATCGCCgtttaagagctatgctggaaacagc |
| Timeless | Timeless-5 | CGCATCAAAGGGAACGACG | tatcttgtggaaaggacgaaacaccGCGCATCAAAGGGAACGACGgtttaagagctatgctggaaacagc |
| Trex1 | Trex1-2 | GACAGCATCGCTGCCCTAA | tatcttgtggaaaggacgaaacaccGGACAGCATCGCTGCCCTAAgtttaagagctatgctggaaacagc |
| Trex1 | Trex1-3 | AAGGTACCATCTAGGGGAC | tatcttgtggaaaggacgaaacaccGAAGGTACCATCTAGGGGACgtttaagagctatgctggaaacagc |
| Trex1 | Trex1-4 | TCCACCACACGGGGCGGTC | tatcttgtggaaaggacgaaacaccGTCCACCACACGGGGCGGTCgtttaagagctatgctggaaacagc |
| Trex2 | Trex2-1 | CTTTTGCGGCCTTGAGCCC | tatcttgtggaaaggacgaaacaccGCTTTTGCGGCCTTGAGCCCgtttaagagctatgctggaaacagc |
| Trex2 | Trex2-2 | TGCACGGAGCTACAACGTC | tatcttgtggaaaggacgaaacaccGTGCACGGAGCTACAACGTCgtttaagagctatgctggaaacagc |
| Trex2 | Trex2-5 | TCCCTGGAGAACCCAGAAC | tatcttgtggaaaggacgaaacaccGTCCCTGGAGAACCCAGAACgtttaagagctatgctggaaacagc |
| Trrap | Trrap-3 | AACGCCCAGATCGGACTGA | tatcttgtggaaaggacgaaacaccGAACGCCCAGATCGGACTGAgtttaagagctatgctggaaacagc |
| Trrap | Trrap-4 | GATCCATCGTGAAGAGTCG | tatcttgtggaaaggacgaaacaccGGATCCATCGTGAAGAGTCGgtttaagagctatgctggaaacagc |
| Trrap | Trrap-5 | TCAATGCTGCGATCCGTAA | tatcttgtggaaaggacgaaacaccGTCAATGCTGCGATCCGTAAgtttaagagctatgctggaaacagc |
| Vps72 | Vps72-3 | GTAGGTGATTATTGGCCCA | tatcttgtggaaaggacgaaacaccGGTAGGTGATTATTGGCCCAgtttaagagctatgctggaaacagc |

| | | | |
|---|---|---|---|
| Vps72 | Vps72-4 | AAGCGGCACCGTCACTGAG | tatcttgtggaaaggacgaaacaccGAAGCGGCACCGTCACTGAGgtttaagagctatgctggaaacagc |
| Vps72 | Vps72-5 | GGCCGGGCTCCCCAACAAG | tatcttgtggaaaggacgaaacaccGGGCCGGGCTCCCCAACAAGgtttaagagctatgctggaaacagc |
| Wapl | Wapl-2 | ATAAAGCGAGATCCTCACA | tatcttgtggaaaggacgaaacaccGATAAAGCGAGATCCTCACAgtttaagagctatgctggaaacagc |
| Wapl | Wapl-4 | GCTGGCCGGCTCAGAAAAA | tatcttgtggaaaggacgaaacaccGGCTGGCCGGCTCAGAAAAAgtttaagagctatgctggaaacagc |
| Wapl | Wapl-5 | AGTCTAGTACTAATAATGC | tatcttgtggaaaggacgaaacaccGAGTCTAGTACTAATAATGCgtttaagagctatgctggaaacagc |
| Wrn | Wrn-2 | GTTGGGTTCCGGCAATAAA | tatcttgtggaaaggacgaaacaccGGTTGGGTTCCGGCAATAAAgtttaagagctatgctggaaacagc |
| Wrn | Wrn-3 | CGTGATTTTGACGTCAAGT | tatcttgtggaaaggacgaaacaccGCGTGATTTTGACGTCAAGTgtttaagagctatgctggaaacagc |
| Wrn | Wrn-4 | GGCCGCCCATATACAAGCC | tatcttgtggaaaggacgaaacaccGGGCCGCCCATATACAAGCCgtttaagagctatgctggaaacagc |
| Xpa | Xpa-2 | GTGTAATCAAACTCCATGA | tatcttgtggaaaggacgaaacaccGGTGTAATCAAACTCCATGAgtttaagagctatgctggaaacagc |
| Xpa | Xpa-3 | GAAACATTGTTCATGAACC | tatcttgtggaaaggacgaaacaccGGAAACATTGTTCATGAACCgtttaagagctatgctggaaacagc |
| Xpa | Xpa-4 | CCAAAATGATTGACACCAA | tatcttgtggaaaggacgaaacaccGCCAAAATGATTGACACCAAgtttaagagctatgctggaaacagc |
| Xrcc1 | Xrcc1-2 | ACACGAAACTGGCCGAGCT | tatcttgtggaaaggacgaaacaccGACACGAAACTGGCCGAGCTgtttaagagctatgctggaaacagc |
| Xrcc1 | Xrcc1-4 | TACGACTAGGGACAGACAC | tatcttgtggaaaggacgaaacaccGTACGACTAGGGACAGACACgtttaagagctatgctggaaacagc |
| Xrcc1 | Xrcc1-5 | GTCCAGTGCGAGCCCCCCT | tatcttgtggaaaggacgaaacaccGGTCCAGTGCGAGCCCCCCTgtttaagagctatgctggaaacagc |
| Zbtb38 | Zbtb38-1 | TGCAACAAGGTCTTTGCGC | tatcttgtggaaaggacgaaacaccGTGCAACAAGGTCTTTGCGCgtttaagagctatgctggaaacagc |
| Zbtb38 | Zbtb38-4 | TGGCGAACTATCCAGCGCG | tatcttgtggaaaggacgaaacaccGTGGCGAACTATCCAGCGCGgtttaagagctatgctggaaacagc |
| Zbtb38 | Zbtb38-5 | ACCGTTGTGGTCAAAAGAC | tatcttgtggaaaggacgaaacaccGACCGTTGTGGTCAAAAGACgtttaagagctatgctggaaacagc |
| Zfp384 | Zfp384-1 | GAGAAGGGAGTCAATCAAG | tatcttgtggaaaggacgaaacaccGGAGAAGGGAGTCAATCAAGgtttaagagctatgctggaaacagc |
| Zfp384 | Zfp384-3 | GTAGCGTCAACCCTAACTG | tatcttgtggaaaggacgaaacaccGGTAGCGTCAACCCTAACTGgtttaagagctatgctggaaacagc |
| Zfp384 | Zfp384-4 | ACCCTTACGTCCTTGCCCC | tatcttgtggaaaggacgaaacaccGACCCTTACGTCCTTGCCCCgtttaagagctatgctggaaacagc |
| Fance | Fance-A1 | GAACGGCTGCGGCTTCGGG | tatcttgtggaaaggacgaaacaccGGAACGGCTGCGGCTTCGGGgtttaagagctatgctggaaacagc |
| Fance | Fance-A2 | GACAGCGCGGGAACGGCTG | tatcttgtggaaaggacgaaacaccGGACAGCGCGGGAACGGCTGgtttaagagctatgctggaaacagc |
| Fance | Fance-A3 | TTGGGAGACCGGAGAAACC | tatcttgtggaaaggacgaaacaccGTTGGGAGACCGGAGAAACCgtttaagagctatgctggaaacagc |
| Lig1 | Lig1-A1 | AATGCCGTGGCGCGAACTT | tatcttgtggaaaggacgaaacaccGAATGCCGTGGCGCGAACTTgtttaagagctatgctggaaacagc |
| Lig1 | Lig1-A2 | CGCGAACTTGGGACTGCAG | tatcttgtggaaaggacgaaacaccGCGCGAACTTGGGACTGCAGgtttaagagctatgctggaaacagc |
| Lig1 | Lig1-A3 | CGTCTGCGCGCGAATGCCG | tatcttgtggaaaggacgaaacaccGCGTCTGCGCGCGAATGCCGgtttaagagctatgctggaaacagc |
| Nhej1 | Nhej1-A1 | CCCGCTCGCGCAAACCGAA | tatcttgtggaaaggacgaaacaccGCCCGCTCGCGCAAACCGAAgtttaagagctatgctggaaacagc |
| Nhej1 | Nhej1-A2 | CCGAAAGGCCTAGAGTAAG | tatcttgtggaaaggacgaaacaccGCCGAAAGGCCTAGAGTAAGgtttaagagctatgctggaaacagc |
| Nhej1 | Nhej1-A3 | TGCGTGCGGCTAAGAGAGT | tatcttgtggaaaggacgaaacaccGTGCGTGCGGCTAAGAGAGTgtttaagagctatgctggaaacagc |
| NonTargeting | NonTargeting-0565 | CTCGCTGATGTGTAATCCG | tatcttgtggaaaggacgaaacaccGCTCGCTGATGTGTAATCCGgtttaagagctatgctggaaacagc |
| NonTargeting | NonTargeting-0760 | ATCACTCCCGAAAGTAGTG | tatcttgtggaaaggacgaaacaccGATCACTCCCGAAAGTAGTGgtttaagagctatgctggaaacagc |
| NonTargeting | NonTargeting-0203 | AGATTAAATTAACGCCGGC | tatcttgtggaaaggacgaaacaccGAGATTAAATTAACGCCGGCgtttaagagctatgctggaaacagc |
| NonTargeting | NonTargeting-0750 | TGCCAGCGTCGATTGACGC | tatcttgtggaaaggacgaaacaccGTGCCAGCGTCGATTGACGCgtttaagagctatgctggaaacagc |
| NonTargeting | NonTargeting-0833 | AACCTTCGCGCAACTTAAC | tatcttgtggaaaggacgaaacaccGAACCTTCGCGCAACTTAACgtttaagagctatgctggaaacagc |
| NonTargeting | NonTargeting-0436 | CTTATCGTCATGCGGGTGA | tatcttgtggaaaggacgaaacaccGCTTATCGTCATGCGGGTGAgtttaagagctatgctggaaacagc |
| NonTargeting | NonTargeting-0510 | TAGTGCGTGTGATGTCGGG | tatcttgtggaaaggacgaaacaccGTAGTGCGTGTGATGTCGGGgtttaagagctatgctggaaacagc |
| NonTargeting | NonTargeting-0220 | ATAAGACTCCGCGAGCTTC | tatcttgtggaaaggacgaaacaccGATAAGACTCCGCGAGCTTCgtttaagagctatgctggaaacagc |
| NonTargeting | NonTargeting-0558 | GTGTGGACCGCTTTTACGC | tatcttgtggaaaggacgaaacaccGGTGTGGACCGCTTTTACGCgtttaagagctatgctggaaacagc |
| NonTargeting | NonTargeting-0150 | ATTCATTATCGGCATACGG | tatcttgtggaaaggacgaaacaccGATTCATTATCGGCATACGGgtttaagagctatgctggaaacagc |
| NonTargeting | NonTargeting-0986 | CGACAAGCGTTGAGTTCGC | tatcttgtggaaaggacgaaacaccGCGACAAGCGTTGAGTTCGCgtttaagagctatgctggaaacagc |
| NonTargeting | NonTargeting-0902 | CCGCAACGTTAGATGTATA | tatcttgtggaaaggacgaaacaccGCCGCAACGTTAGATGTATAgtttaagagctatgctggaaacagc |
| NonTargeting | NonTargeting-0205 | GGACGCTCATCGAGTGACG | tatcttgtggaaaggacgaaacaccGGGACGCTCATCGAGTGACGgtttaagagctatgctggaaacagc |
| NonTargeting | NonTargeting-0642 | TCGCTCGAACTCACGTAGC | tatcttgtggaaaggacgaaacaccGTCGCTCGAACTCACGTAGCgtttaagagctatgctggaaacagc |
| NonTargeting | NonTargeting-0385 | GATATTCGCGCGGTCTTCA | tatcttgtggaaaggacgaaacaccGGATATTCGCGCGGTCTTCAgtttaagagctatgctggaaacagc |
| NonTargeting | NonTargeting-0356 | CGTACCTATCGATAAACCA | tatcttgtggaaaggacgaaacaccGCGTACCTATCGATAAACCAgtttaagagctatgctggaaacagc |
| NonTargeting | NonTargeting-0181 | GCTTATAGACGAGACTCGA | tatcttgtggaaaggacgaaacaccGGCTTATAGACGAGACTCGAgtttaagagctatgctggaaacagc |
| NonTargeting | NonTargeting-0077 | CTACAGATTTGCGTTCGAG | tatcttgtggaaaggacgaaacaccGCTACAGATTTGCGTTCGAGgtttaagagctatgctggaaacagc |
| NonTargeting | NonTargeting-0891 | GCGGGTTAGGCCCCGTTTT | tatcttgtggaaaggacgaaacaccGGCGGGTTAGGCCCCGTTTTgtttaagagctatgctggaaacagc |
| NonTargeting | NonTargeting-0176 | CGGCCCGTGACCGTTCAAT | tatcttgtggaaaggacgaaacaccGCGGCCCGTGACCGTTCAATgtttaagagctatgctggaaacagc |
| NonTargeting | NonTargeting-0872 | ACCGTCGGGATCCTTAATT | tatcttgtggaaaggacgaaacaccGACCGTCGGGATCCTTAATTgtttaagagctatgctggaaacagc |
| NonTargeting | NonTargeting-0801 | CAGGGTTATTCCGCTTCTA | tatcttgtggaaaggacgaaacaccGCAGGGTTATTCCGCTTCTAgtttaagagctatgctggaaacagc |
| NonTargeting | NonTargeting-0432 | ACAGTTGACGCGACGGAGA | tatcttgtggaaaggacgaaacaccGACAGTTGACGCGACGGAGAgtttaagagctatgctggaaacagc |
| NonTargeting | NonTargeting-0684 | AGCGCCAAGCGTGCGTTGA | tatcttgtggaaaggacgaaacaccGAGCGCCAAGCGTGCGTTGAgtttaagagctatgctggaaacagc |
| NonTargeting | NonTargeting-0836 | CCCGAGCGGATTCACCGCG | tatcttgtggaaaggacgaaacaccGCCCGAGCGGATTCACCGCGgtttaagagctatgctggaaacagc |
| NonTargeting | NonTargeting-0799 | TCGTTAACGACCCGGATTC | tatcttgtggaaaggacgaaacaccGTCGTTAACGACCCGGATTCgtttaagagctatgctggaaacagc |
| NonTargeting | NonTargeting-0308 | ATGGCCAGTAACGGCGTCA | tatcttgtggaaaggacgaaacaccGATGGCCAGTAACGGCGTCAgtttaagagctatgctggaaacagc |
| NonTargeting | NonTargeting-0911 | GTAACATAGGCGGCGATCT | tatcttgtggaaaggacgaaacaccGGTAACATAGGCGGCGATCTgtttaagagctatgctggaaacagc |
| NonTargeting | NonTargeting-0282 | CCCGATAGAATTACCCATT | tatcttgtggaaaggacgaaacaccGCCCGATAGAATTACCCATTgtttaagagctatgctggaaacagc |
| NonTargeting | NonTargeting-0709 | GTAAAAAGCGTAACGAGTA | tatcttgtggaaaggacgaaacaccGGTAAAAAGCGTAACGAGTAgtttaagagctatgctggaaacagc |
| NonTargeting | NonTargeting-0106 | CGACGTGTATCGATCACTC | tatcttgtggaaaggacgaaacaccGCGACGTGTATCGATCACTCgtttaagagctatgctggaaacagc |
| NonTargeting | NonTargeting-0281 | AACGTAACGGCATGCATCA | tatcttgtggaaaggacgaaacaccGAACGTAACGGCATGCATCAgtttaagagctatgctggaaacagc |
| NonTargeting | NonTargeting-0763 | CCACGACCTGCGAATAATT | tatcttgtggaaaggacgaaacaccGCCACGACCTGCGAATAATTgtttaagagctatgctggaaacagc |
| NonTargeting | NonTargeting-0964 | GTCTAGATAATCTTACCGA | tatcttgtggaaaggacgaaacaccGGTCTAGATAATCTTACCGAgtttaagagctatgctggaaacagc |
| NonTargeting | NonTargeting-0035 | ACGTCTAATTTCTGGCCGT | tatcttgtggaaaggacgaaacaccGACGTCTAATTTCTGGCCGTgtttaagagctatgctggaaacagc |
| NonTargeting | NonTargeting-0010 | CTTCTACTCGCAACGTATT | tatcttgtggaaaggacgaaacaccGCTTCTACTCGCAACGTATTgtttaagagctatgctggaaacagc |
| NonTargeting | NonTargeting-0333 | TTGCGTGTGTCCGTACAAA | tatcttgtggaaaggacgaaacaccGTTGCGTGTGTCCGTACAAAgtttaagagctatgctggaaacagc |
| NonTargeting | NonTargeting-0172 | GTCTGTCCGTTGCGACCAC | tatcttgtggaaaggacgaaacaccGGTCTGTCCGTTGCGACCACgtttaagagctatgctggaaacagc |
| NonTargeting | NonTargeting-0925 | CTCCAGGTGCGAGATAGGC | tatcttgtggaaaggacgaaacaccGCTCCAGGTGCGAGATAGGCgtttaagagctatgctggaaacagc |
| NonTargeting | NonTargeting-0415 | TTCTTCAAAGACGGGCGGC | tatcttgtggaaaggacgaaacaccGTTCTTCAAAGACGGGCGGCgtttaagagctatgctggaaacagc |
| NonTargeting | NonTargeting-0236 | CCGCTCTTGATAACGACGC | tatcttgtggaaaggacgaaacaccGCCGCTCTTGATAACGACGCgtttaagagctatgctggaaacagc |
| NonTargeting | NonTargeting-0914 | CGATTATCAGTAGTCCACG | tatcttgtggaaaggacgaaacaccGCGATTATCAGTAGTCCACGgtttaagagctatgctggaaacagc |
| NonTargeting | NonTargeting-0464 | CGCGAGTGCCAAACGAGTG | tatcttgtggaaaggacgaaacaccGCGCGAGTGCCAAACGAGTGgtttaagagctatgctggaaacagc |
| NonTargeting | NonTargeting-0383 | TACCATGATAACCGTACTA | tatcttgtggaaaggacgaaacaccGTACCATGATAACCGTACTAgtttaagagctatgctggaaacagc |
| NonTargeting | NonTargeting-0367 | CTTCATTATTAACCGGCGT | tatcttgtggaaaggacgaaacaccGCTTCATTATTAACCGGCGTgtttaagagctatgctggaaacagc |
| NonTargeting | NonTargeting-0028 | GGGATGGCCTTACGTCGCG | tatcttgtggaaaggacgaaacaccGGGGATGGCCTTACGTCGCGgtttaagagctatgctggaaacagc |
| NonTargeting | NonTargeting-0975 | TGCGCGCGTATAGAAAAAA | tatcttgtggaaaggacgaaacaccGTGCGCGCGTATAGAAAAAAgtttaagagctatgctggaaacagc |

| | | | |
|---|---|---|---|
| NonTargeting | NonTargeting-0765 | ACTAGCACGAACTGCGGTT | tatcttgtggaaaggacgaaacaccGACTAGCACGAACTGCGGTTgtttaagagctatgctggaaacagc |
| NonTargeting | NonTargeting-0895 | TCACAAAAAACGAGCGTTA | tatcttgtggaaaggacgaaacaccGTCACAAAAAACGAGCGTTAgtttaagagctatgctggaaacagc |
| NonTargeting | NonTargeting-0977 | GCCCGAAGGGTCACAAGCG | tatcttgtggaaaggacgaaacaccGGCCCGAAGGGTCACAAGCGgtttaagagctatgctggaaacagc |
| NonTargeting | NonTargeting-0118 | CAAATAGGTCGGAGCGTGT | tatcttgtggaaaggacgaaacaccGCAAATAGGTCGGAGCGTGTgtttaagagctatgctggaaacagc |
| NonTargeting | NonTargeting-0297 | CTAGTTCTCCCGGGCGAAA | tatcttgtggaaaggacgaaacaccGCTAGTTCTCCCGGGCGAAAgtttaagagctatgctggaaacagc |
| NonTargeting | NonTargeting-0737 | CTTGAGGCAACGCGACTGA | tatcttgtggaaaggacgaaacaccGCTTGAGGCAACGCGACTGAgtttaagagctatgctggaaacagc |
| NonTargeting | NonTargeting-0579 | TACCACTTATCGACCTTGC | tatcttgtggaaaggacgaaacaccGTACCACTTATCGACCTTGCgtttaagagctatgctggaaacagc |
| NonTargeting | NonTargeting-0331 | CTAAACGTATTTTACGGGC | tatcttgtggaaaggacgaaacaccGCTAAACGTATTTTACGGGCgtttaagagctatgctggaaacagc |
| NonTargeting | NonTargeting-0859 | GGCAACGTCATCTGGCGAC | tatcttgtggaaaggacgaaacaccGGGCAACGTCATCTGGCGACgtttaagagctatgctggaaacagc |
| NonTargeting | NonTargeting-0211 | TACATGCGGCAAGTCGACT | tatcttgtggaaaggacgaaacaccGTACATGCGGCAAGTCGACTgtttaagagctatgctggaaacagc |
| NonTargeting | NonTargeting-0858 | GGGTTGGATACGCGATTTA | tatcttgtggaaaggacgaaacaccGGGGTTGGATACGCGATTTAgtttaagagctatgctggaaacagc |
| NonTargeting | NonTargeting-0566 | GATACATCGGCGCGCTAGT | tatcttgtggaaaggacgaaacaccGGATACATCGGCGCGCTAGTgtttaagagctatgctggaaacagc |
| NonTargeting | NonTargeting-0323 | TTCCGGATATATACGGTTA | tatcttgtggaaaggacgaaacaccGTTCCGGATATATACGGTTAgtttaagagctatgctggaaacagc |
| NonTargeting | NonTargeting-0628 | TTACTTACCTGACCGGCAA | tatcttgtggaaaggacgaaacaccGTTACTTACCTGACCGGCAAgtttaagagctatgctggaaacagc |
| NonTargeting | NonTargeting-0981 | AGGCGGGCTCTTCTATCGT | tatcttgtggaaaggacgaaacaccGAGGCGGGCTCTTCTATCGTgtttaagagctatgctggaaacagc |
| NonTargeting | NonTargeting-0243 | GAGTCTCACGCAATTAGCG | tatcttgtggaaaggacgaaacaccGGAGTCTCACGCAATTAGCGgtttaagagctatgctggaaacagc |
| NonTargeting | NonTargeting-0554 | TTAATATTGTGGCCCGCAC | tatcttgtggaaaggacgaaacaccGTTAATATTGTGGCCCGCACgtttaagagctatgctggaaacagc |
| NonTargeting | NonTargeting-0560 | ATACGTGAGGTTGCCGGTG | tatcttgtggaaaggacgaaacaccGATACGTGAGGTTGCCGGTGgtttaagagctatgctggaaacagc |
| NonTargeting | NonTargeting-0753 | TGTGGTATATCCCCGTTTT | tatcttgtggaaaggacgaaacaccGTGTGGTATATCCCCGTTTTgtttaagagctatgctggaaacagc |
| NonTargeting | NonTargeting-0060 | ACCGGTGTGTTTACGCGTG | tatcttgtggaaaggacgaaacaccGACCGGTGTGTTTACGCGTGgtttaagagctatgctggaaacagc |
| NonTargeting | NonTargeting-0193 | CGTGCCATGGGAGGCCGTT | tatcttgtggaaaggacgaaacaccGCGTGCCATGGGAGGCCGTTgtttaagagctatgctggaaacagc |
| NonTargeting | NonTargeting-0488 | ATCCCCTACGCTTCAACCT | tatcttgtggaaaggacgaaacaccGATCCCCTACGCTTCAACCTgtttaagagctatgctggaaacagc |
| NonTargeting | NonTargeting-0413 | CATTCTTGAGCTCCGCGCC | tatcttgtggaaaggacgaaacaccGCATTCTTGAGCTCCGCGCCgtttaagagctatgctggaaacagc |
| NonTargeting | NonTargeting-0610 | CACGAACCCGTTCGTATGG | tatcttgtggaaaggacgaaacaccGCACGAACCCGTTCGTATGGgtttaagagctatgctggaaacagc |
| NonTargeting | NonTargeting-0202 | CTTGTAATCTAAAGACGCG | tatcttgtggaaaggacgaaacaccGCTTGTAATCTAAAGACGCGgtttaagagctatgctggaaacagc |
| NonTargeting | NonTargeting-0932 | TCGTGGGGCACGATGCGTA | tatcttgtggaaaggacgaaacaccGTCGTGGGGCACGATGCGTAgtttaagagctatgctggaaacagc |
| NonTargeting | NonTargeting-0078 | GACTGAAACCGATAGTATC | tatcttgtggaaaggacgaaacaccGGACTGAAACCGATAGTATCgtttaagagctatgctggaaacagc |
| NonTargeting | NonTargeting-0951 | GGTAGCACCCCGCCGACAC | tatcttgtggaaaggacgaaacaccGGGTAGCACCCCGCCGACACgtttaagagctatgctggaaacagc |
| NonTargeting | NonTargeting-0147 | GACCAATGTTACCGTAGGT | tatcttgtggaaaggacgaaacaccGGACCAATGTTACCGTAGGTgtttaagagctatgctggaaacagc |
| NonTargeting | NonTargeting-0946 | AGATTCGATACTGCGGTCA | tatcttgtggaaaggacgaaacaccGAGATTCGATACTGCGGTCAgtttaagagctatgctggaaacagc |
| NonTargeting | NonTargeting-0823 | CATTAAGACGCCCACATAT | tatcttgtggaaaggacgaaacaccGCATTAAGACGCCCACATATgtttaagagctatgctggaaacagc |
| NonTargeting | NonTargeting-0302 | ATTACGGGGTTCCGTAACT | tatcttgtggaaaggacgaaacaccGATTACGGGGTTCCGTAACTgtttaagagctatgctggaaacagc |
| NonTargeting | NonTargeting-0688 | AGCGCGGGCCAGCGAACGT | tatcttgtggaaaggacgaaacaccGAGCGCGGGCCAGCGAACGTgtttaagagctatgctggaaacagc |
| NonTargeting | NonTargeting-0024 | ACCAACCTTACGGTAACTC | tatcttgtggaaaggacgaaacaccGACCAACCTTACGGTAACTCgtttaagagctatgctggaaacagc |
| NonTargeting | NonTargeting-0363 | CGTGACGCGATCAAACGGT | tatcttgtggaaaggacgaaacaccGCGTGACGCGATCAAACGGTgtttaagagctatgctggaaacagc |
| NonTargeting | NonTargeting-0706 | TACTGACGTTACGTTCATA | tatcttgtggaaaggacgaaacaccGTACTGACGTTACGTTCATAgtttaagagctatgctggaaacagc |
| NonTargeting | NonTargeting-0115 | TGTGCCTCGACAGCGTAAG | tatcttgtggaaaggacgaaacaccGTGTGCCTCGACAGCGTAAGgtttaagagctatgctggaaacagc |
| NonTargeting | NonTargeting-0133 | TATCCTCGCAATCGTTAGG | tatcttgtggaaaggacgaaacaccGTATCCTCGCAATCGTTAGGgtttaagagctatgctggaaacagc |
| NonTargeting | NonTargeting-0518 | GACACTCGCCGACCCCACT | tatcttgtggaaaggacgaaacaccGGACACTCGCCGACCCCACTgtttaagagctatgctggaaacagc |
| NonTargeting | NonTargeting-0145 | GACGTAGATTAGGGCGTAA | tatcttgtggaaaggacgaaacaccGGACGTAGATTAGGGCGTAAgtttaagagctatgctggaaacagc |
| NonTargeting | NonTargeting-0740 | CTTCGACCACGGGTACTGC | tatcttgtggaaaggacgaaacaccGCTTCGACCACGGGTACTGCgtttaagagctatgctggaaacagc |
| NonTargeting | NonTargeting-0313 | GTTAAAAATTAAGCGGTCC | tatcttgtggaaaggacgaaacaccGGTTAAAAATTAAGCGGTCCgtttaagagctatgctggaaacagc |
| NonTargeting | NonTargeting-0800 | AGCTAATCTGTAGACGTCG | tatcttgtggaaaggacgaaacaccGAGCTAATCTGTAGACGTCGgtttaagagctatgctggaaacagc |
| NonTargeting | NonTargeting-0797 | GAGTTCAAGCGTAACTTCG | tatcttgtggaaaggacgaaacaccGGAGTTCAAGCGTAACTTCGgtttaagagctatgctggaaacagc |
| NonTargeting | NonTargeting-0587 | GCAACGCACGCTGGGTTGT | tatcttgtggaaaggacgaaacaccGGCAACGCACGCTGGGTTGTgtttaagagctatgctggaaacagc |
| NonTargeting | NonTargeting-0906 | ACAGTTGGATAACCGTGTC | tatcttgtggaaaggacgaaacaccGACAGTTGGATAACCGTGTCgtttaagagctatgctggaaacagc |
| NonTargeting | NonTargeting-0196 | CTGATTAGACCCGGCGTAA | tatcttgtggaaaggacgaaacaccGCTGATTAGACCCGGCGTAAgtttaagagctatgctggaaacagc |
| NonTargeting | NonTargeting-0738 | AACCAGTATCTAACCCGAC | tatcttgtggaaaggacgaaacaccGAACCAGTATCTAACCCGACgtttaagagctatgctggaaacagc |
| NonTargeting | NonTargeting-0136 | TTTCCCGGGACTGTCGCGT | tatcttgtggaaaggacgaaacaccGTTTCCCGGGACTGTCGCGTgtttaagagctatgctggaaacagc |
| NonTargeting | NonTargeting-0497 | TCCTAGATCCTATCGGGAG | tatcttgtggaaaggacgaaacaccGTCCTAGATCCTATCGGGAGgtttaagagctatgctggaaacagc |
| NonTargeting | NonTargeting-0110 | CAGAGCGTAATCGGCATCG | tatcttgtggaaaggacgaaacaccGCAGAGCGTAATCGGCATCGgtttaagagctatgctggaaacagc |
| NonTargeting | NonTargeting-0901 | CATGCGCTCGCGTGGACTG | tatcttgtggaaaggacgaaacaccGCATGCGCTCGCGTGGACTGgtttaagagctatgctggaaacagc |
| NonTargeting | NonTargeting-0046 | CGCGAGGGCACCGACAAGT | tatcttgtggaaaggacgaaacaccGCGCGAGGGCACCGACAAGTgtttaagagctatgctggaaacagc |
| NonTargeting | NonTargeting-0574 | TTCGCGGGGGCTTCTATCA | tatcttgtggaaaggacgaaacaccGTTCGCGGGGGCTTCTATCAgtttaagagctatgctggaaacagc |
| NonTargeting | NonTargeting-0457 | CCTTCCTCGCAGACCCGAC | tatcttgtggaaaggacgaaacaccGCCTTCCTCGCAGACCCGACgtttaagagctatgctggaaacagc |
| NonTargeting | NonTargeting-0556 | CGAGCGCTATCCCGGTGGA | tatcttgtggaaaggacgaaacaccGCGAGCGCTATCCCGGTGGAgtttaagagctatgctggaaacagc |
| NonTargeting | NonTargeting-0916 | GGCTAGTGCGTTGTTACGA | tatcttgtggaaaggacgaaacaccGGGCTAGTGCGTTGTTACGAgtttaagagctatgctggaaacagc |
| NonTargeting | NonTargeting-0559 | TTATACCACCTACTATGAC | tatcttgtggaaaggacgaaacaccGTTATACCACCTACTATGACgtttaagagctatgctggaaacagc |
| NonTargeting | NonTargeting-0325 | GGCGGGACGTAATATTATG | tatcttgtggaaaggacgaaacaccGGGCGGGACGTAATATTATGgtttaagagctatgctggaaacagc |
| NonTargeting | NonTargeting-0793 | GTCTCGTAATGTCCTATAT | tatcttgtggaaaggacgaaacaccGGTCTCGTAATGTCCTATATgtttaagagctatgctggaaacagc |
| NonTargeting | NonTargeting-0074 | CGGATCTGACGGTTACTTA | tatcttgtggaaaggacgaaacaccGCGGATCTGACGGTTACTTAgtttaagagctatgctggaaacagc |
| NonTargeting | NonTargeting-0818 | ATGCGGACCCACGTTAAGC | tatcttgtggaaaggacgaaacaccGATGCGGACCCACGTTAAGCgtttaagagctatgctggaaacagc |
| NonTargeting | NonTargeting-0758 | CATTCCCGCTTACGTAATT | tatcttgtggaaaggacgaaacaccGCATTCCCGCTTACGTAATTgtttaagagctatgctggaaacagc |
| NonTargeting | NonTargeting-0624 | AGACCCATTATGATCCTAG | tatcttgtggaaaggacgaaacaccGAGACCCATTATGATCCTAGgtttaagagctatgctggaaacagc |
| NonTargeting | NonTargeting-0349 | TGCCTGATAGTGTGAAGCG | tatcttgtggaaaggacgaaacaccGTGCCTGATAGTGTGAAGCGgtttaagagctatgctggaaacagc |
| NonTargeting | NonTargeting-0927 | GGTCGCACCCCAATAGCTG | tatcttgtggaaaggacgaaacaccGGGTCGCACCCCAATAGCTGgtttaagagctatgctggaaacagc |
| NonTargeting | NonTargeting-0899 | CGTGACGTCCCACCCTGAG | tatcttgtggaaaggacgaaacaccGCGTGACGTCCCACCCTGAGgtttaagagctatgctggaaacagc |
| NonTargeting | NonTargeting-0761 | CAACCCACGGACGGCACCG | tatcttgtggaaaggacgaaacaccGCAACCCACGGACGGCACCGgtttaagagctatgctggaaacagc |
| NonTargeting | NonTargeting-0652 | CGAGTTATTTCGCTATAGG | tatcttgtggaaaggacgaaacaccGCGAGTTATTTCGCTATAGGgtttaagagctatgctggaaacagc |
| NonTargeting | NonTargeting-0269 | ATTGAGAAGCCGCGGTATC | tatcttgtggaaaggacgaaacaccGATTGAGAAGCCGCGGTATCgtttaagagctatgctggaaacagc |
| NonTargeting | NonTargeting-0994 | TAGCTTCATGTCCGGTCGG | tatcttgtggaaaggacgaaacaccGTAGCTTCATGTCCGGTCGGgtttaagagctatgctggaaacagc |
| NonTargeting | NonTargeting-0031 | TGGTTACGTTAACGACTAC | tatcttgtggaaaggacgaaacaccGTGGTTACGTTAACGACTACgtttaagagctatgctggaaacagc |
| NonTargeting | NonTargeting-0687 | CCCGTTGTGAGCGGCATGC | tatcttgtggaaaggacgaaacaccGCCCGTTGTGAGCGGCATGCgtttaagagctatgctggaaacagc |
| NonTargeting | NonTargeting-0762 | TGTTGTCCGAGAACCCGAC | tatcttgtggaaaggacgaaacaccGTGTTGTCCGAGAACCCGACgtttaagagctatgctggaaacagc |
| NonTargeting | NonTargeting-0999 | AACTCGTTAGGCCGTGAAG | tatcttgtggaaaggacgaaacaccGAACTCGTTAGGCCGTGAAGgtttaagagctatgctggaaacagc |
| NonTargeting | NonTargeting-0475 | CTTACCTACTCCGCCCCGC | tatcttgtggaaaggacgaaacaccGCTTACCTACTCCGCCCCGCgtttaagagctatgctggaaacagc |

| | | | |
|---|---|---|---|
| NonTargeting | NonTargeting-0184 | ATGCGGGTGGAAAACGTTA | tatcttgtggaaaggacgaaacaccGATGCGGGTGGAAAACGTTAgtttaagagctatgctggaaacagc |
| NonTargeting | NonTargeting-0051 | GGGTAGGCCTAATTACGGA | tatcttgtggaaaggacgaaacaccGGGGTAGGCCTAATTACGGAgtttaagagctatgctggaaacagc |
| NonTargeting | NonTargeting-0014 | TCAAGCCGAACGCTGCCGG | tatcttgtggaaaggacgaaacaccGTCAAGCCGAACGCTGCCGGgtttaagagctatgctggaaacagc |
| NonTargeting | NonTargeting-0130 | CCATAGCCAATCGCTAGTT | tatcttgtggaaaggacgaaacaccGCCATAGCCAATCGCTAGTTgtttaagagctatgctggaaacagc |
| NonTargeting | NonTargeting-0742 | ACTTCGACGAGAAAACGTT | tatcttgtggaaaggacgaaacaccGACTTCGACGAGAAAACGTTgtttaagagctatgctggaaacagc |
| NonTargeting | NonTargeting-0634 | TACGCAGGGTGTAGCGACC | tatcttgtggaaaggacgaaacaccGTACGCAGGGTGTAGCGACCgtttaagagctatgctggaaacagc |
| NonTargeting | NonTargeting-0991 | TTAAGTAGATCGGTGACAT | tatcttgtggaaaggacgaaacaccGTTAAGTAGATCGGTGACATgtttaagagctatgctggaaacagc |
| NonTargeting | NonTargeting-0808 | ACCGGCCAACGGTAGCGGC | tatcttgtggaaaggacgaaacaccGACCGGCCAACGGTAGCGGCgtttaagagctatgctggaaacagc |
| NonTargeting | NonTargeting-0538 | TCTAACATCGGCGCACGTG | tatcttgtggaaaggacgaaacaccGTCTAACATCGGCGCACGTGgtttaagagctatgctggaaacagc |
| NonTargeting | NonTargeting-0602 | CGATCGCCGGTATAGCTTT | tatcttgtggaaaggacgaaacaccGCGATCGCCGGTATAGCTTTgtttaagagctatgctggaaacagc |
| NonTargeting | NonTargeting-0861 | ACGGTCCCCCCTATGAAAT | tatcttgtggaaaggacgaaacaccGACGGTCCCCCCTATGAAATgtttaagagctatgctggaaacagc |
| NonTargeting | NonTargeting-0929 | CCAGCCTCGGTATAACTCG | tatcttgtggaaaggacgaaacaccGCCAGCCTCGGTATAACTCGgtttaagagctatgctggaaacagc |
| NonTargeting | NonTargeting-0079 | CAATTCTGCAACGCACGTC | tatcttgtggaaaggacgaaacaccGCAATTCTGCAACGCACGTCgtttaagagctatgctggaaacagc |
| NonTargeting | NonTargeting-0001 | CGAGGTATTCGGCTCCGCG | tatcttgtggaaaggacgaaacaccGCGAGGTATTCGGCTCCGCGgtttaagagctatgctggaaacagc |
| NonTargeting | NonTargeting-0111 | CCGCGCCATAATATGCCAT | tatcttgtggaaaggacgaaacaccGCCGCGCCATAATATGCCATgtttaagagctatgctggaaacagc |
| NonTargeting | NonTargeting-0185 | CAAAAAGCGGACACGCGAC | tatcttgtggaaaggacgaaacaccGCAAAAAGCGGACACGCGACgtttaagagctatgctggaaacagc |
| NonTargeting | NonTargeting-0194 | GCTTTACGTAAGGAGCGTA | tatcttgtggaaaggacgaaacaccGGCTTTACGTAAGGAGCGTAgtttaagagctatgctggaaacagc |
| NonTargeting | NonTargeting-0159 | CGCGTACATATAAATAGGT | tatcttgtggaaaggacgaaacaccGCGCGTACATATAAATAGGTgtttaagagctatgctggaaacagc |
| NonTargeting | NonTargeting-0248 | AGTCGAGTTAATAACGCTC | tatcttgtggaaaggacgaaacaccGAGTCGAGTTAATAACGCTCgtttaagagctatgctggaaacagc |
| NonTargeting | NonTargeting-0792 | CAGGTAGGCTCGATACTTG | tatcttgtggaaaggacgaaacaccGCAGGTAGGCTCGATACTTGgtttaagagctatgctggaaacagc |
| NonTargeting | NonTargeting-0612 | ACCGACGGTATACCCTACT | tatcttgtggaaaggacgaaacaccGACCGACGGTATACCCTACTgtttaagagctatgctggaaacagc |
| NonTargeting | NonTargeting-0289 | AACCGGCGTGCGTTAGCGG | tatcttgtggaaaggacgaaacaccGAACCGGCGTGCGTTAGCGGgtttaagagctatgctggaaacagc |
| NonTargeting | NonTargeting-0639 | ACAGCCGGTTGACCGGGTC | tatcttgtggaaaggacgaaacaccGACAGCCGGTTGACCGGGTCgtttaagagctatgctggaaacagc |
| NonTargeting | NonTargeting-0681 | TAGGCCGCTGATCGAATAA | tatcttgtggaaaggacgaaacaccGTAGGCCGCTGATCGAATAAgtttaagagctatgctggaaacagc |
| NonTargeting | NonTargeting-0002 | CTTTCACGGAGGTTCGACG | tatcttgtggaaaggacgaaacaccGCTTTCACGGAGGTTCGACGgtttaagagctatgctggaaacagc |
| NonTargeting | NonTargeting-0507 | AACCCGGGAAACACGTCCG | tatcttgtggaaaggacgaaacaccGAACCCGGGAAACACGTCCGgtttaagagctatgctggaaacagc |
| NonTargeting | NonTargeting-0040 | ACCTCGCAATTGAGCGCTC | tatcttgtggaaaggacgaaacaccGACCTCGCAATTGAGCGCTCgtttaagagctatgctggaaacagc |
| NonTargeting | NonTargeting-0655 | ACCACTATGTACTTATCGC | tatcttgtggaaaggacgaaacaccGACCACTATGTACTTATCGCgtttaagagctatgctggaaacagc |
| NonTargeting | NonTargeting-0057 | GGTGCACACGCCGGCCTAT | tatcttgtggaaaggacgaaacaccGGGTGCACACGCCGGCCTATgtttaagagctatgctggaaacagc |
| NonTargeting | NonTargeting-0246 | CCACTCCGCTCGTTCTAGA | tatcttgtggaaaggacgaaacaccGCCACTCCGCTCGTTCTAGAgtttaagagctatgctggaaacagc |
| NonTargeting | NonTargeting-0643 | GGCGCGTCATAGACACACG | tatcttgtggaaaggacgaaacaccGGGCGCGTCATAGACACACGgtttaagagctatgctggaaacagc |
| NonTargeting | NonTargeting-0045 | CGTATCTACCCTACCGCCG | tatcttgtggaaaggacgaaacaccGCGTATCTACCCTACCGCCGgtttaagagctatgctggaaacagc |
| NonTargeting | NonTargeting-0989 | AGGACCTAGGTTCCCGGAT | tatcttgtggaaaggacgaaacaccGAGGACCTAGGTTCCCGGATgtttaagagctatgctggaaacagc |
| NonTargeting | NonTargeting-0508 | GAAACGTTACATTCGACGC | tatcttgtggaaaggacgaaacaccGGAAACGTTACATTCGACGCgtttaagagctatgctggaaacagc |
| NonTargeting | NonTargeting-0638 | CGGGACCCCCGGTATAGTC | tatcttgtggaaaggacgaaacaccGCGGGACCCCCGGTATAGTCgtttaagagctatgctggaaacagc |
| NonTargeting | NonTargeting-0708 | AGCGTCCCGTGATTTAAAG | tatcttgtggaaaggacgaaacaccGAGCGTCCCGTGATTTAAAGgtttaagagctatgctggaaacagc |
| NonTargeting | NonTargeting-0992 | GCGTCGTAACAAGAACTTG | tatcttgtggaaaggacgaaacaccGGCGTCGTAACAAGAACTTGgtttaagagctatgctggaaacagc |
| NonTargeting | NonTargeting-0209 | TTATTTGTCTGTCGAAACG | tatcttgtggaaaggacgaaacaccGTTATTTGTCTGTCGAAACGgtttaagagctatgctggaaacagc |
| NonTargeting | NonTargeting-0575 | GCACCGCGTTTATTGCACT | tatcttgtggaaaggacgaaacaccGGCACCGCGTTTATTGCACTgtttaagagctatgctggaaacagc |
| NonTargeting | NonTargeting-0578 | ACGAAAGGTCCTACGAAGT | tatcttgtggaaaggacgaaacaccGACGAAAGGTCCTACGAAGTgtttaagagctatgctggaaacagc |
| NonTargeting | NonTargeting-0154 | ACCATACGCCTCGTATGCC | tatcttgtggaaaggacgaaacaccGACCATACGCCTCGTATGCCgtttaagagctatgctggaaacagc |
| NonTargeting | NonTargeting-0286 | TGGGCTGACCGTTCTCGAC | tatcttgtggaaaggacgaaacaccGTGGGCTGACCGTTCTCGACgtttaagagctatgctggaaacagc |
| NonTargeting | NonTargeting-0330 | TTACCGTGACGATAAGAAT | tatcttgtggaaaggacgaaacaccGTTACCGTGACGATAAGAATgtttaagagctatgctggaaacagc |
| NonTargeting | NonTargeting-0263 | TTGCGTCCATTCCGTCGCC | tatcttgtggaaaggacgaaacaccGTTGCGTCCATTCCGTCGCCgtttaagagctatgctggaaacagc |
| NonTargeting | NonTargeting-0143 | TAGACATATTGCGTAATCG | tatcttgtggaaaggacgaaacaccGTAGACATATTGCGTAATCGgtttaagagctatgctggaaacagc |
| NonTargeting | NonTargeting-0105 | ATATTTACCGGCGATAAGA | tatcttgtggaaaggacgaaacaccGATATTTACCGGCGATAAGAgtttaagagctatgctggaaacagc |
| NonTargeting | NonTargeting-0298 | TTCGGTTGCAGCTTACACG | tatcttgtggaaaggacgaaacaccGTTCGGTTGCAGCTTACACGgtttaagagctatgctggaaacagc |

### *Supplementary Table S5*

| **Cell-lines** | **Origin** | **Plasmid** | **sgRNA** | **Source** |
|---|---|---|---|---|
| mES IB10 | *M. musculus* | n.a. | n.a. | [18] |
| mES IB10-Cas9 | *M. musculus* | pLenti-Cas9-2A-Blast | n.a. | This study |
| mES IB10-Cas9 Wrn-/- sg-2cC3 | *M. musculus* | pKLV2-U6gRNA5(BbsI)-PGKpuro2ABFP-W | sgmmWrn-2 | This study |
| mES IB10-Cas9 Wrn-/- sg-4cD11 | *M. musculus* | pKLV2-U6gRNA5(BbsI)-PGKpuro2ABFP-W | sgmmWrn-4 | This study |
| mES IB10Hprt-eGFP parental c1 | *M. musculus* | n.a. | n.a. | [39] |
| mES IB10Hprt-eGFP parental c2 | *M. musculus* | n.a. | n.a. | [39] |
| mES IB10Hprt-eGFP Polq-/- c1 | *M. musculus* | n.a. | n.a. | [39] |
| mES IB10Hprt-eGFP Polq-/- c2 | *M. musculus* | n.a. | n.a. | [39] |
| mES IB10Hprt-eGFP Wrn-/- sg-4cH9 | *M. musculus* | pU6-(BbsI)_CBh-Cas9-T2A-mCherry | sgmmWrn-4 | This study |
| mES IB10Hprt-eGFP Wrn-/- sg-4cG11 | *M. musculus* | pU6-(BbsI)_CBh-Cas9-T2A-mCherry | sgmmWrn-4 | This study |
| HEK293T | *H. sapiens* | n.a. | n.a. | [39] |
| HEK293T-Cas9 | *H. sapiens* | pLenti-Cas9-2A-Blast | n.a. | This study |
| U2OS | *H. sapiens* | n.a. | n.a. | [39] |
| U2OS-Cas9 | *H. sapiens* | pLenti-Cas9-2A-Blast | n.a. | This study |
| HeLa | *H. sapiens* | n.a. | n.a. | [39] |
| HeLa-Cas9 | *H. sapiens* | pLenti-Cas9-2A-Blast | n.a. | This study |
| RPE1hTert | *H. sapiens* | n.a. | n.a. | [39] |
| RPE1hTert-Cas9 | *H. sapiens* | pLenti-Cas9-2A-Blast | n.a. | This study |
